# Reliability and accuracy of single-molecule FRET studies for characterization of structural dynamics and distances in proteins

**DOI:** 10.1101/2022.08.03.502619

**Authors:** Ganesh Agam, Christian Gebhardt, Milana Popara, Rebecca Mächtel, Julian Folz, Benjamin Ambrose, Neharika Chamachi, Sang Yoon Chung, Timothy D. Craggs, Marijn de Boer, Dina Grohmann, Taekjip Ha, Andreas Hartmann, Jelle Hendrix, Verena Hirschfeld, Christian G. Hübner, Thorsten Hugel, Dominik Kammerer, Hyun-Seo Kang, Achillefs N. Kapanidis, Georg Krainer, Kevin Kramm, Edward Lemke, Eitan Lerner, Emmanuel Margeat, Kristen Martens, Jens Michaelis, Jaba Mitra, Gustavo G. Moya Muñoz, Robert Quast, Nicole B. Robb, Michael Sattler, Michael Schlierf, Jonathan Schneider, Tim Schröder, Anna Sefer, Piau Siong Tan, Johann Thurn, Philip Tinnefeld, John van Noort, Shimon Weiss, Nicolas Wendler, Niels Zijlstra, Anders Barth, Claus A. M. Seidel, Don C. Lamb, Thorben Cordes

**Affiliations:** Department of Chemistry, Ludwig-Maximilians-Universitaet Muenchen, Butenandtstr. 5-13, 81377 München, Germany; Physical and Synthetic Biology, Faculty of Biology, Ludwig-Maximilians-Universität München, Groβhadernerstr. 2-4, 82152 Planegg-Martinsried, Germany; Molecular Physical Chemistry, Heinrich-Heine-Universität Düsseldorf, 40225 Düsseldorf, Germany; Department of Chemistry, University of Sheffield, S3 7HF, UK; B CUBE – Center for Molecular Bioengineering, TU Dresden, Tatzberg 41, 01307 Dresden, Germany; Cluster of Excellence Physics of Life, Technische Universität Dresden, 01062 Dresden, Germany; Molecular Microscopy Research Group, Zernike Institute for Advanced Materials, University of Groningen, Nijenborgh 4, 9747 AG Groningen, The Netherlands; Department of Biochemistry, Genetics and Microbiology, Institute of Microbiology, Single-Molecule Biochemistry Lab, University of Regensburg, Regensburg, Germany; Department of Biophysics and Biophysical Chemistry, Johns Hopkins University School of Medicine, and Howard Hughes Medical Institute, Baltimore, MD 21205; Dynamic Bioimaging Lab, Advanced Optical Microscopy Center and Biomedical Research Institute, Hasselt University, Agoralaan C (BIOMED), B-3590 Hasselt, Belgium; Department of Chemistry, KU Leuven, Celestijnenlaan 200F, B-3001 Leuven, Belgium; Institute of Physics, University of Lübeck, Germany; Institute of Physical Chemistry, University of Freiburg, Germany; Signalling Research Centers BIOSS and CIBSS, University of Freiburg, Schänzlestrasse 18, 79104 Freiburg, Germany; Department of Physics, Clarendon Laboratory, University of Oxford, Oxford OX1 3PU, UK, Kavli Institute of Nanoscience Discovery, University of Oxford, Oxford OX1 3QU, UK; Bayerisches NMR Zentrum, Department Chemie, Technische Universität München, School of Natural Sciences, Lichtenbergstr. 4, DE-85747 Garching, Germany; Institute of Structural Biology, Helmholtz Center Munich, Ingolstaedter Landstr. 1, 85764 Neuherberg Biomolecular NMR Spectroscopy; Biocenter, Johannes Gutenberg University Mainz; Mainz, 55128, Germany, Institute of Molecular Biology; Mainz, 55128, Germany; Structural and Computational Biology Unit, European Molecular Biology Laboratory, 69117 Heidelberg, Germany; Department of Biological Chemistry, The Alexander Silberman Institute of Life Sciences, and The Center for Nanoscience and Nanotechnology, Faculty of Mathematics & Science, The Edmond J. Safra Campus, The Hebrew University of Jerusalem, Jerusalem, 91904, Israel; Centre de Biologie Structurale (CBS), Univ. Montpellier, CNRS, INSERM, Montpellier, France; Biological and Soft Matter Physics, Huygens–Kamerlingh Onnes Laboratory, Leiden University, Leiden, The Netherlands; Institute for Biophysics, Ulm University, Albert-Einstein-Allee 11, 89081 Ulm, Germany; Materials Science and Engineering, University of Illinois Urbana-Champaign, Urbana IL 61801, USA; Department of Chemistry and Biochemistry, University of California, Los Angeles, CA 90095, USA; California NanoSystems Institute, University of California, Los Angeles, CA 90095, USA

## Abstract

Single-molecule FRET (smFRET) has become an established tool to study biomolecular structure and dynamics *in vitro* and in live cells. We performed a worldwide blind study involving 19 labs to assess the uncertainty of FRET experiments for proteins with respect to the measured FRET efficiency histograms, determination of distances, and the detection and quantification of structural dynamics. Using two protein systems that undergo distinct conformational changes, we obtained an uncertainty of the FRET efficiency of less than ± 0.06, corresponding to an interdye distance precision of ≤ 0.2 nm and accuracy of ≤ 0.5 nm. We further discuss the limits for detecting distance fluctuations with sensitivity down to ≲ 10% of the Förster distance and provide guidelines on how to detect potential dye perturbations. The ability of smFRET experiments to simultaneously measure distances and avoid averaging of conformational dynamics slower than the fluorescence lifetime is unique for dynamic structural biology.

## Introduction

Single-molecule Förster resonance energy transfer (smFRET) studies have become a mature and widely-used approach that is complementary to classical structural biology techniques^1,2^. SmFRET provides information on the structure and conformational dynamics of biomolecules over a distance range of 3 to 12 nm in space and on a timescale of nanoseconds to seconds^1–8^. It allows for the quantitative assessment of structural dynamics and the heterogeneity of conformational ensembles, which are not easily accessible by x-ray crystallography, cryoelectron microscopy and techniques such as cross-linking mass-spectrometry, that provide structural information of solution structures, but lack temporal information. It can also be used to resolve parts of structures or even full structures of biomolecules in an integrative manner (for examples, see refs 9–15) and has the unique ability to provide correlated information on structure and dynamics^1,2^.

Hellenkamp *et al.* presented a quantitative multi-laboratory smFRET blind study of static double-stranded DNA (dsDNA) oligonucleotide rulers that demonstrated a high reproducibility between 19 different labs with an uncertainty of less than 6 Å for the FRET-derived distances^16^. Although the optimal procedure for determining correction factors involved in converting setup-dependent FRET efficiency values into accurate distances remains a topic of active discussion^1^, the results presented by Hellenkamp *et al.* strongly support the idea that standardized smFRET measurements are a useful addition for integrative modelling of static biomolecular structures^10,17,18^.

Here, we take the next step by assessing whether the established procedures translate to more flexible biomacromolecules such as proteins, which often undergo conformational fluctuations. Compared to dsDNA, proteins are generally more challenging systems to study because the local chemical environments of the tethered dyes can vary significantly, which is further amplified by conformational dynamics. Moreover, protein samples require careful handling and storage, due to sample instability and aggregation, and their sensitivity to the biochemical environment and experimental conditions such as buffer composition, pH, temperature or interaction with surfaces. In a blind study involving 19 labs, we investigated how reliably smFRET efficiency histograms of diffusing proteins can be measured by confocal detection of freely-diffusing molecules, and how well structural dynamics can be detected and quantified. As realistic and challenging test cases, we chose two proteins, the maltose-binding protein (MalE) and the U2 Auxiliary Factor 2 (U2AF2), which display conformational dynamics on different timescales that are modulated by ligand binding. Fluorescently labeled protein samples were prepared by stochastically labeling protein double-cysteine variants at positions that will report on specific intramolecular distances. Two key questions are addressed here: (i) How consistently can smFRET efficiency histograms (and the derived distances) be determined by different labs for protein samples? (ii) How reliably can smFRET measurements detect and quantify structural dynamics in proteins? In this context, we investigated the minimal structural fluctuations detectible by smFRET measurements and discuss how to achieve this sensitivity. Our comparison study confirmed the reproducibility of measuring accurate FRET efficiency histograms and the ability of smFRET to detect and quantify conformational dynamics on the sub-millisecond timescale. We demonstrate reproducible FRET efficiency values with uncertainties of less than ± 0.06 corresponding to a distance precision of ≤ 2 Å and an accuracy ≤ 5 Å in MalE. Moreover, we compare the variability of setup-dependent detection parameters and characterize the calibration uncertainty. To push the detection limits for structural dynamics, we refined established experimental and data analysis procedures for the characterization of dynamics and studied a series of distinct dye pairs to identify and eliminate dye-specific effects. With this refinement, we could detect distance fluctuations on the order of 5 Å in the FRET sensitive range. Our work demonstrates the capability of smFRET experiments to study challenging and realistic protein systems with conformational dynamics on timescales from nanoseconds to seconds, highlighting their importance in the expanding toolbox of dynamic integrative structural biology^17–19^.

## Results

In this study, we chose two prototypic protein systems that exhibit conformational dynamics on different timescales. Our first target was MalE of *E. coli.* It is a periplasmic component of the ATP binding cassette transporter MalFGK_2_-E^20,21^ and has been widely studied and applied in biochemistry and molecular biology^22^. MalE exhibits a type II periplasmic-binding protein fold^23,24^ composed of two rigid domains connected by a flexible two-segment hinge (Fig. 1a). This domain arrangement enables an allosterically-driven motion from an open to closed state upon maltose binding with conformational dynamics on the sub-second timescale (Supplementary Fig. 1). As a second system, we chose the large subunit of the U2 auxiliary factor (U2AF2) of the pre-mRNA splicing machinery (spliceosome)^25^. The two RNA recognition motif domains (RRM1,2) of U2AF2 are connected by a long flexible linker and bind single-stranded Py-tract RNA with an affinity of *K_d_* ~1.3 μM for the U9 RNA used in this study^26^. For U2AF2, the two domains fluctuate between an ensemble of detached conformations and a compact conformation in the apo state^27^, whereas ligand binding stabilizes an open conformation (Fig. 2a)^28^.

**Fig. 1.**
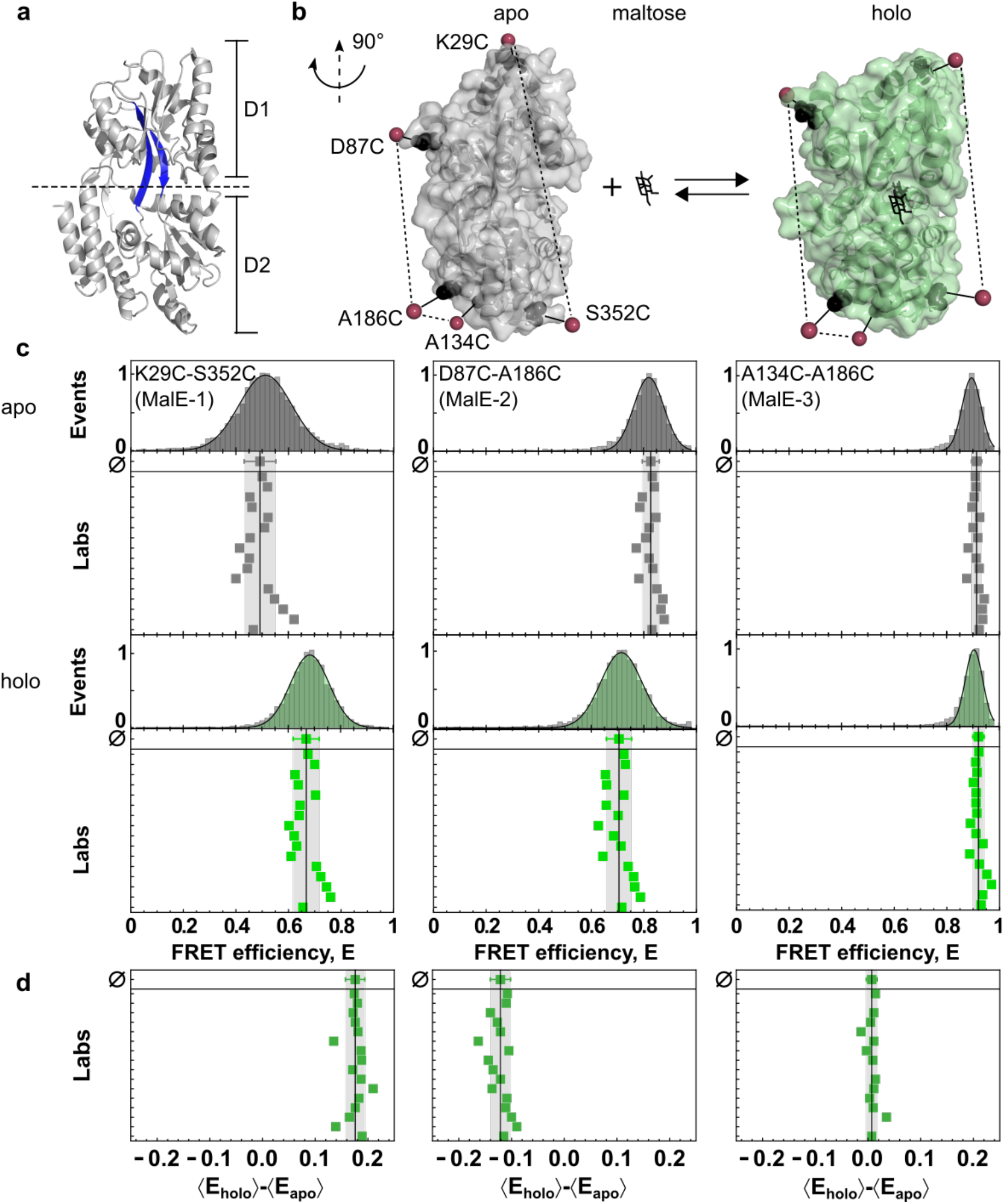
Experimental design of MalE as a protein model system for smFRET studies: **(a)** Crystal structure of MalE in its ligand-free apo state (PDB-ID: 1OMP) with domains D1 and D2 linked by flexible beta-sheets (highlighted in blue). **(b)** The crystal structure of MalE (rotated by 90° as compared to **a**) in the apo (grey, PDB-ID: 1OMP) and holo (green, PDB-ID: 1ANF) states with mutations at K29C / S352C (MalE-1), D87C / A186C (MalE-2), and A134C / A186C (MalE-3) indicated in black. The estimated mean position of the fluorophores from AV calculations are shown as red spheres. **(c)** FRET efficiency *E* histograms for three MalE mutants, MalE-1 (left), MalE-2 (middle), and MalE-3 (right), in the absence and presence of 1 mM maltose (bottom, green) for one exemplary dataset measured in lab 1. The distribution is fitted to a Gaussian distribution. The reported mean FRET efficiencies for 16 labs are shown below (due to experimental difficulties, the results of three labs were excluded; see Supplementary Table 1). The mean FRET efficiency and the standard deviation of all 16 labs are given by the black line and grey area. **(d)** Individual FRET efficiency differences for each lab, between the apo and holo states, 〈*E*_holo_〉 – 〈*E*_apo_〉, for MalE-1 (left), MalE-2 (middle), and MalE-3 (right).

**Fig. 2.**
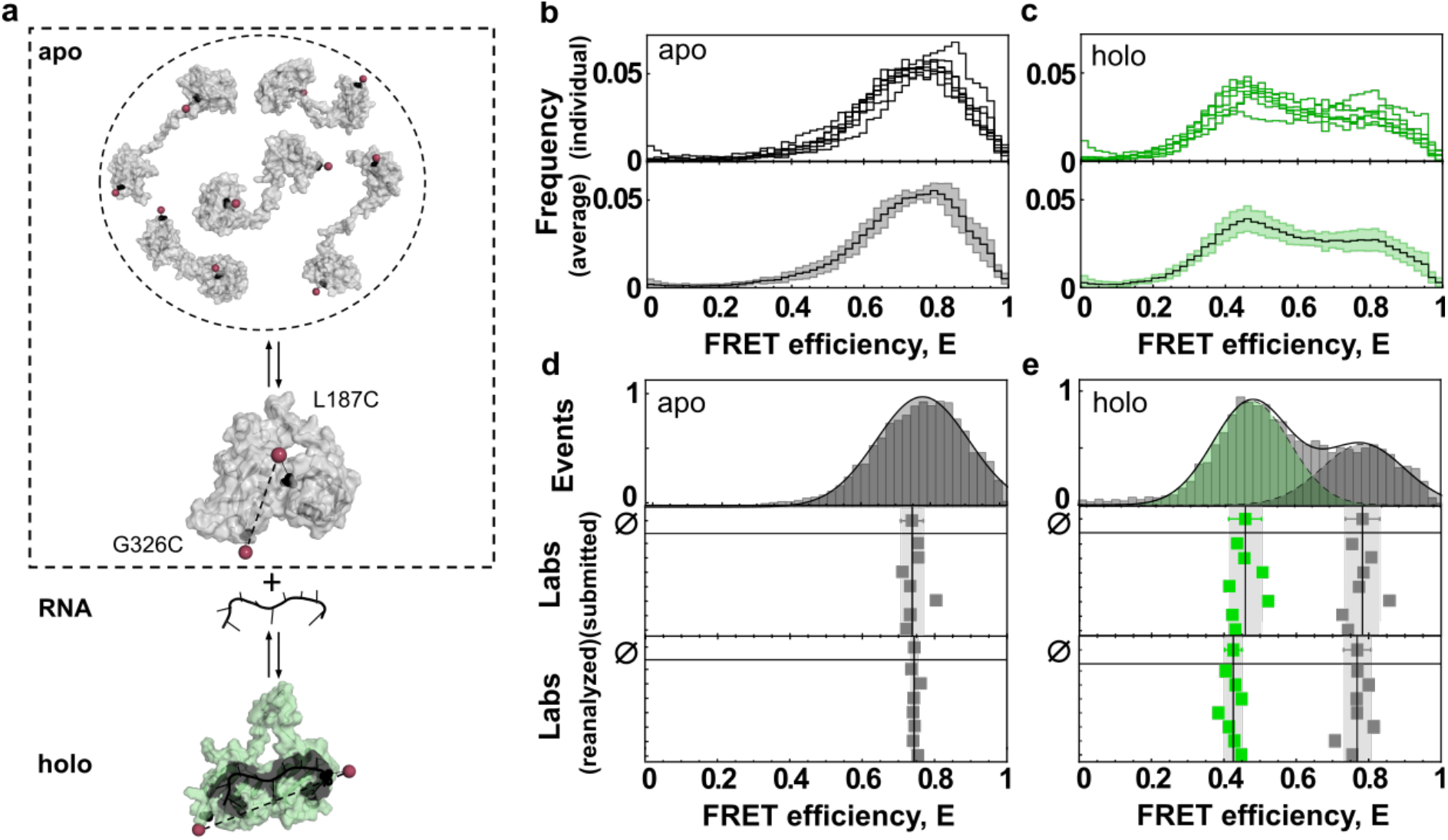
The experimental system of U2AF2 (RRM1,2) and a comparison of FRET efficiency histograms from 7 different laboratories: **(a)** Schematic of the dynamics of U2AF2. U2AF2 is comprised of two tandem RNA-binding motifs, RRM1 and RRM2, which are connected by a flexible linker. The apo state (in grey, top) undergoes fast exchange between an ensemble of detached structures of which 5 representative structures are displayed. A slower exchange occurs between the dynamic detached ensemble and a compact conformation (PDB-ID: 2YHO) shown below. The holo state (in green, PDB-ID: 2YH1), shown with a bound U9 RNA ligand, assumes a well-defined, open conformation. Positions used for introducing cysteine mutations for labeling at L187 in RRM1 and G326 in RRM2 are depicted as black spheres with the mean dye position determined by AV calculations indicated by red spheres. **(b-c)** FRET efficiency histograms reported by participating 7 labs for (**b**) apo and (**c**) holo measurements of U2AF2. Top: Individual FRET efficiency histograms and bottom: the average FRET efficiency histogram from the 7 reporting labs (solid line) with standard deviation (light area). **(d)** FRET efficiency *E* histograms of U2AF2 in the apo state. Top: A representative 1-D FRET efficiency histogram of lab 1 fitted to a Gaussian distribution with a mean FRET efficiency of 0.75. Middle: The reported mean FRET efficiencies reported by 7 labs. The mean value from all data sets is 0.739±0.029, shown above with the corresponding standard deviation in grey. Bottom: The extracted mean FRET values after reanalysis of the collected data. After reanalysis, the agreement improved to 0.742±0.008. **(e)** FRET efficiency histogram comparisons of U2AF2 in the holo state. 5 μM of U9 RNA was used to obtain the holo state FRET histogram for U2AF2. Note the decrease in FRET efficiency after binding of RNA to U2AF2. Top: A representative 1-D FRET efficiency histogram of lab 1 fitted to two Gaussian distributions to determine the FRET efficiencies of the different subpopulations, yielding mean FRET efficiencies of 0.44 for RNA-bound and 0.76 for the RNA-free conformation. Middle: The mean FRET efficiencies reported by the 7 labs. The mean values from all 7 the data sets were 0.45±0.04 for the RNA-bound conformation (in green) and 0.78±0.04 for the RNA-free conformation (in grey). Bottom: Reanalysis of the holo measurements yielding values of 0.42 ± 0.02 and 0.77 ± 0.03 for RNA-bound and RNA-free fractions respectively.

SmFRET experiments were blindly performed by 19 laboratories for MalE and by seven laboratories for U2AF2 using different implementations of solution-based confocal spectroscopy with alternating excitation, μs-ALEX^29^ for intensity-based analysis and ns-ALEX^30^ or PIE^31^ for intensity- and lifetime-based analyses (Supplementary Fig. 2). We adapted a data analysis routine similar to that of Hellenkamp *et al.^16^* to determine setupindependent accurate FRET efficiency *E* values from the photon counts detected in the donor (D) and acceptor (A) detection channels during a single-molecule event. The implementation of ALEX or PIE^2,6,7,16,32,33^ (see Supplementary Note 1 for a detailed comparison of ALEX and PIE) was crucial for: (i) careful corrections of the registered photon counts to reflect the actual donor and acceptor signal; and (ii) exclusion of single-molecule events from further analysis that originate from incompletely labeled molecules, or showed photo-blinking or bleaching. The correction procedures for reporting accurate FRET efficiencies are described in the Online Methods and include subtraction of background signal from all channels and the determination of four correction factors: (α) for spectral crosstalk of D fluorescence into the A channel, (β) for normalization of direct D and A excitation fluxes, (*γ*) for differences in donor and acceptor quantum yields and detection efficiencies, and (δ) for the ratio of indirect and direct A excitation (Supplementary Fig. 3 and Supplementary Tables 1 and 2)^32^.

### MalE

For smFRET investigations of MalE, we prepared three different cysteine variants that cover a large part of the dynamic range of FRET and monitor the conformational change in the protein upon maltose binding (Fig. 1b, see Online methods and Supplementary Fig. 4). The variants were designed such that MalE-1 (K29C-S352C) shows a decrease in the inter-dye distance upon maltose binding, MalE-2 (D87C-A186C) shows an increase in distance and MalE-3 (A134C-A186C) shows no distance change upon substrate binding. All variants of MalE were stochastically labeled at the given positions with Alexa Fluor 546 (Alexa546) as the donor and Alexa Fluor 647 (Alexa647) as the acceptor fluorophore. We confirmed the functionality of the labeled protein by ligand titrations using smFRET and microscale thermophoresis and ensured that the ligand maltose does not affect the photophysical properties of the dyes (Supplementary Fig. 5 and 6). For the sake of comparison, participants were asked to provide the mean FRET efficiencies using the fit to a Gaussian distribution for estimating the peak of the apo and holo FRET efficiency histograms (as shown in Fig. 1c). For this study, we asked 19 laboratories to determine a common (global) *γ*-value using all three MalE for both the apo and holo measurement conditions (Supplementary Note 2 and Supplementary Fig. 3). To execute this workflow, participants used many different in-house or publicly available software packages following the given guidelines.

FRET efficiency histograms for representative experiments of the MalE variants in the apo state and in the presence of 1 mM maltose are shown in Fig. 1c together with values reported by 16 labs, showing very good agreement and reproducibility. All labs observed the expected maltose-induced conformational change for MalE-1 and MalE-2, and no significant change for MalE-3. This indicates that the samples did not degrade during shipment on dry ice and storage in the labs at 4 °C. MalE-1 showed an average FRET efficiency of 0.49±0.06 in the apo- and 0.67±0.05 in the holo state due to the hinge motion of the protein upon ligand binding. MalE-2 showed the expected decrease in FRET efficiency from 0.83±0.03 to 0.71±0.05 in the apo and holo states, respectively (Fig. 1c). MalE-3, with both labels on one lobe, showed no significant change in FRET efficiency (*E*_apo_ = 0.91±0.02, *E*_holo_ = 0.92±0.02).

The standard deviation of the determined mean FRET efficiency over all labs was less than ±0.06, similar to the precision found for dsDNA (Table 1 and Supplementary Table 3)^16^. We observe the highest standard deviation for MalE-1 and the lowest values of ±0.02 for MalE-3, which also has the highest FRET efficiency. As will be discussed in detail below, the observed spread of the reported FRET efficiencies depends less on the measurement statistics, but on the uncertainty in the calibration factors. This effect is largest at intermediate FRET efficiencies, which explains the higher spread of values for the MalE-1 mutant. Interestingly, for most labs, we observed systematic deviations of the reported FRET efficiency values for the apo and holo states from the mean value. This suggests that changes of the FRET efficiency are measured even more accurately than absolute values. In Fig. 1d, we analyze the individual FRET efficiency differences, 〈*E*_holo_〉 – 〈*E*_apo_〉, between the apo and holo states for the different labs. Here, the distribution of values indeed narrows approximately by a factor of two for all three mutants because systematic deviations cancel out (standard deviations σ〈*E*_holo_〉–〈*E*_apo_〉 for MalE-1: ±0.02, MalE-2: ±0.02, MalE-3: ±0.01, Fig. 1d, Table 1 and Supplementary Table 3).

**Table 1.** Average of mean FRET efficiency and standard deviation for MalE and U2AF2 samples reported by the participating laboratories. The calculated average *μ*_〈*E*〉_ and standard deviation *σ*_〈*E*〉_ of the mean FRET efficiency values provided by the participating labs are given for all three studied mutants of MalE labeled with Alexa546 and Alexa647 under both apo and holo conditions (see Supplementary Table 3). The calculated mean and standard deviation of the difference in the reported mean FRET efficiency between the apo and holo (〈*E*_holo_〉 – 〈*E*_apo_〉) for the three MalE mutants are given by *μ*_〈*E*_holo_〉–〈*E*_apo_〉_ and *σ*_〈*E*_holo_〉–〈*E*_apo_〉_ respectively (see Supplementary Table 3). The calculated average *μ*_*R*_〈*E*〉__ and standard deviation *σ*_*R*_〈*E*〉__ of the mean distances were derived according to Eq. 2. The modeled distances 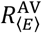 and 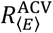 are derived using accessible volume (AV) and accessible contact volume (ACV) calculations respectively, as described in the Online Methods. We also give the average and standard deviation for the FRET values determined for U2AF2 labeled with Atto532-Atto643 under both apo and holo conditions (Supplementary Table 4). *Only studied by two labs. **Due to the fast-structural dynamics in the sample, only 7 labs studied this mutant and distances were not determined. *** Only the holo state under holo condition was considered.

| Sample | Condition | Experimental Values |  |  | Modeled Distances |  |
| --- | --- | --- | --- | --- | --- | --- |
| | | $\mu_{\langle E \rangle} \pm \sigma_{\langle E \rangle}$ | $\mu_{\langle E_{\text{holo}} \rangle - \langle E_{\text{apo}} \rangle} \pm \sigma_{\langle E_{\text{holo}} \rangle - \langle E_{\text{apo}} \rangle}$ | $\mu_{R_{\langle E \rangle}} \pm \sigma_{R_{\langle E \rangle}} [\text{\AA}]$ | $R_{\langle E \rangle}^{\text{AV}} [\text{\AA}]$ | $R_{\langle E \rangle}^{\text{ACV}} [\text{\AA}]$ |
| MalE-1 | apo | 0.49±0.06 | 0.177±0.019 | 65.4±2.6 | 72.0 | 67.7 |
|  | holo | 0.67±0.05 |  | 57.8±2.1 | 62.1 | 58.3 |
| MalE-2 | apo | 0.83±0.03 | -0.121±0.019 | 50.0±1.9 | 50.1 | 48.8 |
|  | holo | 0.71±0.05 |  | 56.1±2.1 | 56.5 | 55.0 |
| MalE-3 | apo | 0.913±0.019 | 0.007±0.010 | 43.8±1.7 | 39.9 | 38.9 |
|  | holo | 0.920±0.021 |  | 43.0±2.4 | 40.8 | 39.8 |
| MalE-4* | apo | 0.442±0.025 |  | 67.6±1.2 | 67.8 | 64.3 |
|  | holo | 0.678±0.017 |  | 57.4±0.7 | 56.9 | 54.6 |
| MalE-5* | apo | 0.613±0.003 |  | 60.22±0.15 | 61.8 | 59.3 |
|  | holo | 0.821±0.001 |  | 50.43±0.08 | 49.3 | 48.2 |
| U2AF2** | apo | 0.74±0.03 |  | 49.6±1.3 |  |  |
|  | holo*** | 0.46±0.04 |  | 60.8±1.7 |  |  |

### U2AF2

For the second protein, U2AF2, we chose the previously published double cysteine variant (L187C / G326C) of the minimal RRM1,2 construct, where we verified that protein properties of the used variant are not affected by labeling (Fig. 2a)^34,35^. The construct contains one cysteine on each RRM domain, which were labeled stochastically with the dye pair Atto532-Atto643. A subset of seven groups measured the second protein. To investigate the consistency of the shape of the obtained FRET efficiency histograms, we plot in Fig. 2b/c the smFRET histograms from the individual laboratories (row 1) as well as the average FRET efficiency distribution illustrated by the mean and standard deviation (row 2). All groups found a single broad distribution (Fig. 2b, row 1) in the apo state with an average FRET efficiency of *E*= 0.74±0.03 (row 2). In the presence of 5 μM ligand, a second narrower peak at lower *E* appears (Fig. 2c, row 1) with an average FRET efficiency of *E* = 0.46±0.04 (row 2) as expected for the open configuration of the holo state^28,34^. Notably, a significant fraction (~ 15%) of ligand-free protein remains in the sample at the RNA concentration used (Supplementary Fig. 7).

For the apo state, we obtained a similar standard deviation of ±0.03 as found for Mal-E, however a clear outlier was apparent (Supplementary Table 4). To test whether user bias affected the reported results, we had the datasets reanalyzed by a single person. While analyzing the different data sets, this person could determine an optimal procedure for determining the correction factors for this challenging sample (Supplementary Note 3). Hereby, the person could improve the agreement to a standard deviation of ±0.008 with no change in the average FRET efficiency value (Fig. 2d/e, Supplementary Table 4). The reanalysis revealed that the detection correction factor γ was the main cause of the deviations between the measurements as the single population of the apo state did not allow for a robust determination of the γ-factor^32,33^. In this case, it was best to estimate the γ-factor from a global analysis of the apo and holo measurements, which was possible due to the absence of any detected changes in the quantum yield of the fluorophores upon binding of the RNA (as measured using PIE) (Supplementary Table 5). We also reanalyzed data from the same seven laboratories for the MalE-1 apo measurements. Nearly identical mean FRET efficiencies and standard deviations were determined upon reanalysis (0.49 ± 0.05 versus 0.47 ± 0.06, Supplementary Fig. 8) indicating that user bias was less significant when a global, well-defined analysis procedure was provided over several samples covering a significant fraction of the FRET range (Supplementary Note 2).

For the holo state of U2AF2, good agreement was obtained for the peak positions with a standard deviation of ±0.03 and ±0.02 for the high-FRET and low-FRET peaks, respectively, and only a minimal improvement resulted from the reanalysis (Supplementary Table 4). In this case, the two populations allowed for a more robust determination of the γ-factor, which can be performed by analyzing the FRET efficiency versus stoichiometry, *S*. In contrast to the good agreement in FRET efficiency, we observed larger variations in the relative amplitudes of the two populations: 0.58±0.08 for the holo state and 0.42±0.08 for the apo population (Fig. 2c, Supplementary Table 4). Such differences are not unexpected due to potentially reduced protein activity, degradation of the ligand, and the high sensitivity of biomolecular dynamics to the experimental conditions, e.g., temperature, ligand concentration, buffer composition, salt concentration or the presence of stabilizers such as BSA (Supplementary Fig. 7).

### Characterizing setup-dependent parameters and correction factors

The quality of smFRET experiments is determined by the statistics of the measurement and the performance of the setup to maximize photon collection and thereby minimize shot noise. To this end, we quantified the number of bursts, average count rate, burst duration and number of photons in the donor and FRET channels for the reported MalE measurements from eight labs (Fig. 3a, Supplementary Fig. 9). On average, participants collected 6000 bursts (min: 500, max: 21,000) of molecules carrying both the donor and acceptor fluorophore. The required number of bursts for a smFRET analysis depends on the goal of the experiment. For a simple estimation of an average FRET efficiency from a single population, as performed for MalE, a low number of double-labeled bursts of ~1000 may be sufficient. However, if advanced analysis methods such as time-correlated single photon counting (TCSPC) detection for lifetime analysis, burst-wise fluorescence correlation spectroscopy (FCS) or a photon distribution analysis (PDA) are to be applied to sub-ensembles, higher burst numbers of (>5000) are desired for a robust analysis. Typical count rates per single-molecule event were found to be 60±20 kHz, and an average of 90±40 photons were detected over a typical burst duration of 1.7±0.9 ms (Fig. 3a, Supplementary Fig. 9). The average count rate and burst duration depend on the size of the confocal volume, where smaller sizes typically result in higher count rates but shorter burst durations. Indeed, for the collected data, we observe a negative correlation between the burst duration and the average count rate (Fig. 3b, Pearson’s r = −0.58, Supplementary Fig. 10). The large spread of the burst duration arises from the fact that some participants applied a diffraction limited observation volume while others intentionally underfilled the objective lens to create a larger confocal volume with a diameter of ~1 μm (assuming that the labs have adjusted their detection pinhole to correspond with the enlarged excitation volume). We also observed a small positive correlation between the number of detected photons and the burst duration (Fig. 3c, Pearson’s r = 0.54, Supplementary Fig. 10). These results indicate that larger confocal volumes, in combination with high irradiances, yield the highest number of photons per burst^36^. Smaller observation volumes generally yield higher count rates and thus shorter inter-photon times, enabling fast transitions on the sub-μs timescale to be resolved^37,38^. Longer burst durations offer the benefit that slower dynamics can be studied.

**Fig. 3.**
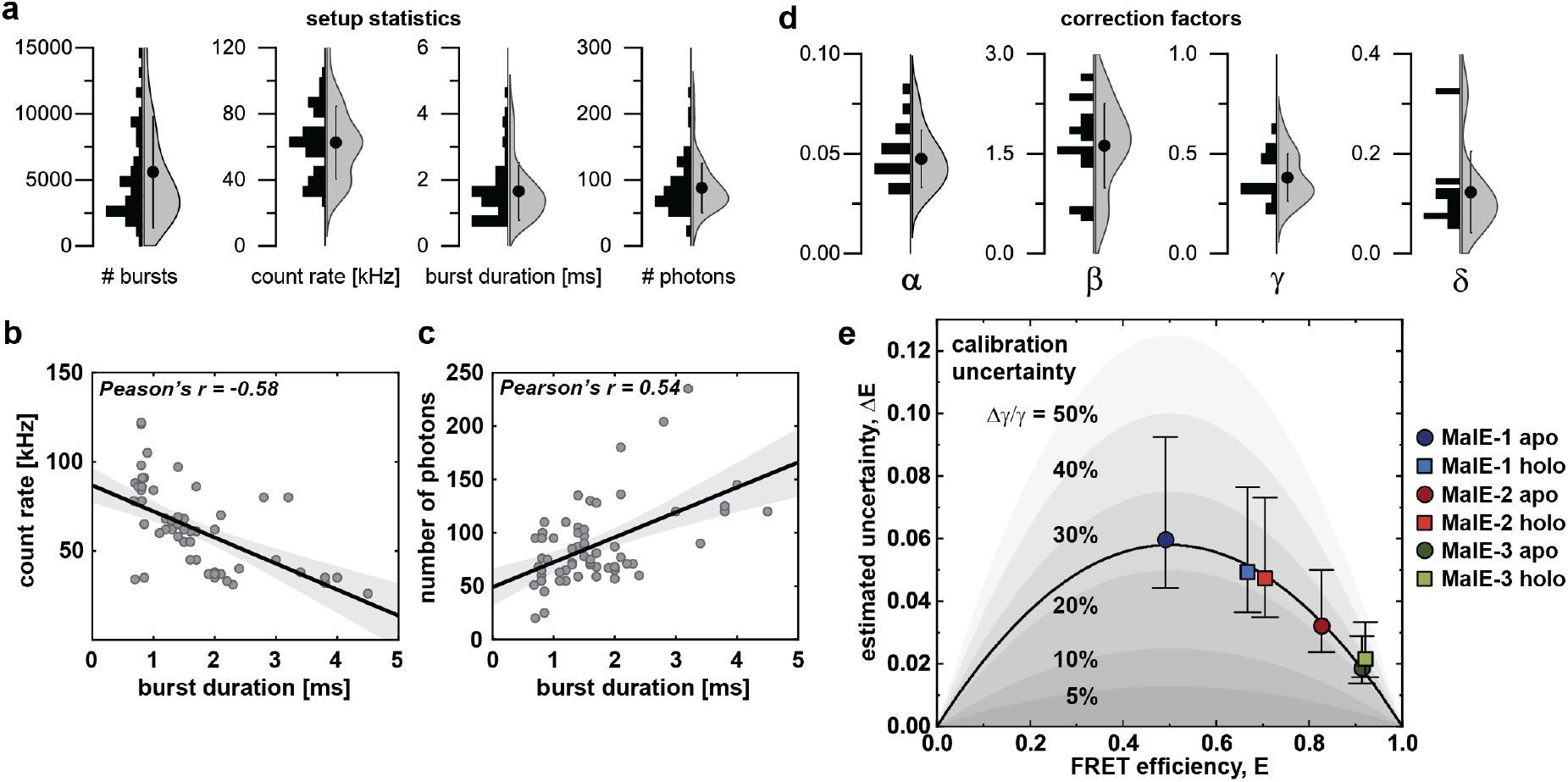
Setup-dependent parameters and calibration uncertainty. **(a)** The distribution of the parameters quantifying the statistics of the measurements and the performance of the setups used for both MalE and U2AF2 measurements are shown as histograms and violin plots for the measurements from 8 labs. The circle and whiskers in the violin plot indicate the mean and standard deviation. Sample-dependent distributions of the shown parameters are given in Supplementary Fig. 9. **(b-c)** Pairwise plots of the average count rate **(b)** and number of photons **(c)** against the burst duration. The same datasets are plotted as used for panel **(a).** While the count rate decreases slightly for longer burst durations, a positive correlation is observed for the acquired number of photons per burst and the burst duration, indicating that larger observation volumes result in a higher accumulated signal per molecule. Correlations between all parameters are shown in Supplementary Fig. 10. **(d)** The distributions of the correction factors for the calculation of accurate FRET efficiencies for all the MalE measurements are shown as histograms and violin plots for the measurements from all labs. **(e)** A plot of the standard deviation of the reported FRET efficiencies (as a measure of the experimental uncertainty) against the average FRET efficiency for the MalE mutants 1-3 reveals that lower uncertainties are observed for higher FRET efficiencies. The black line represents a fit of the estimated uncertainties under the assumption that the variations arise solely due to an uncertainty in the γ-factor (see Eq. 1). The inferred relative uncertainty of the γ-factor is ~23%. Shaded areas indicate relative uncertainties of 5-50%. Error bars indicate 95% confidence intervals.

For an accurate analysis of the data, the correction factors for spectral crosstalk (α), excitation flux (β), detection efficiency (*γ*) and direct excitation (δ) must be determined. Based on data from 16 labs, we plot the distribution of the correction factors used to determine accurate FRET efficiencies for the MalE system in Fig. 3d (Supplementary Table 1). Besides fluorophore properties, the correction factors also depend on setup-specific parameters such as the dichroic mirrors, the emission filters, the detectors, and the excitation wavelengths and power. Nonetheless, we observed a defined distribution for the crosstalk correction factors *α* of 0.05±0.01, which is mainly determined by the emission filters and type of detectors used for the donor and acceptor detection channels. A larger spread was observed for the correction factor for the excitation flux β of 1.6±0.6 and direct excitation *δ* of 0.12±0.08. Both factors depend on the ratio of the excitation powers for the donor and acceptor fluorophores. Most participants used about half the laser power for direct acceptor excitation (45±27 μW) as they used for excitation of the donor fluorophore (78±58 μW) to achieve similar count rates after donor and acceptor excitation. The agreement between the reported FRET efficiency values clearly shows that the various experimental settings are compensated for by the self-consistent correction procedure applied here.

For the detection efficiency correction factor *γ*, we observed an average of 0.4±0.1. The *γ*-factor is arguably the most difficult to determine (see Supplementary Fig. 3b). It depends on the acceptor to donor ratio of the detection efficiencies, *g*, and the effective fluorescence quantum yields 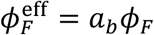, where *a_b_* represents the fractions of molecules in the bright state, as 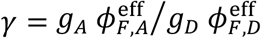^16^. Like the crosstalk correction term, the detection efficiencies strongly depend on the emission filters and the type of detectors used. Due to the relatively low fluorescence quantum yield of the acceptor, *ϕ_F,A_* ~ 0.32, compared to that of the donor, *ϕ_F,D_* ~ 0.72, all labs reported *γ*-factors below 1. Despite the large spread in the different correction factors, we observed very good agreement for the reported FRET efficiencies in our blind study. However, *γ* is also the key factor that limits the consistency between laboratories. This notion is supported by two observations: (i) In Fig. 1d, the spread of FRET efficiency differences, 〈*E*_holo_〉 – 〈*E*_apo_〉, is smaller (e.g., 0.06 to 0.02 for MalE-1) than for the absolute *E* values in Fig. 1c, suggesting that errors in *E* are systematic rather than random. (ii) The observed spread in reported FRET efficiencies depends on the absolute FRET efficiency measured for MalE (Fig 1c,d). We also calculated the uncertainty due to all parameters in the FRET efficiency calculation using error propagation for cross-talk, direct acceptor excitation, background correction in the donor and acceptor channels. The reported uncertainty can be attributed mainly to the uncertainty in the *γ*-factor (Fig. 3e, Supplementary Note 4). The error of the *γ*-factor, Δ*γ*, propagates into an uncertainty in the reported FRET efficiencies, Δ*E*, as follows:

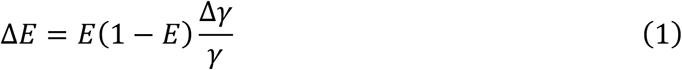

Notably, the observed Δ*E* is well described by Eq. 1 (black line in Fig. 3e), yielding a relative uncertainty of Δγ/γ = 23% (corresponding to an approximate absolute error of Δγ = 0.07). The improved agreement between the measurements upon reanalysis by a single person for U2AF2 (Fig. 2d) indicates that the accuracy of the analysis could be further improved by establishing robust and standardized procedures for the determination of all experimental correction factors, which differ depending on how many populations are present in the measurement and whether the FRET efficiency peak is dynamically averaged (Supplementary Note 2).

### Detection and quantification of conformational dynamics in proteins via smFRET

For immobilized molecules, the analysis of dwell-times from the fluorescence trajectories provides access to kinetics on the millisecond to second time-scales (Supplementary Fig. 1)^39–41^. When performing smFRET experiments using confocal detection of freely diffusing molecules, millisecond dynamics can also be measured from a direct analysis of the intensity trajectories (for slowly diffusing molecules)^42,43^. A number of additional approaches can be used for detecting and quantifying faster sub-millisecond conformational dynamics (with the maximum timescale limited by the burst duration) such as FRET-FCS^42,44,45^, filtered-FCS^46,47^, burst-variance analysis (BVA)^48^, FRET-2CDE^49^, dynamic PDA^50^, FRET efficiency *E* versus fluorescence-weighted average donor lifetime 〈*τ*_*D*(*A*)_〉_*F*_ analysis (*E*-*τ* plots)^50,51^, nanosecond-FCS^52^, recurrence analysis of single particles^53^, photon-by-photon maximum likelihood approaches^38,54–57^ and Monte Carlo diffusion-enhanced photon inference (MC-DEPI)^58^. To assess how consistently dynamics can be detected, we asked the various groups in this blind study to evaluate whether the protein systems they studied were static or dynamic on the millisecond timescale and which method they used to come to this conclusion (Supplementary Table 6).

The most frequently used methods to evaluate dynamics were the BVA and *E*-*τ* plots. Both techniques visualize FRET dynamics by comparing the measured data to theoretical expectations for static systems (Fig. 4a/b). BVA detects dynamics by estimating the standard deviation of the FRET efficiency over the time course of the individual bursts, using a predefined photon window (typically around ≳ 100 μs depending on the molecular brightness). Due to FRET dynamics, the standard deviation of the FRET signal within a burst (red line in Fig. 4a) is higher than expected from shot noise (black line in Fig. 4a), which becomes visible as a deviation (apparent dynamic shift, ds) from the shot-noise limited standard deviation of the apparent FRET efficiency (which is a semi-circle in shape)^48^.

**Fig. 4.**
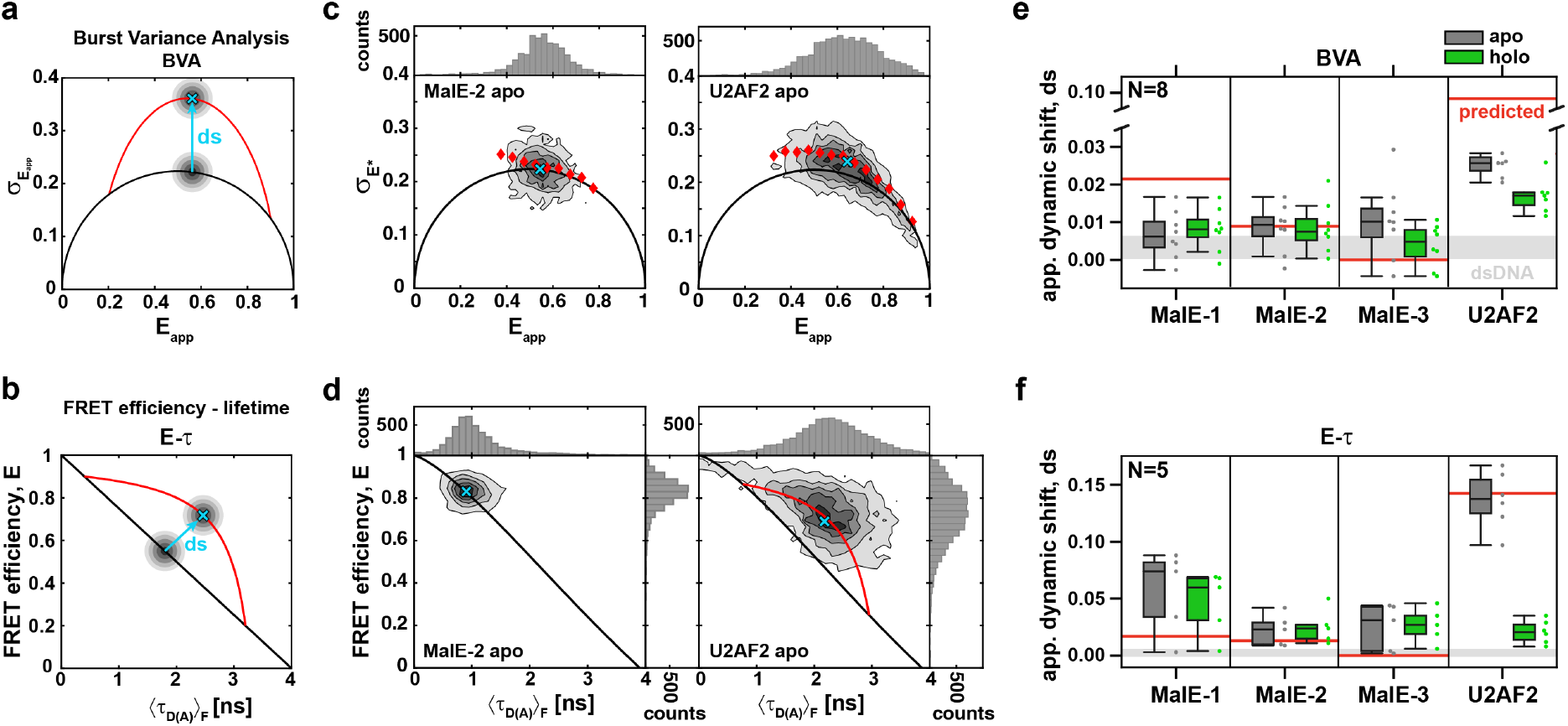
Detection and characterization of conformational dynamics on the sub-millisecond timescale in MalE and U2AF2: **(a-b)** Schematic representations of burst variance analysis (BVA) and *E*-*τ* plot. (a) In BVA, the standard deviation σ_*E*_app__ of the apparent FRET efficiency *E*_app_ is compared to the shot-noise limit. Singlemolecule events with conformational dynamics show increased variance and follow the dynamic line (red). The dynamic shift ds is defined as the excess standard deviation compared to the static line. (b) In the *E*-*τ* plot, the intensity-based FRET efficiency *E* is plotted against the intensity-weighted average donor fluorescence lifetime, 〈*τ*_(*D*(*A*)_〉_*F*_. Molecules undergoing dynamics are shifted from the static line (black) and follow a dynamic FRET-line (red, see text)^59^. For a given population, the dynamic shift is defined as the displacement of the population orthogonal to the static FRET-line. See Supplementary Note 5 for details. **(c)**BVA of MalE-2 labeled with Alexa546-Alexa647 without maltose (apo, left) and U2AF2 labeled with Atto532-Atto643 without RNA (apo, right). Here, the BVA is based on a photon binning of 5 photons. Red diamonds indicate the average standard deviation of all bursts within a FRET efficiency range of 0.05. The mean positions of the populations (cyan crosses) were determined by fitting a two-dimensional Gaussian distribution to the data (Supplementary Note 5). **(d)**The plots of the FRET efficiency *E* versus intensity-weighted average donor lifetime 〈*τ*_*D*(*A*)_〉_*F*_ of the same measurement as in **(c)**. The static lines are slightly curved as they account for the flexibility of the dye linkers^50^. The donor-only population was excluded from the plot. For MalE-2, the population falls on the static FRET-line, while a clear dynamic shift is observed for U2AF2. The end points of the dynamic FRET-line for U2AF2 were determined from a sub-ensemble analysis of the fluorescence lifetime decay. **(e-f)** The dynamic shift of the peak of the population was determined graphically from BVA (8 labs for MalE, and 7 labs for U2AF2 respectively) and *E*-*τ* (5 labs) plots (see Online Methods). For U2AF2 in the holo state, the dynamic shift was assessed only for the low-FRET RNA-bound population. All labs consistently detected the highest dynamic shift for U2AF2 in the apo state. A significant dynamic shift was also consistently detected for MalE-1. Boxes indicate the median and 25%/75% quartiles of the data. Whiskers extend to the lowest or highest data point within 1.5-times the interquartile range. The grey area indicates the dynamic shift obtained for the double-stranded DNA used in a previous benchmark study^16^ based on measurements performed in lab 1 for BVA (ds_DNA_ = 0.0033 ± 0.0033) and lab 2 for the *E*-*τ* plot (ds_DNA_ =0.0026 ± 0.0044). The horizontal red lines indicate the expected dynamic shift for a potential conformational exchange between apo and holo states. We computed the expected change of FRET efficiency using their structural models in the PDB (Supplementary Note 6 and Supplementary Table 9).

In the *E*-*τ* plots, the observed FRET efficiency determined via intensity (the *y*-axis in Fig. 4b) is a species-weighted average and, in the presence of dynamics, the position along this axis depends on the fraction of time spent in the respective states. The fluorescence lifetime of the donor (〈*τ*_*D*(*A*)_〉_*F*_, the *x*-axis in Fig. 4b) is a photon-weighted average because only a single lifetime can be determined from the single-molecule lifetime data. It is weighted towards the lifetime of the lower FRET state as the majority of photons are emitted from the donor in the low FRET efficiency state^50,51^, shifting the data to the right of the ‘static’ FRET-line. *E*-*τ* plots can detect dynamics on the ns to ms timescale. Note that, for the experimental data, we have included an additional correction to the ‘static’ FRET-line that accounts for the distance fluctuations due to the flexible dye linkers of 6 Å, resulting in a slightly curved line^59^. To quantify the dynamics between two distinct states, a theoretical ‘dynamic’ FRET-line (red line in Fig. 4b) can be calculated and overlaid on the plots. Again, the apparent dynamic shift, ds, is defined as the deviation of the observed data from the theoretical static line (Fig. 4b). See Supplementary Note 6 for details.

We previously showed that MalE exhibits slow ligand-driven dynamics on the sub-second timescale between high- and low-FRET states (Supplementary Fig. 1)^60^. Here, we investigated whether the apo and holo states of MalE are undergoing dynamics faster than or on the timescale of the diffusion time of 1-3 ms. Both techniques reveal that the conformations of MalE exhibit no large FRET-fluctuations on the ms timescale (Fig. 4c,d; Supplementary Fig. 11). Almost all groups confirmed this assessment for all MalE samples (Supplementary Table 6). Three groups concluded that MalE is dynamic without further justification. To investigate potential dynamics in more detail, we determined the dynamic shifts for a subset of the data (eight labs for BVA, Fig. 4e and five for *E*-*τ*, Fig. 4f, Supplementary Note 5, Supplementary Table 7). As a static control, we determined the dynamic shift of the dsDNA rulers used in Hellenkamp *et al.^16^* (mean ± one standard deviation as determined from Labs 1 and 2) shown in grey in Fig. 4e,f (Supplementary Table 8). Interestingly, no apparent dynamic shift exceeding the dsDNA reference was observed when using BVA for all MalE mutants. From the *E*-*τ* plots, however, there is an apparent dynamic shift, especially for MalE-1 of ~0.05, that clearly exceeds what would be expected for a static system or even what is predicted for potential dynamics between the apo and holo conformation (Fig. 4f, red lines, Supplementary Note 6). Hence, some labs categorized MalE as dynamic. The origin of this apparent shift, which must originate from dynamics that are faster than ~100 μs, will be discussed in detail below.

In contrast to MalE, all groups found U2AF2 to be dynamic as was expected for two domains connected by a flexible linker (Fig. 4c-f, Supplementary Table 6). The ligand-free apo state shows pronounced deviations from the behavior for static molecules both in the BVA and *E*-*τ* plots, while the RNA-bound holo state shows a significant apparent dynamic shift for BVA but not for the *E*-*τ* analysis (Fig. 4c-f). Due to the existence of a significant fraction of apo-protein and the overlap between the apo and holo populations, it was challenging to assess whether the holo state is truly static or dynamic, although a clear apparent shift is observed. In summary, even though U2AF2 is a very challenging test case, dynamics were unambiguously detected in all labs demonstrating the reliability of smFRET for investigating dynamic systems.

### Accuracy of FRET-derived distances in proteins with respect to structural models

After determining FRET efficiencies of different conformations in a protein, the next step is (often) to convert these FRET efficiencies into distances and compare them to what is expected from structures or to use them as distance constraints in integrative FRET-assisted structural modeling^1,3,5,13,61^. The smFRET experiments yield the FRET efficiency as a result of dynamically, non-linearly averaged distances due to the flexible linker used to attached the fluorophore to the molecule. Fast and robust ways of accurately modeling the fluorophore positions and volumes accessible to the fluorophore attached to the biomacromolecule is a topic of ongoing investigation^3,61,62^. To assess the accuracy of our measurements, we measured the fluorescence lifetimes and time-resolved anisotropies for each labeling position and applied the accessible volumes (AV) approach^3,4,63^ that employs a coarse-grained dye model to estimate the FRET efficiency averaged model distance 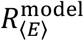 between the two dyes. For this, all possible positions of the donor and acceptor fluorophores are averaged, taking into account distinct linker conformations and steric hindrances of the protein (Fig. 5a-c; see Online Methods and reference 4 for details).

**Fig. 5.**
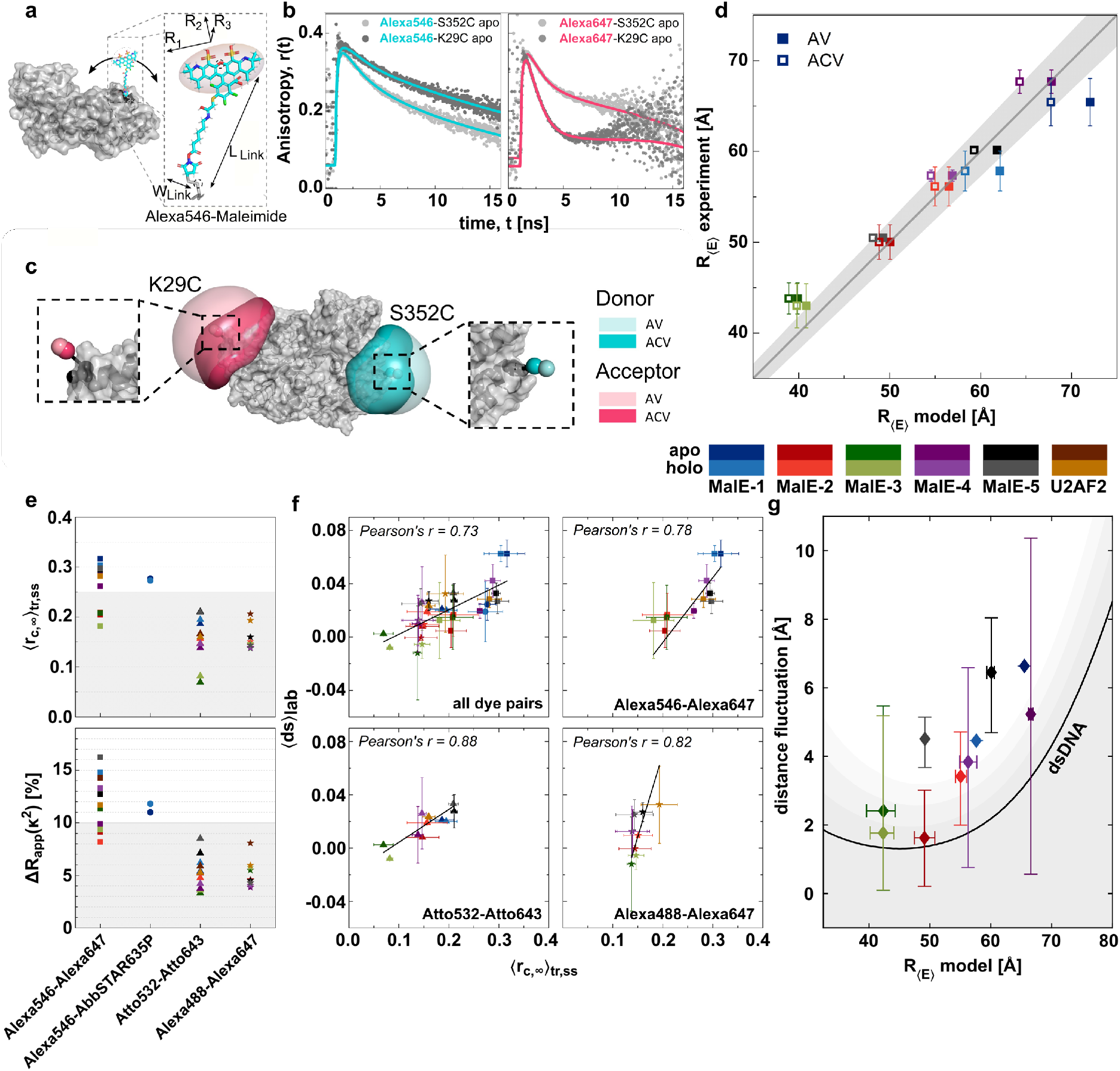
Assessing the accuracy of FRET-derived distances in MalE. **(a-d)** Accessible volume calculations and model-based interdye distances: **(a)** Scheme of the Alexa546 fluorophore attached to MalE (PDB 1OMP) showing the parameters used for the accessible volume calculations. The fluorophore is modeled as an ellipsoid with a flexible linker using the AV3 model^4^ (Supplementary Table 10). **(b)** Fluorescence anisotropy decays of the donor fluorophore (Alexa546, left) and acceptor fluorophore (Alexa647, right) at the two labeling positions K29C and S352C on mutant MalE-1. The anisotropy decays were obtained from single-cysteine mutants labeled only with the donor or acceptor fluorophore respectively. Strong hindrance of the rotation due to sticking to the protein surface, as indicated from the high vertical offset and slow decay of the anisotropy, are detected for the Alexa546 at both positions and Alexa647 at position S352C. Solid lines represent fits to a model of two or three rotational components (Supplementary Tables 11 and 12). The analysis was performed as described in Supplementary Note 8. **(c)** Accessible volumes for Alexa546 (cyan) and Alexa647 (pink) at labeling positions 352 and 29 calculated using the AV model as described in panel (a). The contact volume close to the biomolecular surface (shown as a darker shade) is used in the ACV model by weighting the occupancy of the contact volume based on the residual anisotropy. The zoom-ins show the mean positions of the dyes based on the AV (light shade) and ACV (darker shade) model. In the ACV model, the position of the dyes is biased towards the protein surface, resulting in a reduction of the interdye distance for the given labeling positions. See Online Methods for details. **(d)** Comparison of the experimentally obtained FRET-averaged distance *R*_〈*E*〉_, with the theoretical model distances using the AV (filled squares) and ACV (empty squares) calculations. Errors of the experimental distances represent the standard deviation over all labs. The solid line represents a 1:1 relation and the grey area indicates an uncertainty of ± 3 Å for a Förster radius of R_0_ = 6.5 nm. The two additional mutants, MalE-4 and 5, labeled at positions K34C-N205C and T36C-N205C, were measured by two labs. The agreement between the model and experiment (determined using the average root-mean-square deviation) decreases from 3 Å for the AV model to 2 Å for the ACV model. **(e)** Detection of dye-specific protein interactions. (Top) The five MalE mutants and U2AF2 were labeled with four different dye combinations (Alexa546-Alexa647, Alexa546-AbbSTAR635P, Atto532-Atto643 and Alexa488-Alexa647) and measured by three different labs to determine the donor-acceptor-combined residual anisotropy from time-resolved (tr) and steady state (ss) anisotropy measurements, 〈*r*_c, ∞_〉_tr,ss_. (Bottom) From the residual anisotropy of the donor and acceptor fluorophores, the distance uncertainty relating to the orientation factor *κ*^2^, Δ*R*_app_(*κ*^2^), was estimated for the different dye pairs, as described in the Supplementary Note 8. Based on the distance uncertainty, Δ*R*_app_(*κ*^2^), a threshold is derived to filter the datasets based on the sticking propensity of the dyes, a maximum allowed distance uncertainty of ≤ 10% (shaded grey region) leads to a dye-independent threshold for 〈*r*_c, ∞_〉 of 0.25. **(f)** The apparent dynamic shift 〈ds〉 shows a strong correlation with the combined residual anisotropy 〈*r*_c, ∞_〉 over all measured dye pairs (top left, Pearson correlation coefficient 0.73), indicating that sticking interactions can lead to a false-positive detection of conformational dynamics. A higher correlation between 〈ds〉 and 〈*r*_c, ∞_〉 is observed when the dyes pairs are analyzed separately. (**g**) The structural flexibility of MalE was estimated based on the residual dynamic shift after filtering using the distance uncertainty threshold shown in **e**. The residual dynamic shift is converted into a corresponding distance fluctuation assuming a two-state dynamic exchange that is symmetric around the center distance *R*_〈*E*〉_ (see methods). The residual distance fluctuations obtained for control measurements on double-stranded DNA (ds_DNA_ = 0.0026 ± 0.0044) is shown as a black line (gray areas represent the 1σ, 2σ and 3σ confidence intervals).

The experimental FRET efficiencies 〈*E*〉 for MalE from all labs (Fig. 2) were utilized to determine *R*_〈*E*〉_ for each lab (Table 1 and Supplementary Table 3) using the Förster equation (Eq. 2):

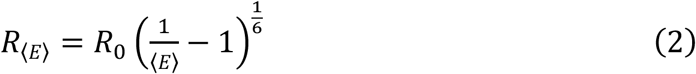

The Förster radius of Alexa546-Alexa647 on MalE was determined to be *R*_0_ = 6.5±0.3 nm (see Supplementary Note 7). In Fig. 5d, we display the correlation between the experimental observable *R*_〈*E*〉_ and predicted 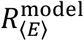 using the known, well-defined structures of the apo and holo states showing an uncertainty of 3-5 Å over all mutants. In agreement with the predictions by Peulen et al.^64^, this accuracy is achieved despite stochastic protein labeling, which could result in drastically different charge environments and accessible volumes of the fluorophores depending on their locations on the protein, as evidenced by the varying dye behavior at different labeling positions (Fig. 5). Over the course of this study, three labs studied two additional MalE mutants (MalE-4: K34C-N205C and MalE-5: T36C-N205C) which were designed to provide a larger FRET efficiency contrast between the apo and holo states, complementing the results of the other variants (Table 1).

Closer inspection of Fig. 5d reveals the largest deviation for MalE-1, which also showed a significant dynamic shift in the *E*-*τ* plot (Fig. 4f and Supplementary Fig. 11). Therefore, we investigated whether dye-protein interactions play a role for the donor or acceptor dye by measuring the fluorescence lifetime and the time-resolved and steady state anisotropies of single-cysteine variants of MalE (Supplementary Note 8, Supplementary Table 5, 11 and 12, and Fig. 5b). These results show that labeling at residue 352 strongly promotes sticking to the protein surface indicated by multiexponential fluorescence lifetimes and a high residual anisotropy, *r*_∞_, for both the donor and acceptor fluorophores (*r*_∞_ > 0.25), while labeling at residue 29 only shows sticking for the donor (*r*_∞, *D*_ > 0.30, *r*_∞, *A*_ ~ 0.12). However, at other positions (e.g., residue 186), free rotation is possible for both dyes (Supplementary Tables 5 and 11). These position-specific interactions can cause the deviations of the experimentally determined distances from the AV model (Fig. 5d, Table 1) and the apparent dynamic shift for mutant MalE-1 (Fig. 4f). A more accurate prediction of the model distances is obtained when the dye sticking is accounted for using the accessible contact volume, ACV^61^, approach (Fig. 5c). When labeling the protein on opposite sides, the dye-surface interactions in the ACV model generally results in a reduced model distance (Fig. 5d, Table 1), which leads to a significant improvement of the accuracy for the outlying mutants.

It has been previously suggested to use the combined residual anisotropy of D and A, computed via 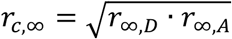, as a criterion for filtering out dye-related artifacts in FRET-assisted structural modeling with an empirical threshold of *r*_c, ∞_ < 0.2^11,65^. To further investigate dye-specific sticking, three labs performed measurements of different MalE mutants with additional dye pairs (Alexa546-AbbSTAR635P, Atto532-Atto643 and Alexa Fluor 488 (Alexa488)-Alexa647) and determined the residual anisotropies and distance uncertainties based on the orientation factor κ^2^ (Fig. 5e, top; Supplementary Table 13 and 14, Supplementary Note 8 and 9). These results depend on the dye-pair, protein and labeling position investigated and have to be addressed individually for the specific system being studied. In this case, the dye pair Alexa546-Alexa647 showed the highest combined anisotropies (Supplementary Fig. 12a, Supplementary Table 13). This is attributed mainly to the donor dye Alexa546 as the combined anisotropy remains high for a different acceptor (Alexa546-AbbSTAR635P) but is reduced markedly for another donor fluorophore (Alexa488-Alexa647). To derive a robust and well-defined threshold for recognizing measurements with dye artifacts, we determined the uncertainty in the FRET-derived distances, Δ*R*_app_(κ^2^), that originates from the uncertainty of the orientation factor κ^2^. Previous approaches have estimated the uncertainty in κ^2^ from the residual anisotropy in terms of rotational restrictions (wobbling-in-a-cone model)^65–68^. Here, we used a ‘diffusion with traps’ model suggested by S. Kalinin, which assumes two dye populations (free and trapped) and relates the residual anisotropies to the fraction of dyes sticking to the surface of the labeled biomolecule (for details, see Supplementary Note 9). Based on the estimated distance uncertainty, we propose a threshold of Δ*R*_app_(κ^2^) < 10% to identify measurements with dye-related artifacts (Fig. 5e, bottom). This threshold corresponds to a combined residual anisotropy of 0.25, similar to the previous empirical threshold value of ~0.211,65.

Next, we investigated whether dye sticking could indeed cause an apparent dynamic shift in the *E*-*τ* plot as seen for MalE-1 with the dye pair Alexa546-Alexa647 (Fig. 4f). For the effect to be observable in the *E*-*τ* plot, the exchange between the free and trapped species must occur faster than the diffusion time of ~1 ms, otherwise the two species would be observable as individual peaks. We observed a correlation between the lab-averaged apparent dynamic shift (ds) and combined residual anisotropy 〈*r*_c, ∞_〉 over all dye pairs (Pearson’s r = 0.73), with a stronger correlation being observed for each dye pair individually (Fig. 5f). As conformational dynamics should be independent of the labels used, we conclude that dye sticking is responsible for the observed apparent dynamic shift. Interestingly, the *x*-intercept of the linear fit is between 0.1 and 0.2, suggesting a dye-dependent anisotropy threshold needs to be considered. When applying the criteria 〈*r*_c, ∞_〉 < 0.25 to MalE-1 (Supplementary Fig. 12b), only the dye pair Atto532-Atto643 could be used, which also showed a significantly reduced apparent dynamic shift (Supplementary Fig. 12c). A lifetime analysis of Alexa546 donor only molecules from MalE-1 showed donor quenching that is not observed at other positions, which confirms that labeling at position 352 is especially problematic (see Supplementary Fig. 12c, Supplementary Note 10 and Supplementary Table 5).

Using the above criteria of 〈*r*_c, ∞_〉 < 0.25 to minimize the influence of dye artifacts on the dynamic shift, we were interested in finding out whether the observed dynamic shift for the other MalE mutants could be indicative of low-amplitude, fast conformational fluctuations. A *p*-test analysis between the dynamic shift for DNA rulers and protein samples (*p* < 0.05) indicated that the dynamic shift calculated after filtering out dye artifacts is still significant for various protein variants (Supplementary Note 11, Supplementary Table 8). To estimate the magnitude of the conformational fluctuations necessary to generate the observed dynamic shifts (Fig. 4f and Supplementary Table 8), we assume that the dynamics occur between two nearby states with interdye distances of *R*_〈*E*〉_, ± *δR*, where δ*R* is the amplitude of the fluctuation^59^ (Fig. 5g, Supplementary Note 12 and Supplementary Table 8). This inferred distance fluctuation must be interpreted as an upper bound for the conformational flexibility because other factors are likely to contribute to the dynamic shift such as calibration errors, dye blinking or photoisomerization. To account for experimental errors that induce falsepositive dynamic shifts, we consider the dynamic shift obtained from dsDNA molecules as the lower limit (black line in Fig. 5g, ds_DNA_ = 0.0026±0.0044, see Supplementary Note 12), which defines the current detection limit for dynamics in smFRET experiments. The MalE variants 1, 4 and 5 clearly exceed the dynamic shift observed for dsDNA by 2-3 Å (Fig. 5g, Supplementary Fig. 13, Supplementary Table 8). Consistent with the smFRET results, all-atom MD simulations of MalE using the ff14SB force field^69^, which is widely used for folded proteins, clearly indicate the presence of small thermally-induced conformational fluctuations with a standard deviation of up to ~3 Å at the labeling locations used for MalE-1, MalE-4 and MalE-5. This is larger than the typical fluctuations on the order of 1 Å^70^ and leads to a broadening of the inter-residue distance distributions for the used FRET pairs (Supplementary Note 13). We conclude that the residual dynamic shift observed in the experiments can be sufficiently explained by a combination of measurement uncertainty and small-scale structural fluctuations. Note that such small-scale distance fluctuations can be amplified in FRET experiments because the dye linker can act as a lever arm, leading to an enhancement in the FRET contrast if labeling positions are chosen appropriately. A detailed discussion of the theoretical limits for detecting dynamics in smFRET experiments using BVA or the *E*-*τ* is given in Supplementary Note 14.

### Quantitative analysis of U2AF2

The structural characterization of conformationally flexible U2AF2 is much more complex and a simple distance comparison as for MalE is not possible. Nonetheless, we asked ourselves what information smFRET measurements could provide for such a dynamic system. We first surveyed the structural information available on the conformational ensemble of apo U2AF2 determined using NMR and SAXS^27^. The high flexibility of the linker allows for a heterogeneous ensemble of possible conformations (Fig. 6a). To assess how this conformational heterogeneity translates into the expected smFRET distributions, we quantified the FRET efficiency for each of the 200 conformers available from the NMR/SAXS derived full ensemble of apo U2AF2^27^ using AV calculations. Notably, conformations with similar center-of-mass (COM) distances between the domains could show vastly different FRET efficiencies (Fig. 6a-b). This occurs because rotations of the domains can result in the dyes pointing towards or away from each other (Fig. 6a, right). Due to this degeneracy, the single-distance information provided here is insufficient to capture the full structural complexity of the apo state.

**Fig. 6.**
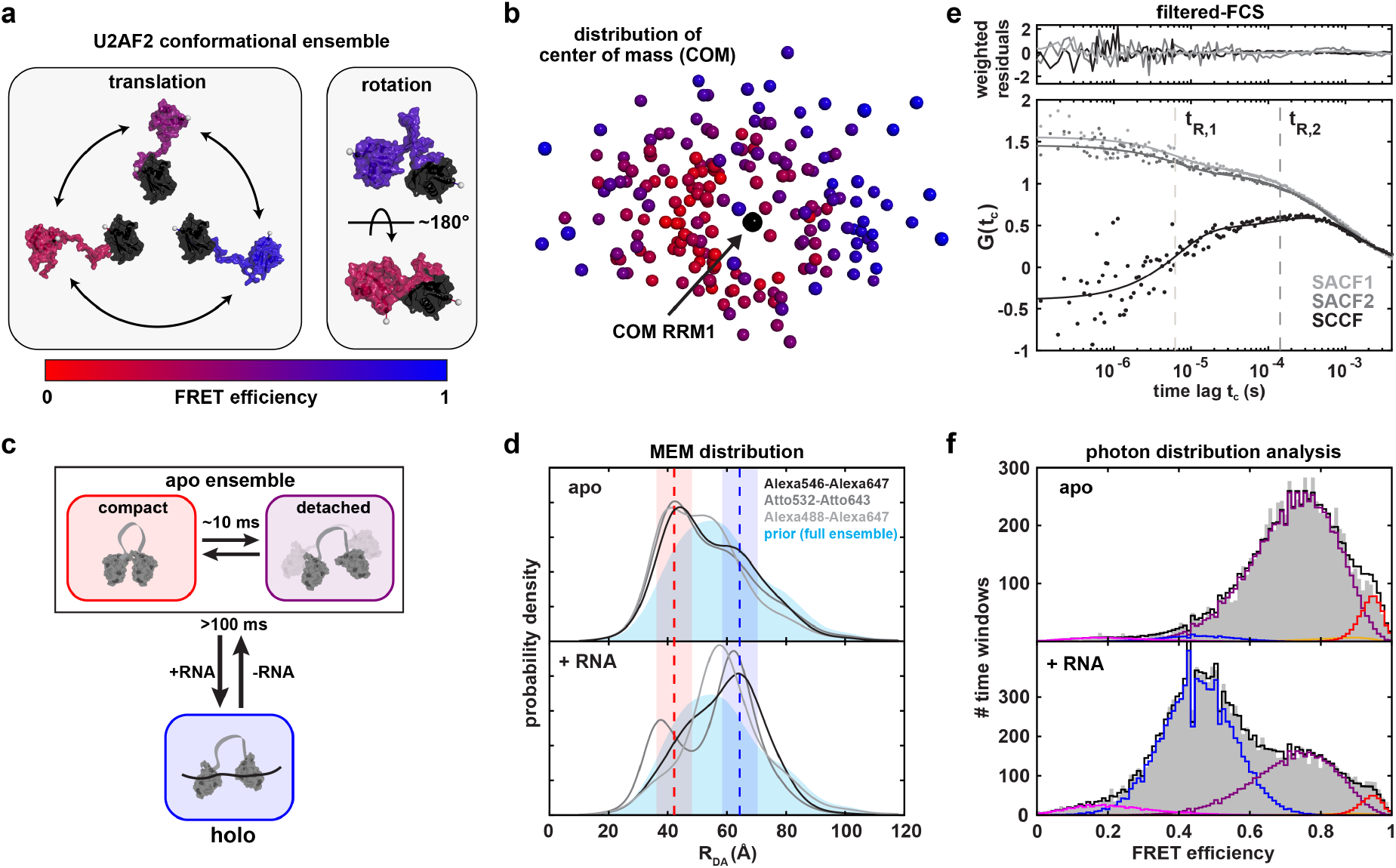
Structural characterization of U2AF2 labeled with Atto532-Atto643 at positions 187 and 326. **(a)** Structural flexibility in the conformational ensemble of U2AF2 is given by translational (left) and rotational (right) movement of the two domains. Representative structures are taken from the ensemble determined using NMR and SAXS measurements by Huang et al.^27^. **(b)** Degeneracy of structural states in FRET measurements. The position of the two domains of U2AF2 is illustrated by the center-of-masses of the C_α_ atoms (COM) in RRM2 (residues 150-227, colored) with respect to RRM1 (residues 260-329, black) for the 200 structures of the conformational ensemble from Huang et al.^27^. The COM of RRM2 is color-coded according to the FRET efficiency of the conformation based on AV3 calculations. Structures with similar COM distances can exhibit different FRET efficiencies due to rotation of the domains. **(c)** A schematic of the kinetic model used for the global dynamic PDA of U2AF2. In the apo state, the protein fluctuates between two states, a defined compact state and the detached ensemble. The rapid dynamics within the detached ensemble are not seen in PDA due to kinetic averaging and the degeneracy of the ensemble with respect to the FRET efficiency (see panel b). The holo state is populated by binding of RNA. Exchange between the apo and holo states occurs on timescales >100 ms (estimated using the known *K_d_*) and is thus too slow to be visible in the diffusion-based smFRET experiments. **(d)** Distance distributions obtained from a donor fluorescence decay analysis by a model-free MEM approach (Supplementary Note 15). The distance distribution from the full NMR/SAXS ensemble^27^ (shown in light blue) was used as the prior distribution. The expected interdye distances for the resolved structure of the compact apo and open holo states are shown as red and blue dashed lines (PDB: 2YH0, 2YH1) with the shaded areas indicating the distance broadening due to the flexible dye linkers of 6 Å. The distribution in the donor-acceptor distance *R_DA_* obtained by the MEM analysis for different dye pairs (see grey shading) is shown. See Supplementary Note 15 for details. **(e)** Filtered fluorescence correlation spectroscopy reveals conformational dynamics in the U2AF2 apo ensemble on two timescales, *t*_*R*, 1_ = 9±3 μs and *t*_*R*, 2_ = 300±90 μs, averaged over all reporting labs (results from lab 1 are shown). The two species were defined at the lower and upper edge of the FRET efficiency histogram shown in Fig. 2b, top panel, by selecting bursts with *E* ≤ 0.6 and *E* ≥ 0.9 respectively. The two species-autocorrelation functions (SACF) and the two (one correlated with two and two correlated with one) species-cross-correlation functions (SCCF) were globally fit to a single-component diffusion model with two kinetic relaxation times (see methods and Supplementary Note 16 for details). For clarity, only one of the two SCCFs is shown. The weighted residuals are shown above. **(f)** The global PDA analysis was performed globally over both apo (top) and holo (bottom) measurements using time windows of 0.5, 1, 1.5 and 2 ms (the displayed histograms correspond to a 1 ms time window), resulting in a global reduced *χ*^2^ of 1.69. A relaxation time of ~10 ms for the dynamics between the detached ensemble and compact apo state with a small amplitude was determined (orange curve). See Supplementary Fig. 16 for all histograms and Supplementary Note 17 for details of the analysis.

As expected, the significant dynamic shift observed in the smFRET experiments clearly supports the presence of conformational dynamics in the apo state (Fig. 4d-f). To decipher the different kinetics involved and their temporal hierarchy, we applied three analyses that are sensitive to dynamics on different timescales. First, the full conformational heterogeneity of U2AF2 was investigated using the donor fluorescence decay. We infer the distribution of interdye distances for the apo and holo states using a model-free maximum entropy method (MEM, Supplementary Note 15)^71^. To test the consistency between the distance distributions provided by the FRET lifetime analysis and by the NMR/SAXS data, we used the NMR/SAXS full structural ensemble as the prior distribution for the MEM (Fig. 6c-d and Supplementary Note 15). This analysis yielded consistent results for the three dye pairs studied for U2AF2. Notably, the MEM analysis revealed peaks in the probability density at the expected distances for the compact apo conformation and RNA-bound holo structure for all dye combinations (Fig. 6d, dashed lines and Supplementary Note 15). We note that the fluorescence lifetime analysis resolves states on the ns time scale and is therefore less sensitive to dynamic averaging.

Secondly, to assess the dynamics on the microsecond timescale, three groups performed a FRET-FCS or filtered-FCS analysis for the Atto532-Atto643 labeled protein and found at least two relaxation times of 9±3 μs and 300±90 μs (Fig. 6e, Supplementary Table 15, Supplementary Note 16). Control experiments using different dye combinations revealed consistent dynamic timescales (Supplementary Fig. 15). We assign these processes to the fast dynamics of the detached domains and the slower interconversion between compact conformations within the conformational ensemble.

Lastly, we investigated dynamics on the millisecond timescale using a dynamic photon distribution analysis (PDA). Here, we performed a global analysis of the apo and holo measurements using the kinetic model shown in Fig. 6c (Supplementary Note 17 and Supplementary Table 16). We treat the apo state as a two-state system with slow dynamics between a detached ensemble and a well-defined, compact apo conformation. The rapid dynamics within the detached ensemble is empirically described using a broad, static distribution. For the holo measurement, we account for the residual population of apo molecules. Exchange between the holo and apo state is not relevant as the binding and dissociation of the RNA occurs on timescales >100 ms^34^. This global model including all known information provides an approximate description of the measured FRET efficiency histograms (Fig. 6f, Supplementary Fig. 16). The dynamic PDA analysis returned a relaxation time of ~10 ms for the dynamics between the detached ensemble and compact apo state with a very small amplitude (orange curve, Supplementary Fig. 16, Supplementary Table 16). We were also able to accurately determine an interdye distance of = 61 Å in the RNA-bound holo state, which is in very good agreement with the RNA-bound conformation of 63 Å (PDB: 2YH1).

## Discussion

The presented results of our blind study involving 19 labs clearly demonstrate that smFRET can consistently provide accurate distances of conformational states and reliable information on dynamics in proteins. All experiments were performed using established experimental procedures and analyzed with freely-available data analysis routines^3,4,32,33,72–74^, indicating that the presented experiments and the conclusions drawn are accessible to groups with similar technical expertise. Despite the challenges of dealing with proteins samples, we could achieve a similar precision in measured FRET efficiencies for both systems over a large part of the dynamic range of FRET as reported previously for stable oligonucleotide structures^16^ (between ± 0.02 and ± 0.06) (Table 1). The high level of consistency for qualitative detection of large-scale sub-millisecond dynamics in U2AF2 and exclusion thereof for MalE shows that the community is well positioned to deal with dynamic protein systems. In addition, we could establish the wide range of timescales and hierarchy of the exchange dynamics observed in U2AF2. The investigation of the complex dynamics could be improved by using multiple labeling positions to measure additional intramolecular distances^3,11,15,75–77^. Consistent results regarding the dynamic timescales were provided by different laboratories using a correlation analysis, and further improvements would be expected when the experimental conditions are better controlled (Supplementary Fig. 7 and Supplementary Table 15).

The high level of consistency is especially notable given the diversity of the setups (Fig. 3 and Supplementary Fig. 2) and the number of difficulties and pitfalls that can occur. The largest contribution to the spread in the reported mean FRET efficiencies was caused by differences in data analysis that can introduce systematic errors. This is demonstrated by investigating the FRET efficiency changes (〈*E*_holo_〉 – 〈*E*_apo_〉) instead of the absolute FRET efficiency values (Fig. 1d), which reduced the spread by a factor of ~ 3. Having a single person reanalyze the data lead to a similar decrease in the uncertainty of the FRET efficiency for the apo state of U2AF2 (Fig. 2d). The most commonly used calibration procedures are γ determination according to Lee *et al.^32^*, using a linear regression of *1/S_app_* versus *E_app_*, and γ determination according to Kudryavtsev *et al*.^33^ via *E*-*τ* calibration (Supplementary Note 2). In the first approach, multiple samples are needed for the calibration and either requires uniform fluorophore properties across all samples or individual corrections made to the samples that deviate. In the second approach, the system needs to be static. A generalized protocol with unambiguous instructions for each of the calibration steps and minimized number of user-dependent steps would alleviate calibration related issues and further enhance the accuracy of FRET measurements. However, the optimal approach depends on the properties of the measured system, making determination of a generalized protocol challenging.

Upon determination of an accurate FRET efficiency, the next step is to convert FRET-efficiency values to inter-dye and inter-base distances as discussed previously in reference 16. Using the structural model for MalE, we obtained reproducible distances with a precision of 3 Å and an accuracy of 5 Å against structural models (Table 1), values similar to what was determined for dsDNA samples. This is a very positive outcome, given that the DNA standards featured a consistent, homogenous chemical environment for the DNA labeling positions, which was in strong contrast to the much more variable dye environment experienced in the measured proteins.

To improve the distance determination further, two important factors were shown to be useful. First, the interaction of the fluorophores with the surface needs to be included in the accessible volume calculations (Fig 5d)^61^. Secondly, only dye-pairs with a combined residual anisotropy of *r*_c, ∞_ < 0.25 should be used (Fig 5f). By comparing measurements on several dye-pairs, we now give experimental support for the value of *r*_c, ∞_ ≤ 0.25, in line with previously given empirical thresholds^11,65^. In addition, proteins often exist within a family of conformations and thus a distribution of distances is necessary to properly describe the system. This can be seen, for example, by the lifetime distribution of U2AF2, where a MEM was used to estimate the conformational distribution of the ensemble (Fig. 6d). Determining how to best deal with distance distributions for conformational ensembles is one of the challenges for structural biology.

Investigating different dye pairs allowed us to select samples where dye artifacts are minimized, thereby leading to more accurate and robust FRET efficiencies as well as the reliable detection and quantification of the dynamics. A careful inspection of the *E*-*τ* plots for MalE raised the question whether a significant deviation from the static FRET-line was observable, implying the existence of FRET dynamics. Therefore, we investigated the detection limits for conformational dynamics using FRET with a subset of laboratories. We note that dynamic FRET shifts can have several origins and may not be due to conformational motions of the protein. For example, linkers used to attach the fluorophore to the molecule or structural instabilities induced by the labeling, as shown by Sánchez-Rico *et al.* for U2AF2, can lead to FRET dynamics^35^. Additional influences come from donor and acceptor quenching, acceptor blinking and photobleaching, and dye sticking (as shown in Fig. 5f)^42^. Thus, in some cases, it can be advisable to verify the key findings in smFRET measurements with at least two dye pairs and/or with different residue combinations in the protein. Contributions to the dynamics shift that have origins other than FRET dynamics can have both positive as well as negative influences on the apparent dynamics shift values. There, we verified with dsDNA structures, which we treat as relatively static biomolecules, that the observed average shift was small (~0.003 in both the BVA and *E*-*τ* plots). Once the non-FRET-dynamic contributions could be minimized, we still observed significant residual dynamic shifts for MalE. Interpreting these shifts as coming from small-scale conformational dynamics, we establish here a current lower limit for the detection of structural changes via smFRET on the order of ≤ 5 Å. Fluctuations having a similar magnitude were also observed at the labeling positions of various MalE variants in an all-atom MD simulation (Supplementary Note 13).

## Conclusions

The consensus of the smFRET data from 19 laboratories on two protein systems exhibiting dynamic behavior on different timescales offers strong support for the use of smFRET as a robust, versatile and quantitative tool for protein distances and dynamics. Deviations in FRET efficiency measured by the various groups were similar to what was determined using DNA standards. One factor that could improve the consistency between laboratories would be a more robust determination of the detection-correction factors required for calculation of setup-independent accurate FRET efficiencies. We also demonstrated that smFRET allows one to detect and characterize conformational dynamics in proteins and can disentangle the latter from dye quenching, blinking, photobleaching, and sticking. A correlation between the observed dynamic indicator and the combined residual anisotropy allowed us to experimentally validate the threshold criterion of both dyes *r*_c, ∞_ < 0.25 when performing accurate FRET measurements. We also present indications that, when artifacts can be excluded, smFRET allows the sensitive detection of small-scale conformational fluctuations on the Ångström level. Hence, FRET can be used to investigate the dynamic behavior of biomolecular complexes on a wide range of time scales and is a powerful tool for the coming age of dynamic structural biology.

## Methods

Methods, including statements of data availability and any associated accession codes and references, are available in the online methods.

## Acknowledgements

Work in the lab of T.C. was financed by an ERC Starting Grant (ERC-StG 638536—SM-IMPORT), Deutsche Forschungsgemeinschaft within GRK2062 (project C03) and SFB863 (project A13) and an Alexander von Humboldt postdoctoral fellowship (to N.Z.). T.C. and D. C.L acknowledge the support of the LMUexcellent and the Center for integrated protein science Munich (CiPSM). D.C.L. acknowledges the support of the Nanosystems Initiative Munich (NIM) and the LMU via the LMUinnovative program BioImaging Network (BIN). We also acknowledge support via the SFB1035 (German Research Foundation DFG, Sonderforschungsbereich 1035 Projektnummer 201302640, Project A11 to D.C.L. and Project B03 to M.Sa.). C.A.M.S. acknowledges the support by the European Research Council (ERC; grant No. 671208 (hybridFRET)) and by the Deutsche Forschungsgemeinschaft (DFG, grant SE 1195/17-1). Research in the contributing authors labs was financed by the following sources: P.T acknowledges the support by the Deutsche Forschungsgemeinschaft (DFG, German Research Foundation) – Project-ID 201269156 – SFB 1032 (A13) and Project-ID 267681426. T.D.C was supported by the BBSRC (BB/T008032/1) and EPSRC (EP/V034804/1), B.A. was supported by an EPSRC Prize Fellowship. BMBF grant 03Z2EN11 and 03Z22E511 as well as DFG grant SCHL 1896/4-1 (to M.Sc.). SFB960 Project A7 (to D.G.) US National Institutes of Health grant GM122569 (to T.Ha). J.H. acknowledges the Research Foundation Flanders (FWO, projects G0B4915, G0B9922N and G0H3716N) and is indebted to Johan Hofkens at KU Leuven for the used smFRET infrastructure. ERC grant agreement no. 681891 (Prosint) and German Research Foundation (DFG) under Germany’s Excellence Strategy (CIBSS EXC-2189 Project ID 390939984) (to T.Hu.). Royal Society Dorothy Hodgkin Research Fellowship DKR00620 and Research Grant for Research Fellows RGF\R1\180054 (to N.C.R.), by the Wellcome Trust (110164/Z/15/Z to A.N.K.). The Israel Science Foundation (grant 3565/20 to E.L., within the KillCorona – Curbing Coronavirus Research Program), the NIH (grant R01 GM130942 to S.W. and to E.Ler. as a subaward), by the Milner Fund (to E.Ler.), and by the Hebrew University of Jerusalem (start-up funds to E. Ler.). Agence Nationale pour la Recherche (ANR 18-CE11-0004-02, ANR-19-CE44-0009-02, ANR-21-CE11-0034-01, ANR-21-CE11-0026-03 and ANR-10-INBS-04, “Investments for the future” to E.M.)].

## Competing interests

The authors declare no competing interests.

## Author contributions

C.G. and T.C. initiated the study. C.G., G.A., A.B., C.A.M.S., D.C.L. and T.C. designed research. A.B., C.A.M.S., D.C.L. and T.C. supervised the project. R.M. cloned and purified MalE. H.S.K. and M.S. provided U2AF2. C.G. and G.A. performed labeling of MalE and U2AF2 variants for shipment to participating labs. M.P., J.F. and C.A.M.S. designed MalE mutants 4 and 5 in silico. M.P. and J.F. performed initial measurements on MalE mutants 4 and 5. G.A. reperformed the analysis on the provided raw data for U2AF2 and MalE-1 from 8 laboratories. G.A., A.B. and M.P. performed dynamic shift estimation. C.G. performed FCS experiments on MalE variants, time-resolved anisotropy experiments and *R*_0_-determination. M.P. performed time-resolved anisotropy analysis of single labelled MalE cysteine mutants from ensemble measurements as well of all MalE and U2AF2 dye combinations from smFRET measurements. C.G., G.A. and M.P. performed measurements of MalE mutants with additional dye combinations and M.P. and A.B. performed statistical analysis of dynamic shifts and anisotropies. G.A. and A.B. performed estimation of setup-dependent parameters and PDA of U2AF2. G.A. performed the filtered-FCS and A.B. performed TCSPC analysis of U2AF2. C.G., G.A. and M.P. performed smFRET measurements on dsDNA rulers. M.P. performed AV and ACV modelling of dye distributions for MalE and U2AF2. G.M. performed MST experiments. M.d.B. performed confocal scanning experiments for surface-immobilized MalE. All authors were involved in performing comparison experiments and analyzing smFRET data. C.G. and G.A. consolidated data collection of participating labs. C.G. and G.A. designed Fig. 1. C.G., G.A. and M.P. designed Fig. 2. A.B. designed Fig.s 3 and 4. A.B. and M.P. designed Figs. 5 and 6. C.G., G.A., M.P., A.B., C.A.M.S., D.C.L. and T.C. interpreted data and wrote the manuscript in consultation with all authors.

## Online Methods

### Sample preparation of proteins

Double-cysteine mutants of MalE were prepared and labeled using established protocols^60^. Human RRM1,2 L187C-G326C mutant (U2AF2-148-342) was obtained and purified as described in Mackereth et al.^28^.

### Fluorescence labeling of proteins

All fluorophores were purchased as maleimide derivatives from commercial suppliers as listed in Supplementary Table 19. MalE was stochastically labeled as described previously^78^ with fluorophores as indicated in the text with a combined labeling efficiency higher than 70% resulting in a donor-acceptor pairing of at least 20%. Protein stability and functionality (ligand binding) was verified by affinity measurements using microscale thermophoresis^79^. All preparations, i.e., MalE-wildtype, unlabeled cysteine mutants and fluorophore-labeled variants, showed an affinity for maltose between ~1-2 μM (Supplementary Fig. 5) consistent with previously published *K_d_*-values for wild type MalE^80,81^. The stability and labeling of the sample were verified by fluorescence correlation spectroscopy (Supplementary Fig. 18), which excluded the presence of larger aggregates in the samples and confirms that MalE is functional.

U2AF2 was stochastically labeled as described previously in Voith von Voithenberg et al.^34^. The combined labeling efficiency for labeling reactions were 20% and 14% for Alexa546-Alexa647 and Atto532-Atto643 pairs, respectively. For Alexa488-Alexa647, the combined labeling efficiency was found to be 10%. The functionality of the labeled U2AF protein was checked with affinity measurement for U9 RNA, which was found to be 1.2 μM^28^, consistent with the previous reports^34^ (Supplementary Fig. 7d).

### Sample handling

Both protein systems required special handling due to sample instability or aggregate formation, which are both problematic for long-term storage and shipping. The labeled MalE proteins were stored in 50 mM Tris-HCl pH 7.4, 50 mM KCl with 1 mg/ml bovine serum albumin (BSA) at 4°C for less than 7 days. U2AF2 was stored in 20 mM potassium phosphate buffer pH 6.5, 50 mM NaCl and kept in the fridge until used. Both samples were loaded in low-binding Eppendorf tubes (Eppendorf Germany, Catalog number 0030108094) and shipped on ice in a cooling box with overnight shipping to avoid unnecessary freezing and thawing. MalE stock solutions were on the order of 10 to 100 nM concentration and the sent stock solution of U2AF2 was 5-10 μM concentration. Dilution buffer for apo and holo measurement were provided. SmFRET experiments were carried out by diluting the labeled proteins to concentrations of ≈50 pM in 50 mM Tris-HCl pH 7.4, 50 mM KCl supplemented with the ligand maltose at 1 mM concentration. Labeled U2AF2 protein was measured at ~40-100 pM in 20 mM potassium phosphate buffer pH 6.5, 50 mM NaCl. Purchased U9 RNA (Biomers.net GmbH, Ulm, Germany, IBA Solutions for Life Sciences, Göttingen, Germany) was dissolved in RNA-free water and added directly to the solution at a final concentration of 5 μM for the holo measurements. Both proteins were studied on coverslips typically passivated with 1 mg/ml BSA in buffer before adding the sample. The measurements were performed without any photo-stabilizer to keep the measurements as simple as possible to avoid any further source for discrepancies between the groups, e.g., degradation of photostabilizer or use of different photostabilizer concentrations.

### SmFRET data acquisition and analysis

Data acquisition and correction procedures were performed for confocal measurements as described by Hellenkamp *et al*.^16^. The samples were measured using alternating laser excitation mode (ALEX) or Pulsed Interleaved Excitation (PIE) on a confocal microscope as sketched in Supplementary Fig. 2. A description of experimental procedures of all labs are given in Supplementary Note 18.

Briefly, the three recorded intensity time traces for each single-molecule event are:

donor emission after donor excitation: i_*I*_Dem_|_Dex__,
acceptor emission after donor excitation (FRET signal): i_*I*__Aem|Dex_,
and acceptor emission after acceptor excitation: i_*I*_Aem|Aex__.

The apparent (raw) FRET efficiency is computed as:

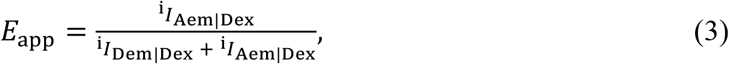

Recorded intensities were corrected for background contributions as:

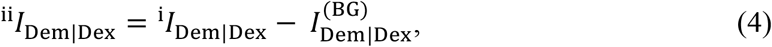

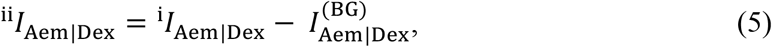

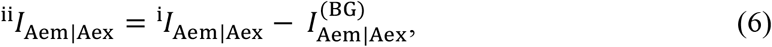

where 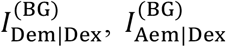 and 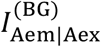 are the respective background signals. Correction factors for spectral crosstalk, *α*, and direct excitation, *δ*, were determined from the donor-only and acceptor-only populations^32^. The corrected acceptor fluorescence after donor excitation, *F_A|D_*, is computed as:

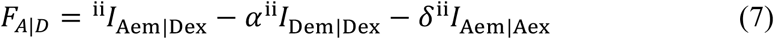

The γ and β factors, correcting for differences in the detection yield and excitation fluxes of the donor and acceptor dyes, were estimated using a global correction procedure using the approach of Lee *et al.* (Supplementary Fig. 3)^32^. Alternatively, when pulsed excitation was used and the sample is known to be static, the γ factor can be determined by fitting the measured population to the static FRET line^33,82^. This allows a good determination of the γ factor when only a single species is present but requires a static sample and the appropriate static FRET line (Supplementary Note 2).

The accurate FRET efficiency *E* and stoichiometry *S* values were then calculated as:

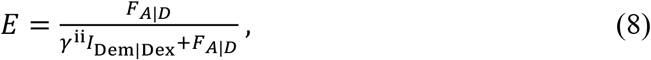

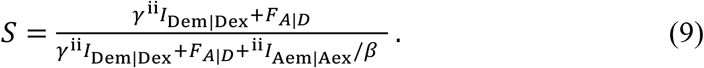

Conversion of accurate FRET efficiencies into distances were done using Eq. 2 with Förster radii determined as described in Supplementary Note 7.

### Detection of Protein Dynamics

In this work, we used the following two approaches to detect conformational dynamics:

#### Burst Variance Analysis (BVA)

In BVA, the presence of dynamics is determined by looking for excess variance in the FRET efficiency data beyond the shot-noise limit. The the standard deviation (σ_*E*_app__) of the apparent FRET efficiency (*E_app_*) is calculated using a fixed photon window of 5 (*n*) over the time period of the individual bursts given by:

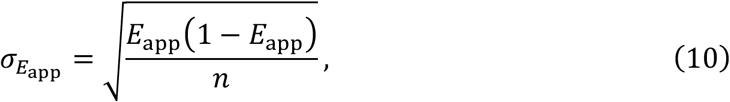

The shot-noise limited standard deviation of the apparent FRET efficiency is generally described by a semi-circle^48^ (Fig. 4a and Supplementary Fig. 11a-d). In the presence of dynamics, the standard deviation for the FRET efficiency within a burst becomes higher than that expected from shot noise. Photophysical effects like bleaching and blinking also give rise to the higher standard deviation beyond the shot-noise limit. Typically, BVA is sensitive to fluctuations in FRET signal of ≳ 100 μs, but these depends on the brightness of the burst and the photon window used.

#### FRET efficiency versus fluorescence-weighted average donor lifetime analysis (E-τ plots)

Two-dimensional histograms of the FRET efficiency *E* and donor fluorescence lifetime 〈*τ*(_D_(*A*)〉_*F*_ (Fig. 4b and Supplementary Fig. 11e-h) were created for single molecule measurements using MFD in combination with pulsed-interleaved excitation (PIE)^33^, described below. Static FRET lines were calculated using the following equation:

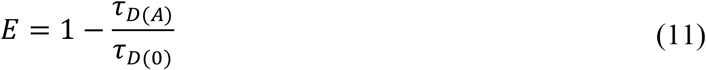

and further modified for linker dynamics^59^. Deviations of FRET populations from the static FRET line can indicate FRET dynamics, which can be due to conformational fluctuations or photophysical dynamics. In addition, a time-resolved FRET analysis of TCSPC data can accurately resolve the distance heterogeneities by revealing multiple components in the decay of the curve and recovers their specific species fractions and FRET rate constants^64^. Dynamics are thus detected from the presence of multiple components in the sub-ensemble decay of a single FRET population. In addition, dynamics that are slower than the fluorescence lifetime (~ 5 ns) are not averaged in the FRET lifetime analysis leading to detection of the full conformational distribution.

### Multiparameter fluorescence detection (MFD) with Pulsed Interleaved Excitation (PIE)

MFD, introduced by Eggeling et al.^83^, combines spectral and polarized detection with picosecond pulsed lasers and time correlated single photon counting (TCSPC), allowing the simultaneous detection of intensity, lifetime, anisotropy and spectral range of the fluorescence signal of single molecules. nsALEX of PIE additionally provide the acceptor lifetime information^33^. Due to the availability of the lifetime information when using pulsed excitation, this approach is well suited for using *E*-*τ*-based analyses.

### AV simulations

The AV approach employs a simple coarse-grained dye model^63^ defined by five parameters: the width and length of the linker, and three radii that define the fluorophore volume (Fig. 5a, Supplementary Table 10). Using these parameters, AV simulations for both fluorophores were calculated by considering the linker flexibility and steric hindrances of the labeled molecule (Fig. 5a). In the ACV model^61^, the residual anisotropy was used to estimate the fraction of sticking dyes. In the computation of the FRET-averaged model distances, the occupancy of a thin surface layer (~3 Å) was then increased such that its fraction matches the amount of interacting dye detected in the experiment (Fig. 5b Supplementary Table 10).

## Notes

### Competing Interest Statement

Tim Craggs and Achilles Kapanidis, two of the authors are founders of different companies selling single-molecule fluorescence microscopes (Exciting Instruments, Oxford Nanoimager).

## References

1. Lerner, E. et al. FRET-based dynamic structural biology: Challenges, perspectives and an appeal for open-science practices. eLife 10, e60416 (2021).

2. Lerner, E. et al. Toward dynamic structural biology: Two decades of single-molecule Förster resonance energy transfer. Science. 359, eaan1133 (2018).

3. Muschielok, A. et al. A nano-positioning system for macromolecular structural analysis. Nat. Methods 5, 965–971 (2008).

4. Kalinin, S. et al. A toolkit and benchmark study for FRET-restrained high-precision structural modeling. Nat. Methods 9, 1218–1225 (2012).

5. Craggs, T. D. & Kapanidis, A. N. Six steps closer to FRET-driven structural biology. Nat. Methods 9, 1157–1159 (2012).

6. Voith von Voithenberg, L. & Lamb, D. C. Single Pair Förster Resonance Energy Transfer: A Versatile Tool To Investigate Protein Conformational Dynamics. BioEssays 40, 1–14 (2018).

7. Hohlbein, J., Craggs, T. D. & Cordes, T. Alternating-laser excitation: Single-molecule FRET and beyond. Chem. Soc. Rev. 43, 1156–1171 (2014).

8. Krainer, G., Hartmann, A. & Schlierf, M. FarFRET: Extending the Range in Single-Molecule FRET Experiments beyond 10 nm. Nano Lett. 15, 5826–5829 (2015).

9. Muschielok, A. & Michaelis, J. Application of the nano-positioning system to the analysis of fluorescence resonance energy transfer networks. J. Phys. Chem. B 115, 11927–11937 (2011).

10. Sali, A. et al. Outcome of the First wwPDB Hybrid/Integrative Methods Task Force Workshop. Structure 23, 1156–1167 (2015).

11. Hellenkamp, B., Wortmann, P., Kandzia, F., Zacharias, M. & Hugel, T. Multidomain structure and correlated dynamics determined by self-consistent FRET networks. Nat. Methods 14, 176–182 (2017).

12. Choi, U. B. et al. Single-molecule FRET-derived model of the synaptotagmin 1-SNARE fusion complex. Nat. Struct. Mol. Biol. 17, 318–324 (2010).

13. Dimura, M. et al. Automated and optimally FRET-assisted structural modeling. Nat. Commun. 11, 1–14 (2020).

14. Lerner, E., Ingargiola, A. & Weiss, S. Characterizing highly dynamic conformational states: The transcription bubble in RNAP-promoter open complex as an example. J. Chem. Phys. 148, 123315 (2018).

15. Craggs, T. D. et al. Substrate conformational dynamics facilitate structure-specific recognition of gapped DNA by DNA polymerase. Nucleic Acids Res. 47, 10788–10800 (2019).

16. Hellenkamp, B. et al. Precision and accuracy of single-molecule FRET measurements—a multi-laboratory benchmark study. Nat. Methods 15, 669–676 (2018).

17. Rout, M. P. & Sali, A. Principles for Integrative Structural Biology Studies. Cell 177, 1384–1403 (2019).

18. Sali, A. From integrative structural biology to cell biology. J. Biol. Chem. 296, 100743, 1–15 (2021).

19. Burley, S. K. et al. PDB-Dev: a Prototype System for Depositing Integrative/Hybrid Structural Models. Structure 25, 1317–1318 (2017).

20. Davidson, A. L., Dassa, E., Orelle, C. & Chen, J. Structure, Function, and Evolution of Bacterial ATP-Binding Cassette Systems. Microbiol. Mol. Biol. Rev. 72, 317–364 (2008).

21. Mächtel, R., Narducci, A., Griffith, D. A., Cordes, T. & Orelle, C. An integrated transport mechanism of the maltose ABC importer. Res. Microbiol. 170, 321–337 (2019).

22. Malik, A. Protein fusion tags for efficient expression and purification of recombinant proteins in the periplasmic space of E. coli. 3 Biotech 6, 1–7 (2016).

23. Berntsson, R. P. A., Smits, S. H. J., Schmitt, L., Slotboom, D. J. & Poolman, B. A structural classification of substrate-binding proteins. FEBS Lett. 584, 2606–2617 (2010).

24. Fukami-Kobayashi, K., Tateno, Y. & Nishikawa, K. Domain dislocation: A change of core structure in periplasmic binding proteins in their evolutionary history. J. Mol. Biol. 286, 279–290 (1999).

25. Banerjee, H., Rahn, A., Davis, W. & Singh, R. Sex lethal and U2 small nuclear ribonucleoprotein auxiliary factor (U2AF65) recognize polypyrimidine tracts using multiple modes of binding. Rna 9, 88–99 (2003).

26. Sickmier, E. A. et al. Structural Basis for Polypyrimidine Tract Recognition by the Essential Pre-mRNA Splicing Factor U2AF65. Mol. Cell 23, 49–59 (2006).

27. Huang, J. R. et al. Transient electrostatic interactions dominate the conformational equilibrium sampled by multidomain splicing factor U2AF65: A combined NMR and SAXS study. J. Am. Chem. Soc. 136, 7068–7076 (2014).

28. MacKereth, C. D. et al. Multi-domain conformational selection underlies pre-mRNA splicing regulation by U2AF. Nature 475, 408–413 (2011).

29. Kapanidis, A. N. et al. Fluorescence-aided molecule sorting: Analysis of structure and interactions by alternating-laser excitation of single molecules. Proc. Natl. Acad. Sci. U. S. A. 101, 8936–8941 (2004).

30. Kapanidis, A. N. et al. Alternating-laser excitation of single molecules. Acc. Chem. Res. 38, 523–533 (2005).

31. Müller, B. K., Zaychikov, E., Bräuchle, C. & Lamb, D. C. Pulsed interleaved excitation. Biophys. J. 89, 3508–3522 (2005).

32. Lee, N. K. et al. Accurate FRET measurements within single diffusing biomolecules using alternating-laser excitation. Biophys. J. 88, 2939–2953 (2005).

33. Kudryavtsev, V. et al. Combining MFD and PIE for accurate single-pair Förster resonance energy transfer measurements. ChemPhysChem 13, 1060–1078 (2012).

34. Von Voithenberg, L. V. et al. Recognition of the 3′ splice site RNA by the U2AF heterodimer involves a dynamic population shift. Proc. Natl. Acad. Sci. U. S. A. 113, E7169–E7175 (2016).

35. Sánchez-Rico, C., Voith von Voithenberg, L., Warner, L., Lamb, D. C. & Sattler, M. Effects of Fluorophore Attachment on Protein Conformation and Dynamics Studied by spFRET and NMR Spectroscopy. Chem. - A Eur. J. 23, 14267–14277 (2017).

36. Eggeling, C., Widengren, J., Rigler, R. & Seidel, C. A. M. Photobleaching of Fluorescent Dyes under Conditions Used for Single-Molecule Detection: Evidence of Two-Step Photolysis. Anal. Chem. 70, 2651–2659 (1998).

37. Chung, H. S., McHale, K., Louis, J. M. & Eaton, W. A. SI-Single-Molecule Fluorescence Experiments Determine Protein Folding Transition Path Times. Science. 335, 981–984 (2012).

38. Ramanathan, R. & Muñoz, V. A Method for extracting the free energy surface and conformational dynamics of fast-folding proteins from single molecule photon trajectories. J. Phys. Chem. B 119, 7944–7956 (2015).

39. McKinney, S. A., Joo, C. & Ha, T. Analysis of single-molecule FRET trajectories using hidden Markov modeling. Biophys. J. 91, 1941–1951 (2006).

40. Liu, Y., Park, J., Dahmen, K. A., Chemla, Y. R. & Ha, T. A comparative study of multivariate and univariate hidden Markov modelings in time-binned single-molecule FRET data analysis. J. Phys. Chem. B 114, 5386–5403 (2010).

41. Bronson, J. E., Fei, J., Hofman, J. M., Gonzalez, R. L. & Wiggins, C. H. Learning rates and states from biophysical time series: A Bayesian approach to model selection and single-molecule FRET data. Biophys. J. 97, 3196–3205 (2009).

42. Margittai, M. et al. Single-molecule fluorescence resonance energy transfer reveals a dynamic equilibrium between closed and open conformations of syntaxin 1. Proc. Natl. Acad. Sci. U. S. A. 100, 15516–15521 (2003).

43. Diez, M. et al. Proton-powered subunit rotation in single membrane-bound F 0F1-ATP synthase. Nat. Struct. Mol. Biol. 11, 135–141 (2004).

44. Torres, T. & Levitus, M. Measuring conformational dynamics: A new FCS-FRET approach. J. Phys. Chem. B 111, 7392–7400 (2007).

45. Felekyan, S., Sanabria, H., Kalinin, S., Kühnemuth, R. & Seidel, C. A. M. Analyzing Förster resonance energy transfer with fluctuation algorithms. in Methods in Enzymology (ed. Tetin, S. Y. B. T.-M. in E.) 519, 39–85 (Academic Press, 2013).

46. Felekyan, S., Kalinin, S., Sanabria, H., Valeri, A. & Seidel, C. A. M. Filtered FCS: Species auto-and cross-correlation functions highlight binding and dynamics in biomolecules. ChemPhysChem 13, 1036–1053 (2012).

47. Olofsson, L. et al. Fine tuning of sub-millisecond conformational dynamics controls metabotropic glutamate receptors agonist efficacy. Nat. Commun. 5, 5206 (2014).

48. Torella, J. P., Holden, S. J., Santoso, Y., Hohlbein, J. & Kapanidis, A. N. Identifying molecular dynamics in single-molecule fret experiments with burst variance analysis. Biophys. J. 100, 1568–1577 (2011).

49. Tomov, T. E. et al. Disentangling subpopulations in single-molecule FRET and ALEX experiments with photon distribution analysis. Biophys. J. 102, 1163–1173 (2012).

50. Kalinin, S., Valeri, A., Antonik, M., Felekyan, S. & Seidel, C. A. M. Detection of Structural Dynamics by FRET: A Photon Distribution and Fluorescence Lifetime Analysis of Systems with Multiple States. J. Phys. Chem. B 114, 7983–7995 (2010).

51. Gopich, I. V. & Szabo, A. Theory of the energy transfer efficiency and fluorescence lifetime distribution in single-molecule FRET. Proc. Natl. Acad. Sci. U. S. A. 109, 7747–7752 (2012).

52. Nettels, D., Gopich, I. V., Hoffmann, A. & Schuler, B. Ultrafast dynamics of protein collapse from single-molecule photon statistics. Proc. Natl. Acad. Sci. U. S. A. 104, 2655–2660 (2007).

53. Hoffmann, A. et al. Quantifying heterogeneity and conformational dynamics from single molecule FRET of diffusing molecules: Recurrence analysis of single particles (RASP). Phys. Chem. Chem. Phys. 13, 1857–1871 (2011).

54. Gopich, I. V. & Szabo, A. Decoding the pattern of photon colors in single-molecule FRET. J. Phys. Chem. B 113, 10965–10973 (2009).

55. Chung, H. S. & Gopich, I. V. Fast single-molecule FRET spectroscopy: Theory and experiment. Phys. Chem. Chem. Phys. 16, 18644–18657 (2014).

56. Pirchi, M. et al. Photon-by-Photon Hidden Markov Model Analysis for Microsecond Single-Molecule FRET Kinetics. J. Phys. Chem. B 120, 13065–13075 (2016).

57. Harris, P. D. et al. Multi-parameter photon-by-photon hidden Markov modeling. Nat. Commun. 13, 1000 (2022).

58. Ingargiola, A., Weiss, S. & Lerner, E. Monte Carlo Diffusion-Enhanced Photon Inference: Distance Distributions and Conformational Dynamics in Single-Molecule FRET. J. Phys. Chem. B 122, 11598–11615 (2018).

59. Barth, A. et al. Unraveling multi-state molecular dynamics in single-molecule FRET experiments. I. Theory of FRET-lines. J. Chem. Phys. 156, 141501 (2022).

60. De Boer, M. et al. Conformational and dynamic plasticity in substrate-binding proteins underlies selective transport in ABC importers. eLife 8, e44652 (2019).

61. Dimura, M. et al. Quantitative FRET studies and integrative modeling unravel the structure and dynamics of biomolecular systems. Current Opinion in Structural Biology 40, 163–185 (2016).

62. Beckers, M., Drechsler, F., Eilert, T., Nagy, J. & Michaelis, J. Quantitative structural information from single-molecule FRET. Faraday Discuss. 184, 117–129 (2015).

63. Sindbert, S. et al. Accurate distance determination of nucleic acids via Förster resonance energy transfer: Implications of dye Linker length and rigidity. J. Am. Chem. Soc. 133, 2463–2480 (2011).

64. Peulen, T. O., Opanasyuk, O. & Seidel, C. A. M. Combining Graphical and Analytical Methods with Molecular Simulations to Analyze Time-Resolved FRET Measurements of Labeled Macromolecules Accurately. J. Phys. Chem. B 121, 8211–8241 (2017).

65. Dale, R. E., Eisinger, J. & Blumberg, W. E. The orientational freedom of molecular probes. The orientation factor in intramolecular energy transfer. Biophys. J. 26, 161–193 (1979).

66. Dale, R. E. & Eisinger, J. Intramolecular distances determined by energy transfer. Dependence on orientational freedom of donor and acceptor. Biopolymers 13, 1573–1605 (1974).

67. Ivanov, V., Li, M. & Mizuuchi, K. Impact of emission anisotropy on fluorescence spectroscopy and FRET distance measurements. Biophys. J. 97, 922–929 (2009).

68. Eilert, T., Kallis, E., Nagy, J., Röcker, C. & Michaelis, J. Complete Kinetic Theory of FRET. J. Phys. Chem. B 122, 11677–11694 (2018).

69. Maier, J. A. et al. ff14SB: Improving the Accuracy of Protein Side Chain and Backbone Parameters from ff99SB. J. Chem. Theory Comput. 11, 3696–3713 (2015).

70. Zaccai, G. How soft is a protein? A protein dynamics force constant measured by neutron scattering. Science. 288, 1604–1607 (2000).

71. Vinogradov, S. A. & Wilson, D. F. Recursive maximum entropy algorithm and its application to the luminescence lifetime distribution recovery. Appl. Spectrosc. 54, 849–855 (2000).

72. Ingargiola, A., Lerner, E., Chung, S. Y., Weiss, S. & Michalet, X. FRETBursts: An open source toolkit for analysis of freely-diffusing Single-molecule FRET. PLoS One 11, 39198 (2016).

73. Schrimpf, W., Barth, A., Hendrix, J. & Lamb, D. C. PAM: A Framework for Integrated Analysis of Imaging, Single-Molecule, and Ensemble Fluorescence Data. Biophys. J. 114, 1518–1528 (2018).

74. Ambrose, B. et al. The smfBox is an open-source platform for single-molecule FRET. Nat. Commun. 11, 5641 (2020).

75. Knight, J. L., Mekler, V., Mukhopadhyay, J., Ebright, R. H. & Levy, R. M. Distance-restrained docking of rifampicin and rifamycin SV to RNA polymerase using systematic FRET measurements: Developing benchmarks of model quality and reliability. Biophys. J. 88, 925–938 (2005).

76. Kapanidis, A. N. et al. Initial transcription by RNA polymerase proceeds through a DNA-scrunching mechanism. Science. 314, 1144–1147 (2006).

77. Sanabria, H. et al. Resolving dynamics and function of transient states in single enzyme molecules. Nat. Commun. 11, 1231 (2020).

78. Gouridis, G. et al. Conformational dynamics in substrate-binding domains influences transport in the ABC importer GlnPQ. Nat. Struct. Mol. Biol. 22, 57–64 (2015).

79. Jerabek-Willemsen, M. et al. MicroScale Thermophoresis: Interaction analysis and beyond. J. Mol. Struct. 1077, 101–113 (2014).

80. Hall, J. A., Gehring, K. & Nikaido, H. Two modes of ligand binding in maltose-binding protein of Escherichia coli: Correlation with the structure of ligands and the structure of binding protein. J. Biol. Chem. 272, 17605–17609 (1997).

81. Kim, E. et al. A single-molecule dissection of ligand binding to a protein with intrinsic dynamics. Nat. Chem. Biol. 9, 313–318 (2013).

82. Sisamakis, E., Valeri, A., Kalinin, S., Rothwell, P. J. & Seidel, C. A. M. Accurate Single-Molecule FRET Studies Using Multiparameter Fluorescence Detection. Methods Enzymol. 475, 455–514 (2010).

83. Eggeling, C. et al. Data registration and selective single-molecule analysis using multiparameter fluorescence detection. J. Biotechnol. 86, 163–180 (2001).

