## Supplementary Information for "Reliability and accuracy of single-molecule FRET studies for characterization of structural dynamics and distances in proteins"

Supplementary Figure 2: Schematics of the experimental setups. 64
Supplementary Figure 3: Correction procedure when using a global γ 65
Supplementary Figure 4: Primer design and sequences for creation of MalE variants 66
Supplementary Figure 5: Binding affinity measurements of maltose to MalE using microscale thermophoresis 67
Supplementary Figure 6: Millimolar maltose concentration does not influence fluorophore properties 69
Supplementary Figure 7: Temperature and concentration dependence of RNA binding to U2AF2. 70
Supplementary Figure 8: Comparison of the FRET efficiency reported for different labs and after reanalysis for the MalE-1 mutant under apo conditions. 70
Supplementary Figure 9: Sample-dependent setup statistics 71
Supplementary Figure 10: Correlations between setup-dependent parameters 72
Supplementary Figure 11: Overview of conformational dynamics and determination of the dynamic shift on the sub-millisecond timescale in MalE labeled with Alexa546-Alexa647 and U2AF2 labeled with Atto532-Atto643. 73
Supplementary Figure 12: Dye-specific sticking interactions in proteins and residual apparent dynamic shift upon removal of sticky dyes 75
Supplementary Figure 13: Estimated conformational flexibility of MalE based on the residual dynamic shift for different donor-acceptor pairs 76
Supplementary Figure 14: Comparison of the distance distributions obtained using different donor-acceptor dye pairs. 77
Supplementary Figure 15: Filtered fluorescence correlation spectroscopy curves for U2AF2. 78
Supplementary Figure 16: A global dynamic photon distribution analysis (PDA) of apo and holo measurements of U2AF2 labeled with the Atto532-Atto643 dye pair 79
Supplementary Figure 17: The very-high-FRET population in U2AF2. 80
Supplementary Figure 18: FCS experiments to rule out the presence of large aggregates 81
Supplementary Figure 19: A global dynamic photon distribution analysis (PDA) of the apo and holo measurements of U2AF2 labeled with the Alexa546-Alexa647 dye-pair. 82
*Supplementary Tables* 83
Supplementary Table 1: FRET efficiency correction factors reported by the participating labs for MalE constructs labeled with Alexa546-Alexa647 83
Supplementary Table 2: FRET efficiency correction factors reported by the 7 labs for apo and holo measurements on U2AF2 labeled with Atto532-Atto643 84
Supplementary Table 3: Reported mean FRET Efficiency and standard deviation for MalE samples by the participating laboratories 85
Supplementary Table 4: Reported and reanalyzed mean FRET efficiency and standard deviations for the U2AF2 experiments: 86

### Supplementary Notes

##### Supplementary Note 1: Comparison between µsALEX and PIE.

With the lifetime information, it is possible to directly access distances and distance distributions via analysis of the donor fluorescence decay[1,2] and detect dynamics on the nano- to millisecond timescale in the E-τ plot, as we have shown for the apo state of U2AF2. The advantage of µsALEX is that it is less expensive to implement than nsALEX/PIE and that the continuous wave excitation exerts less photophysical stress on the fluorophores (e.g., bleaching or blinking) compared to the high peak irradiance for pulsed excitation[3]. NsALEX/PIE have the advantage that they provide the fluorescence lifetime information of the donor and acceptor fluorophore[4]. Via the lifetime information, it is also possible to detect changes of the donor and acceptor quantum yields due to protein or ligand interactions with the dye. In addition, the faster alternation timescale of nsALEX/PIE enables higher time-resolution for fluorescence correlation spectroscopy and reduces the spread in the stoichiometry distribution due to diffusion of molecules through the confocal volume during the alternation period. Combining pulsed excitation with polarized excitation and detection[4,5] further adds the fluorescence anisotropy information on the single-molecule or sub-ensemble level, allowing one to monitor size changes of the biomolecule (e.g., through the binding of interaction partners) and detect changes of the local environment of the fluorophore such as dye sticking (as seen for MalE-1) directly from the single-molecule dataset.

The spread in the results in Fig. 3d indicates that the direct probing of the acceptor in ALEX/PIE experiments was applied differently by the participating labs. Part of the participants either used high laser power for acceptor excitation to obtain reliable information on the labeling stoichiometry, acceptor lifetime and anisotropy by acquiring a maximum number of photons. Others kept the acceptor excitation power as low as possible to minimize acceptor saturation and photobleaching[6]. Regardless of which approach was used, accurate FRET efficiencies can be determined from the experimentally determined correction factors. Acceptor photobleaching results in a higher amount of donor-only molecules. In addition, acceptor saturation leads to dark states that still quench the donor via FRET and appears below the static FRET line. Acceptor blinking leads to a mixture of donor-only signal, which, in turn, results in false-positive dynamics.

**Supplementary Note 2: Discussion of γ-factor estimation.**

Obtaining a reliable $\gamma$-factor is very crucial for smFRET data analysis and proper determination of accurate FRET values. While correction for background, spectral crosstalk and direct excitation can be performed reliably, the detection efficiency correction factor γ is much more difficult to determine and the best approach depends on the sample at hand and the measurement modality (see Supplementary Note 1, Supplementary Note 18). An incorrect γ-factor introduces systematic errors in the FRET efficiency and the derived distances, especially for intermediate FRET efficiencies. The difficulties arising from the $\gamma$-factor can be illustrated with data from one lab that used sub-optimal filter combinations for the dye pair Alexa Fluor 546 (Alexa546)/Alexa Fluor 647 (Alexa647). This resulted in an inefficient detection of the red fluorescent signal and a $\gamma$-factor of 0.09 (Supplementary Table 1; lab #18). Consequently, the determined accurate FRET efficiencies were unreliable, i.e., deviating largely from the expected values, and were hence excluded for the calculation of the averages in Fig. 1. For another laboratory, the correction factors could not be determined due to lack of a functional red laser (Supplementary Table 1; lab #19). Hence, accurate FRET values could not be calculated. An incorrect $\gamma$-factor can additionally distort the dynamic shift in the $E$-$\tau$ plot resulting either in unphysically negative or artificially positive apparent dynamic shifts. The absence of negative values for the apparent dynamic shifts in Fig. 4f indicates that the five participating groups estimated the $\gamma$-factor well.

In this study, we asked all groups to use a global $\gamma$-factor for the data analysis of MalE, meaning that one common $\gamma$-factor was determined for all six data sets (Supplementary Figure 3). A global $\gamma$-factor works well when all samples have the same photophysical behavior. However, when the labeling position and/or conformation of the biomolecule induces a change of the fluorescence quantum yield of the fluorophore by local quenching or enhancement, a global $\gamma$-factor is no longer strictly appropriate. Here, we observed variations of the fluorescence lifetimes of the dyes on the order of 5-10% (Supplementary Table 11). This indicates no large variations of the fluorescence quantum yields between the different samples and justifies the global $\gamma$ correction. On the other hand, a local $\gamma$ correction can only be done when multiple conformations with different FRET efficiencies are present in a single measurement, or by using the lifetime information when the sample is known to be static. The $\gamma$-factors reported in this study for the MalE system with dye pair Alexa546-Alexa647 were generally low in the range from 0.2 to 0.6, with an average of 0.39±0.12 (Supplementary Table 1). Interestingly, the spread in FRET efficiency between the different groups can be described by an apparent uncertainty in the $\gamma$-factor of ~23% (Fig. 3e) corresponding to an absolute error of ±0.07. This suggests that the uncertainty in the determination of the $\gamma$-factor contribute significantly to the uncertainty in the overall FRET efficiency (see Supplementary Note 4).

**Supplementary Note 3: Data re-evaluation procedure for U2AF2.**

The apo population for U2AF2 is a single, dynamic population. Hence, *E*-*S* plots or *E*-τ plots are not suitable for determination of the γ-factor. In addition, there was an additional subpopulation with an unusually high stoichiometry. Therefore, we had to find another approach, discussed below, to accurately estimate the γ-factor for this system. In general, the best protocol for γ-factor determination will depend on the properties of the sample being measured. Reanalysis of the collected raw data of U2AF2 from different labs was performed using the PAM software[7] (Supplementary Table 17). First, a burst search was performed using an all photon burst search with a threshold of 50-100 photons per sliding time window of 500 µs depending on the dataset. For one set of measurements, a lower threshold of 20 photons per 500 µs time window was necessary. To remove blinking and bleaching events, an ALEX-2CDE filter with a lower limit of 5 and an upper limit of 25 was used depending on the data set. Values may differ depending on the excitation intensities and sample concentrations used for the measurements[8]. After burst selection, background subtraction and correction for crosstalk and direct excitation were performed as discussed in the data analysis section. Briefly, for background subtraction, the background signal was obtained from the buffer measurement and subtracted it from the burst signal. Crosstalk and direct excitation corrections were performed by calculating the signal in the acceptor channel for donor only and acceptor only species respectively, after donor excitation and subtracting them from the burst signal. To determine the detection correction factor, we used the approach of Lee *et al.*[9] (i.e. fitting a line to $1/S_{PR}$ vs $E_{PR}$) as the sample is dynamic and a lifetime approach is not possible. The apo configuration shows a single, dynamically averaged population so that we had to combine data from both the holo and apo measurements. We verified that there was no significant change in quantum yield of the donor and acceptor fluorophores in the absence and presence of ligand by measuring the fluorescence lifetime of the donor-only species and the acceptor lifetime with direct excitation. However, we did observe an additional subpopulation with a higher stoichiometry value (Supplementary Figure 17a). The acceptor is slightly quenched in this population, but not enough to explain the stoichiometry shift. As the origins of this population are unclear and simple explanations are insufficient to describe the observed properties, this population was not incorporated in the calculation of the γ-value. The average values of $S_{PR}$ and $E_{PR}$ for the three populations (apo, holo – low FRET, holo – high FRET) were determined from the peak values of a 2D-Gaussian fit in the ES-histograms for the respective populations. From these peak values, a straight line was fit to the three data points of $1/S_{PR}$ vs $E_{PR}$. Reanalysis of the data by a single person led to a further improvement of the consistency between laboratories (Fig. 2d-e). Part of the discrepancy came from the fact that the individual labs used a global γ approach but did not compensate for the presence of the second population. Further reasons for the discrepancies in the measured data from different laboratories arise from the fact that the dynamics and RNA binding are temperature dependent (and temperature was not specified) and that the RNA concentration used in the holo experiments was insufficient to saturate protein binding leading to a mixture of apo and holo proteins in the "holo" measurements.

**Supplementary Note 4: Error propagation of the uncertainty of the $\gamma$-factor on the FRET efficiency $E$.**

To support the hypothesis that the spread of the reported FRET efficiency $E$ values for the different MalE mutants is caused by inaccuracies of the experimental calibration, we derive below how the uncertainty of the detection efficiency correction factor $\gamma$ propagates into the uncertainty of the measured FRET efficiency. We also investigated the propagation of error for all other correction factors (donor leakage, acceptor direct excitation, and background in the donor and acceptor channels) to show that only the propagated uncertainty of the $\gamma$-factor follows the parabolic trend of the observed uncertainty in Fig. 3e. Lastly, we discuss the relative contributions of the uncertainties of the different correction factors.

**I. Error propagation of the $\gamma$-factor on $E$**

To support the hypothesis that the spread of the reported FRET efficiency $E$ values for the different MalE mutants is caused by inaccuracies of the experimental calibration, we derive how the uncertainty of the detection efficiency correction factor $\gamma$ propagates into the uncertainty of the measured FRET efficiency. Using the nomenclature introduced by Hellenkamp et al.[10] , the apparent FRET efficiency before $\gamma$-factor correction ${}^{\text{iii}}E_{\text{app}}$ is given by:

$$ {}^{\text{iii}}E_{\text{app}} = \frac{F_{A|D}}{{}^{\text{ii}}I_{\text{D}_{\text{em}}|\text{D}_{\text{ex}}} + F_{A|D}}, \tag{4.1} $$

where ${}^{\text{ii}}I_{\text{D}_{\text{em}}|\text{D}_{\text{ex}}}$ is the background-corrected donor intensity after donor excitation, and $F_{A|D}$ is the cross-talk and direct excitation corrected acceptor fluorescence after donor excitation, given by:

$$ F_{A|D} = {}^{\text{ii}}I_{\text{A}_{\text{em}}|\text{D}_{\text{ex}}} - \alpha\,{}^{\text{ii}}I_{\text{D}_{\text{em}}|\text{D}_{\text{ex}}} - \delta\,{}^{\text{ii}}I_{\text{A}_{\text{em}}|\text{A}_{\text{ex}}}. \tag{4.2} $$

Here, ${}^{ii}I_{\text{A}_{\text{em}}|\text{D}_{\text{ex}}}$ is the background-corrected acceptor intensity after donor excitation, ${}^{\text{ii}}I_{\text{A}_{\text{em}}|\text{A}_{\text{ex}}}$ is the background-corrected acceptor intensity after acceptor excitation, and $\alpha$ and $\delta$ are the correction factor for donor crosstalk and acceptor direct excitation respectively.

The fully corrected FRET efficiency $E$ is then given by:

$$ E = \frac{F_{A|D}}{\gamma\,{}^{\text{ii}}I_{\text{D}_{\text{em}}|\text{D}_{\text{ex}}} + F_{A|D}} = \left(\gamma\left(\frac{1}{{}^{\text{iii}}E_{\text{app}}} - 1\right) + 1\right)^{-1} \tag{4.3} $$

Using standard error propagation, the uncertainty of the FRET efficiency $E$ due to the uncertainty of the $\gamma$-factor is given by:

$$ \Delta E = \left|\frac{\partial E}{\partial \gamma}\right| \Delta\gamma, \tag{4.4} $$

where $\Delta\gamma$ is the uncertainty in $\gamma$, and $|\partial E/\partial \gamma|$ is the partial derivative of $E$ with respect to $\gamma$. The partial derivative is given by:

$$\left|\frac{\partial E}{\partial \gamma}\right| = \frac{E(1-E)}{\gamma}$$

Thus, the propagated uncertainty is given by:

$$\Delta E = E(1-E)\frac{\Delta\gamma}{\gamma} \tag{4.5}$$

This equation was fit to the experimental data in Fig. 3e of the main text, yielding an estimated uncertainty of the $\gamma$-factor calibration of $\Delta\gamma/\gamma = 0.23$.

To estimate the absolute (rather than the relative) error in $\gamma$ from the data, we calculated $\Delta\gamma$ directly from the reported FRET efficiency $E$ and $\gamma$-factor for each lab by rearranging Eq. 4.5:

$$\Delta\gamma = \gamma\frac{\Delta E}{E(1-E)} \tag{4.6}$$

From the obtained values for $\Delta\gamma$ for the different labs, we obtain an average value of $\Delta\gamma$ = 0.071 ± 0.051 (mean ± standard deviation).

**II. Error propagation of the remaining correction factors on *E***

Hellenkamp et al.[10] have provided explicit expressions for the propagation of all relevant error sources into the derived FRET-averaged distance $R_{DA}^{\langle E\rangle}$. Analogous expressions have been provided by Peulen et al.[1] Here, we compare the uncertainty of the FRET efficiency $\Delta E$, which relates to the uncertainty of the distance as:

$$\Delta E = \left|\frac{\partial E}{\partial R_{DA}^{\langle E\rangle}}\right| \Delta R_{DA}^{\langle E\rangle}, \tag{4.7}$$

where $\Delta R_{DA}^{\langle E\rangle}$ is the distance uncertainty and $\left|\frac{\partial E}{\partial R_{DA}^{\langle E\rangle}}\right|$ is the absolute value of the partial derivative with:

$$E = \left(1 + \left(\frac{R_{DA}^{\langle E\rangle}}{R_0}\right)^6\right)^{-1} \tag{4.8}$$

and:

$$\left|\frac{\partial E}{\partial R_{DA}^{\langle E\rangle}}\right| = \frac{6E(1-E)}{R_{DA}^{\langle E\rangle}}. \tag{4.9}$$

For the propagated uncertainty of the correction factors for donor leakage ($\alpha$) and acceptor direct excitation ($\delta$) on $E$, we obtain according to Hellenkamp et al.[10]:

$$\Delta E_\alpha = \frac{(1-E)^2}{\gamma}\Delta\alpha, \tag{4.10}$$

$$\Delta E_\delta = (1-E)\beta\Delta\delta, \tag{4.11}$$

where β is the correction factor for the different excitation flux of the donor and acceptor fluorophores. For the contributions of constant background signal in the donor or acceptor channel (bgD and bgA, respectively), we obtain according to Hellenkamp et al.[10]:

$$\Delta E_{\mathrm{bgD}} = [\gamma E + \alpha(1-E)]\frac{\Delta I_D^{\mathrm{BG}}}{\langle F \rangle}, \tag{4.12}$$

$$\Delta E_{\mathrm{bgA}} = (1-E)\frac{\Delta I_A^{\mathrm{BG}}}{\langle F \rangle}, \tag{4.13}$$

where $\Delta I_D^{\mathrm{BG}}$ and $\Delta I_A^{\mathrm{BG}}$ are the uncertainties of the estimated background signal in the donor and acceptor detection channels and $\langle F \rangle$ is the average sum of the corrected donor and acceptor fluorescence collected during a single-molecule event.

The total uncertainty of the FRET efficiency is then given by:

$$\Delta E = \sqrt{(\Delta E_\alpha)^2 + \left(\Delta E_\gamma\right)^2 + (\Delta E_\delta)^2 + \left(\Delta E_{\mathrm{bgD}}\right)^2 + \left(\Delta E_{\mathrm{bgA}}\right)^2} \tag{4.14}$$

**III. Identification of the uncertainty of the $\gamma$-factor as the dominant error source**

Our assumption that the uncertainty of the $\gamma$-factor is the dominant source of calibration error is based on the following experimental, methodical, and theoretical arguments.

*a) Inaccuracies of other correction factors propagate into the determined $\gamma$-factor*

The corrections applied here and in Hellenkamp et al.[10] proceed through a step-wise workflow of background subtraction and correction for donor crosstalk and acceptor direct excitation, after which the correction factors $\gamma$ and $\beta$ are estimated from the apparent FRET efficiency and stoichiometry (see Online Methods). Any inaccuracies of the background count rates and correction factors $\alpha$ and $\delta$ must thus propagate to the correction factor $\gamma$. Hence, we argue that the $\gamma$-factor effectively consolidates the uncertainties of all correction factors and serves as an adequate measure for the total calibration uncertainty.

*b) The $\gamma$-factor adds the largest uncertainty to E over the probed distance range*

To compare the relative contributions of the different correction factors to the total uncertainty $\Delta E$, we assess the expected propagated errors according to Eq. 4.5 and 4.10-14. For the absolute values of the correction factors $\alpha$, $\beta$, $\gamma$ and $\delta$, we use the average values of the participating labs reported for the MalE system (Fig. 3d and Supplementary Table 1). For the relative uncertainties, we follow the estimates given in Hellenkamp et al.[10] of $\Delta\alpha/\alpha = 0.1$, $\Delta\gamma/\gamma = 0.1$ and $\Delta\delta/\delta = 0.1$. This results in the following values for the correction factors and their uncertainties: $\alpha = 0.048 \pm 0.005$, $\beta = 1.6$ (no uncertainty needed for propagation), $\gamma = 0.38 \pm 0.04$, and $\delta = 0.12 \pm 0.01$.

The uncertainty of the background signals $\Delta I_D^{\mathrm{BG}}$ and $\Delta I_A^{\mathrm{BG}}$ has previously been estimated to be of the order of ~1 photon per burst with a mean number of fluorescence photons per burst of $\langle F \rangle = 50$.[10] Here, we argue that the error due to background signal was previously overestimated. We found that an average of 90 ± 40 photons were detected by the participants over a typical burst duration of 1.7 ± 0.9 ms. Background count rates can generally be quantified from buffer measurements or even extracted from burst measurements[4,11,12] with a high accuracy and reproducibility of $\Delta I^{\mathrm{BG}}/I^{\mathrm{BG}} \leq 0.1$. Given typical background count rates of 1-2 kHz, we expect an average of 2-3 background photons per burst with an absolute uncertainty of 0.2-0.3. Therefore, we assume $\Delta I_D^{\mathrm{BG}} = \Delta I_A^{\mathrm{BG}} = 0.25$ and $\langle F \rangle = 90$, leading to a lower value for the factor $\Delta I^{\mathrm{BG}}/\langle F \rangle$ in Eq. 4.12-13 of 0.003 compared to the value used in Hellenkamp et al.[10] of $\Delta I^{\mathrm{BG}}/\langle F \rangle = 0.02$. When the background is not constant but fluctuates, the uncertainty in the background will increase.

Based on the estimated correction factor values and uncertainties, we plot the propagated error of the different calibration factors in Supplementary Fig. SN4.1. Over the range of mean FRET efficiency values probed in this study for the MalE system (i.e. between 0.49-0.92), clearly the uncertainty caused by the $\gamma$-factor provides the largest contribution to the total calibration uncertainty. At FRET efficiency values below 0.25, the contribution of the correction factors for direct excitation ($\delta$) and crosstalk ($\alpha$) become significant. Note that we assumed a lower contribution of the background signal to the uncertainty compared to Hellenkamp et al.[10] as discussed above.

*c) The observed trend of $\Delta E$ can only be described by an uncertainty in $\gamma$*

The dependence of the observed uncertainty of the FRET efficiency $\Delta E$ on the FRET efficiency, $E$, clearly follows a parabolic trend (Fig. 3e). Based on the error propagation of the different correction factors, only the $\gamma$-factor follows a parabolic shape of the form $\Delta E_\gamma \propto E(1-E)$ (Eq. 4.5), while the remaining propagated uncertainties show either a monotonic decreasing or increasing behavior (Eq. 4.10-13). This supports the notion that the uncertainty of $E$ is dominated by a calibration uncertainty of $\gamma$.

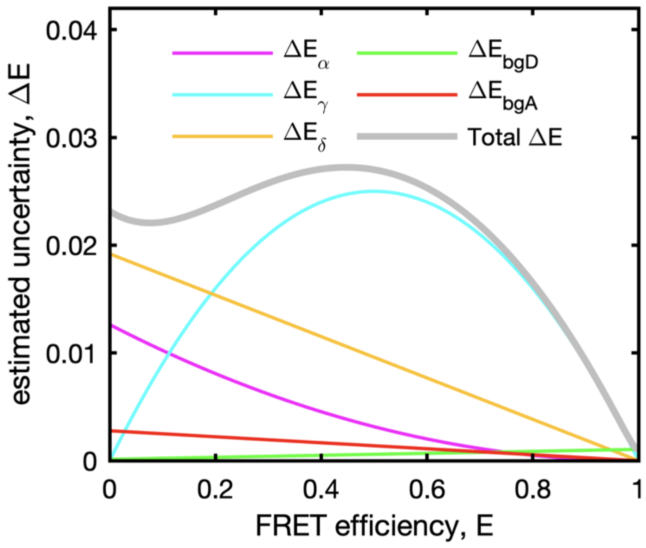

**Supplementary Figure SN4.1. Error propagation of calibration uncertainties on the FRET efficiency, $\Delta E$.** The contributions of the uncertainty in different correction factors to the total uncertainty are given as colored lines ($\Delta E_\alpha$, crosstalk; $\Delta E_\delta$, direct excitation; $\Delta E_\gamma$, $\gamma$-factor; and $\Delta E_{\mathrm{bgD}}$ and $\Delta E_{\mathrm{bgA}}$, background in donor and acceptor channels). The total uncertainty, $\Delta E$, is given as a gray line, evaluated according to Eq. 4.14. The following values and uncertainties were used for the correction factors: $\alpha = 0.048 \pm 0.005$, $\beta = 1.6$, $\gamma = 0.38 \pm 0.04$, $\delta = 0.12 \pm 0.01$, and $\Delta I^{\mathrm{BG}}/\langle F \rangle = 0.003$.

**Supplementary Note 5: Estimation of the experimental dynamic shift.**

In BVA, the standard deviation of the apparent FRET efficiency, $\sigma_{E_{\text{app}}}$, is plotted against the apparent FRET efficiency, $E_{\text{app}}$ (Fig. 4a). For the $E$-$\tau$ plot, usually the accurate FRET efficiency $E$ is plotted against the intensity-averaged donor fluorescence lifetime $\langle\tau_{D(A)}\rangle_F$ (Fig. 4b). For the estimation of the dynamic shift, $\langle\tau_{D(A)}\rangle_F$ is normalized with respect to the lifetime of the donor in the absence of the acceptor, $\tau_{D(0)}$, to constrain both axes to the interval of [0,1].

To determine the peak position of the single-molecule population, we fit the distribution of single molecule events in the $E$-$\langle\tau_{D(A)}\rangle_F/\tau_{D(0)}$ or BVA histograms using a superposition of $M$ two-dimensional Gaussian distributions. Each Gaussian distribution is described by an amplitude $A_i$, a vector of the central coordinates $\boldsymbol{\mu_i}$, and a covariance matrix $\boldsymbol{\Sigma_i}$:

$$f(\boldsymbol{x}) = \sum_{i=1}^{M} A_i \cdot (2\pi)^{-1}|\boldsymbol{\Sigma_i}|^{-\frac{1}{2}}\exp\left(-\frac{1}{2}(\boldsymbol{x}-\boldsymbol{\mu_i})\boldsymbol{\Sigma_i}^{-1}(\boldsymbol{x}-\boldsymbol{\mu_i})'\right), \quad (5.1)$$

where $\boldsymbol{\mu_i}$ is a row vector of length 2, and $\boldsymbol{\Sigma}$ is a *2*-by-*2* symmetric matrix whose diagonal elements and non-diagonal elements are the variances and covariances, respectively. Depending on whether the analysis is applied to the $E$-$\tau$ or BVA plot, the coordinate row vector $\boldsymbol{x}$ is given by $\boldsymbol{x} = \left(\langle\tau_{D(A)}\rangle_F/\tau_{D(0)}, E\right)$ or $\boldsymbol{x} = \left(E_{\text{app}}, \sigma_{E_{\text{app}}}\right)$, respectively. The model parameters are determined using a maximum likelihood estimation. The required number of populations needed to adequately describe the data is determined visually. When more than one bivariate normal distributions ($M > 1$) is needed to describe the data, the position of the component with the highest amplitude is taken. After the position of the major population is determined, we estimate the dynamic shift as described below.

For BVA, the dynamic shift is defined as the distance of the population center along the vertical ($\sigma_{BVA}$) axis from the static FRET-line, given by:

$$\sigma_{E_{\text{app}}} = \sqrt{\frac{E_{\text{app}}\left(1-E_{\text{app}}\right)}{n}}, \quad (5.2)$$

where $n$ is the number of photons per window used to estimate $\sigma_{E_{\text{app}}}$ (here, $n = 5$). To provide consistency with the procedure proposed by Torella et al.[13], we compute the dynamic shift for the main population based on the average value of $\sigma_{E_{\text{app}}}$ over a FRET efficiency interval given by its estimated mean and width, $[\mu_{E_{\text{app}}} - w_{E_{\text{app}}}, \mu_{E_{\text{app}}} + w_{E_{\text{app}}}]$, where $\mu_{E_{\text{app}}}$ and $w_{E_{\text{app}}}$ are the center position and width of the main population in the $\left(E_{\text{app}}, \sigma_{E_{\text{app}}}\right)$ plot.

For the $E$-$\tau$ plot, the dynamic shift is defined as the minimum distance from the peak of the population to the static FRET line, given by:

$$E = 1 - \frac{\langle\tau_{D(A)}\rangle_F}{\tau_{D(0)}}. \quad (5.3)$$

Generally, a linker correction for the static FRET-line is performed[14]. As a result, the static FRET-line is slightly curved. The dynamic shift is thus not necessarily the orthogonal distance to the static FRET line and was determined numerically. Lastly, the dynamic shift is assigned a positive or negative sign depending on whether the population lies above (positive) or below (negative) the static FRET line.

**Supplementary Note 6: Prediction of the expected dynamic shifts.**

Analytical expressions were derived to predict the maximum expected dynamic shift for a dynamic exchange between two conformational states with FRET efficiencies $E^{(1)}$ and $E^{(2)}$ in the $E$-$\tau$ and BVA plots.

##### I. The dynamic shift in the E-τ plot

For the $E$-$\tau$ plot, i.e. a plot of the intensity averaged FRET efficiency $E$ against the normalized intensity averaged donor fluorescence lifetime $\langle\tau_{D(A)}\rangle_F/\tau_{D(0)}$, the maximum dynamic shift is given[15] :

$$ds_{(E-\tau)} = \frac{1}{\sqrt{2}}\left(\sqrt{1-E^{(1)}} - \sqrt{1-E^{(2)}}\right)^2. \tag{6.1}$$

This equation is derived by considering the maximum deviation between the ideal static FRET-line, given by:

$$E_{(E-\tau)}^{(\mathrm{stat})} = 1 - \frac{\langle\tau_{D(A)}\rangle_F}{\tau_{D(0)}}, \tag{6.2}$$

where $\langle\tau_{D(A)}\rangle_F$ and $\tau_{D(0)}$ are the intensity-weighted average donor fluorescence lifetimes in the presence and absence of the acceptor, respectively. Binary dynamic exchange between two limiting conformational states with FRET efficiencies $E^{(1)}$ and $E^{(2)}$ and the corresponding donor fluorescence lifetimes $\tau_{D(A)}^{(1)}$ and $\tau_{D(A)}^{(2)}$ is described by the dynamic FRET-line for exchange between two states[14]:

$$E_{(E-\tau)}^{(\mathrm{dyn})} = 1 - \frac{\tau_{D(A)}^{(1)}\tau_{D(A)}^{(2)}}{\tau_{D(0)}\left(\tau_{D(A)}^{(1)} + \tau_{D(A)}^{(2)} - \langle\tau_{D(A)}\rangle_F\right)}. \tag{6.3}$$

Using the relations:

$$\tau_{D(A)}^{(i)} = \tau_{D(0)}\left(1-E^{(i)}\right);\ \ i = 1,2 \tag{6.4}$$

we can then write Eq. 6.3 as a function of the FRET efficiencies:

$$E_{(E-\tau)}^{(\mathrm{dyn})} = 1 - \frac{\left(1-E^{(1)}\right)\left(1-E^{(2)}\right)}{\left(2 - E^{(1)} - E^{(2)} - \frac{\langle\tau_{D(A)}\rangle_F}{\tau_{D(0)}}\right)} \tag{6.5}$$

The difference between the static and dynamic FRET-lines along the FRET efficiency axis, $\Delta_E$, as a function of $\langle\tau_{D(A)}\rangle_F$ is then given by:

$$\Delta_E = E_{(E-\tau)}^{(\mathrm{dyn})} - E_{(E-\tau)}^{(\mathrm{stat})} \tag{6.6}$$

$$\Delta_E(\langle\tau_{D(A)}\rangle_F) = \frac{\langle\tau_{D(A)}\rangle_F}{\tau_{D(0)}} - \frac{\left(1-E^{(1)}\right)\left(1-E^{(2)}\right)}{\left(2-E^{(1)}-E^{(2)}-\frac{\langle\tau_{D(A)}\rangle_F}{\tau_{D(0)}}\right)} \tag{6.7}$$

The function $\Delta_E(\langle\tau_{D(A)}\rangle_F)$ is unimodal and the maximum depends on the FRET efficiencies of the limiting states, $E^{(1)}$ and $E^{(2)}$:

$$\Delta_{E,max} = \left(\sqrt{1-E^{(1)}} - \sqrt{1-E^{(2)}}\right)^2. \tag{6.8}$$

We define the *dynamic shift*, $ds_{(E-\tau)}$, as the maximum deviation of the dynamic FRET-line measured orthogonal to the static FRET-line (see Fig. 4b), which introduces the factor of $1/\sqrt{2}$:

$$ds_{(E-\tau)} \stackrel{\text{def}}{=} \frac{\Delta_{E,max}}{\sqrt{2}} = \frac{1}{\sqrt{2}}\left(\sqrt{1-E^{(1)}} - \sqrt{1-E^{(2)}}\right)^2. \tag{6.9}$$

#### II. The dynamic shift in BVA

For the BVA plot, the maximum dynamic shift is defined as the maximum distance of the dynamic FRET-line from the static FRET-line along the y-axis, i.e., the estimated standard deviation of the apparent FRET efficiency, $\sigma_{E_{\text{app}}}$ (see Fig. 4a). The static FRET-line in BVA, describing the shot noise variance, is given by[13] :

$$\sigma_{E_{\text{app}}}^{(\text{stat})} = \sqrt{\frac{E_{\text{app}}(1-E_{\text{app}})}{n}}, \tag{6.10}$$

where $n$ is the photon averaging window used to estimate the standard deviation. For the dynamic FRET-line in BVA describing the exchange between two limiting conformational states with apparent FRET efficiencies $E_{\text{app}}^{(1)}$ and $E_{\text{app}}^{(2)}$, we need to consider the excess variance caused by the conformational exchange. The contributions of shot noise, described by Eqn. 6.10, and the conformational exchange are not additive. The variance of the signal does not depend on the time dependence of the fluctuations. Thus, it is not important to know at what time points transitions between the limiting states occurred. It is only important what fraction of time the molecule spent in each individual state. For a two-state system, the expected variance in the presence of shot-noise and conformational dynamics is given by:

$$\begin{aligned}\text{Var}^{(\text{dyn})}(E_{\text{app}}) &= f_1\left[\frac{E_{\text{app}}^{(1)}\left(1-E_{\text{app}}^{(1)}\right)}{n} + E_{\text{app}}^{(1)\,2}\right] \\ &+ f_2\left[\frac{E_{\text{app}}^{(2)}\left(1-E_{\text{app}}^{(2)}\right)}{n} + E_{\text{app}}^{(2)\,2}\right] - \left(f_1 E_{\text{app}}^{(1)} + f_2 E_{\text{app}}^{(2)}\right)^2,\end{aligned} \tag{6.11}$$

where $f_1$ and $f_2$ are the fraction of time spent in the respective state with $f_1 + f_2 = 1$, and the apparent FRET efficiency is given by $E_{\text{app}} = f_1 E_{\text{app}}^{(1)} + f_2 E_{\text{app}}^{(2)}$. The standard deviation of the apparent FRET efficiency for a dynamic exchange is then given by:

$$\sigma_{E_{\text{app}}}^{(\text{dyn})} = \sqrt{\text{Var}^{(\text{dyn})}(E_{\text{app}})}. \tag{6.12}$$

The dynamic FRET-line in BVA is then obtained by varying the fraction of time spent in the limiting states, $f_1 \in [0,1]$.

To derive Eq. 6.11, we start by expressing the variance as the difference between the expected value of the squared apparent FRET efficiency $\text{E}[E_{\text{app}}^2]$ and the square of the expected value $\text{E}[E_{\text{app}}]^2$, that is:

$$\text{Var}(E_{\text{app}}) = \text{E}[E_{\text{app}}^2] - \text{E}[E_{\text{app}}]^2. \tag{6.13}$$

Here, $\text{E}[X^m]$ is the expected value of a continuous random variable $X^m$ with $m \in \{1,2\}$, defined by:

$$\text{E}[X^m] = \int X^m P(X)\, dX. \tag{6.14}$$

In BVA, the number of photons per sampling window, $n$, is constant. The probability to observe an apparent FRET efficiency, $E_{\text{app}}$ , is thus equal to the probability of observing $n_A = nE_{\text{app}}$ acceptor photons among the $n$ detected photons, which is given by a binomial distribution:

$$P(E_{\text{app}}) = P\left(n_A = nE_{\text{app}} \middle| E_{\text{app}}^{(i)}, n\right) = \binom{n}{n_A} {E_{\text{app}}^{(i)}}^{n_A} \left(1 - E_{\text{app}}^{(i)}\right)^{n-n_A}, \tag{6.15}$$

where $n_A$ is the number of acceptor photons and $E_{\text{app}}^{(i)}$ is the ideal apparent FRET efficiency of the molecule in state $i$.

We assume that the conformational dynamics are slow compared to the sampling frequency of the standard deviation, which is defined by the average time needed to detect $n$ photons. Then, in the presence of conformational dynamics between two states with apparent FRET efficiencies $E_{\text{app}}^{(1)}$ and $E_{\text{app}}^{(2)}$, the probability to observe a given average apparent FRET efficiency $E_{\text{app}}$ in a sampling window of $n$ photons is given by the weighted average of the binomial distributions for the two limiting states:

$$P(E_{\text{app}}) = f_1 P\left(n_A = n\, E_{\text{app}} \middle| E_{\text{app}}^{(1)}, n\right) + f_2 P\left(n_A = n\, E_{\text{app}} \middle| E_{\text{app}}^{(2)}, n\right), \tag{6.16}$$

where $f_1$ and $f_2$ are the probabilities that the molecule is in state 1 or 2, respectively (with $f_1 + f_2 = 1$). This follows because the molecule is found exclusively in one of the two limiting states during each sampling window of $n$ photons.

We can now calculate the expected value of the apparent FRET efficiency $E_{\text{app}}$ as:

$$\mathrm{E}\left[E_{\mathrm{app}}\right] = f_1 E_{\mathrm{app}}^{(1)} + f_2 E_{\mathrm{app}}^{(2)}. \tag{6.17}$$

The expected value of the squared apparent FRET efficiency, $\mathrm{E}\left[E_{\mathrm{app}}^2\right]$, is given by:

$$\begin{aligned} \mathrm{E}\left[E_{\mathrm{app}}^2\right] &= \int E_{\mathrm{app}}^2 P\left(E_{\mathrm{app}}\right) dE_{\mathrm{app}} \\ &= f_1 \int E_{\mathrm{app}}^2 P\left(n_A = n\, E_{\mathrm{app}} \middle| E_{\mathrm{app}}^{(1)}, n\right) dE_{\mathrm{app}} \\ &+ f_2 \int E_{\mathrm{app}}^2 P\left(n_A = n\, E_{\mathrm{app}} \middle| E_{\mathrm{app}}^{(2)}, n\right) dE_{\mathrm{app}} \end{aligned} \tag{7.18}$$

The integrals represent the expected value of the square, $\mathrm{E}[X^2]$, i.e., the second moments, for a binomial random variable. Using the definition of the variance as given in Eq. 6.13, $\mathrm{E}[X^2]$ can be calculated as:

$$\mathrm{E}[X^2] = \mathrm{Var}(X) + \mathrm{E}[X]^2 = np(1-p) + n^2p^2, \tag{6.19}$$

where $n$ and $p$ are the attempt number (i.e. number of photons in the averaging window, $n$) and success probability of the binomial process ($p = E_{\mathrm{app}}^{(i)}$), and $X$ is the number of successes (here, $X = n_A = nE_{\mathrm{app}}$ ). The apparent FRET efficiency is given by $E_{\mathrm{app}} = n_A/n$. Thus, for state $i$, the expected value of the squared apparent FRET efficiency, $\mathrm{E}^{(i)}\left[E_{\mathrm{app}}^2\right]$, is given by:

$$\mathrm{E}^{(i)}\left[E_{\mathrm{app}}^2\right] = \frac{E_{\mathrm{app}}^{(i)}\left(1 - E_{\mathrm{app}}^{(i)}\right)}{n} + E_{\mathrm{app}}^{(i)}. \tag{6.20}$$

Finally, the variance of $E_{\mathrm{app}}$ is given by:

$$\begin{aligned} \mathrm{Var}^{(\mathrm{dyn})}\left(E_{\mathrm{app}}\right) &= \mathrm{E}\left[E_{\mathrm{app}}^2\right] - \mathrm{E}\left[E_{\mathrm{app}}\right]^2 \\ &= f_1 \left[\frac{E_{\mathrm{app}}^{(1)}\left(1 - E_{\mathrm{app}}^{(1)}\right)}{n} + {E_{\mathrm{app}}^{(1)}}^2\right] \\ &+ f_2 \left[\frac{E_{\mathrm{app}}^{(2)}\left(1 - E_{\mathrm{app}}^{(2)}\right)}{n} + {E_{\mathrm{app}}^{(2)}}^2\right] - \left(f_1 E_{\mathrm{app}}^{(1)} + f_2 E_{\mathrm{app}}^{(2)}\right)^2, \end{aligned} \tag{6.21}$$

where the fractions of the limiting states are defined by the observed average apparent FRET efficiency $E_{\mathrm{app}} = f_1 E_{\mathrm{app}}^{(1)} + f_2 E_{\mathrm{app}}^{(2)}$. This is Eqn. 6.11. The fractions are given by:

$$f_1 = \frac{E_{\mathrm{app}} - E_{\mathrm{app}}^{(2)}}{E_{\mathrm{app}}^{(1)} - E_{\mathrm{app}}^{(2)}}; \quad f_2 = 1 - f_1. \tag{6.22}$$

The dynamic shift in BVA is then defined as the maximum difference between the static and dynamic FRET-lines:

$$ds_{(BVA)} = \max_{E_{\mathrm{app}}} \left[\sigma_{E_{\mathrm{app}}}^{(\mathrm{dyn})}\left(E_{\mathrm{app}}\right) - \sigma_{E_{\mathrm{app}}}^{(\mathrm{stat})}\left(E_{\mathrm{app}}\right)\right]. \tag{6.23}$$

Since no analytical expression was found for Eq. 6.23, it was evaluated numerically.

#### Supplementary Note 7: $R_0$ determination for the Alexa546-Alexa647 dye pair for MalE.

The Förster radius, $R_0$, is given by[16]

$$R_0{}^6 = \frac{9\, ln\,(10)}{128\, \pi^5\, N_A} \frac{\kappa^2}{n_{im}} Q_D \frac{\int_0^\infty F_D(\lambda)\varepsilon_A(\lambda)\,\lambda^4\, d\lambda}{\int_0^\infty F_D(\lambda)\, d\lambda}. \quad (7.1)$$

where $\kappa^2$ is the orientation factor, $Q_D$ the quantum yield of the donor, $F_D(\lambda)$ is the fluorescence emission spectrum of the donor, $\varepsilon_A(\lambda)$ is the absorption spectrum of the acceptor (scaled with the appropriate absorption coefficient), $N_A$ is Avogadro's number and $n_{im}$ is the index of refraction in the intervening medium. For an appropriate $R_0$ determination, it is necessary to determine the fluorescence quantum yield of the donor labeled to the molecule of interest as well as the overlap integral.

##### 7.1 Fluorescence quantum yield $\Phi_{F,D}$

The fluorescence quantum yield of the donor dye Alexa546 covalently bound to the protein was determined using Rhodamine 6G as a reference.

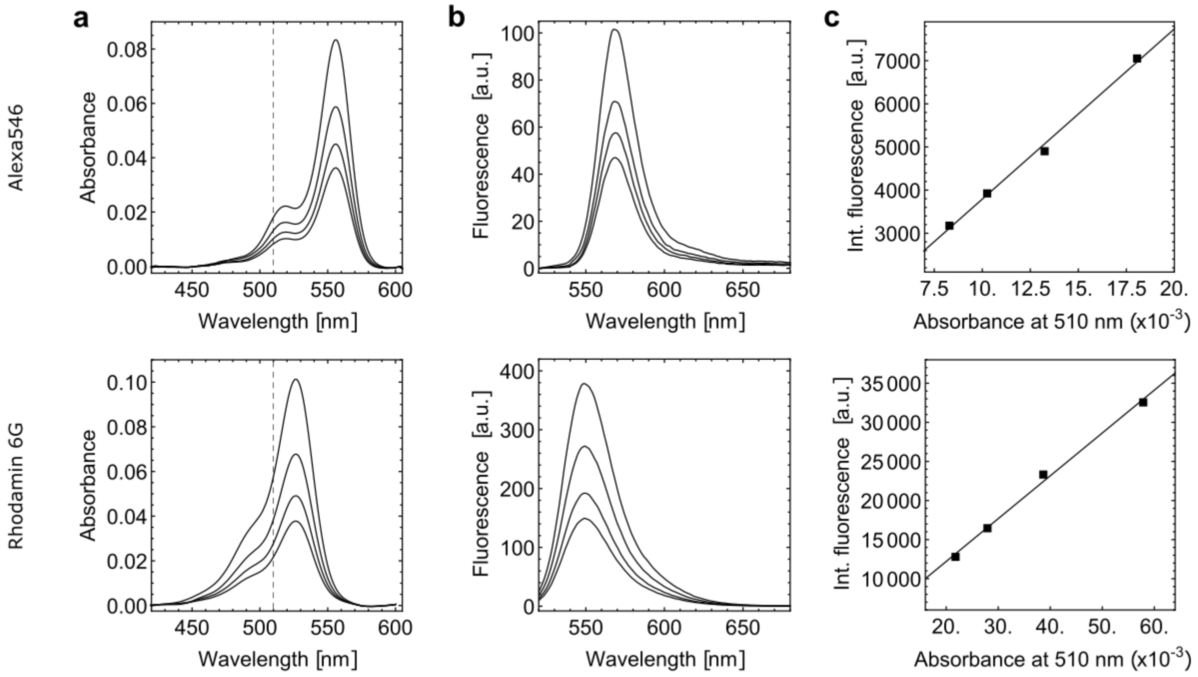

**Supplementary Figure SN7.1. Exemplarily fluorescence quantum yield determination for MalE-1. (a)** The absorption spectrum of MalE-1 labeled with Alexa546 (top) and Rhodamine 6G (bottom) at 0.5, 0.625, 0.75 and 1 µM concentration. **(b)** The emission spectrum of MalE-1 labeled with Alexa546 (top) and Rhodamine 6G (bottom) at 0.5, 0.625, 0.75 and 1 µM concentration excited at 510 nm. **(c)** The integrated fluorescence of MalE-1 labeled with Alexa546 (top) and Rhodamine 6G (bottom) plotted against absorbance at 510 nm at 0.5, 0.625, 0.75 and 1 µM concentration and linear fit to the data points (line). Assuming a fluorescence quantum yield of 91% for Rhodamine 6G, the quantum yield for MalE-1 labeled with Alexa546 was determined using the ratio of the slopes.

The absorption (Supplementary Figure SN7.1a) and emission spectra (Supplementary Figure SN7.1b) at 510 nm excitation wavelength were measured for four different dye concentrations in the range of 0.5-1.0 µM for the sub-stoichiometrically labeled donor only samples and compared to Rhodamine 6G in water. A linear fit to the integrated fluorescence intensity $\int_0^\infty F_D(\lambda)\,d\lambda$ for 510 nm excitation as a function of concentration provides two slopes: $m_{Al546}$ and $m_{R6G}$ (Supplementary Figure SN7.1c).

The fluorescence quantum yield of the donor Alexa546 is calculated from the slopes of the linear fit as

$$\boldsymbol{\Phi}_{F,D} = \frac{m_{Al546}}{m_{R6G}} \boldsymbol{\Phi}_{F,R6G} \tag{7.2}$$

where $\boldsymbol{\Phi}_{F,R6G} = 91 \pm 2\%$ is taken from literature[17]. The quantum efficiency of Alexa546 bound to the protein was found to be $72 \pm 4\%$. This is similar to the quantum yield of Alexa546 alone in PBS given in the molecular probes handbook of 0.79[18].

#### 7.2 Overlap integral J

The overlap integral was calculated using the emission spectrum $F_D(\lambda)$ of the donor only sample and a normalized absorption spectrum of the acceptor $\bar{\varepsilon}_A$ scaled to the literature extinction coefficient $\varepsilon_A(\lambda) = \varepsilon_{A_{max}}\, \bar{\varepsilon}_A$ using:

$$J = \frac{\int_0^\infty F_D(\lambda)\varepsilon_A(\lambda)\,\lambda^4\,d\lambda}{\int_0^\infty F_D(\lambda)\,d\lambda}. \tag{7.3}$$

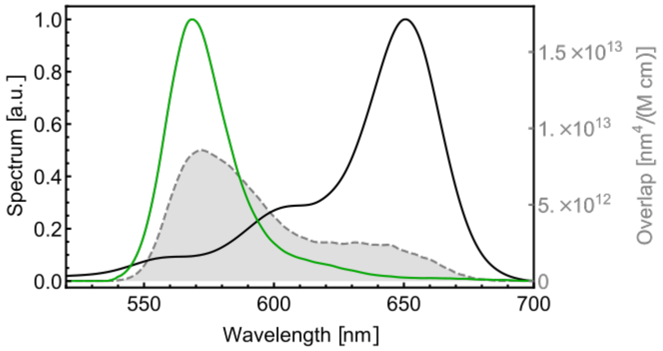

**Supplementary Figure SN7.2. The overlap integral of Alexa546-Alexa647.** The normalized emission spectrum of Alexa546 (green) and absorption spectrum of Alexa647 (black) plotted along with the overlap integral shown in gray for the MalE-1 mutant.

The overlap integral is illustrated in Supplementary Figure SN7.2 (gray area) resulting from the donor emission (green) and acceptor absorption (black) spectra.

#### 7.3 Förster radius calculation

For calculating $R_0$, we have used the following values:

Orientation factor $\kappa^2$: 2/3
Averaged refractive index $n_{im}$ (ref. [19]): 1.4
Extinction coefficient at maximum $\varepsilon_{A_{max}}$ (ref. [19]): 265,000 $OD/(M\,cm)$
Fluorescence quantum yield $\boldsymbol{\Phi}_{F,D}$: $72 \pm 4\%$
Overlap integral $J$: $7.0 \pm 0.1 \times 10^{15}\, nm^4/(M\,cm)$

(assuming that the given extinction coefficient is precise) and determined to be $R_0 = 65 \pm 3$ Å considering uncertainties in $\kappa^2$ of 10% and in $n_{im}$ of 5%. This differs significantly from the manufacture's published $R_0$ of 74 Å[20].

The absorption and emission spectra were measured using singly labeled donor and acceptor mutants for MalE-1, MalE-2 and MalE-3. For other dye combinations used in this study, the following Förster radii were used: $R_0$ (Alexa546-Abberior STAR635P) = 62.5 Å, $R_0$ (Atto532-Atto643) = 59.0 Å and $R_0$ (Alexa488-Alexa647) = 52.0 Å.

**Supplementary Note 8: Estimation of the residual anisotropies.**

To improve the robustness of the analysis, the residual anisotropies were determined via two approaches, from the steady-state intensities ("*ss*") and from the time-resolved data ("*tr*"). The values reported in Fig. 5e-f of the main text correspond to an average value of the two estimates.

*a) Residual anisotropies from steady-state intensities*

Using single-molecule multiparameter fluorescence detection (MFD)[5], steady-state anisotropies, $r_{ss}$, in each detected single-molecule event were calculated from the background-corrected photon counts registered in the parallel and perpendicular detection channels as described in detail in reference [21]. Next, we plotted the obtained $r_{ss}$ versus the intensity averaged fluorescence lifetime $\langle\tau\rangle_F$ to analyze their relation by the Perrin equation[22,23]. Here, we assume that the anisotropy decay can be described by two depolarization components[24]: (i) the global rotation of the protein, $\rho_{\text{global}}$, of approx. 20 ns, and (ii) a second component of ~ 0.5 ns describing the local dye motions, $\rho_{\text{linker}}$, with amplitudes $x_{\text{global}} = 1 - x_{\text{linker}}$ and $x_{\text{linker}}$, respectively:

$$r_{ss} = r_0 \left( \frac{x_{\text{linker}}}{1 + \frac{\langle\tau\rangle_F}{\rho_{\text{linker}}}} + \frac{1 - x_{\text{linker}}}{1 + \frac{\langle\tau\rangle_F}{\rho_{\text{global}}}} \right). \tag{8.1}$$

The residual anisotropy is defined as the amplitude of the global rotational motion as:

$$r_{\infty,ss} = r_0(1 - x_{\text{linker}}), \tag{8.2}$$

where $x_{\text{linker}}$ is given by:

$$x_{\text{linker}} = \left( \frac{r_{ss}}{r_0} - \frac{1}{1+\frac{\langle\tau\rangle_F}{\rho_{\text{global}}}} \right) \Bigg/ \left( \frac{1}{1+\frac{\langle\tau\rangle_F}{\rho_{\text{linker}}}} - \frac{1}{1+\frac{\langle\tau\rangle_F}{\rho_{\text{global}}}} \right), \tag{8.3}$$

and the fundamental anisotropy $r_0$ is assumed to be 0.38 for all dyes used here.

*b) Residual anisotropies from the time-resolved anisotropy analysis*

For the single-cysteine variants, the time-resolved anisotropy decay, $r(t)$, was obtained from ensemble TCSPC experiments. For some variants where single-cysteine mutants were not available, $r(t)$ was obtained from the single-molecule MFD experiments using a sub-ensemble selection of double-labeled molecules by applying an adequate value of the ALEX-2CDE filter. When needed, the bursts are additionally filtered using the stoichiometry value. From the selected double-labeled molecules, the fluorescence decays were obtained for the donor emission channel after donor excitation ($D_{ex}|D_{em}$) and the acceptor emission channel after acceptor excitation ($A_{ex}|A_{em}$). Note that, using this selection, the donor fluorescence lifetime is shortened by FRET to the acceptor. The anisotropy information was obtained from a global fit of the fluorescence decays $F_{VV}(t)$, $F_{VH}(t)$ and $F_{VM}(t)$, where $VV$ and $VH$ denote vertically (V) and horizontally (H) polarized emitted light upon vertical excitation (V), respectively and $VM$ represents the fluorescence lifetime decay measured at the magic angle (~54.7°) of the emission polarizer after vertical excitation. Here, the $VM$ decay was not measured but rather constructed by mixing the $VV$ and $VH$ decays and with the experimentally determined $g$-factor and correcting for polarization mixing in the beam path of the high NA objective using the factors $l_1$ and $l_2$[21,25] as follows:

$$F_{VM}(t) = (1 - 3l_2)F_{VV}(t) + g_{VH/VV}(2 - 3l_1)F_{VH}(t) \tag{8.4}$$

By including the magic angle decay $F_{VM}(t)$ in a global analysis, the stability of the fit is significantly improved as it increases the robustness of the fluorescence lifetime estimation.

The experimental decay curves $F(t)$ are described based on the idealized model decays $f(t)$. The relation between time-resolved anisotropy and polarization resolved fluorescence decays $f(t)$ is given as follows[25] :

$$f_{VV}(t) = \frac{1}{3}f_{VM}(t)\big(1 + (2 - 3l_1)r(t)\big); \tag{8.5}$$

$$f_{VH}(t) = \frac{1}{3}f_{VM}(t)\big(1 - (1 - 3l_2)r(t)\big), \tag{8.6}$$

The magic angle decay, $f_{VM}(t)$, is modelled by a multi-exponential function:

$$f_{VM}(t) = F_0 \sum_i x_i \exp\left(-\frac{t}{\tau_i}\right), \tag{8.7}$$

where $\tau_i$ denotes the fluorescence lifetimes and $x_i$ are the fraction of molecules with the corresponding lifetime, and $F_0$ is a scaling factor that normalizes the integral of the model decay to the integrated experimental signal (i.e., the total measured counts). Similarly, the time-resolved anisotropy is modelled as:

$$r(t) = r_0\left(x_{\text{dye}}\exp\left(-\frac{t}{\rho_{\text{dye}}}\right) + x_{\text{linker}}\exp\left(-\frac{t}{\rho_{\text{linker}}}\right) + x_{\text{global}}\exp\left(-\frac{t}{\rho_{\text{global}}}\right)\right), \tag{9.8}$$

where $\rho_j$ is the depolarization time of species $j$ with anisotropy amplitude $b_j = r_0 x_j$. As above, we consider the global rotation of the molecule on timescales longer than 10 ns ($\rho_{\text{global}} >$ 10 ns) and the local mobility of the tethered label including the linker on the nanosecond scale, which is accounted for using two components, one for the linker movement ($\rho_{\text{linker}}$ ~ 1-5 ns) and the other for fast rotation of the dye ($\rho_{\text{dye}}$ ~ 0.3-0.5 ns). Due to the slow, global rotation, this equation becomes equivalent to the cone-in-cone model[26]. The contribution of a specific process to the overall depolarization of the fluorescence signal is determined by the species fraction $x_j$, with $\sum_j x_j = 1$. The fundamental anisotropy is taken to be $r_0 = 0.38$ for all dyes. The contribution of global rotation of the macromolecule to the total depolarization of emitted light represents the residual anisotropy, $r_{\infty,tr}$, given by $r_{\infty,tr} = r_0 x_{\text{global}}$.

The model decays are then convolved with the corresponding polarized component of the instrument response function (IRF):

$$F_{VV}(t) = f_{VV}(t) \otimes IRF_{VV}(t) + sc_{VV} IRF_{VV}(t) + B_{VV}, \tag{8.9}$$

$$F_{VH}(t) = g_{VH/VV}\, f_{VH}(t) \otimes IRF_{VH}(t) + sc_{VH} IRF_{VH}(t) + B_{VH}, \tag{8.10}$$

$$F_{VM}(t) = f_{VM}(t) \otimes IRF_{VM}(t) + sc_{VM} IRF_{VM}(t) + B_{VM}, \tag{8.11}$$

Here, $B_{VV}$, $B_{VH}$ and $B_{VM}$ account for constant background signal, the factors $sc_{VV}$, $sc_{VH}$ and $sc_{VM}$ describe the contribution of scattered laser light, and the factor $g_{VH/VV}$ corrects for the different detection efficiencies in the two detection channels $VV$ and $VH$ and $\otimes$ denotes a circular convolution. Here we apply a circular convolution because the full microtime range is used for the analysis, resulting in a periodic signal. Note that the idealize curves, $f(t)$, have been normalized to the total measured number of counts in Eqs. 8.5 and 8.6 via Eq. 8.7. Prior to the convolution, the IRF is corrected for uncorrelated background signal, e.g., due to the detector dark counts, and a time shift of the IRF is applied to correct for shifts that arise due to count rate differences between the IRF and fluorescence decay measurements. Similar to the $F_{VM}(t)$ decay, its corresponding $IRF_{VM}(t)$ was constructed by mixing $IRF_{VV}(t)$ and $IRF_{VH}(t)$ according to Eq. 8.4.

The quality of the fit was judged using the reduced chi-squared, $\chi_r^2$. Analysis of the time-resolved anisotropy was done using the software package ChiSurf, a GUI-based suite composed of a collection of Python scripts for various types of fluorescence data analysis, available at https://github.com/Fluorescence-Tools/chisurf.

##### c) *Computation of the average residual and combined anisotropies from steady-state and time-resolved measurements*

For the steady-state anisotropy, assumptions regarding the rotational times are needed, which improves the robustness of this calculation. Time-resolved anisotropy requires no assumptions, but results in higher uncertainties. Hence, to increase the overall robustness of the analysis, we take the average of the residual anisotropies from the two approaches to compute the average residual anisotropy, $\langle r_\infty \rangle_{tr,ss}$:

$$\langle r_\infty \rangle_{tr,ss} = \frac{r_{\infty,tr} + r_{\infty,ss}}{2} \tag{8.12}$$

The uncertainty $\Delta\langle r_\infty \rangle_{tr,ss}$ was obtained as the standard deviation of the two estimates, corresponding to:

$$\Delta\langle r_\infty \rangle_{tr,ss} = \frac{\left| r_{\infty,tr} - r_{\infty,ss} \right|}{\sqrt{2}} \tag{8.13}$$

The combined residual anisotropy was then computed as a geometric average:

$$\langle r_{c,\infty} \rangle_{tr,ss} = \sqrt{\langle r_{\infty,D} \rangle_{tr,ss} \cdot \langle r_{\infty,A} \rangle_{tr,ss}}, \tag{8.14}$$

and the corresponding error was obtained using standard error propagation as:

$$\Delta\left(\langle r_{c,\infty} \rangle_{tr,ss}\right) = \frac{1}{2}\sqrt{\frac{\langle r_{\infty,D} \rangle_{tr,ss}}{\langle r_{\infty,A} \rangle_{tr,ss}} (\Delta\langle r_{\infty,A} \rangle_{tr,ss})^2 + \frac{\langle r_{\infty,A} \rangle_{tr,ss}}{\langle r_{\infty,D} \rangle_{tr,ss}} (\Delta\langle r_{\infty,D} \rangle_{tr,ss})^2}. \tag{8.15}$$

**Supplementary Note 9: Calculation of distance uncertainties according to the "diffusion with traps" model.**

Anisotropy measurements of fluorophores tethered to the surface of biomolecules indicate that the dye motion is commonly not isotropic despite the use of long, flexible linkers. Estimating a precise value of the orientational factor $\kappa^2$ is generally impossible as it would require knowledge of the mutual orientations of the transition dipole moments of the donor and acceptor molecules and their orientation with respect to the inter-dye distance vector. However, one can restrict the possible $\kappa^2$ values to a range that is compatible with the experiments with the help of the experimental anisotropy data. Previous approaches have estimated the uncertainty in $\kappa^2$ from the residual anisotropy in terms of rotational restrictions[27,28]. Here, we adopt a different approach termed the *diffusion with traps* (DWT) model[29] that provides an uncertainty in the measured donor-acceptor separation due to the unknown orientation factor, $\kappa^2$, based on the experimentally obtained fractions of trapped dye species and FRET efficiencies. The fraction of trapped dye species is obtained from the fluorescence anisotropy decays of the donor and acceptor fluorophores as a ratio of the residual, $r_\infty$, to the fundamental anisotropy, $r_0$, which is often referred to as the second rank order parameter, $S^{(2)}_{A/D}$:

$$S^{(2)}_{A/D} = \frac{r_{\infty,A/D}}{r_{0,A/D}}, \tag{9.1}$$

where the subscript denotes that the order parameter is computed either for the donor ($D$) or acceptor ($A$) fluorophore.

Inspired by observations from MD simulations, the DWT model assumes that the fluorophores are either completely free, i.e., undergoing isotropic reorientation, or remain immobile for an extended period of time, meaning that they are trapped. It is also assumed that the exchange between these two species is slower than the donor fluorescence lifetime (i.e., static on the timescale of FRET) but faster than the diffusion time through the confocal volume in smFRET experiments of ~1-5 ms. This approximation is supported from MD data where the residence times of these states are found to be on the order of 10-100 ns or longer.

The DWT model assumes a mixture of trapped and free dye species and considers all four mobility scenarios for the donor and acceptor when computing the FRET efficiency, that is:

(I) $D_{\text{free}} - A_{\text{free}}$
(II) $D_{\text{free}} - A_{\text{trapped}}$
(III) $D_{\text{trapped}} - A_{\text{free}}$ and
(IV) $D_{\text{trapped}} - A_{\text{trapped}}$.

To calculate the range of possible $\kappa^2$ values, we generate random orientations of the dipole moment vectors, $\boldsymbol{\mu_D}$ and $\boldsymbol{\mu_A}$. For a given orientation of dipole vectors, the orientation factor $\kappa^2$ is given by[29]:

$$\kappa^2 = \frac{2}{3} + \frac{2}{3}S_D^{(2)}S_{\beta_1}^{(2)} + \frac{2}{3}S_A^{(2)}S_{\beta_2}^{(2)} + \frac{2}{3}S_D^{(2)}S_A^{(2)}\Big[S_\delta^{(2)} + 6S_{\beta_1}^{(2)}S_{\beta_2}^{(2)} + 1 + 2S_{\beta_1}^{(2)} + 2S_{\beta_2}^{(2)} - 9\cos\beta_1\cos\beta_2\cos\delta\Big] \tag{9.2}$$

where the angles $\beta_1$ and $\beta_2$ define the dipole orientations with respect to the donor-acceptor distance vector $\mathbf{R_{DA}}$ for D and A, respectively, and $\delta$ corresponds to the angle between the D and A transition dipole vectors, as illustrated below:

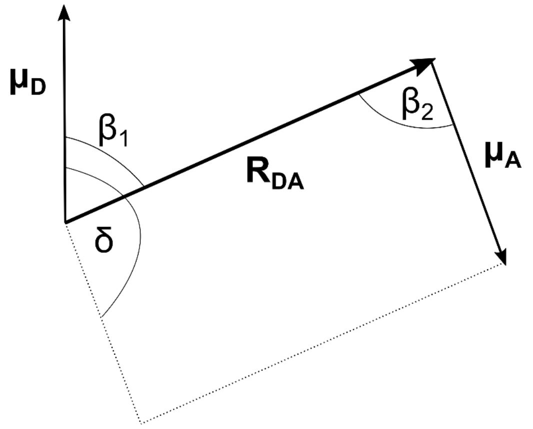

The second-rank order parameters for the angles are defined as follows:

$$S_\delta^{(2)} = \frac{1}{2}(3cos^2\delta - 1) \tag{9.3}$$

$$S_{\beta_1}^{(2)} = \frac{1}{2}(3cos^2\beta_1 - 1) \tag{9.4}$$

$$S_{\beta_2}^{(2)} = \frac{1}{2}(3cos^2\beta_2 - 1) \tag{9.5}$$

In the first step, generated are $N$ random vectors, which represent random orientations of transition dipole vectors of donor and acceptor dyes. From those vectors, the angles $\beta_1$, $\beta_2$ and $\delta$ are calculated. Since we do not use the FRET sensitized acceptor anisotropy decay, $\delta$ is allowed to vary between 0° and 90°. However, it is possible to limit the range of possible δ - and $\kappa^2$- values by using the residual FRET sensitized acceptor anisotropy, $r_{\infty,A(D)}$.

In the next step, the FRET efficiency is computed as a function of $\kappa^2$ by considering all possible scenarios (I - IV) weighted by their respective probability of occurrence based on the estimated fractions of free/trapped dyes from the order parameters:

$$E(\kappa^2) = \left(1 - S_D^{(2)}\right)\left(1 - S_A^{(2)}\right)E + S_D^{(2)}S_A^{(2)}E\left(\kappa^2(D_{\text{trapped}} - A_{\text{trapped}})\right) + S_D^{(2)}\left(1 - S_A^{(2)}\right)E\left(\kappa^2(D_{\text{trapped}} - A_{\text{free}})\right) + \left(1 - S_D^{(2)}\right)S_A^{(2)}E\left(\kappa^2(D_{\text{free}} - A_{\text{trapped}})\right) \tag{9.6}$$

where $E = E(\kappa^2(D_{\text{free}} - A_{\text{free}})$ is the experimentally measured FRET efficiency, and:

$$E\left(\kappa^2(D_{\text{trapped/free}} - A_{\text{trapped/free}})\right) = \frac{1}{1 + \frac{2/3}{\kappa^2(D_{\text{trapped/free}} - A_{\text{trapped/free}})}\left(\frac{1}{E} - 1\right)}, \tag{9.7}$$

where the factor $2/3$ represents the isotropically averaged $\kappa^2$ value. The value of $\kappa^2(D_{\text{trapped/free}} - A_{\text{trapped/free}})$ is calculated according to Eq. 9.2 using the following assignment of the order parameters:

$$\begin{aligned} &\kappa^2\left(D_{\text{trapped}} - A_{\text{trapped}}\right):\ S_D^{(2)} = S_A^{(2)} = 1 \\ &\kappa^2\left(D_{\text{trapped}} - A_{\text{free}}\right):\ S_D^{(2)} = 1, \qquad S_A^{(2)} = 0 \\ &\kappa^2\left(D_{\text{free}} - A_{\text{trapped}}\right):\ S_D^{(2)} = 0, \qquad S_A^{(2)} = 1 \end{aligned} \tag{9.8}$$

The dynamically averaged $\langle\kappa^2\rangle$ that corresponds to the obtained FRET efficiency $E(\kappa^2)$ is then obtained as:

$$\langle\kappa^2\rangle = \frac{2}{3}\frac{\frac{1}{E} - 1}{\frac{1}{E(\kappa^2)} - 1}, \tag{9.9}$$

where again the factor $2/3$ represents the isotropic average of $\kappa^2$.

This procedure is repeated for 10,000 randomly generated orientations of the dye vectors for a pre-defined distance, $R_{DA}(2/3) = R_0(1/E - 1)^{\frac{1}{6}}$, where $E$ is the measured FRET efficiency and $R_0$ is the Förster radius assuming a $\kappa^2$ of $2/3^{\text{rds}}$. The obtained dynamically-averaged $\langle\kappa^2\rangle$ values are then converted into a normalized distance $R_{\text{app}}$ according to the following equation:

$$R_{\text{app}} = \frac{R_{DA}(\langle\kappa^2\rangle)}{R_{DA}(2/3)} = \left(\frac{3}{2}\langle\kappa^2\rangle\right)^{-1/6} \tag{9.10}$$

The relative standard error in distance due to $\kappa^2$, $\Delta R_{\text{app}}(\kappa^2)$, is then defined as the standard deviation of the obtained $R_{\text{app}}$ distribution normalized to its mean (Fig. 5e). Here, we assume all orientations of the fluorophore in the trapped states are possible. A more precise determination would require a detailed knowledge of the trapped states and the orientation distribution function of the dyes in the various traps.

**Supplementary Note 10: Donor quenching estimation at different labeling positions on the MalE protein.**

To explain the observed quenching at particular locations in MalE, we used coarse-grained Brownian dynamics simulations as described previously[1]. First, we identified the positions of the amino acids that tend to quench. These are typically MET, TYR, TRP and HIS. Next, we assigned a quenching rate constant to each of them. We found that the experimental data is explained best with a quenching rate constant of $k_q$= 2 $ns^{-1}$. After the position and quenching rate of the problematic amino acids are identified, we determine all sterically allowed positions of the donor dye using Accessible Volume simulations[24]. Within the accessible volume, we simulate diffusion of a dye using Brownian dynamics. As it is known that dyes diffuse slower in the vicinity of biomolecule's surface due to the non-specific sticking interactions, a heterogeneous diffusion model was applied. In this model, the diffusion coefficient of D = 10 $Å^2\ ns^{-1}$ is decreased by a factor of 10 in the vicinity of the surface. For each dye position within the AV, we measured the distance between the dye and all $C_\beta$ atoms of the protein. Whenever the minimum distance was below the threshold of $R_{surface}$ = 8 Å, the slow diffusion coefficient (D = 1 $Å^2\ ns^{-1}$) was used. Next, for each dye position, we estimated the fluorescence properties by measuring the distance of the fluorophore to the amino acids that can quench. The quenching was approximated by a step function such that whenever the dye was found within a distance of $R_{rad}$ = 8.5 Å, quenching occurs at the given rate of $k_q$. Hence, during quenching, the emission rate of the donor is given by $k_D = \tau_0^{-1} + k_q$ and, for distances > $R_{rad}$, the donor is unquenched $k_D = \tau_0^{-1}$. Simulations were performed using the open-source GUI version of *QuEST – Quenching Estimator* software (GitHub page: https://github.com/Fluorescence-Tools/quest). Parameters used in the simulations are summarized in the Supplementary Table SN10.1 and our estimated fluorescence quantum yields for the donor at different labeling positions of MalE protein are provided in Supplementary Table SN10.2.

**Supplementary Table SN10.1.** Parameters used for the coarse-grained MD simulations of Alexa546 quenching at the different labeling positions of MalE in the apo and holo states.

<table>
<tr><th>Dye species</th><th>$\tau_0$ / ns</th><th colspan="2">Dye properties/ Å</th><th>$D_{free}$ / Å$^2$ ns$^{-1}$</th><th>$R_{surface}$ / Å</th><th>$D_{surface}$ / Å$^2$ ns$^{-1}$</th><th>$R_{rad}$ / Å</th><th>Quencher</th><th>$k_q$ /ns$^{-1}$</th><th colspan="2">BD simulations</th></tr>
<tr><td rowspan="4">Alexa Fluor 546</td><td rowspan="4">4.1</td><td rowspan="2">$L_{length}$</td><td rowspan="2">20.5</td><td rowspan="4">10</td><td rowspan="4">8</td><td rowspan="4">1</td><td rowspan="4">8.5</td><td>MET</td><td rowspan="4">2</td><td>Simulation time / µs</td><td>10</td></tr>
<tr><td>TYR</td><td>Time step / ps</td><td>2</td></tr>
<tr><td>$L_{width}$</td><td>4.5</td><td>TRP</td><td>Grid size / Å</td><td>0.5</td></tr>
<tr><td>$R_{dye}$</td><td>3.5</td><td>HIS</td><td></td><td></td></tr>
</table>

**Supplementary Table SN10.2.** Simulated fluorescence quantum yield $\phi_{F,D}$ and fraction of dye states that collided with quenching amino acids for Alexa546 dye on the MalE protein. Using the coarse-grained Brownian dynamics simulations (see used simulation parameters in Supplementary Table SN10.1), we estimated quenching of the donor dye at the labeling positions of MalE. The positions that show a prominent quenching in BD simulations are S352C (apo/holo), K34C(apo/holo) and T36C (apo/holo), as given by frequent collisions with quenching amino acids. Using species averaged fluorescence lifetime $\langle\tau\rangle_x$ obtained by fitting ensemble time-resolved lifetime measurements of single labelled cysteine mutants (Supplementary Table 11), one can determine experimental fluorescence quantum yields, according to $\phi_{F,D} = \phi_{F,D,ref} \frac{\langle\tau_{D(0)}\rangle_x}{\tau_{D(0)}}$, with $\phi_{F,D,ref}$ = 0.72 (see Supplementary Note 7) and $\tau_{D(0)}$ = 4.1 ns. As predicted by BD simulations, experiments confirm that at position S352C dye is prone to more frequent collisions with quenchers, resulting in lower fluorescence quantum yield compared to other labelling positions.

| Position | State | Fraction of collisions with quenchers / % | Simulated $\phi_{F,D}$ | Experimental $\phi_{F,D}$* |
|---|---|---|---|---|
| **K29C** | apo | 10.84 | 0.64 | 0.71 |
| | holo | 14.45 | 0.60 | 0.71 |
| **S352C** | apo | 54.77 | 0.50 | 0.66 |
| | holo | 42.73 | 0.52 | 0.67 |
| **D87C** | apo | 23.16 | 0.57 | 0.68 |
| | holo | 28.65 | 0.53 | 0.69 |
| **A186C** | apo | 20.95 | 0.62 | 0.69 |
| | holo | 20.94 | 0.61 | 0.69 |
| **A134C** | apo | 7.92 | 0.64 | 0.69 |
| | holo | 5.96 | 0.65 | 0.70 |
| **K34C** | apo | 38.62 | 0.58 | - |
| | holo | 32.88 | 0.58 | - |
| **T36C** | apo | 49.94 | 0.50 | - |
| | holo | 48.02 | 0.53 | - |
| **N205C** | apo | 28.94 | 0.52 | - |
| | holo | 25.08 | 0.55 | - |

*Ensemble lifetime measurements of the single-labeled cysteine mutants K34C, T36C and N205C are not available.

**Supplementary Note 11: Determination of statistical significance of the dynamic shifts of MalE.**

To assess the statistical significance of the excess dynamic shift obtained for the measured MalE mutants, we compared the distribution of the measured dynamic shift between the different labs to the dynamic shifts observed for dsDNA. We computed the $p$-value for each of them using a chi-square score. With the $p$-value, we test whether distributions of ds values differ between DNA samples (reference, *r*) and the measured MalE and U2AF2 samples (*s*). Our null hypothesis is that measured MalE and U2AF2 samples appear as static as DNA. The $p$-value is the probability that the data is observed given the null hypothesis and can be used to decide whether the null hypothesis is valid or should be rejected given. Small $p$-values, particularly those below 0.05 (corresponding to a 2σ confidence interval), leads to the rejection of the null hypothesis.

Assuming that the reference DNA samples and samples under the tested hypothesis (MalE and U2AF2 variants) both have normally distributed *ds* values with mean $\langle ds\rangle_r$, $\langle ds\rangle_s$ and standard error of the mean $SEM_r$, $SEM_s$ respectively, the chi-square test is then performed as follows:

$$\chi^2 = \left(\frac{\langle ds\rangle_r - \langle ds\rangle_{tot}}{SEM_r}\right)^2 + \left(\frac{\langle ds\rangle_s - \langle ds\rangle_{tot}}{SEM_s}\right)^2 \quad (11.1)$$

with $\langle ds\rangle_{tot}$ being:

$$\langle ds\rangle_{tot} = \frac{\frac{\langle ds\rangle_r}{SEM_r{}^2} + \frac{\langle ds\rangle_s}{SEM_s{}^2}}{\frac{1}{SEM_r{}^2} + \frac{1}{SEM_s{}^2}} \quad (11.2)$$

Using the probability density function of $\chi^2$ distribution:

$$f(\chi^2|N_{dof}) = \frac{1}{2^{\frac{N_{dof}}{2}}\Gamma\left(\frac{N_{dof}}{2}\right)}(\chi^2)^{\frac{N_{dof}}{2}-1}e^{-\frac{\chi^2}{2}} \quad (12.3)$$

where $N_{dof}$ is number of degrees of freedom ($N_{dof}$ $=N_{measurements} - N_{fit.params}$; here, $N_{dof} = 1$), one can determine the $p$-value, also referred to as *significance* or *certainty* that a reference distribution (DNA) does not match the sampled one (MalE, U2AF2). The p-value quantifies how unlikely it is to obtain a $\chi^2$ value that is larger than the one observed between reference and measured sample. Commonly, the $p$-value is defined as the area under the right tail of the $f(\chi^2|N_{dof})$ function:

$$p = \int_{\chi^2}^{+\infty} f(\chi^2|N_{dof})d\,\chi^2 \quad (11.4)$$

Using the described methodology in Supplementary Note 5, three labs estimated the apparent dynamic shift, *ds*, for different dye combinations of MalE and U2AF2 mutants (Supplementary

Table 8). Additionally, one lab determined *ds* values for the DNA rulers, that would serve as a reference value of what would be the observed *ds* value for static systems. Furthermore, using *ds* values reported from all three labs, we calculated *p*-values for all measured variants (Supplementary Table SN11.1). *p*-values below 0.05 are obtained for the apo state of MalE-1, MalE-5 and U2AF2, and for the holo state of MalE-1, MalE-2, MalE-4, MalE-5 and U2AF2. For those samples, the distribution of *ds* values cannot be explained with the one of reference DNA sample and the samples appear to be more dynamic compared to dsDNA.

Furthermore, *p*-values were computed using the residual dynamic shift values after filtering out dyes with pronounced sticking interactions. Significant shrinking in average *ds* value was obtained mainly for MalE-1, where the *p*-value could not be estimated due to only one point being left after dye filtering. An increase in the *p*-value can be observed for a few mutants, meaning that, after removal of sticking artifacts, they do not appear significantly different compared to a static DNA sample. This effect is particularly pronounced for the MalE-4 apo/holo sample.

**Supplementary Table SN11.1.** *p*-values for MalE and U2AF2 samples before and after filtering dye-pairs with pronounced sticking interactions. *p*-values were computed according to Eq. 11.4.

| | ***p*-value all dyes** | | ***p*-value dyes ($\Delta R_{app}(\kappa^2) \leq 10\%$)** | |
|---|---|---|---|---|
| **sample \ condition** | **apo** | **holo** | **apo** | **holo** |
| **MalE-1** | 0.0004 | 0.0024 | n.a* | n.a* |
| **MalE-2** | 0.7841 | 0.0376 | 0.7841 | 0.0376 |
| **MalE-3** | 0.9290 | 0.8973 | 0.5959 | 0.8973 |
| **MalE-4** | 0.0679 | 0.0051 | 0.4009 | 0.2024 |
| **MalE-5** | 1.20e-6 | 1.38e-6 | 0.0002 | 1.08e-5 |
| **U2AF2** | 1.11e-52 | 0.0009 | 6.91e-36 | 0.0260 |

**p*-value could not be estimated due to only one point being left after filtering dyes with pronounced sticking interactions

**Supplementary Note 12: Estimation of conformational flexibility from the residual dynamic shift for MalE.**

The dynamic shift of a population in the $E$-$\tau$ plot is defined as the minimum distance to the reference static FRET-line given by:

$$E = 1 - \frac{\langle \tau_{D(A)} \rangle_F}{\tau_{D(0)}}, \tag{12.1}$$

where $\langle \tau_{D(A)} \rangle_F$ is the intensity-weighted average donor fluorescence lifetime and $\tau_{D(0)}$ is the donor-only lifetime (see Supplementary Note 5). We use the normalized donor fluorescence lifetime $\langle \tau_{D(A)} \rangle_F / \tau_{D(0)}$ to quantify the dynamic shift as this ensures that both axes in the $E$-$\tau$ plot range from 0 to 1. We have previously derived an expression for the maximum dynamic shift of a two-state system from the static FRET-line in a plot of E against $\langle \tau_{D(A)} \rangle_F / \tau_{D(0)}$ as a function of the limiting FRET efficiencies of the two states, $E_1$ and $E_2$ (Eq. 6.1 and ref. [15]):

$$\mathrm{ds} = \frac{1}{\sqrt{2}}\left(\sqrt{1-E_1} - \sqrt{1-E_2}\right)^2 \tag{12.2}$$

Here, we use Eq. 12.2 to estimate the conformational flexibility from the measured dynamic shift under the following assumptions.

1) The dynamics are fast compared to the burst duration (< 100 µs) so that complete averaging occurs.

2) The two states are equally populated, that is the equilibrium constant of the dynamics, $K = k_{12}/k_{21}$, is equal to 1.

3) The dynamics are symmetric around the FRET-averaged mean interdye distance $R_{\langle E \rangle}$, i.e. the two states have interdye distances of $R_{\langle E \rangle} \pm \delta R$.

Under these assumptions, the dynamic population will be at a defined position off the static line that falls between the two limiting states. The dynamic shift then only depends on the amplitude of the distance fluctuation, $\delta R$:

$$\mathrm{ds}(\delta R) = \frac{1}{\sqrt{2}}\left(\sqrt{1+\left(\frac{R_0}{R_{\langle E \rangle}-\delta R}\right)^6} - \sqrt{+\left(\frac{R_0}{R_{\langle E \rangle}+\delta R}\right)^6}\right)^2. \tag{12.3}$$

We numerically solve this equation to translate the measured dynamic shift of a given dye pair into an apparent distance fluctuation given the Förster radius $R_0$ and FRET-averaged interdye distance $R_{\langle E \rangle}$. The estimated distance fluctuations $\delta R$ are given in Supplementary Table 8.

**Supplementary Note 13: Molecular dynamics (MD) simulations of MalE.**

**Setup of topology and coordinate files.** All-atom MD simulations were performed for MalE using the Amber18 suite[30] of programs with the FF14SB protein force-field[31] and the TIP4P-Ew water model[32]. For the initial structure, the crystal structure from PDB ID 1OMP was used. Using the LEaP program, the initial structure was solvated using TIP4P-Ew water molecules in an octahedral box such that the distance between the edge of water box and closest solute atom was at least 11Å. $K^+$/$Cl^-$ ions were added to neutralize the system and then additional ions were added to reach a concentration of 50 mM for both ions. To allow for a larger time step of 4 fs, Hydrogen Mass Repartitioning (HMR)[33] was applied using ParmEd in the Amber software suite.

**Minimization and Thermalization.** After the topology and coordinate files were created and solvated, energy minimization of the system was performed while the positional restraints were applied to the solute atoms. Minimization was performed in two phases: first with high and then with low force constants for the positional restraints. In the first phase, the positional restraint was applied using a harmonic potential with a force constant of 25 kcal $mol^{-1}$ $Å^{-2}$ and minimization was done through a total of 15,000 steps (5,000 steps of steepest descent minimization followed by 10,000 steps of conjugate gradient minimization). In the second phase, the force constant was reduced to 5 kcal $mol^{-1}$ $Å^{-2}$ and minimization was done in the same manner as the first phase with 5,000 steps of steepest descent minimization followed by 10,000 steps of conjugate gradient minimization. After minimization, the system was heated from 100-300 K over 100 ps using NVT-MD simulations with the force constant of 5 kcal $mol^{-1}$ $Å^{-2}$. The temperature change was applied in steps as follows. In the first 80 ps, the temperature was gradually raised from 100 K to the target 300 K. In the last 20 ps, the system was kept at the target temperature. To adjust the solvent density after increasing the temperature, heating was followed with a 300 ps of MD simulations using the NPT ensemble and the same force constant. Lastly, the force constant of the harmonic restraints was gradually reduced from 5 kcal $mol^{-1}$ $Å^{-2}$ to 1 kcal $mol^{-1}$ $Å^{-2}$ over 5×20 ps of NVT-MD simulations and then unrestrained NVT-MD production runs were performed.

**Production runs.** In total, 5 independent production replicas with a length of 2 µs each were performed. Conformations were saved every 20 ps. The SHAKE algorithm[34] was used in the production runs to constrain the bond lengths involving hydrogen atoms and long-range electrostatic interaction were treated using the Particle Mesh Ewald method[35] with a non-bonded cutoff of 8 Å.

Trajectories were analyzed using the MDTraj[36] library. The distances between the $C_\beta$ atoms of the labeling positions for the five MalE mutants were calculated (Supplementary Figure SN13.1). For mutants MalE-1, MalE-4 and MalE-5, larger $C_\beta$-$C_\beta$ distance fluctuations were observed.

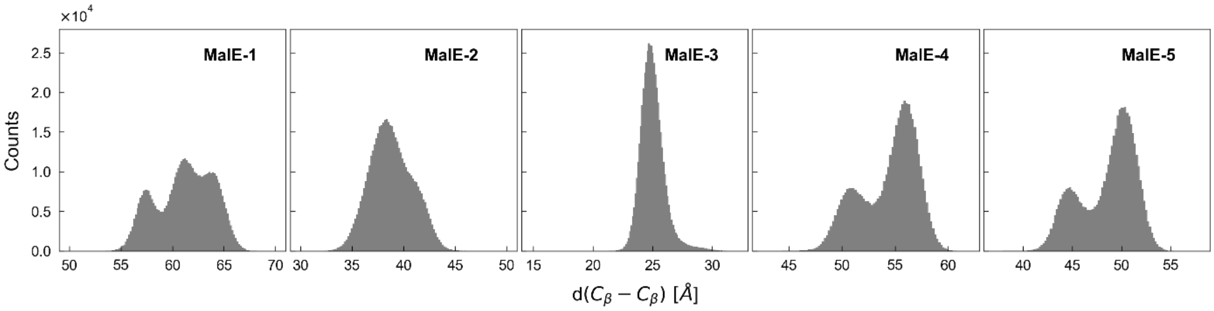

**Supplementary Figure SN13.1. Distributions of $C_\beta$-$C_\beta$ distances from MD simulations of MalE proteins in the apo state.** For each of the MalE mutants in the absence of fluorescent labels, the distance distribution between $C_\beta$ atoms of the labeling positions is shown from a total of five production runs of 2μs length each. Consistent with smFRET experiments, a higher backbone-to-backbone distance fluctuations are detected for mutants MalE-1, MalE-4 and MalE-5. The obtained mean and standard deviations are $\langle d(C_\beta - C_\beta)\rangle_{MalE-1} =$ (61.2 ± 2.7) Å, $\langle d(C_\beta - C_\beta)\rangle_{MalE-2} =$ (38.8 ± 2.1) Å, $\langle d(C_\beta - C_\beta)\rangle_{MalE-3} =$ (25.0 ± 1.0) Å, $\langle d(C_\beta - C_\beta)\rangle_{MalE-4} =$ (54.4 ± 2.7) Å and $\langle d(C_\beta - C_\beta)\rangle_{MalE-5} =$ (48.6 ± 2.8) Å.

**Supplementary Note 14: Theoretical limits for detecting dynamics.**

The theoretical description of the dynamic shifts in the $E$-$\tau$ and BVA plots as given in Supplementary Note 5 allows us to assess the detection limit for conformational dynamics with respect to the uncertainty of the experiment. To define the detectability of a given shift from the static FRET-line in an experiment, it is necessary to quantify the experimental uncertainties associated with the FRET indicators $E$ and $\langle\tau_{D(A)}\rangle_F$ for the $E$-$\tau$ plot, and $E_{\mathrm{app}}$ and $\sigma_{E_{\mathrm{app}}}^{(\mathrm{stat})}$ for the BVA plot. The experimental uncertainty contains contributions from the photon counting statistics of the experiment and systematic errors due to the calibrations or timescales of the dynamic exchange.

We first consider only the statistical error based on the standard error of the mean ($\sigma_{SEM}$) of a population, given by:

$$\sigma_{SEM} = \frac{\sigma_{SN}}{\sqrt{N_b}}, \tag{14.1}$$

where $\sigma_{SN}$ is the shot-noise broadened width of a population and $N_b$ is the number of single molecule bursts in the population. A given dynamic shift is considered detectable if it is larger than the SEM of a population on the static FRET-line.

#### I. Detection limit for dynamics in the $E$-$\tau$ plot

##### Shot-noise broadening in the $E$-$\tau$ plot

To simplify the expressions for the shot-noise broadening of a population in the $E$-$\tau$ plot, we assume that all detected single-molecule events have the same number of photons, $N_p$. For the FRET efficiency, the population width is then given by[37]:

$$\sigma_{SN}^{E} = \sqrt{\frac{E(1-E)}{N_p}}. \tag{14.2}$$

For the donor fluorescence lifetime $\tau_{D(A)}$, the variance of the estimator in the absence of background and for an instrument response with zero width ($\delta$-function) is (ref [38,39]):

$$\mathrm{Var}\left(\tau_{D(A)}\right) = \frac{1}{N_p}\tau_{D(A)}{}^2\frac{k^2}{r^2}(1-e^{-r})\left(\frac{e^{\frac{r}{k}}(1-e^{-r})}{\left(e^{\frac{r}{k}}-1\right)^2} - \frac{k^2}{e^r-1}\right)^{-1}, \tag{14.3}$$

where $k$ is the number of TCSPC detection channels and $r = T/\tau_{D(A)}$ is the ratio of the detection time window $T$ and the fluorescence lifetime $\tau_{D(A)}$. The complex expression for the variance arises because only a part of the decay is seen in the finite time window $T$ and due to the discretization of the time axis into $k$ bins. In the limit of a large time window $T$ and a high number of bins $k$, the expression simplifies to:

$$\lim_{r\to\infty,\ k\to\infty} \mathrm{Var}(\tau_{D(A)}) = \frac{1}{N_p}\tau_{D(A)}{}^2 \tag{14.4}$$

The same result is obtained by considering that the sum of independent exponential random variables $\delta t_i$ (i.e. delay times) follows the Erlang distribution with variance:

$$\mathrm{Var}\left(\sum_{i=1}^{N_p} \delta t_i\right) = N_p \tau_{D(A)}{}^2 \tag{14.5}$$

from which the variance of the lifetime estimate is obtained as:

$$\mathrm{Var}\left(\frac{1}{N_p}\sum_{i=1}^{N_p} \delta t_i\right) = \frac{1}{N_p^2}\mathrm{Var}\left(\sum_{i=1}^{N_p} \delta t_i\right) = \frac{1}{N_p}\tau_{D(A)}{}^2 \tag{14.6}$$

Thus, for the normalized donor lifetime, we obtain the shot-noise limited width of the burst-wise distribution as:

$$\sigma_{SN}^{\tau} = \frac{\tau_{D(A)}}{\sqrt{N_p}\ \tau_{D(0)}} \tag{14.7}$$

To obtain the width of the distribution along the vector of the dynamic shift, i.e., orthogonal to the static FRET-line, the two contributions are combined geometrically:

$$\sigma_{SN}^{(E-\tau)} = \frac{1}{\sqrt{2}}\sqrt{\sigma_{SN}^{E}{}^2 + \sigma_{SN}^{\tau}{}^2}, \tag{14.8}$$

The standard error of the mean of the population, determined for example by Gaussian fitting, depends on the number of single molecule events in the population $N_b$, and is given by:

$$\sigma_{SEM}^{(E-\tau)} = \frac{\sigma_{SN}^{(E-\tau)}}{\sqrt{N_b}} \tag{14.9}$$

**Effect of calibration error in the $\gamma$-factor**

An incorrect calibration of the detection efficiency correction factor $\gamma$ can lead to the detection of false-positive apparent dynamic shifts by shifting the population away from the static FRET line. However, only an underestimation of the $\gamma$-factor, leading to an overestimation of the FRET efficiency $E$, will shift the population above the static FRET-line, whereas an overestimation of $\gamma$ will result in an unphysical shift below the static FRET-line. The propagated uncertainty of the $\gamma$-factor on the FRET efficiency is derived in Supplementary Note 4 (Eq. 4.5) and is given by:

$$\sigma_{E,\gamma} = E(1-E)\frac{\Delta\gamma}{\gamma}. \tag{14.10}$$

The calibration error, $\sigma_{E,\gamma}$, is combined with the shot-noise related uncertainty $\sigma_{SN}^{(E-\tau)}$ as:

$$\sigma_{SN,\gamma}^{(E-\tau)} = \sqrt{{\sigma_{SN}^{(E-\tau)}}^2 + \sigma_{E,\gamma}^2}, \tag{14.11}$$

leading to an estimation of the combined standard error of the mean of:

$$\sigma_{SEM}^{(E-\tau)} = \frac{\sigma_{SN,\gamma}^{(E-\tau)}}{\sqrt{N_b}}. \tag{14.12}$$

#### II. Detection limit for dynamics in the BVA plot

##### Shot-noise broadening in the BVA plot

For BVA, we only have to consider the broadening along the y-axis, i.e., the uncertainty in the variance estimate. Assuming the apparent FRET efficiency, $E_{\text{app}}$, follows a Gaussian distribution with width parameter $\sigma_{E_{\text{app}}}$, the estimated standard deviation in BVA, $\tilde{\sigma}_{E_{\text{app}}}$, follows a scaled chi-squared distribution:

$$(M-1)\frac{\tilde{\sigma}_{E_{\text{app}}}^2}{\sigma_{E_{\text{app}}}^2} \sim \chi_{M-1}^2. \tag{14.13}$$

Here, $\sigma_{E_{\text{app}}} = \sqrt{\frac{E_{\text{app}}(1-E_{\text{app}})}{n}}$ and $M$ is the number of samples for the standard deviation estimate given by $M = N_p/n$, where $N_p$ is the number of photons and $n$ is the photon averaging window used for BVA. The upper $1\sigma$ confidence interval was used for the standard deviation estimate $\tilde{\sigma}_{E_{\text{app}}}$ and is given by:

$$P\left((M-1)\frac{\tilde{\sigma}_{E_{\text{app}}}^2}{\sigma_{E_{\text{app}}}^2} \leq (M-1)\frac{\tilde{\sigma}_{E_{\text{app}},\text{UL}}^2}{\sigma_{E_{\text{app}}}^2}\right) = T_M(1), \tag{14.14}$$

where $\tilde{\sigma}_{E_{\text{app}},\text{UL}}^2$ is the $1\sigma$ upper limit and $T_M(x)$ is the cumulative Student's $t$-distribution with $M$ degrees of freedom. Assuming an average of $N_p$ = 100 photons per burst and a photon averaging window of $n$ = 5, then $M$ = 20 and $T_{20}(1) \approx 0.84$. The upper limit $\tilde{\sigma}_{E_{\text{app}},\text{UL}}$ is given by:

$$\tilde{\sigma}_{E_{\text{app}},\text{UL}} = \sqrt{\frac{E_{\text{app}}(1-E_{\text{app}})}{n}}\sqrt{\frac{\chi_{\text{inv},M-1}^2\left(T_M(1)\right)}{M-1}}, \tag{14.15}$$

where $\chi_{\text{inv},M-1}^2(x)$ is the inverse cumulative chi-squared distribution with $M-1$ degrees of freedom.

The shot-noise broadening around the true value on the static FRET-line is then given by:

$$\sigma_{SN}^{(BVA)} = \tilde{\sigma}_{E_{\text{app}},\text{UL}} - \sqrt{\frac{E_{\text{app}}(1 - E_{\text{app}})}{N}}, \tag{14.16}$$

from which the standard error of the mean is obtained by dividing by the square root of the number of bursts (Eqn. 14.9).

**Kinetic averaging in BVA**

As BVA relies on the sampling of the FRET efficiency distribution from the photon time trace, it is subject to kinetic averaging during the sampling time, reducing the detectable variance. Thus, the dynamic shift is decreased for dynamics that occurs on the order of the sampling time or faster.

The sampling time, $T$, depends on the observed signal count rate, $S$, and the photon number, $n$, used to sample the FRET efficiency. The delay time between subsequent photon detection events is exponentially distributed. The time that it takes to detect $n$ photons is then described by the Erlang distribution with a shape parameter given by $n$ and the rate $S$. The average sampling time $\langle T \rangle$ is then given by:

$$\langle T \rangle = n/S. \tag{14.17}$$

For typical experiments, $S$ = 100 kHz and $n$ = 5, yielding a sampling time of $\langle T \rangle$ = 50 µs.

For BVA, it is assumed that the molecule is either in state 1 or 2 (with apparent FRET efficiencies $E_{\text{app}}^{(1)}$ or $E_{\text{app}}^{(2)}$) at each sampling point of the FRET efficiency. Here, we consider a two-state system with rates $k_{12}$ and $k_{21}$:

$$E_{\text{app}}^{(1)} \underset{k_{21}}{\overset{k_{12}}{\rightleftarrows}} E_{\text{app}}^{(2)} \tag{14.18}$$

If the dynamics are fast, there is a certain probability that the molecule interconverts during the sampling time $T$, which reduces the observed variance. The average time for the molecule to convert from any state is given by the kinetic relaxation time $\tau_r$:

$$\tau_r = \frac{1}{k_{12} + k_{21}} \tag{14.19}$$

The number of interconversions $k$ during the sampling time $T$ is Poisson distributed with the average value $\mu = T/\tau_r$:

$$P(k) = \frac{(T/\tau_r)^k}{k!} e^{-T/\tau_r}. \tag{14.20}$$

The probability to convert at least once during a time interval $T$ is then given by:

$$P_{\text{convert}} = P(k \geq 1) = 1 - P(k = 0) = 1 - e^{-\frac{T}{\tau_r}}. \tag{14.21}$$

For simplification, we assume that, when an interconversion occurred, the sampling window converges to the average apparent FRET efficiency $\langle E_{\text{app}} \rangle$. Then, any sampling of the apparent

FRET efficiency in which an interconversion occurred will have zero contribution to the variance and the dynamic contribution to the variance is simply reduced by the probability that no interconversion event occurred:

$$\mathrm{Var}'_{\mathrm{dyn}} = (1 - P_{\mathrm{convert}})\mathrm{Var}_{\mathrm{dyn}} = e^{-\frac{T}{\tau_r}}\mathrm{Var}_{\mathrm{dyn}}. \tag{14.22}$$

By mixing the shot-noise and conformational contributions to the variance of the FRET efficiency, the dynamic FRET-line for BVA in the presence of fast conformational dynamics becomes:

$$\begin{aligned}\mathrm{Var}(E_{\mathrm{app}}, P_{\mathrm{convert}}) \\ &= (1 - P_{\mathrm{convert}})\left(f_1\left[\frac{E_{\mathrm{app}}^{(1)}\left(1 - E_{\mathrm{app}}^{(1)}\right)}{n} + E_{\mathrm{app}}^{(1)\,2}\right]\right. \\ &\left.+ (1 - f_1)\left[\frac{E_{\mathrm{app}}^{(2)}\left(1 - E_{\mathrm{app}}^{(2)}\right)}{n} + E_{\mathrm{app}}^{(2)\,2}\right]\right) + P_{\mathrm{convert}}\frac{\langle E_{\mathrm{app}}\rangle(1 - \langle E_{\mathrm{app}}\rangle)}{n} \\ &- \langle E_{\mathrm{app}}\rangle^2\end{aligned}$$

where $f_1$ and $(1 - f_1)$ are the fraction of time the molecule spends in state 1 and 2, respectively.

##### III. Discussion of detection limits

Using the described formalism, we tested whether hypothetical conformational dynamics between the apo and holo states of the studied systems, MalE-1 to 5 and U2AF, could be detected in a given situation. The FRET efficiencies of the apo and holo states were obtained from AV simulations on the available crystal structures as described in the online methods (Supplementary Table 10).

We first visualized the expected dynamic shift for the $E$-$\tau$ and BVA plots as a function of the FRET efficiencies of the limiting states (Supplementary Figure SN14.1). Along the diagonal, dynamic shifts are low (<0.025) because the FRET efficiency contrast is small whereas off-diagonal combinations of FRET efficiencies result in large dynamic shifts. While the BVA plot is symmetric between low and high FRET efficiencies, the $E$-$\tau$ plot is better at resolving a dynamic exchange at high FRET efficiencies of the limiting states compared to the low FRET efficiency region. This asymmetry arises because the uncertainty in the fluorescence lifetime is largest for long fluorescence lifetimes and thus for low FRET efficiencies.

The experimentally studied MalE mutants all showed dynamic shifts below 0.03 while, for U2AF, a dynamic shift of > 0.1 is expected for both the $E$-$\tau$ and BVA plots. To put these expected shifts into perspective with respect to the experimental uncertainty, we plot the dynamic shift normalized to the expected standard error of the mean for different numbers of detected photons per burst and number of bursts in Supplementary Figure SN14.2 for the $E$-$\tau$ plot and Supplementary Figure SN14.3 for the BVA plot. Dynamics are detectable when the ratio of the dynamic shift to the experimental uncertainty, $ds/\sigma_{SEM}$, exceeds one. Clearly, the sensitivity to detect a dynamic exchange increases for both the $E$-$\tau$ and BVA plot with increasing number of photons and bursts. For typical experimental values (100 photons per burst and 1000 bursts), the dynamic exchange is predicted to be detectable as $ds/\sigma_{SEM} > 10$ for all experimental systems.

The sensitivity of the $E$-$\tau$ plot crucially depends on the accuracy of the correction factors used to compute accurate FRET efficiencies, in particular the detection efficiency correction factor $\gamma$. With higher uncertainty, $\Delta\gamma/\gamma$, the sensitivity of the $E$-$\tau$ plot decreases (Supplementary Figure SN14.4). At a relative uncertainty of $\Delta\gamma/\gamma = 0.1$, the sensitivity reduces to an extent that potential dynamics between the apo and holo states of the different MalE mutants would become undetectable. On the other hand, the large-scale dynamics for U2AF would remain detectable even at a high calibration uncertainty of $\Delta\gamma/\gamma = 0.3$. Note, the uncertainty discussed here is the uncertainty of the detection correction factor in a single lab and not the distribution of $\Delta\gamma/\gamma$ values calculated between labs in Fig. 3e.

The BVA plot loses sensitivity when the timescale of dynamics approaches the sampling window used for the estimation of the variance of the FRET efficiency distribution (Supplementary Figure SN14.5). For slow conformational dynamics with a relaxation time of $\tau_r = 10$ ms, the potential exchange between the apo and holo states is detectable with $ds/\sigma_{SEM} > 10$. As the relaxation time approaches the sampling window (here, $T \approx 50$ µs), the sensitivity is reduced significantly. At $\tau_r = 20$ µs, the exchange for most MalE mutants is on the border of the detection limit while, for $\tau_r = 10$ µs, even potential dynamics of U2AF would become undetectable in BVA.

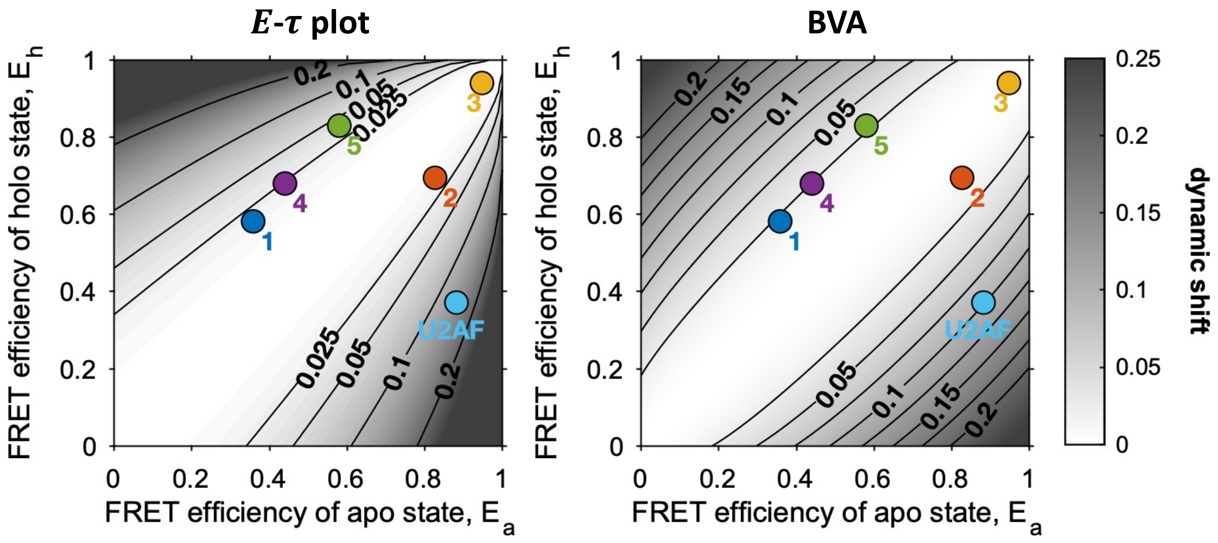

**Supplementary Figure SN14.1:** The dynamic shift in the $E$-$\tau$ (left) and the BVA (right) plots as a function of the FRET efficiencies of the limiting states (apo and holo) undergoing hypothetical dynamic exchange. The positions of the studied experimental systems are shown as colored markers (dark blue: MalE-1, red: MalE-2, yellow: MalE-3, purple: MalE-4, green: MalE-5, light blue: U2AF). The theoretical FRET efficiencies of the apo and holo states for the experimental systems were estimated from the PDB structures using AV simulations (see Online methods, Supplementary Tables 9 and 10).

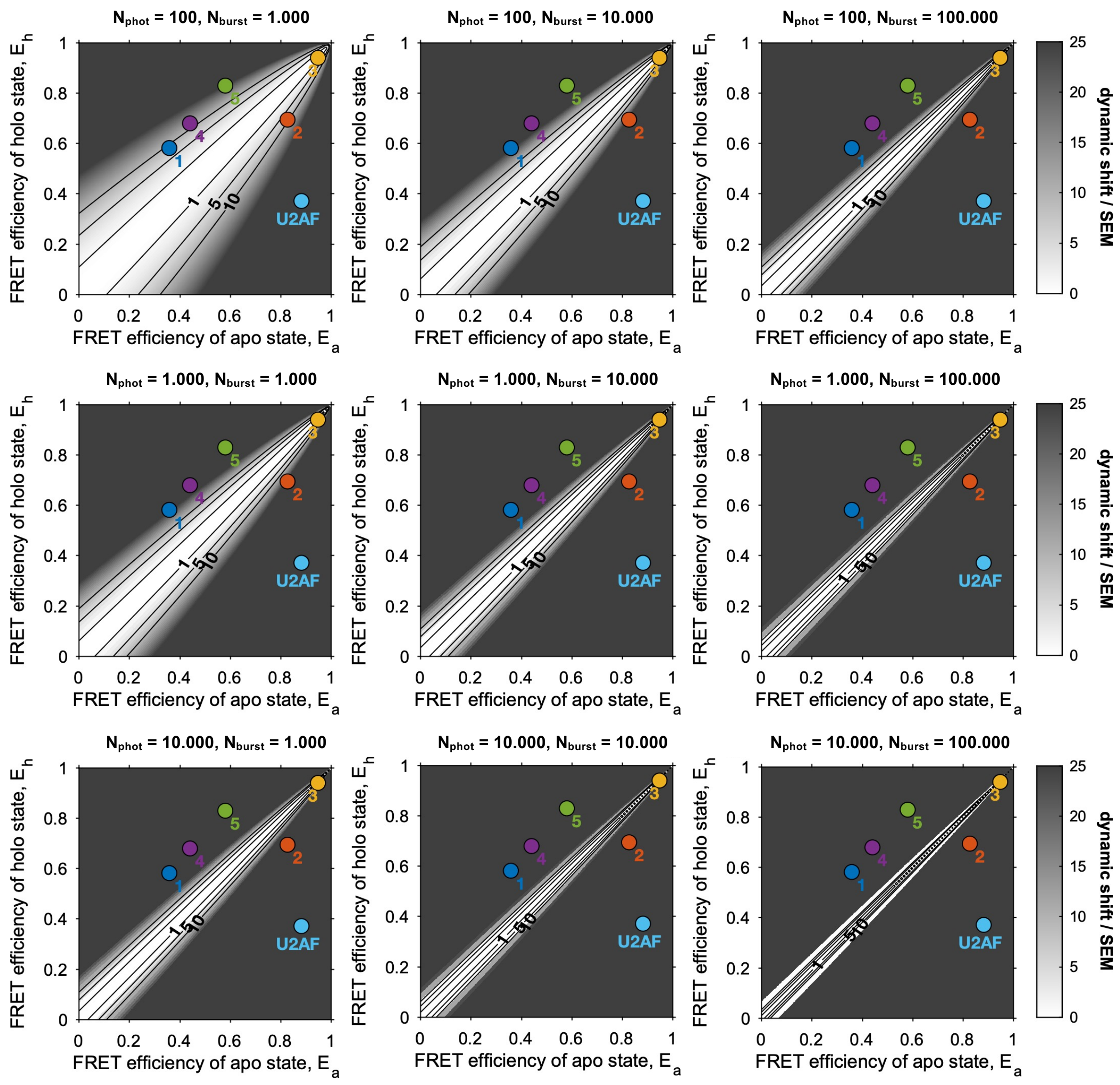

**Supplementary Figure SN14.2:** Detectability of conformational dynamics in the $E$-$\tau$ plot as a function of the photon-counting statistics. The detectability is defined as the ratio of the dynamic shift over the theoretical measurement uncertainty given by the standard error of the mean of the dynamic population. Dynamics are undetectable for ratios below one. The positions of the studied experimental systems are shown as colored markers (dark blue: MalE-1, red: MalE-2, yellow: MalE-3, purple: MalE-4, green: MalE-5, light blue: U2AF). The theoretical FRET efficiencies of the apo and holo states for the experimental systems were estimated from the PDB structures using AV simulations (see Online methods, Supplementary Tables 9 and 10).

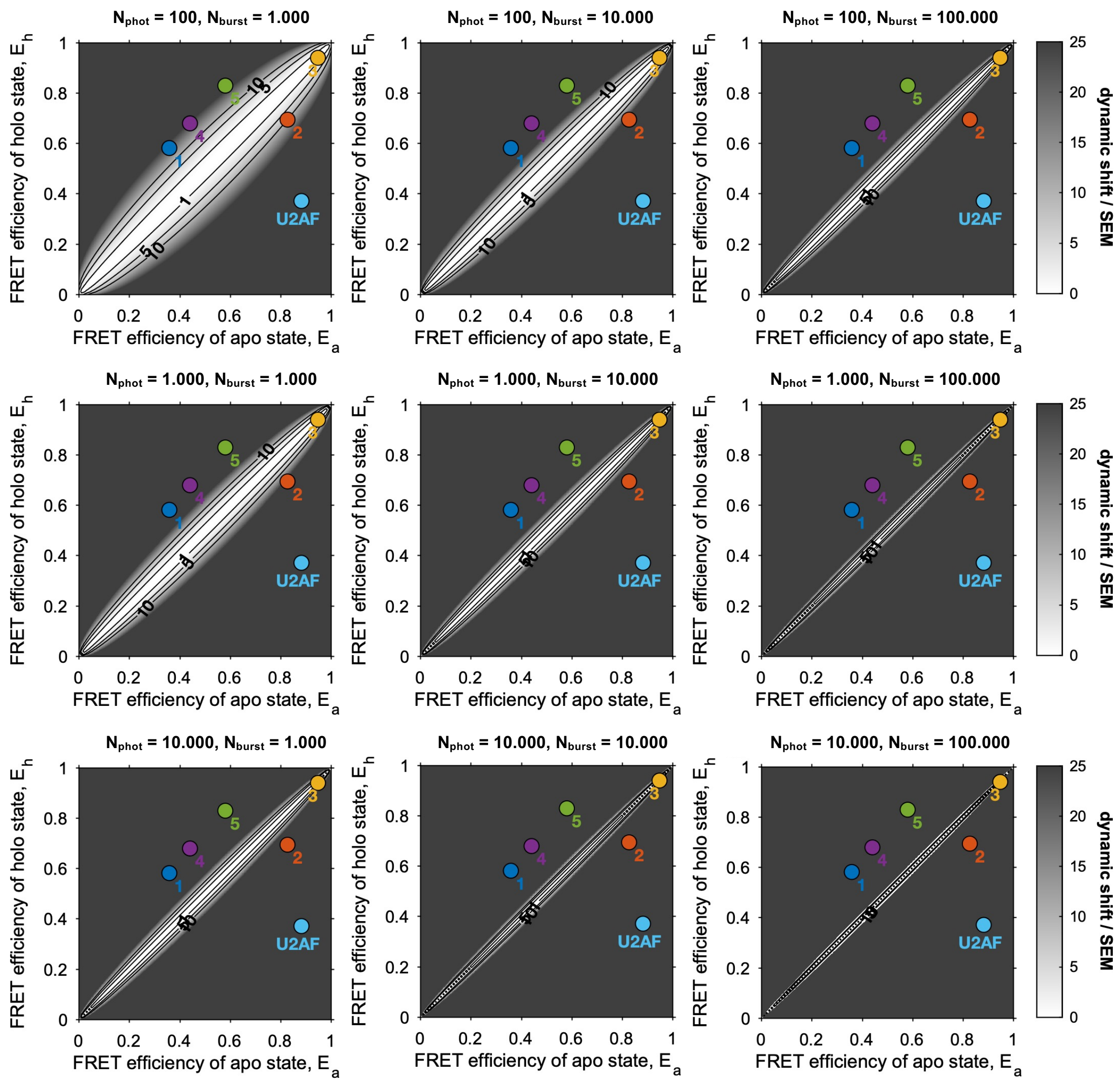

**Supplementary Figure SN14.3:** Detectability of conformational dynamics in the BVA plot as a function of the photon-counting statistics. The detectability is defined as the ratio of the dynamic shift over the theoretical measurement uncertainty given by the standard error of the mean of the dynamic population. Dynamics are undetectable for ratios below one. The positions of the studied experimental systems are shown as colored markers (dark blue: MalE-1, red: MalE-2, yellow: MalE-3, purple: MalE-4, green: MalE-5, light blue: U2AF). The theoretical FRET efficiencies of the apo and holo states for the experimental systems were estimated from the PDB structures using AV simulations (see Online methods, Supplementary Tables 9 and 10).

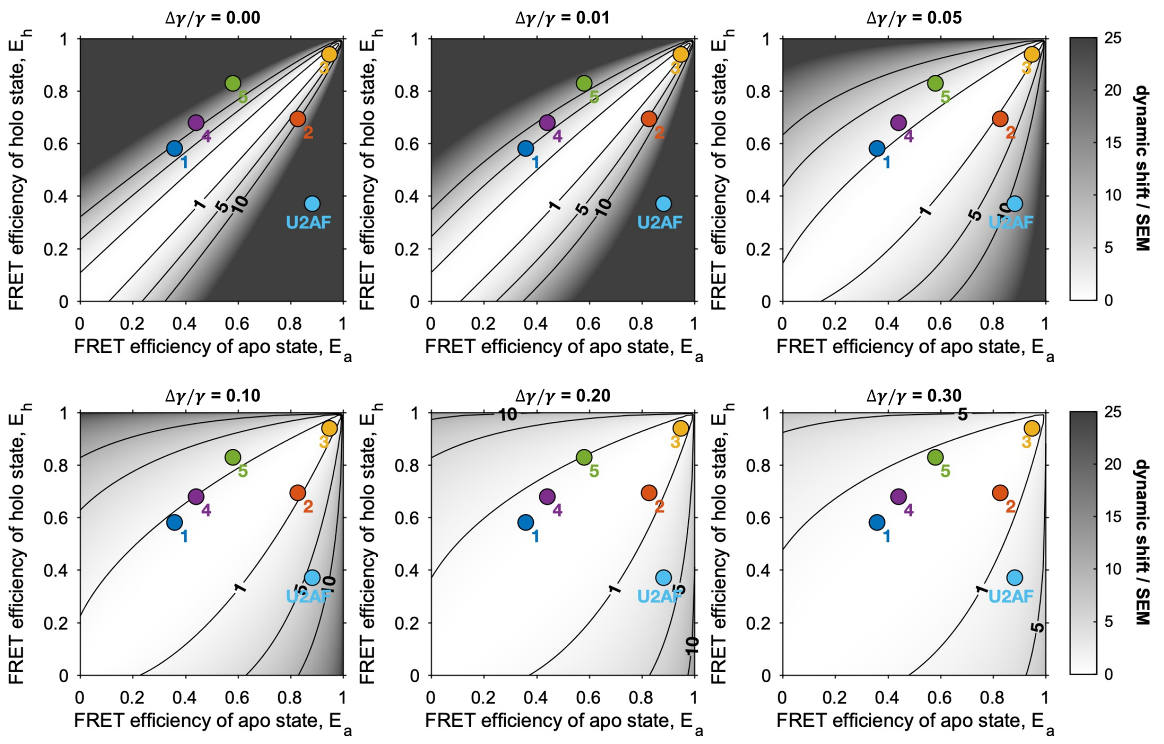

**Supplementary Figure SN14.4:** Detectability of conformational dynamics in the $E$-$\tau$ plot as a function of the calibration error of the detection efficiency correction factor $\gamma$. The relative uncertainty of the $\gamma$-factor is given by $\Delta\gamma/\gamma$. The detectability is defined as the ratio of the dynamic shift over the theoretical measurement uncertainty given by the standard error of the mean of the dynamic population. Dynamics are undetectable for ratios below one. The positions of the studied experimental systems are shown as colored markers (dark blue: MalE-1, red: MalE-2, yellow: MalE-3, purple: MalE-4, green: MalE-5, light blue: U2AF). It is assumed that the dynamic population contains 1000 bursts of 100 photons. The theoretical FRET efficiencies of the apo and holo states for the experimental systems were estimated from the PDB structures using AV simulations (see Online methods, Supplementary Tables 9 and 10).

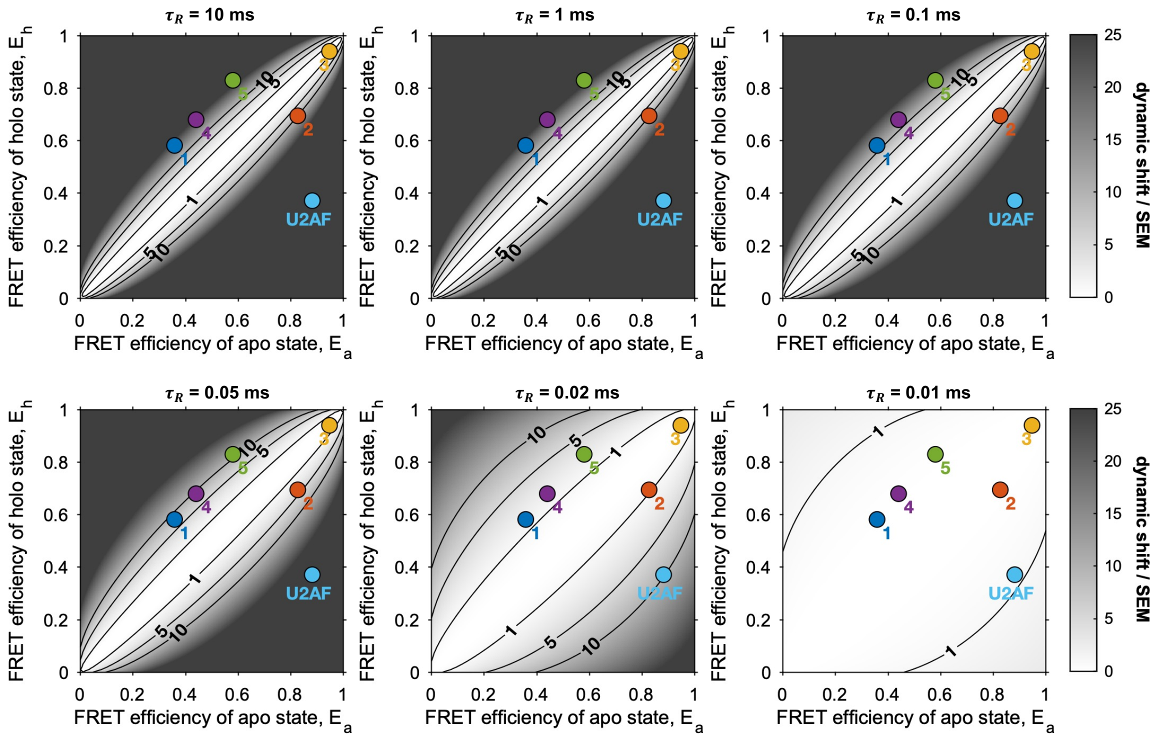

**Supplementary Figure SN14.5:** Detectability of conformational dynamics in the BVA plot as a function of the time scale of the dynamic exchange, quantified by the relaxation rate $\tau_r$. The detectability is defined as the ratio of the dynamic shift over the theoretical measurement uncertainty given by the standard error of the mean of the dynamic population. Dynamics are undetectable for ratios below one. The positions of the studied experimental systems are shown as colored markers (dark blue: MalE-1, red: MalE-2, yellow: MalE-3, purple: MalE-4, green: MalE-5, light blue: U2AF). It is assumed that the dynamic population contains 1000 bursts of 100 photons and a count rate of 100 kHz, and that an averaging window of 5 photons is used for the BVA analysis. The theoretical FRET efficiencies of the apo and holo states for the experimental systems were estimated from the PDB structures using AV simulations (see Online methods, Supplementary Tables 9 and 10).

**Supplementary Note 15: Model-free analysis of fluorescence decays for U2AF2.**

*Sub-ensemble fluorescence decays.* Sub-ensemble donor fluorescence decays were generated from single molecule experiments by selecting the double-labeled population using the ALEX-2CDE filter[8] with an upper limit of 15. An additional stoichiometry cut was applied (0.4 ≤ S ≤ 0.55) to remove the dye-related artifact at high FRET efficiency that showed a higher stoichiometry of ~0.6 (Supplementary Figure 17). For the donor-only reference decays, a stoichiometry threshold of S ≥ 0.98 was used.

*Fitting procedure.* The ideal fluorescence decays $f(t)$ were convoluted with the instrument response function and corrected for background contributions to the TCSPC pattern due to, for example, the contribution of scattered laser light, autofluorescence of the buffer, dark counts and uncorrelated background signal, to obtain the model decay $F(t)$:

$$F(t) = F_0 \cdot f(t) \otimes IRF(t - t_{\text{shift}}) + sc\, BG(t), \tag{15.1}$$

where ⊗ here denotes a linear convolution, $F_0$ is the initial amplitude of the fluorescence decay, $IRF(t)$ is the instrument response function (which is shifted by the time $t_{\text{shift}}$), $BG(t)$ is the normalized background/scatter pattern obtained from a buffer measurement and $sc$ is the background/scatter amplitude, which is estimated from the cumulative duration of all analyzed bursts $T_{\text{bursts}}$ and the background count rate $c_{BG}$ as $sc = c_{BG} T_{\text{bursts}}$. Note that, contrary to Supplementary Note 8, we apply the linear convolution operation here for the analysis of smFRET data recorded using PIE because the analysis is performed only a selected time interval of the microtime histogram (i.e., a PIE channel) to restrict the analysis to the fluorescence decay of the donor. Hence, the signal here is not periodic as it is in the ensemble TCSPC experiments.

Fits were optimized using the reduced chi-square defined by:

$$\chi^2_{\text{red}} = \frac{1}{N_{\text{data}} - N_{\text{param}}} \sum \frac{\left(F_{exp}^{(k)} - F_{model}^{(k)}\right)^2}{w_k^2}, \tag{15.2}$$

where $F_{exp}^{(k)}$ and $F_{model}^{(k)}$ are the intensities of the measured and model decay in the TCSPC bin $k$, $w_k$ is the respective weight given by $w_k = \sqrt{F_{exp}^{(k)}}$ based on the Poisson statistics of the detected signal, and $N_{\text{data}}$ and $N_{\text{param}}$ are the number of bins in the TCSPC histogram and the number of independent fit parameters, respectively.

*Pre-fitting to estimate the background parameters.* In a first step, we performed a global fit of the donor-only fluorescence decay $F_{DO}(t)$ and the FRET-induced donor decay $F_{DA}(t)$ using a two-component Gaussian distribution for the interdye distance. This allows us to estimate the fluorescence lifetimes and respective amplitudes of the donor-only sample as well as the parameters $t_{\text{shift}}$ and $F_0$ of the FRET-induced donor decay. The donor-only decay is described using two lifetime components:

$$f_{DO}(t) = \sum_{i=1}^{2} x_{D(0)}^{(i)} \exp\left(-t/\tau_{D(0)}^{(i)}\right), \tag{15.3}$$

where $\tau_{D(0)}^{(i)}$ and $x_{D(0)}^{(i)}$ are the lifetime and fraction of donor-only species $i$. The presence of the acceptor acts as an additional process that depopulates the donor excited state at a rate of:

$$k_{RET}(R_{DA}) = \frac{1}{\tau_{D(0)}}\left(\frac{R_0}{R_{DA}}\right)^6, \tag{15.4}$$

where $R_0$ is the Förster radius and $R_{DA}$ is the donor-acceptor separation distance. Note that $\tau_{D(0)}$ refers here to the lifetime of the species with the respective quantum yield that is used for the calculation of $R_0$, which can be different from the $\tau_{D(0)}^{(i)}$ obtained for the donor-only decay. For a given distance distribution, $p(R_{DA})$, the FRET-induced donor decay is then given by:

$$f_{DA}(t) = (1 - x_{\mathrm{DOnly}})f_{DO}(t)\left(\int p(R_{DA})\exp[-k_{RET}(R_{DA})\,t]\,dR_{DA}\right) + x_{\mathrm{DOnly}}f_{DO}(t), \tag{15.5}$$

where $x_{\mathrm{DOnly}}$ is the contribution of a donor-only signal due to acceptor photoblinking or photobleaching.

The two-component Gaussian distance distribution is given by:

$$p_{2G}(R_{DA}) = \sum_{i=1}^{2} x_{DA}^{(i)}\left(\sqrt{2\pi}\sigma_{DA,i}\right)^{-1}\exp\left[-\frac{\left(R_{DA} - \langle R_{DA}^{(i)}\rangle\right)^2}{2\sigma_{DA,i}^2}\right], \tag{15.6}$$

where $x_{DA}^{(i)}$ is the amplitude, $\langle R_{DA}^{(i)}\rangle$ the average interdye distance and $\sigma_{DA,i}$ the width of component $i$. The model is globally optimized with respect to the amplitude and lifetimes of the donor-only components. All parameters, except for the distance distribution $p(R_{DA})$, were fixed for the maximum entropy method model-free analysis discussed below.

*Model-free analysis.* The maximum entropy method (MEM) is an approach to extract the most unbiased distribution of a given parameter that provides a satisfactory fit to the experimental data[40–42]. Instead of minimizing the reduced chi-square, $\chi^2_{\mathrm{red}}$, the following functional is maximized:

$$\Theta = vS - \chi^2_{\mathrm{red}}, \tag{15.7}$$

where $v$ is a constant scaling factor and $S$ is the entropy functional of the parameter distribution. The entropy, $S$, of a discrete probability distribution $p_i$ is defined by:

$$S = -\sum_i p_i \log\frac{p_\mathrm{i}}{m_i}, \tag{15.8}$$

where $p_i$ is the distribution of the parameter of interest and $m_i$ describes the prior knowledge of the parameter distribution. We applied the MEM analysis to extract the distribution of interdye distances $R_{DA}$, $p(R_{DA})$. The measured FRET-induced donor fluorescence decay $f_{DA}^{exp}(t)$ is described as a superposition of exponential functions given by:

$$f_{DA}^{exp}(\mathrm{t}) = (1 - x_{\mathrm{DOnly}})f_{DO}(t)\left(\sum_j p\left(R_{DA}^{(j)}\right)\exp\left[-k_{RET}\left(R_{DA}^{(j)}\right)t\right]\right) + x_{\mathrm{DOnly}}f_{DO}(t) \tag{15.9}$$

where the summation is performed over a range of interdye distances $R_{DA}^{(j)}$ from 10 to 150 Å using a step size of 0.7 Å. Maximization of Θ is performed as described in Vinogradov and Wilson[43] over a wide range of values for the regularization parameter $v$. The choice of the regularization parameter $v$ was done by visual inspection of the L-curve plot of the negative entropy, -$S$, against the reduced chi-squared $\chi^2_{\text{red}}$. The resulting values for $\chi^2_{\text{red}}$ and the regularization parameter $v$ are given in Supplementary Table SN15.1. All analyses were performed using the *TauFit* module of the PAM software package[7].

*Prior distribution.* The prior distribution is based on the full apo ensemble derived in Huang et al.[44] For each of the 200 structures in the ensemble, the average interdye distance $\langle R_{DA}\rangle$ was determined using AV simulations[24,45]. To account for additional broadening due to the flexible dye linker, a kernel density estimate using a Gaussian kernel with a fixed width of 6 Å was performed to obtain the prior distribution as shown in Fig. 6d of the main text (light blue curve). We also performed a kernel density estimation of the interdye distance distribution without explicitly accounting for linker broadening using a Gaussian kernel by the *ksdensity* function of MATLAB, which computes the theoretically optimal bandwidth for normally distributed data[46]. This procedure returned a similar bandwidth of 5.7 Å. The different histograms and the kernel density estimate of the full apo ensemble are compared in Supplementary Figure SN15.1.

*Deconvolution of the probability distribution obtained by the MEM analysis.* The obtained distribution of the donor-acceptor distance $R_{DA}$ from the MEM analysis is broadened due to the flexible dye linkers. The magnitude of this additional broadening has previously been characterized to be on the order of ~6 Å[24]. To approximate the underlying distribution of mean donor-acceptor distances $\langle R_{DA}\rangle$, we performed a deconvolution of the $R_{DA}$ distribution obtained by MEM.

The deconvolution is performed using a Gaussian kernel with a width of $\sigma_{DA}$ = 6 Å, defined on a distance grid of $\langle R_{DA}\rangle \in [10\text{ Å}, 150\text{ Å}]$ with a resolution of 0.5 Å. The kernel matrix $\boldsymbol{Q}$ is defined as:

$$\boldsymbol{Q} = \begin{pmatrix} g_Q\left(R_{DA}^{(1)}; \langle R_{DA}\rangle^{(1)}\right) & \cdots & g_Q\left(R_{DA}^{(1)}; \langle R_{DA}\rangle^{(M)}\right) \\ \vdots & \ddots & \vdots \\ g_Q\left(R_{DA}^{(N)}; \langle R_{DA}\rangle^{(1)}\right) & \cdots & g_Q\left(R_{DA}^{(N)}; \langle R_{DA}\rangle^{(M)}\right) \end{pmatrix}, \tag{15.10}$$

where the indices $N$ and $M$ represent the number of sampling points of the discrete distributions for $R_{DA}$ and $\langle R_{DA}\rangle$, respectively, and the kernel functions are given by:

$$g_Q\left(R_{DA}^{(n)}; \langle R_{DA}\rangle^{(m)}\right) = \left(\sqrt{2\pi}\sigma_{DA}\right)^{-1} \exp\left[-\frac{\left(R_{DA}^{(n)} - \langle R_{DA}\rangle^{(m)}\right)^2}{2\sigma_{DA}^2}\right].$$

The discretized probability distribution of mean donor-acceptor distances $\langle R_{DA}\rangle$, $\boldsymbol{p}_{\langle \boldsymbol{R_{DA}}\rangle}$, is expressed as a row vector:

$$\boldsymbol{p}_{\langle \boldsymbol{R_{DA}}\rangle} = \begin{pmatrix} p_{\langle R_{DA}\rangle}^{(1)} \\ p_{\langle R_{DA}\rangle}^{(2)} \\ \vdots \end{pmatrix}, \tag{15.11}$$

and the discrete distribution of $R_{DA}$, $\boldsymbol{p}_{R_{DA}}$, is obtained as:

$$\boldsymbol{p}_{R_{DA}} = \boldsymbol{Q} \cdot \boldsymbol{p}_{\langle R_{DA} \rangle}. \tag{15.12}$$

To obtain an estimate of $\boldsymbol{p}_{\langle R_{DA} \rangle}$, we minimized the absolute value of the difference between the measured distribution $\boldsymbol{p}_{R_{DA}}^{(\text{exp})}$ and the distribution obtained by Eq. 15.12 above:

$$\min \left\| \boldsymbol{p}_{R_{DA}}^{(\text{exp})} - \boldsymbol{Q} \cdot \boldsymbol{p}_{\langle R_{DA} \rangle} \right\|, \tag{15.13}$$

under the constraint that all elements of $\boldsymbol{p}_{\langle R_{DA} \rangle}$ must be positive. In addition, we used the kernel density estimate of the distribution of $\langle R_{DA} \rangle$ as obtained from the full apo ensemble as a starting point for $\boldsymbol{p}_{\langle R_{DA} \rangle}$. The optimization is performed using the *fmincon* function of MATLAB. The deconvoluted $\boldsymbol{p}_{\langle R_{DA} \rangle}$ distributions are shown in Supplementary Figure 14.

**Supplementary Table SN15.1:** Reduced chi-squared $\chi^2_{\text{red}}$ and regularization parameter $v$ for the model-free analysis of fluorescence decays of U2AF2 measured in Lab #2.

| | **Dye Pair** | | | | | |
|---|---|---|---|---|---|---|
| | **Atto532-Atto643** | | **Alexa546-Alexa647** | | **Alexa488-Alexa647** | |
| **Sample** | $\chi^2_{\text{red.}}$ | $v$ | $\chi^2_{\text{red.}}$ | $v$ | $\chi^2_{\text{red.}}$ | $v$ |
| **U2AF2 apo** | 1.30 | 5.09 | 1.28 | 3.20 | 1.29 | 2.21 |
| **U2AF2 holo** | 1.03 | 1.05 | 1.24 | 1.05 | 1.13 | 1.05 |

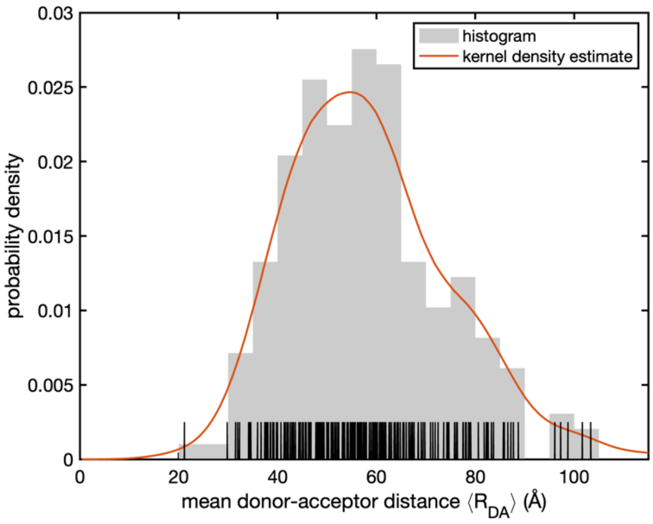

**Supplementary Figure SN15.1:** Probability distribution of the mean donor-acceptor separation obtained for the full apo-ensemble of U2AF2 as reported in Huang et al.[44] For each of the 200 structures in the ensemble, the mean donor-acceptor distance $\langle R_{DA} \rangle$ for the dye pair Atto532-Atto643 was computed from AV simulations. The vertical black lines indicate the individual values of $\langle R_{DA} \rangle$ and the gray bars shows the distance histogram computed with a bin width of 5 Å. The kernel density estimate (red line) was computed using a Gaussian kernel with a bandwidth of 6 Å, providing a smooth estimate of the probability density that is in good agreement with the histogram. The kernel density estimate was used as the prior distribution for the maximum entropy analysis of the fluorescence decays (Fig. 6d).

**Supplementary Note 16: Filtered-FCS Analysis of the U2AF2 kinetics.**

Filtered-FCS (fFCS) calculates the correlation functions for distinct species using statistical weighting based on the TCSPC patterns for the different species[47]. For the fFCS of the apo state of U2AF2 shown in Fig. 6e in the main text, the correlation functions were calculated as described in Barth et al.[48] using the PAM software package[7]. Briefly, two species with low and high FRET efficiency (LF, HF) were selected based on the FRET efficiency thresholds of $E \leq 0.6$ and $E \geq 0.9$, respectively. Filter functions for the fFCS analysis were calculated based on the concatenated TCSPC patterns of the donor and FRET-sensitized acceptor fluorescence decays. As a third species, a background pattern from a buffer measurement was included to account for the contribution of scattered laser light and constant background signal. "Purified" fFCS correlation functions were computed only for the detected and filtered double-labeled bursts. Signal occurring within 5 ms before or after the burst was included to obtain a better estimate of the diffusional part of the correlation function. To avoid detector afterpulsing in the species autocorrelation functions, the signal detected in the parallel and perpendicular channels of the MFD setup was filtered separately and cross-correlated. The two species autocorrelation functions (SACF) and two species cross-correlation functions (SCCF) were globally analyzed using a model with one diffusion term and two kinetic terms:

$$
\begin{aligned}
G(t_c) &= G_{\mathrm{diff}}(t_c) G_{\mathrm{kin}}(t_c) \\
G_{\mathrm{diff}}(t_c) &= \frac{1}{\langle N \rangle}\left(1+\frac{t_c}{t_{\mathrm{diff}}}\right)^{-1}\left(1+\frac{t_c}{p\, t_{\mathrm{diff}}}\right)^{-\frac{1}{2}} \\
G_{\mathrm{kin}}^{SACF}(t_c) &= 1 + A_1 e^{-\frac{t_c}{t_{R,1}}} + A_2 e^{-\frac{t_c}{t_{R,2}}} \\
G_{\mathrm{kin}}^{SCCF}(t_c) &= 1 - A_1 e^{-\frac{t_c}{t_{R,1}}} - A_2 e^{-\frac{t_c}{t_{R,2}}}
\end{aligned}
\qquad (16.1)
$$

Here, $G_{\mathrm{diff}}(t_c)$ and $G_{\mathrm{kin}}(t_c)$ are the diffusion and kinetic part of the correlation function $G(t_c)$ at lag time $t_c$. For the diffusion component, $\langle N \rangle$ is the average number of particles in the observation volume, $t_{\mathrm{diff}}$ is the diffusion time and the structural factor $p$ is given by the ratio of the axial to the lateral width of the observation volume (typically, $p$ = 5-10). For the kinetic component, $t_{R,1}$ and $t_{R,2}$ are the kinetic relaxation times with amplitudes $A_1$ and $A_2$ respectively. For the analysis, the kinetic relaxation times and diffusion time are optimized globally over all four curves. The amplitude of the diffusional part (given by $\langle N \rangle^{-1}$) and the kinetic terms ($A_1$ and $A_2$) were optimized individually for each of the four curves. This is necessary as the amplitude information in fFCS is less reliable compared to the time evolution of the curves due to the effect of imperfect filters[47].

**Supplementary Note 17: Dynamics of U2AF2.**

From previous work, we were aware that U2AF2 is a dynamic protein system whose conformational distribution can be shifted by the addition of an RNA ligand[49]. From the BVA and $E$-$\tau$ plots (Fig.4), it is evident that the apo protein is dynamic on the sub-millisecond timescale. The $E$-$\tau$ plot already contains information regarding the timescale of the dynamics of the sample with respect to the duration of the burst. A single transition is already sufficient to cause a shift for that burst from the static FRET line in the $E$-$\tau$ plot. Hence, dynamics that are up to an order of magnitude slower than the diffusion time can still be detected. In this case, individual "static" populations should be observable on the static FRET line with a smearing between the states coming from the few bursts where transitions occurred during the burst. For faster dynamics, several transitions occur during a burst and the individual "static" populations disappear with only a single, dynamically averaged state observable on the dynamic FRET line. The position of the population along the dynamic FRET line depends on the equilibrium between the different states[15]. For U2AF2, the dynamics are fast enough that we only observed an average FRET state and the individual populations are not visible.

The dynamics of U2AF2 is further complicated by the fact that, as already known from NMR and SAXS data, there is not a single closed or a single open conformation, but a family of conformations[44]. To get an idea of the distribution of FRET efficiencies in the sample, we performed a model-free analysis of the donor fluorescence lifetime using the maximum entropy method[40–42] to infer the underlying distribution of interdye distances (Supplementary Note 15). Our analysis revealed a higher population of compact states compared to the full structural ensemble of Huang et al[44] (Fig. 6d in the main text and Supplementary Fig. 14). We note that smFRET and NMR provide complementary information as the two techniques are sensitive to distinct distance ranges of 1-3 nm for paramagnetic relaxation enhancement (PRE) measurements and 4-12 nm for smFRET[50]. The structural ensemble was selected to satisfy the experimental averages of the measured PREs while the information from SAXS constrains the overall structural extension of the ensemble. The slight deviation of the smFRET information from the published structural ensemble indicates that a further refinement of the structural ensemble could be performed. We speculate that an alternative ensemble could be found that provides better agreement with the smFRET data while still satisfying the NMR/SAXS constraints[50].

After establishing the distribution of FRET efficiencies observable within the sample, we next determined the timescale of the dynamics. To get a model-independent estimation of the timescale of kinetics faster than the burst duration, we utilized an FCS approach. One possibility to visualize the dynamics is by performing a cross-correlation analysis between the donor and acceptor signals (FRET-FCS)[51,52]. We choose to use filtered-FCS (fFCS)[47,53] (Supplementary Note 16) as it increases the contrast of the FCS signal in comparison to FRET-FCS by weighing the correlation signals depending on the difference in the lifetime, color and anisotropy of fluorescence fluctuations between the bursts showing the lowest and highest FRET values. For the Atto532-Atto643 labeled protein, we observed two relaxation times with values of ~ 10 µs and ~ 200 µs (Fig. 6e, Supplementary Table 15). We assign the faster kinetics to dynamics of the detached domains and the slower rate to interconversion between compact conformations within the conformational ensemble.

A second approach for extracting the dynamic timescales from a sample is to use the dynamic photon distribution analysis (PDA) method[14]. Dynamic PDA allows one to delineate the conformational heterogeneity over sub-millisecond transitions from the width of the FRET histograms beyond shot-noise by analyzing the raw photon counts[54,5514]. The FRET efficiency histograms are calculated using different time-windows for the binning and then fit using a global model (Fig. 6f in the main text and Supplementary Figure 16). There is little change in the FRET efficiency histograms when varying the integration time between 0.5 ms, 1.0 ms, 1.5 ms and 2.0 ms. From a global fit to the different histograms for the apo and holo conditions, the states as well as the conversion rates between them (provided they are between ca 100 µs and the timescale of diffusion) can be extracted. To simplify the complexity of the dynamic PDA model, we used a broad Gaussian function to empirically describe the fast, dynamic ensemble observed with fFCS and used dynamic PDA to look for slower kinetics. We observed a slow interconversion on the timescale of 10-100 ms between the very-high-FRET state at $E \sim 0.95$ (consistent with the closed NMR structure, PDB 2YH0[56]) and the main, dynamically averaged population at $E \sim 0.85$. The high FRET population (E = 0.85) is clearly distinct from a second peak in the $E$-$\tau$ plot of E ~ 0.95 and has a very short donor lifetime. The very-high-FRET signal is also reminiscent of what is observed for dye-dye interactions[57]. If this is true, the rates given by dynamic PDA should change when a different dye-pair is used whereas the protein dynamics should remain unchanged. Indeed, when evaluating measurements using the Alexa546 – Alexa647 dye pair, the fast dynamics remains unchanged (90 – 200 µs) whereas the amplitude of this very-high-FRET peak and the rate of the slow dynamics changes slightly (Supplementary Figure 19 and Supplementary Table 18). This demonstrates the importance of performing smFRET measurements with at least two different dye-pairs[57]. Here, the high stoichiometry state at very-high FRET efficiency was excluded and the observed dynamics suggest there is a transient stabilization of the closed structure.

Due to the complexity of the dynamics, we did not ask the various groups to provide a detailed analysis of the kinetics. We did ask laboratories, when possible, to provide an estimate of the timescale of the dynamics, when present. Five labs could contribute to a dynamic quantification of U2AF2 (Supplementary Table 15). These all correctly reported a quasi two-state behavior of the U2AF2 system. A comparison of kinetic rates based on further evaluation with dynamic PDA and fFCS for U2AF2 from different labs showed good consistency. Specifically, the calculated relaxation times for the apo state evaluated with fFCS were consistent with an approximately two-fold variation across labs (~200, ~320 and ~370 µs from three different labs). A dynamic PDA analysis for the holo-state kinetic rate estimations were not fully consistent most likely due to the variation seen in the FRET histograms (perhaps due to differences in temperature). Three among the five groups provided the kinetic rates for the holo -state with only ~20% variation of the reported relaxation times ($\tau_R = 1/(k_{12}+k_{21})$) of 1.25 ms, 1.42 and 1.6 ms (see Supplementary Table 15).

*Challenges with the U2AF2 sample and analysis:*

From the detailed analysis described above, it is clear that U2AF2 is not a simple two-state dynamic system. Apo U2AF2 exhibits dynamics on the microsecond timescale between a family of open and of closed conformations. Both apo and holo measurements exhibit a slow dynamic exchange between a very-high-FRET populations (at $E \sim 0.95$) and the other FRET states.

Beyond the complicated dynamics, other factors also impacted the analysis of the dynamic state. First of all, the measured RNA ligand concentration was not high enough to saturate binding to U2AF2. From the measured binding affinity of $K_d \approx 1.2$ µM (Supplementary Figure 7d), the ligand concentration of 5 µM used in these measurements lead to approximately 85% of the sample having a ligand and 15% remaining in the apo conformation. This corresponds well to the measured ratio of the two populations. An alternative explanation is that the protein exchanges between a quasi-static state and a dynamic conformation where one of the RRM domains transiently releases the RNA strand. However, there is no evidence that the RNA-bound state of U2AF2 interconverts with the apo-like conformation on the timescale of a burst (~ 10 ms). This is consistent with what is expected assuming a $K_d$ of 1.2 µM and a measured $k_{on}$ of 0.7 $\mu M^{-1}\ s^{-1}$, leading to a $k_{off}$ of ~ 1s. This timescale is too slow to be detected with the solution-based measurements. Hence, we attributed the dynamics detected by BVA and the $E$-$\tau$ plots in the holo measurements to the significant presence of proteins not having an RNA ligand bound (Fig.5). Also, the timescale of the kinetics for the dynamic population was similar to that of the apo state (~ 100 - 200 µs). The second FRET peak with a FRET efficiency of $E \sim 0.44$ observed in the presence of RNA is close to the static line suggesting that the RNA bound conformation is static.

A second challenge in analyzing the U2AF2 data comes from the fraction of molecules with a higher than normal stoichiometry (Supplementary Figure 17). The origin of this population is not known. It could be due to dye-dye interactions where a change in fluorescence intensity yields a different stoichiometry value. Two populations with similar FRET efficiencies but different stoichiometries indicates inconsistencies in the dataset and most likely require individual $\gamma$ factors for a proper analysis. At any rate, both populations should not be included in determining a single detection correction factor.

In summary, apo U2AF2 appears to undergo fluctuations between families of open and closed states with dynamics on the timescale of 100-200 µs. Dynamics on the millisecond timescale are also observed due to the formation of stable compact states. Fluctuations between an open and a closed conformation is a fair approximation of the dynamics, but an ensemble of open and closed conformations are needed to fully describe the measured data as shown in Fig. 6c-f.

#### Supplementary Note 18: Overview of set-ups and analysis software used across all labs.

##### Lab#1

All sample solutions were measured in Lab-Tek I chamber slides at a concentration of ~50-100 pM. Single-molecule FRET experiments with MFD-PIE were performed on a homebuilt confocal setup as described previously[4].

For samples labeled with Alexa546-Alexa647 and Atto532-Atto643, our two-color green-red setup was used: Fluorescent donor molecules were excited by an amplified, frequency-doubled diode laser (PicoTA 530, PicoQuant, Berlin) at 532 nm and acceptor molecules were excited by a pulsed diode laser (LDH-D-C 640, PicoQuant) at 640 nm. Both lasers were operated at a power of 50 μW measured at the sample, operated at a repetition rate of 26.66 MHz and synchronized with a delay of 18 ns. The laser light was guided into the epi-illuminated confocal microscope base (Nikon Eclipse TE300) and focused by a 60X water immersion objective (Plan Apo IR 60x/1.27, Nikon). For excitation, only parallel and perpendicular polarized light was selected using a Glan-Thompson polarizer (GTHM Polarizer, Thorlabs) before focusing into the objective. The emitted fluorescence was collected through the objective and spatially filtered using a pinhole with 75 μm diameter. The fluorescence emission was first separated for polarization using a polarizing beam-splitter (PBS3, Thorlabs) before being spectrally separated into donor and acceptor channels by a dichroic mirror (640DCXR; AHF Analysentechnik). Fluorescence emission was filtered (donor: Brightline HQ582/75, acceptor: Brightline HQ700/75, AHF Analysentechnik) and focused on avalanche photodiodes (SPCM-AQR, Perkin-Elmer) for detection. The detector outputs were recorded using four TCSPC cards (SPC154; Becker and Hickl)

For samples labeled with Alexa488-Alexa647, our three-color setup was used as described previously[58]. For this dye-pair combination, we switched off the 560 nm laser and associated detectors. In general, the three-color set-up has three pulsed lasers with a ~20 ns delay between the laser pulses (pulse frequency of 16.7 MHz) (PicoQuant, Germany; LDH-D- C-485, LDH-D-TA-560, LDH-D-C-640). The lasers were synchronized using a laser driver (PicoQuant, Germany; Sepia II). A 60x water immersion objective with 1.27 N.A. (Nikon, Germany; Plan Apo IR 60x 1.27 WI) was used for focusing the lasers into the sample. The measured laser powers before the objective were ~120 μW for blue, ~75 μW for green and ~35 μW for red laser. Fluorescence was collected by the same objective, separated from the laser excitations using a polychroic mirror (AHF Analysentechnik; zt405/488/633, Germany) and confocal geometry was achieved using a 50 μm pinhole. Light passing through the pinhole was further separated by a polarizing beam splitter (Thorlabs, Germany). Separation of blue and red wavelengths were performed with a dichroic mirror (AHF Analysentechnik; 640DCXR). Emission filters (AHF Analysentechnik; ET525/50, ET670/30) were placed right before the APD detectors (LaserComponents, 2x COUNT-100B for blue detection; Perkin Elmer, 2x SPCM-AQR14 for red detection). Photons were recorded and synchronized to the lasers pulses using a TCSPC module (PicoQuant; HydraHarp400).

Data analysis was performed using the PAM (PIE Analysis with Matlab) software package as described elsewhere[7]. Single-molecule events were identified using a sliding time window analysis with a threshold of 50 photons, a time window of 500 μs and a minimum photon number of 10. To remove photo-blinking and -bleaching events, the ALEX-2CDE filter was applied using an upper threshold of 12.

**Lab#2**

All sample solutions were measured in NUNC chambers (Lab-Tek, Thermo Scientific) with 500 µL sample volume and a pM concentration. Single-molecule FRET experiments with PIE were performed on a homebuilt confocal setup as described previously[23].

Setup #1: for samples labelled with Alexa Fluor 546 – Alexa Fluor 647 and Atto 532-Atto 643: The fluorescent donor molecules are excited by a pulsed white light laser source (SuperK EXTREME, NKT Photonics), using a modulator (SuperK Varia, NKT Photonics), operated at 25 MHz, 80 µW. The acceptor molecules are excited by a pulsed diode laser (LDH-D-C 640), operated at 25 MHz and 10 µW. Laser powers were measured at objective. Laser light is guided into the epi-illuminated confocal microscope (Olympus IX71, Hamburg, Germany) by dichroic beamsplitter F68-532_zt532/640NIRrpo (AHF, Germany) and focused on a sample by a water immersion objective (UPlanSApo 60x/1.2w, Olympus Hamburg, Germany). The emitted fluorescence is collected through the objective and spatially filtered using a pinhole with 100 µm diameter and further split into parallel and perpendicular components using polarizing beam splitter cube (VISHT11, Gsänger). Light is then spectrally split into "green" and "red" spectral windows by a dichroic mirror (T640lpxr, AHF, Germany). Fluorescence emission was filtered (donor: 47-595/50 ET, acceptor: HQ 730/140, AHF, Germany) prior to detection using avalanche photodiodes (SPCM-AQRH 14, Excelitas). The detector outputs were recorded by a TCSPC module (HydraHarp 400, PicoQuant).

Setup #2: for samples labelled with Alexa488-Alexa647: Donor molecules were excited by a pulsed diode laser (LDH-D-C 485, PicoQuant) at 485 nm. Acceptor molecules were excited with pulsed diode laser (LDH-D-C 640, PicoQuant) at 635 nm. Lasers were operated with the repetition frequency of 32 MHz, and with a delay with respect to each other of 10.5 ns. Laser powers were measured at the objective and were 60 µW for donor excitation laser and 10 µW for acceptor excitation laser. Laser light is guided into the epi-illuminated confocal microscope (Olympus IX71, Hamburg, Germany) by dichroic beamsplitter FF500/646-Di01 (Semrock, USA), and focused on the sample by a water immersion objective (UPlanSApo 60x/1.2 NA, Olympus Hamburg, Germany). The emitted fluorescence is collected through the objective and focused on a 100 µm pinhole. Using a polarizing beam splitter cube, emitted light is divided into its parallel and perpendicular components. This is then followed by light being split into two spectral windows, "green" and "red", using longpass filter Q595, and then again using 50/50 beam splitters resulting in a total of eight detection channels. Additionally, bandpass filters are placed in front of the detectors (FF01-530/43-25; AHF, Tübingen, Germany for donor molecules and HQ 720/150 nm; AHF, Tübingen, Germany for acceptor molecules). Detection is performed using eight avalanche photodiodes (4 green channels: τ-SPAD (PicoQuant, Germany) and 4 red channels: SPCM-AQR-14 (Perkin Elmer). The detector outputs were recorded by a TCSPC module (HydraHarp 400, PicoQuant). For both setups data analysis was performed using home-written LabView-based software. Burst search was performed as described[12] ,using APBS (All Photon Burst Search) method and inter-photon times as threshold.

**Lab#3**

Sample solutions were measured with 100 µl drop on coverslip with concentration of around 50 pM. Single-molecule FRET experiments with ALEX were performed on a homebuilt confocal microscope as described previously[59]. The fluorescent donor molecules are excited by a diode laser OBIS 532-100-LS (Coherent, USA) at 532 nm operated at 60 µW at the sample in alternation mode. The fluorescent acceptor molecules are excited by a diode laser OBIS 640-100-LX (Coherent, USA) at 640 nm operated at 25 µW at the sample in alternation mode (100 µs alternation period). The lasers are combined by an aspheric fiber port (PAF2S-11A) and coupled into a polarization maintaining single-mode fiber P3-488PM- FC-2 (Thorlabs, USA). The laser light is guided into the epi-illuminated confocal microscope

(Olympus IX71, Hamburg, Germany) by dual-edge beamsplitter ZT532/640rpc (Chroma/AHF) focused by a water immersion objective (UPlanSApo 60x/1.2w, Olympus Hamburg, Germany). The emitted fluorescence is collected through the objective and spatially filtered using a pinhole with 50 µm diameter and spectrally split into donor and acceptor channel by a single-edge dichroic mirror H643 LPXR (AHF). Fluorescence emission was filtered (donor: BrightLine HC 582/75 (Semrock/AHF), acceptor: Longpass 647 LP Edge Basic (Semroch/AHF), focused on avalanche photodiodes (SPCM-AQRH-64, Excelitas). The detector outputs were recorded by a NI-Card PCI-6602 (National Instruments, USA).

Data analysis was performed using home written software package as described[59] .Single-molecule events were identified using a All-Photon-Burst-Search algorithm with a threshold of 15, a time window of 500 µs and a minimum total photon number of 150.

**Lab#4**

All sample solutions are measured in home-built 60 µL chambers with ~100-pM concentration. PIE-FRET experiments are carried out on a home build confocal microscope. The fluorescent donor molecules are excited by a pulsed diode laser (LDH-P- FA-530B, PicoQuant), at 532 nm operated at 20 MHz, with an excitation power of 55 µW at the sample in PIE experiment. The fluorescent acceptor molecules are excited by a pulsed diode laser (LDH-D-C-640, PicoQuant), at 639 nm operated at 20 MHz, with an excitation power of 50 µW at the sample in PIE experiment. The laserpulses are altered on the nanosecond timescale by a multichannel picosecond diode laser driver (PDL 828 “Sepia II”, PicoQuant GmbH) with an oscillator module (SOM 828, PicoQuant GmbH). The lasers were coupled into a single mode fiber (P3-488PM-FC, Thorlabs GmbH) to obtain a Gaussian beam profile. Circular polarized light is obtained by a linear polarizer (LPVISE100-A, Thorlabs GmbH) and a quarter-wave plate (AQWP05M- 600, Thorlabs GmbH). The laser light is guided into the epi-illuminated confocal microscope (Olympus IX71, Hamburg, Germany) by dual-edge beam splitter (z532/633, AHF analysentechnik AG) focused by an oil immersion objective (UPLSAPO100XO, NA 1.40, Olympus Hamburg, Germany). The emitted fluorescence is collected through the objective and spatially filtered using a pinhole with 50 µm diameter and spectrally split into donor and acceptor channel by a single-edge dichroic mirror (640DCXR, AHF Analysentechnik AG, Germany). Fluorescence emission is filtered (donor: Brightline HC582/75 (AHF Analysentechnik AG), RazorEdge LP 532 (Laser 2000 GmbH), acceptor: (Shortpass 750, AHF Analysentechnik AG; RazorEdge LP 647, Laser 2000 GmbH), focused on avalanche photodiodes (SPCM-AQRH-14-TR, Excelitas Technoligies GmbH & Co. KG). The detector outputs were recorded by a TCSPC module (HydraHarp 400, PicoQuant). The setup is controlled by a commercial software package (SymPhoTime64, Picoquant GmbH).

Data analysis is performed using PAM software package as described[7] . Single-molecule events are identified using a two channel APBS-algorithm with a threshold of 5 photons per time window, a time window of 500 µs and a minimum photon number of 20. To remove photo-blinking and -bleaching events, the ALEX-2CDE filter was applied using an upper threshold of $10^8$.

**Lab#5**

All sample solutions were measured in 200 µl PBS buffer with a labelled protein concentration of 25-100 pM. In short: Single-molecule FRET experiments with ALEX were performed on a homebuilt confocal microscope as described previously[59–61] .The fluorescent donor molecules are excited by a spectrally filtered laser beam of a pulsed supercontinuum source (SuperK Extreme, NKT Photonics) with an acousto-optical tunable filter (AOTFnc-VIS, EQ Photonics), at 532 nm and 640 nm. The laser light is guided into a single-mode fiber (PM-S405-XP, Thorlabs) and the collimated beam (Focusing collimator MB06, Q-Optics/Linos) was coupled into an oil-immersion objective (60×, NA 1.35, UPLSAPO 60XO, Olympus) by using a dichroic beam splitter (zt532/642rpc, AHF Analysentechnik) mounted on an inverse microscope body (IX71, Olympus). The emitted fluorescence is collected through the objective and spatially filtered using a pinhole with 50 µm diameter and spectrally split into donor and acceptor channel by a single-edge dichroic mirror (640DCXR, AHF Analysentechnik). Fluorescence emission was filtered (donor: Brightline HC582/75, acceptor ET700/75; AHF Analysentechnik), focused on avalanche photodiodes (Tau-SPAD, PicoQuant). The detector outputs were recorded by a TCSPC module (HydraHarp 400, PicoQuant).

Data analysis was performed as described[59–61]. Single-molecule events were identified using an All Photon Burst Search algorithm with a threshold of 15, a time window of 500 µs and a minimum photon number of 200.

**Lab#6**

Single molecule measurements were carried out on a home-built confocal microscope[62] .Pulsed green and red laser light (532nm, LDH-P-FA-530 and 640nm, LDH-D-C-640, respectively, PicoQuant) was polarised, overlaid and focused on the sample by a 60x water immersion objective (CFI Plan Apo VC 60XC/1.2 WI, Nikon). Excitation light was separated from the emitted light by a dichroic mirror (F53-534 Dual Line beamsplitter z 532/633, AHF). The emitted light was then guided through a further dichroic mirror (F33-647 beam splitter 640 DCXR, AHF) to separate donor and acceptor fluorescence. After spectral separation pinholes with a diameter of 150 micrometer refined the detection volume to about 8fL. Finally, the two photon streams were separated by polarizing beam splitters into their parallel and perpendicular parts and recorded by single-photon detectors (two SPCMAQR-14, PerkinElmer and two PDMseries APDs, Micro Photon Devices). Time-correlated single photon counting with picosecond resolution and data collection was performed by a HydraHarp400 (PicoQuant) and the Symphotime 32 software (PicoQuant).

The sample was measured within a BSA passivated well. Data analysis was performed using the PAM software package. Bursts were identified first using APBS with at minimum of 50 photons per burst, a 500 µs time window and at least 20 photons per time window to determine correction factors $\boldsymbol{\alpha}$ and $\boldsymbol{\delta}$. Then a DCBS was performed with similar parameters and at least 12 photons in the green and 10 in the red channel. The other two correction factors were determined from the bursts identified in the second burst search.

**Lab#7**

All sample solutions were measured a drop on a coverslip, sealed in an airtight chamber, with concentration ~50 pM. The general scheme of the setup is described[63] . In short, Single-molecule FRET experiments with ALEX were performed on a homebuilt confocal setup, the smfBox. The fluorescent donor molecules are excited by a continuous wave laser at 515 nm operated at 100 µW at the sample. The fluorescent acceptor molecules are excited by a continuous wave laser, at 638 nm operated at 230 µW at the sample. The laser light is guided into the custom-built microscope body and focused by an oil immersion objective (UPLSAPO 60× NA = 1.35, Olympus Hamburg, Germany). The emitted fluorescence is collected through the objective and spatially filtered using a pinhole with 20 µm diameter and spectrally split into donor and acceptor channels by a single-edge dichroic mirror (NC395323 - T640lpxr, Chroma, USA). Fluorescence emission was filtered (donor: FF01-571/72-25, acceptor: FF01-679/41-25, Semrock, USA), focused on avalanche photodiodes (SPCM-AQRH-14,

Excelitas). The detector outputs were recorded by a national instruments card (PCIe-6353), with acquisition controlled using custom software available on our github (see smfBox reference above). A full description of the procedure is available[64].

Data analysis was performed using the PAM software package as described[7] .Single-molecule events were identified using a dual channel burst search algorithm with a threshold of 5 photons, a time window of 500 µs and a minimum photon number of 50.

**Lab#8**

All sample solutions were measured as a drop on a passivated with BSA (1mg/ml) coverslip with protein concentration from 20 to 50pM. SmFRET measurements were carried out on a custom built confocal setup using time-correlated single photon counting (TCSPC), which combines pulsed interleaved excitation with multi-parameter fluorescence detection (PIE-MFD) [4]. In brief, the fluorescent donor molecules are excited by a pulsed diode laser (LDH-P-FA 530L, PicoQuant) at 531nm, operated at 20MHz with 60µW power. The fluorescent acceptor molecules are excited by a pulsed diode laser (LDH-D-C 640, PicoQuant) at 640nm, operated at 20MHz with 40µW power. The laser light is coupled into a fiber collimator (60FC-4-RGBV11-47, Schäfter + Kirchhoff GmbH, Hamburg). After the collimator, the laser light is reflected by a dichroic mirror (HC quadband laser beamsplitter R405/488/532/635, AHF Analysentechnik AG, Tübingen) and then focused into the sample by a 1.2 NA water immersion objective (CFI Plan Apochromat VC 60x, Nikon GmbH, Düsseldorf). The laser driver, Sepia II (PicoQuant), operates the lasers such that they are pulsed (20 MHz) and shifted (by ~25ns) with respect to each other. The emitted fluorescence is collected by the objective and spatially filtered using a pinhole with 75µm diameter and spectrally split into two beams: parallel and perpendicular with respect to the excitation light by a polarizing beam splitter cube (PBS201, Thorlabs, Munich). After the PBS, the two beams are then split into donor and acceptor channel according to wavelength (HC BS 649, AHF Analysentechnik), resulting in 2 beams per polarization (green and red). Fluorescence emission was filtered (donor: 582/75 Brightline HC, acceptor: 700/75 ET bandpass, AHF Analysentechnik) and focused on a single-photon avalanche diode (τ-SPAD-100, PicoQuant) by a lens of 100 mm focal length. The detector outputs were recorded by a TCSPC module HydraHarp 400 (PicoQuant) which is synchronized with the laser driver. The photon arrival times were recorded with 16ps resolution for microtime and synchronization period of 50ns for macrotime.

Data analysis was performed using (PAM–PIE analysis with MATLAB v2.0 (develop branch up to commit 320364c4) software package as described [7]. Single-molecule events were identified using APBS as burst search method with a threshold of 100 photons per burst, a time window of 500 µs and a minimum photon number of 5. To remove photo-blinking and -bleaching events, the ALEX-2CDE filter was applied using an upper threshold of 10 [8].

**Lab#9**

The smFRET experiments were performed on a custom-built confocal detection-based microscope. The general scheme of the setup was previously described[65] All sample solutions were measured in 8-well chamber slides with a final volume of 200 µl at sample concentration of 50 pM. A pulsed laser diode (LDH-TA-560, Picoquant, Germany), pulsed at 40 MHz, 35 µW, is used for donor excitation; A pulsed laser diode (LDH-D-C-660, Picoquant, Germany), pulsed at 40 MHz, 100 µW, is used for acceptor excitation. We used pulsed interleaved excitation (PIE) scheme to alternatingly excite donor and acceptor fluorophores to retrieve the stoichiometry (S) information[66] .The lasers were cleaned up with excitation filters (Brightline FF01-572/15 and Brightline FF01-650/13, Semrock), passed through the polarizer (GL-10A, Thorlabs) before entering the objective lens (60×, water immersion, NA=1.27, Nikon) through the central dichroic mirror (ZT405/488/561/660/905rpc-UF3, Chroma). The fluorescence emission signal was collected through the same objective lens and spatially filtered using a pinhole with 100 µm diameter and split into parallel and perpendicular polarization axis. Fluorescence emission was then filtered (donor: Brightline HC612/69, acceptor: ET700/75, Chroma) after separated into donor and acceptor channels via the dichroic mirror (FF650-DI01, Semrock). Photon signals were detected on photon counting detectors (τ-SPAD, Picoquant, Berlin) and recorded by a TCSPC module (Hydraharp400, Picoquant, Berlin). Acquired data were subject to multi-parameter fluorescence analysis and processed burst-wise for fluorescence intensities and lifetime with a threshold of 30 photons per burst[5,23]. All acquired data was performed by a custom written Igor-Program (Wavemetrics)[65] and the burst variance analysis (BVA) was analyzed by using algorithm from PAM-PIE with MATLAB[7] .

**Lab#10**

All sample solutions were measured as drops on a coverslip with a concentration of 50 pM. A custom-built confocal microscope was used for single-molecule FRET experiments as previously described[63-67] and the setup was modified to allow alternating-laser excitation of donor and acceptor fluorophores[72,73]. For this purpose, the fibre-coupled outputs of a 532-nm laser (operated at 250 µW) and a 640-nm laser (operated at 60 µW) were alternated with a modulation frequency of 20 kHz. Both beams were spatially filtered and coupled into an oil-immersion objective (60x 1.35 NA, UPLSAPO 60XO, Olympus). The same objective was used to collect the resulting fluorescence; the emission was separated from excitation light by a dichroic mirror, focused onto a 200-µm pinhole, and subsequently split spectrally on two avalanche photodiodes (SPCM-AQR-14, PerkinElmer, UK) detecting the donor and acceptor fluorescence with two distinct spectral filters (green: 585DF70; red: 650LP). Custom-written LabVIEW software was used to register and evaluate the detected signal. Fluorescence photons were assigned to either donor or acceptor-based excitation with respect to their photon arrival time. Two characteristic ratios, the fluorophore stoichiometry S and apparent FRET efficiency E*, were calculated for each fluorescent burst, yielding a two-dimensional histogram 6-7. One-dimensional E* distributions for donor-acceptor species were obtained using a $0.4 < S < 0.8$ threshold. These E* distributions were fitted with Gaussian functions, yielding the mean E* value for each distribution.

**Lab#11**

All sample solutions were measured in custom made glass chambers at a concentration of 15 pM. The general scheme of the setup is described in Krainer et al. 2018[74]. In short, single-molecule FRET experiments were performed on a custom-built confocal microscope as described previously by Hartmann et al. 2015[75]. The fluorescent donor and acceptor labelled molecules were excited in PIE mode with 25 MHz repetition rate by a green pulsed diode laser (LDH-P-FA 530L, PicoQuant, Berlin, Germany), at 530 nm wavelength, and a red pulsed diode laser (LDH-D-C 640, PicoQuant), at 640 nm wavelength with laser powers of 110 µW and 90 µW before objective, respectively. The laser light is guided into the inverted confocal microscope (Nikon Eclipse Ti - Nikon, Tokyo, Japan) by a dual-edge beam splitter zt532/640rpc (Chroma, Bellows Falls, VT, USA) and focused by a water immersion objective (FI Plan Apo WI 60x (NA 1.2), Nikon). The emitted fluorescence is collected through the same objective and spatially filtered using a pinhole with 50 µm diameter. In order to detect the fluorescence anisotropy, the spatially filtered light is separated according to its polarization by a polarizing beam splitter (PBS201, Thorlabs, Newton, NJ, USA) and guided in the parallel and perpendicular detection channel. In each detection channel the light is spectrally split into donor and acceptor emission by a single-edge dichroic mirror (FF650-Di01, Semrock, New York, NY, USA). The polarization separated and spectrally split fluorescence emission was bandpass filtered (donor: FF01-

582/75, Semrock, acceptor: ET700/75M, Chroma) and focused on avalanche photodiodes (SPCM-AQRH-14, Excelitas, Waltham, MS, USA). The detector outputs were recorded by a TCSPC module (HydraHarp 400, PicoQuant).

Data analysis was performed using a custom written software package as described in Hartmann et al. 2017[76]. Single-molecule events were identified from the acquired photon stream as fluorescence bursts with a maximum inter-photon time of 50 µs containing a minimum total number of 40 photons for protein 1 and 100 photons for protein 2 after background correction and a Lee filter with window size four. To remove photo-blinking and bleaching events, the ALEX-2CDE filter as described by Tomov et al. 2012[8] was applied using an upper threshold of 8.

**Lab#12**

All sample solutions were measured in Cellview chamber slides (Greiner BioOne, Frickenhausen, Germany) with a dilution of 1 to 600 from the received stocks. Single-molecule FRET experiments with PIE were performed on a commercial MicroTime 200 confocal microscope (PicoQuant, Berlin, Germany) as described [77]. The fluorescent donor molecules are excited by a pulsed diode laser (LDH-P-FA 530B, PicoQuant), at 532 nm operated at 20 MHz, 60 µW at the sample in the PIE excitation mode . The fluorescent acceptor molecules are excited by a pulsed diode laser (LDH-D-C-640, PicoQuant, Berlin, Germany; clean up filter: zet636/20x, Chroma, Bellow Falls, VT, USA), at 640 nm operated at 20 MHz, 30 µW at the sample in the PIE excitation mode. The laser light is guided into the inverted IX73 confocal microscope (Olympus, Hamburg, Germany) by a dual-edge beam splitter ZT532/640rpc-UF3 (Chroma, Bellow Falls, VT, USA) and then focused by a water immersion objective (UPlanSApo 60x/1.2w, Olympus, Hamburg, Germany). The emitted fluorescence is collected through the objective and spatially filtered using a pinhole with 50 µm diameter and spectrally split into donor and acceptor channel by a single-edge dichroic mirror (T635lpxr; Chroma, Bellow Falls, VT, USA). Fluorescence emission was filtered (donor: ff01-582/64; Semrock, Rochester, NY, USA; acceptor: H690/70; Chroma, Bellow Falls, VT, USA) and focused on avalanche photodiodes (SPCM-AQRH-14-TR, Excelitas Technologies, Waltham, MS, USA). The detector outputs were recorded by a TCSPC module (HydraHarp 400, PicoQuant, Berlin, Germany). The setup was controlled with the SymPhoTime64 software package (PicoQuant, Berlin, Germany).

Data analysis was performed using the PAM software package as described [7]. Single-molecule events were identified using an APBS and DCBS with a threshold of 100 photons per burst, a sliding time window of 500 µs and a minimum photon number of 10.

**Lab#13**

All sample solutions were measured as a drop on a coverslip (Roth, Karlsruhe, heated to 500 °C for 2 h), covered with a humidity chamber with a dilution of 1 to 300 (sample 1) and 1 to 200 (sample 2 and 3) of the delivered stock solution. The general scheme of the setup is described[78]. In short: Single-molecule FRET experiments with PIE were performed on a homebuilt confocal microscope as described[78] .The fluorescent donor molecules are excited by a cw DPSS "Crysta Laser" (GCL-005-L, Laser2000, Wessling) at 532 nm with 40 µW at the sample in the PIE experiment. The fluorescent acceptor molecules are excited by a pulsed diode laser (LDH-P-C 635, PicoQuant GmbH), at 635 nm operated at 10 MHz, 5 µW at the sample in PIE experiment. The laser light is guided through a single-mode fibre (SMC-460, Schäfter&Kirchoff), directed by dual-band beamsplitter (Z532/633, Chroma, Bellows Falls, USA) and then focussed by a water immersion objective (CFI PlanApochromat 60x WI, Nikon, Japan). The emitted fluorescence is collected through the objective and spatially filtered using a pinhole with 50 µm diameter and spectrally split into donor and acceptor channel by a single-edge dichroic mirror (BS640DCXR, Chroma, Bellows Falls, USA). Fluorescence emission was filtered (donor: FF01_582_75, Semrock, USA) acceptor: HQ685_70, Chroma, Bellows Falls, USA) and focused on avalanche photodiodes (SPCM-AQR 14, Perkin Elmer, Fremont, USA). The detector outputs were recorded by a TCSPC module (TimeHarp200, PicoQuant GmbH, Berlin, Germany).

Data analysis was performed using IgorPro 8 (Wavemetrics, Portland OR, USA). Single-molecule events were identified using a bin-selection algorithm with a threshold of 40 photons in the sum of donor and acceptor channel upon donor excitation and a threshold of 10 photons in the acceptor channel upon acceptor excitation.

**Lab#14**

Our multi-parameter fluorescence detection setup equipped with pulsed interleaved excitation is conceptually identical to the confocal microscope described previously[4]. For excitation, a pulsed supercontinuum laser was used with wavelength selector at 532±5 nm (Solea, Picoquant, Berlin, Germany), and a spectrally filtered (Chroma z635/10x, Picoquant) 635-nm laser diode (LDH-P-C-635B, Picoquant). Both lasers were alternated at 26.67 MHz (PDL 828 Sepia2, Picoquant), delayed ~18-ns with respect to each other and combined via a dichroic mirror (Chroma T560lpxr, F48-559, AHF) in a single-mode optical fiber (coupler: 60FC-4-RGBV11-47, fiber: PMC-400Si-2.6-NA012-3-APC-150-P, Schäfter und Kirchhoff GmbH, Hamburg, Germany). After collimation (60FC-L-4-RGBV11-47, SuK GmbH), the linear polarization was cleaned up (Codixx VIS-600-BC-W01, F22-601, AHF) and the light was reflected on a 3-mm thick excitation polychroic mirror (Laser Beamsplitter zt532/640rpc, F58-PQ09, AHF) upward and into the back port of the microscope (IX70, Olympus Belgium NV, Berchem, Belgium) via two mirrors and upward to the sample (3-mm thick Full Reflective Ag Mirror, F21-005, AHF, mounted in a TIRF Filter Cube for BX2/IX2, F91-960, AHF) to the objective (UPLSAPO-60XW, Olympus). Sample emission was focused through a 75-µm pinhole (P75S, Thorlabs, Munich, Germany) via an achromatic lens (AC254-200-A-ML, Thorlabs), collimated again (AC254-50-A-ML, Thorlabs) and spectrally split (Laser Beamsplitter H 643 LPXR superflat, F48-643, AHF). The yellow range was filtered (582/75 BrightLine HC, F37-582, AHF) and polarization was split (PBS251, Thorlabs). The red range was also filtered (Chroma ET705/100m, AHF) and polarization was split (PBS252, Thorlabs). Photons were detected on four avalanche photodiodes (yellow photons: Laser Components COUNT Blue, red photons: Perkin-Elmer or EG&G SPCM-AQR12/14) all of which were connected to a time-correlated single photon counting (TCSPC) device (SPC-630, Becker & Hickl GmbH, Berlin, Germany) over a router (HRT-82, Becker & Hickl) and power supply (DSN 102, Picoquant). Signals were stored in 12-bit first-in- first-out (FIFO) files. All analyses of experimental data were performed in the software package PAM[7].

**Lab#15**

All sample solutions were measured in Corning 384 well non-binding plates at concentrations ranging between 20-60 pM. The general scheme of the setup is described[79].In short: Single-molecule FRET experiments with PIE - MFD were performed on a homebuilt confocal microscope as described previously[79].Excitation was performed with a pulsed SC450-4-20Mhz supercontinuum source (Fianium, Southampton, UK). It runs at 20MHz, and has a power density >2mW/nm over the 450-800nm range, with an average pulse width of 100-150ps. The collimated, unpolarized output of the source is divided by a 50:50 beamsplitter cube (BS016, Thorlabs, NJ, USA), thus generating two beams. The beams are spectrally filtered using excitation bandpass filters at wavelength 532/10 (prompt beam) and 635/10 (delayed beam) to excite donor and acceptor molecules, respectively. The delayed beam has a path length of ~ 8m relative to the prompt beam, generating a ~24 ns delay in the pulse. The two beam paths are then recombined using a 50:50 beamsplitter cube (BS016, Thorlabs, NJ, USA) and focused using a 10x objective into a single-mode fiber (SMF) (P1-460A-FC, Thorlabs, NJ, USA), by which the beams become spatially overlapped and filtered. The output of the fiber is collimated using a 10x microscope objective lens (04OAS010; CVI Melles Griot, Albuquerque, NM, USA), polarized by a polarizing beamsplitter cube (PBS; PBS201, Thorlabs, NJ, USA) and coupled into an inverted microscope (Eclipse Ti, Nikon, France). Power was controlled using ½ (prompt: WPH05M-532 and delayed WPH05M-633, Thorlabs, NJ, USA) and ¼ waveplates (prompt: WPQ05M-532 and delayed WPQ05M-633, Thorlabs, NJ, USA) placed in the prompt and delayed beams before recombination, resulting in 50 µW for the prompt and 25 µW for the delayed beam at the entrance into the microscope. The light is reflected by a dichroic mirror that matches the excitation/emission wavelengths (FF545/650-Di01, Semrock, Rochester, NY, USA) and coupled into a Nikon 100x, NA1.4 objective. Emitted photons are then collected by the objective and focused by the tube lens on a pinhole of 150µm. The emission photon stream is collimated and divided using a polarizing beamsplitter cube (PBS; PBS201, Thorlabs, NJ, USA). In each created polarization channel, the photons are spectrally separated using dichroic mirrors (BS 649, Semrock, Rochester, NY, USA) and filtered using high quality emission filters (parallel: ET BP 585/65 and ET BP 700/75 and perpendicular: HQ 590/75 M and HQ 660 LP, Chroma, Bellows Falls, VT, USA). Single photons are detected using Single Photon Avalanche Diodes. We use two MPD-1CTC (MPD, Bolzano, Italy) for the donor wavelength channels and two SPCM AQR-14 (Perkin Elmer, Fremont, CA, USA) for the acceptor wavelength channels. The output of the detectors is coupled into a TCSPC counting board (SPC-150, Becker&Hickl, Berlin, Germany), through a HRT41 router (B&H), using appropriate pulse inverters and attenuators. The sync signal that triggers the TCSPC board is provided by picking a small fraction of the light from the prompt path (reflected by a coverslip), and focusing it on an avalanche diode (APM-400, B&H).

Data analysis was performed using the Software Package for Multiparameter Fluorescence Spectroscopy, Full Correlation and Multiparameter Fluorescence Imaging developed in C.A.M. Seidel's lab (http:// www.mpc.uni-duesseldorf.de/seidel/). A single-molecule event was defined as a burst containing of at least 40 photons with a maximum allowed interphoton time of 0.3 ms and a Lee-filter of 20. Photobleaching events were identified base on |TGX-TRR|<1 ms as described [4].

**Lab#16**

All sample solutions were measured in chamber with concentration 100 pM. The general scheme of the setup is similar to Supplementary Figure 2. In short: Single-molecule FRET experiments with ALEX were performed on a homebuilt confocal microscope as described previously. The light from 532nm (Compass 215M-50, Coherent, USA) and 638nm (25mW Red flame, Coherent, USA) cw-laser sources is alternately directed to an IX71 inverted microscope (Olympus, Japan) every 25 µs, reflected on a beamsplitter (Z488-533-633RPC, Chroma, USA) and focused through a water-immersion objective (UPlanApo 60x/1.2w, Olympus, Japan) to excite fluorescent molecules. The light intensities were 100 µW for 532nm and 35 µW for 638nm, measured before the beamsplitter. The emitted fluorescence is collected through the objective, spatially filtered using a 100 µm pinhole, and then spectrally split into two photon streams by a dichroic mirror (635 DCXR, Chroma, USA). Individual photon streams were filtered (for donor: HQ580/60m, for acceptor: HQ665lp, Chroma, USA) and detected by single photon-avalanche photodiodes (SPCM-AQR-14, PerkinElmer, USA). The detector outputs (photon arrival times) were recorded by a counter/timer device module (PCI-6602, National Instruments, USA).

Data analysis was performed using the ALEX-suite software package as described[80].Single-molecule events were identified using an all-photon-burst-search (APBS) and a dual-channel-burst-search (DCBS) algorithm with a threshold of 150, a time window of 50 µs and a minimum photon number of 50.

**Lab#17**

All sample solutions were measured in microscopy coverslip wells (µ-Slide 18 Well, Ibidi, GmbH) as was previously described in the analysis of smFRET within-burst dynamics of a DNA hairpin[81]. Sample excitation in all nsALEX/PIE measurements was achieved using pulsed picosecond fiber laser (λ = 532 nm, pulse width of 100 ps FWHM, operating at 20 MHz repetition rate and 100 μW measured at the back aperture of the objective lens; FL-532-PICO, CNI, China), and pulsed picosecond diode laser (λ = 642 nm, pulse width of 100 ps FWHM, operating at 20 MHz repetition rate and 60 μW measured at the back aperture of the objective lens; QuixX® 642-140 PS, Omicron, GmbH), delayed by 25 ns, for donor and acceptor excitation, respectively. Excitation path: (i) polarization maintaining optical fiber, (ii) beam shaping through quarter waveplate and a linear polarizer, (iii) dichroic beamsplitter with high reflectivity at 532 and 640 nm (ZT532/640rpc, Chroma, USA), (iv) high numerical aperture (NA) super apochromatic objective lens (60X, NA = 1.2, water immersion, Olympus, Japan). Emission path: (i) fluorescence was collected through the same objective lens, (ii) focused with an achromatic lens (f = 100 mm) onto a 100 μm diameter pinhole (variable pinhole, motorized, tunable from 20 μm to 1 mm), (iii) re-collimated with an achromatic lens (f = 100 mm), (iv) donor and acceptor fluorescence were split into two separate detection channels using a dichroic mirror with cutoff wavelength at λ = 652 nm (FF652-Di01-25x36, Semrock Rochester NY, USA), (v) donor and acceptor fluorescence were further filtered using 585/40 nm (FF01-585/40-25, Semrock Rochester NY, USA) and 698/70 nm (FF01-698/70-25, Semrock Rochester NY, USA) bandpass filters, respectively, and (vi) donor and acceptor fluorescence photons were detected using two hybrid photomultipliers (Model R10467U-40, Hamamatsu, Japan). Signal acquisition path: detectors were routed through a 4-to-1 router to a time-correlated single photon counting (TCSPC) module (SPC-150, Becker & Hickl, GmbH) as its START signal (the STOP signal is routed from the laser controller). Data acquisition was performed using the VistaVision software (version 4.2.095, 64-bit, ISS™, USA) in the time-tagged time-resolved (TTTR) file format, and then transformed into the photon HDF5 file format[82] for input in the FRET Bursts[11] analysis software. Data analysis: (i) the data was split into different photon streams, based on the detector they were originating from (donor or acceptor detectors) and on their photon nanotimes, whether it belonged to the donor or acceptor excitation time windows in nsALEX/PIE. The photon streams further used were $D_{ex.}D_{em.}$, $D_{ex.}A_{em.}$ and $A_{ex.}A_{em.}$ (ii) The background for each photon stream was assessed per each 30 s of acquisition. (iii) Photon bursts were identified as time periods were the instantaneous photon count rate of a sliding window of m=20 consecutive photons were at least F=6 times higher than the background rate. Both ACBS and DCBS were tested[83], where ACBS was used for attaining all data including donor- and acceptor-only bursts, and DCBS was used for attaining FRET-active bursts. (iv) After ACBS, bursts were further selected if they included at least 30 photons in all photon streams. After DCBS, bursts were further selected if they included at least 30 photons in the donor excitation photon streams, $D_{ex.}D_{em.}$, $D_{ex.}A_{em.}$, and at least 30 photons in the acceptor excitation stream, $A_{ex.}A_{em.}$. (v) Using the selected bursts after ACBS, donor-only and acceptor-only sub-populations were further selected, and then used for calculating the α and δ. (vi) Using the selected bursts after DCBS, the FRET-active bursts were further used for estimating the β and γ correction factors. (vii) burst-wise parameters, E and S values, as well as mean donor/acceptor nanotime for E-τ maps and E standard deviation for BVA were calculated per each burst and accumulated in 1D and 2D histograms. Sum of gaussian fitting was used for extracting the mean values of sub-populations of burst-wise parameters. All burst-wise parameters were corrected for background and the α, β, γ and δ correction factors.

**Lab#18**

All sample solutions were measured as a 50ul drop on a coverslip covered with a humidity chamber with a dilution of 1 to 1000 (sample 1) and 1 to 400 (sample 2 and 3) of the delivered stock solution. Measurements were performed on a home build confocal microscope, as previously described[84] . Using an ICHROME MLE-SFG laser module (Toptica, Germany) as excitation source, alternating 25 μs excitation pulses of 514 nm (80 μW) and 632 nm (80 μW). The excitation beam was directed via a fiber coupler and a dichroic mirror (z514/640rpc, Chroma, USA) through the water-immersion objective (60x, NA 1.2, Olympus, Japan), and focused 50 μm above the glass-sample interface. Fluorescence was spatially filtered with a 50 μm pinhole in the image plane and split by a second dichroic mirror (640dcxr, Chroma, USA). The fluorescence signals were further filtered (hq570/100nm and hq700/75nm, Chroma, USA) for green and red detection and focused on the active area of two single photon avalanche photodiodes (SPADs, SPCM AQR-14, Perkin Elmer, USA). The photodiodes were read out with a TimeHarp 200 photon counting board (Picoquant, Germany), and the arrival times were stored in t3r (time-tagged to timeresolved) files. Each dataset contained ~3000 bursts and was analyzed using the FRETBursts toolkit[11] .A single-molecule event was defined as a burst containing at least 40 photons with a maximum interphoton time of 100 μs.

**Lab#19**

Single-molecule FRET experiments were performed on a homebuilt confocal microscope as described previously[85]. Briefly, a confocal excitation laser (532 nm, World StarTech) was focused on the sample through the side port of the microscopy (Olympus) with a 100X oil immersion objective (Olympus). The excitation laser intensity was maintained at 50 μW for all experimental conditions. The fluorescence emission was collected through the objective, spatially filtered through a 100 μM pinhole, spectrally filtered through a band pass filter (HQ580/60 m, Chroma), split into donor and acceptor channels by a dichroic mirror (HQ680/60 m, Chroma) and subsequently detected by two avalanche photodiodes (Excelitas Technologies).

All sample solutions were measured in BSA (Bovine serum albumin, 1 mg/mL) passivated chambers with sample concentrations around 50 pM. Baseline fluorescence signal was estimated from imaging chamber containing imaging buffer only. Single molecule events were identified after subtraction of baseline signal, as bursts with minimum 40 counts over an exposure time of 1 ms.

### Supplementary Figures

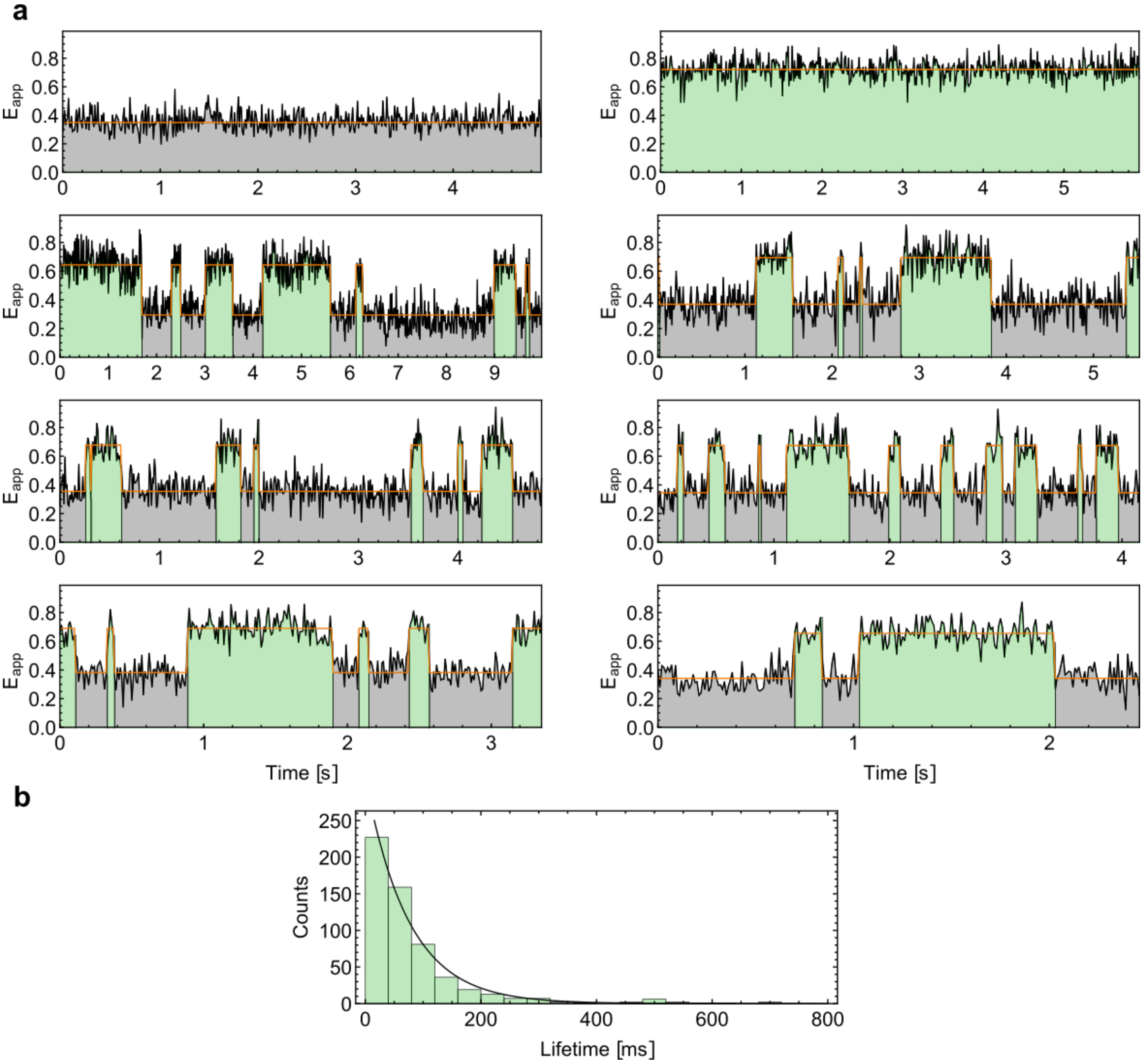

**Supplementary Figure 1: Ligand-induced slow conformational dynamics of MalE switching between the apo- and holo states. (a)** Time traces of immobilized MalE 36C/352C molecules labeled with Alexa555-Alexa647 measured in buffer containing no ligand (top row, left, grey), 1 mM maltose (top row, right, green) and 1 µM maltose , close to the $K_d$ (rows 2-4)[86]. The samples were measured in a scanning confocal microscope as described in reference [60]. FRET states and lifetimes are extracted from a fitted two-state Hidden-Markov-Model as described in reference [87]. The traces show ligand-induced interconversion of states on the >10 millisecond timescale. **(b)** A dwell time histogram of the duration of the holo state at 1 µM maltose with an exponential fit (solid line) shows a mean dwell time of 75 ms.

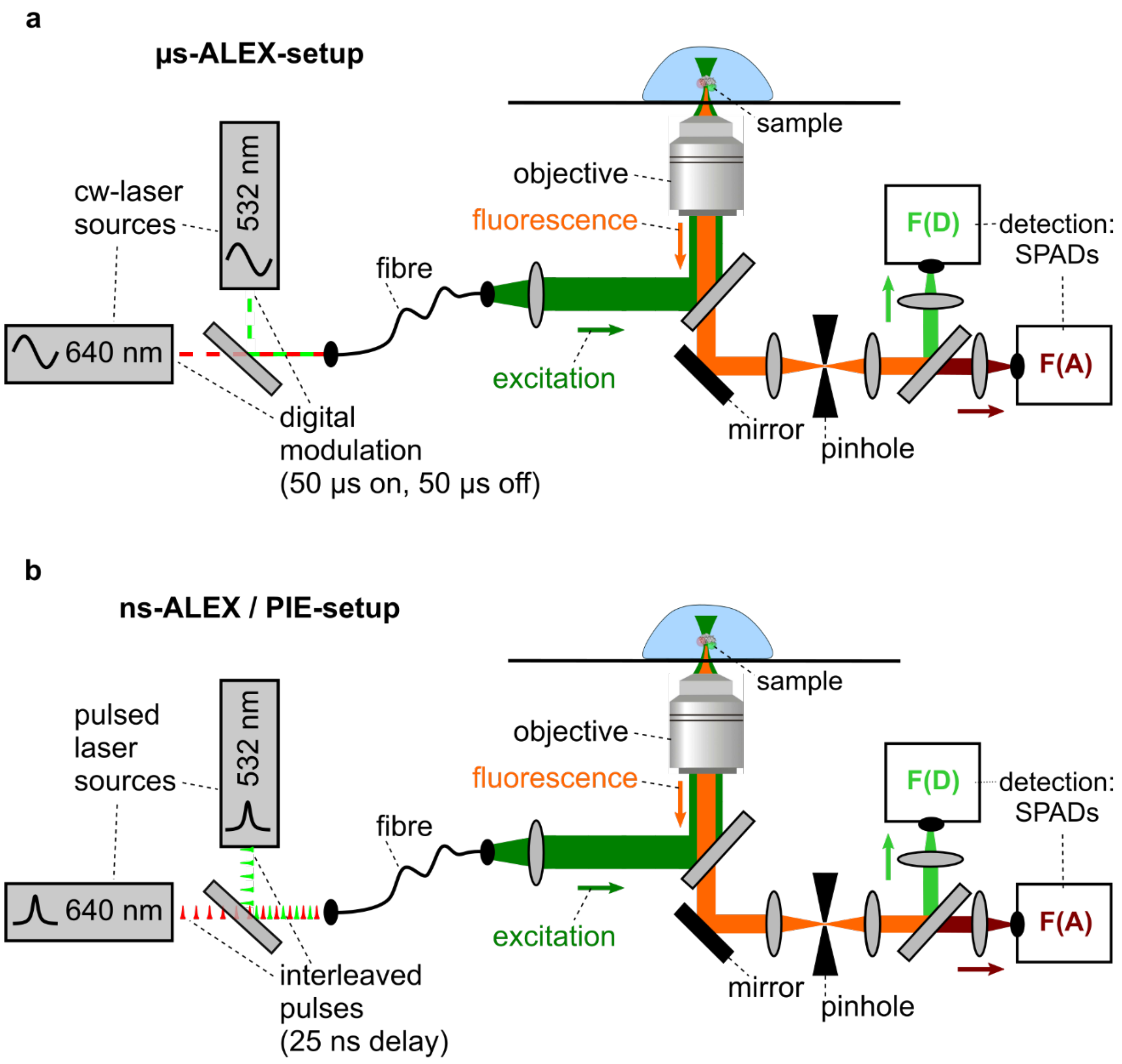

**Supplementary Figure 2: Schematics of the experimental setups. (a)** A schematic of a confocal microscope setup used for the acquisition of diffusion-based smFRET data using alternating-laser excitation (ALEX). Continuous wave laser sources (here with wavelengths of 532 nm and 640 nm) excite the sample alternatively for periods of 50 µs through a microscope objective. F(D) and F(A) indicate the donor and acceptor detection channels, respectively. **(b)** A schematic of a confocal microscope setup used for the acquisition of diffusion-based smFRET data using nsALEX / pulsed interleaved excitation (PIE). Pulsed laser sources (here with wavelengths of 532 nm and 640 nm) alternatingly excite the sample with picosecond pulses delayed by ~25 ns.

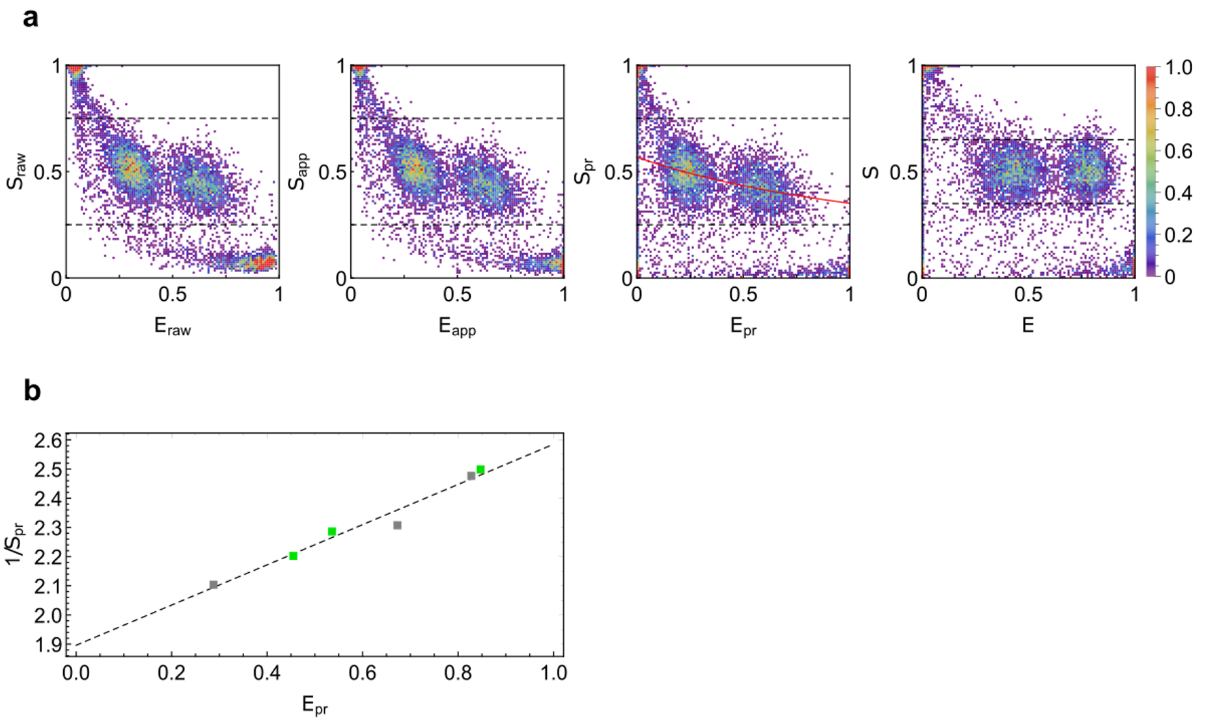

**Supplementary Figure 3: Correction procedure when using a global γ. (a)** *ES*-diagrams of two selected and merged data sets (Alexa546-Alexa647 labeled MalE-1 apo and MalE-2 apo) showing the results from the different correction steps; from left to right: raw data, background corrected apparent FRET efficiency, crosstalk and direct excitation corrected proximity ratio, $E_{pr}$, with fitted γ curve (red), and the γ corrected FRET efficiency versus stoichiometry plots. **(b)** Proximity ratio, $E_{pr}$ and stoichiometry $S_{pr}$ of all MalE mutants in the apo (gray dots) and holo states (green dots) with the linear fit (dashed line) used for a global γ correction. For more details for global γ estimation, please see Supplementary Note 2.

#### Supplementary Figure 4: Primer design and sequences for creation of MalE variants

The sequence of MalE (blue, DNA sequence in black) and the primers (in red) used for generating the mutants are given below. The procedures have been published previously[60].

```
   1                                                                   GTCGGTAAG
   1  K  I  E  E  G  K  L  V  I  W  I  N  G  D  K  G  Y  N  G  L  A  E  V  G  K
   1 AAAATCGAAGAAGGTAAACTGGTAATCTGGATTAACGGCGATAAAGGCTATAACGGTCTCGCTGAAGTCGGTAAG

  76 AAATTCGAGWRMGATACCGG 3' Lys29Cys
  26  K  F  E  K  D  T  G  I  K  V  T  V  E  H  P  D  K  L  E  E  K  F  P  Q  V
  76 AAATTCGAGAAAGATACCGGAATTAAAGTCACCGTTGAGCATCCGGATAAACTGGAAGAGAAATTCCCACAGGTT

  51  A  A  T  G  D  G  P  D  I  I  F  W  A  H  D  R  F  G  G  Y  A  Q  S  G  L
 151 GCGGCAACTGGCGATGGCCCTGACATTATCTTCTGGGCACACGACCGCTTTGGTGGCTACGCTCAATCTGGCCTG

 226                CCGGACAAAGCGTTCCAGKRCAAGCTGTATCCG 3' Asp87Cys
  76  L  A  E  I  T  P  D  K  A  F  Q  D  K  L  Y  P  F  T  W  D  A  V  R  Y  N
 226 TTGGCTGAAATCACCCCGGACAAAGCGTTCCAGGACAAGCTGTATCCGTTTACCTGGGATGCCGTACGTTACAAC

 101  G  K  L  I  A  Y  P  I  A  V  E  A  L  S  L  I  Y  N  K  D  L  L  P  N  P
 301 GGCAAGCTGATTGCTTACCCGATCGCTGTTGAAGCGTTATCGCTGATTTATAACAAAGATCTGCTGCCGAACCCG

 376             GAAGAGATCCCGKSSCTGGATAAAGAAC 3' Ala134Cys
 126  P  K  T  W  E  E  I  P  A  L  D  K  E  L  K  A  K  G  K  S  A  L  M  F  N
 376 CCAAAAACCTGGGAAGAGATCCCGGCGCTGGATAAAGAACTGAAAGCGAAAGGTAAGAGCGCGCTGATGTTCAAC

 151  L  Q  E  P  Y  F  T  W  P  L  I  A  A  D  G  G  Y  A  F  K  Y  E  N  G  K
 451 CTGCAAGAACCGTACTTCACCTGGCCGCTGATTGCTGCTGACGGGGGTTATGCGTTCAAGTATGAAAACGGCAAG

 526                      GTGGATAACKSYGGCGCGAAAGCG 3' Ala186Cys
 176  Y  D  I  K  D  V  G  V  D  N  A  G  A  K  A  G  L  T  F  L  V  D  L  I  K
 526 TACGACATTAAAGACGTGGGCGTGGATAACGCTGGCGCGAAAGCGGGTCTGACCTTCCTGGTTGACCTGATTAAA

 201  N  K  H  M  N  A  D  T  D  Y  S  I  A  E  A  A  F  N  K  G  E  T  A  M  T
 601 AACAAACACATGAATGCAGACACCGATTACTCCATCGCAGAAGCTGCCTTTAATAAAGGCGAAACAGCGATGACC

 226  I  N  G  P  W  A  W  S  N  I  D  T  S  K  V  N  Y  G  V  T  V  L  P  T  F
 676 ATCAACGGCCCGTGGGCATGGTCCAACATCGACACCAGCAAAGTGAATTATGGTGTAACGGTACTGCCGACCTTC

 251  K  G  Q  P  S  K  P  F  V  G  V  L  S  A  G  I  N  A  A  S  P  N  K  E  L
 751 AAGGGTCAACCATCCAAACCGTTCGTTGGCGTGCTGAGCGCAGGTATTAACGCCGCCAGTCCGAACAAAGAGCTG

 276  A  K  E  F  L  E  N  Y  L  L  T  D  E  G  L  E  A  V  N  K  D  K  P  L  G
 826 GCGAAAGAGTTCCTCGAAAACTATCTGCTGACTGATGAAGGTCTGGAAGCGGTTAATAAAGACAAACCGCTGGGT

 301  A  V  A  L  K  S  Y  E  E  E  L  A  K  D  P  R  I  A  A  T  M  E  N  A  Q
 901 GCCGTAGCGCTGAAGTCTTACGAGGAAGAGTTGGCGAAAGATCCACGTATTGCCGCCACCATGGAAAACGCCCAG

 976                                                                  GATCAACGCC
 326  K  G  E  I  M  P  N  I  P  Q  M  S  A  F  W  Y  A  V  R  T  A  V  I  N  A
 976 AAAGGTGAAATCATGCCGAACATCCCGCAGATGTCCGCTTTCTGGTATGCCGTGCGTACTGCGGTGATCAACGCC

1051 GCCWGCGGTCGTCAG 3' Ser352Cys
 351  A  S  G  R  Q  T  V  D  E  A  L  K  D  A  Q  T
1051 GCCAGCGGTCGTCAGACTGTCGATGAAGCCCTGAAAGACGCGCAGACT
```

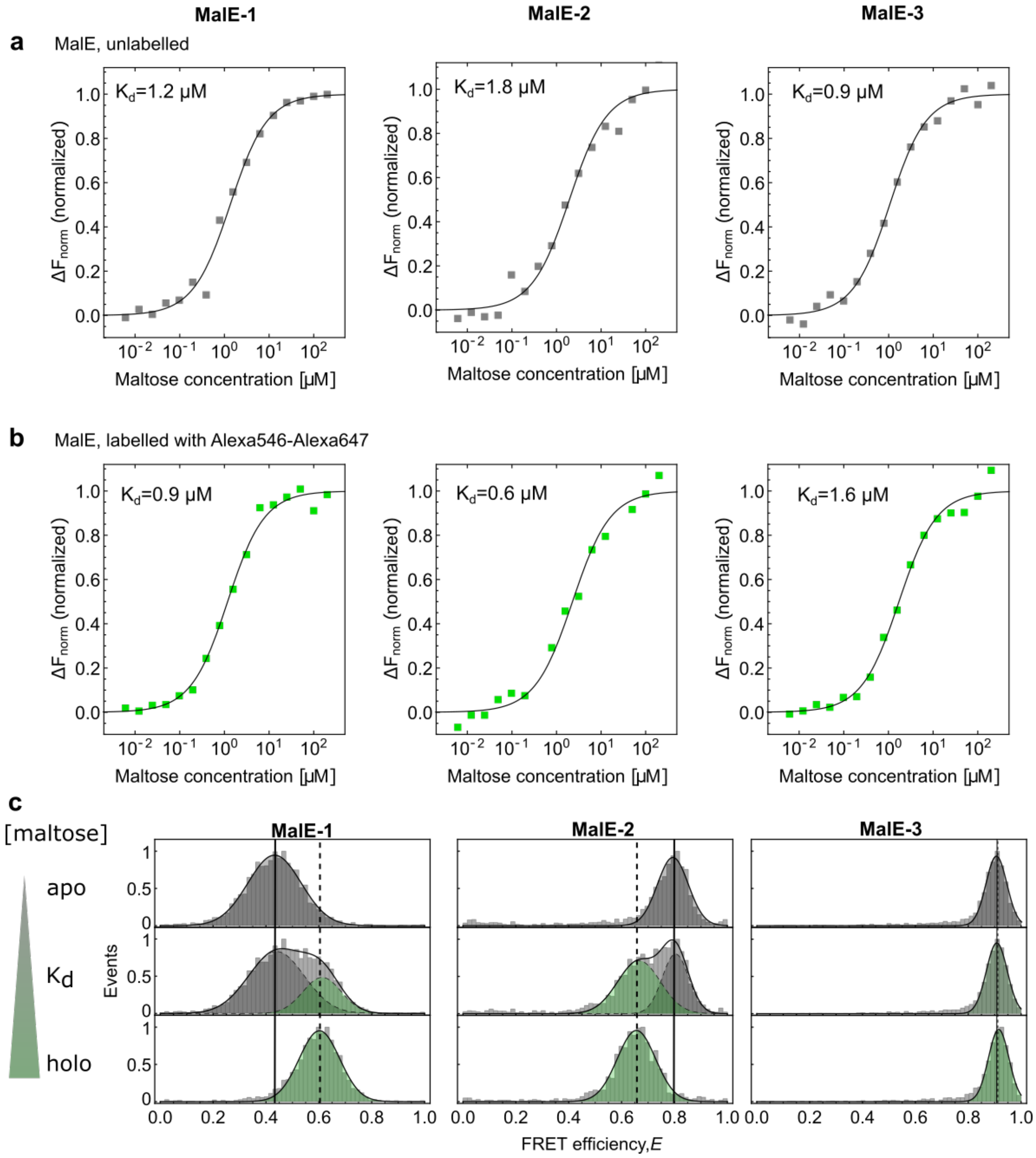

**Supplementary Figure 5: Binding affinity measurements of maltose to MalE using microscale thermophoresis. (a)** The binding affinities of maltose to MalE were measured using microscale thermophoresis (Monolith NT.LabelFree, Nanotemper) where the ratio of fluorescence before and after heating $\Delta F_{norm} = F_{cold}/F_{hot}$ was recorded at different maltose concentrations[88]. Data points show $\Delta F_{norm}$ normalized to the minimal and maximal fluorescence intensities for the unlabeled mutants MalE-1 (left), MalE-2 (middle), and MalE-3 (right). The curves were fitted with a standard model for receptor-ligand kinetics

$$\Delta F_{norm} = \frac{K_d + c_P + c_{malt} - \sqrt{(K_d + c_P + c_{malt})^2 - 4 c_P\, c_{malt}}}{2\, c_P},$$

where $K_d$ is the dissociation constant, $c_P$ the protein concentration (0.25 μM in the experiment), and $c_{malt}$ the maltose concentration. The fits to the binding model (solid line) yield $K_d$-values of 1.2 μM (left), 1.8 μM (middle), and 0.9 μM (right), respectively. This variation in the $K_d$ values is within the typical reproducibility of the measurement determined to be $K_d$ =1.2 ± 0.5 μM from measurements on19 different mutants. **(b)** The binding affinities of maltose to fluorescently-labeled MalE (Alexa547 and Alexa647) mutants MalE-1 (left), MalE-2 (middle), and MalE-3 (right) were measured using microscale thermophoresis. These experiments yielded $K_d$-values of 0.9 μM (left), 0.6 μM (middle), and 1.6 μM (right), respectively. **(c)** FRET efficiency $E$ histogram for the MalE mutant MalE-1 (left), the mutant MalE-2 (middle), and the mutant MalE-3 (right) in the presence of

0 (top), 1 μM (middle), and 1 mM maltose (bottom). The peak position of the apo (solid line) and holo (dashed line) populations, determined for this experiment, are shown. The data is from Lab #3.

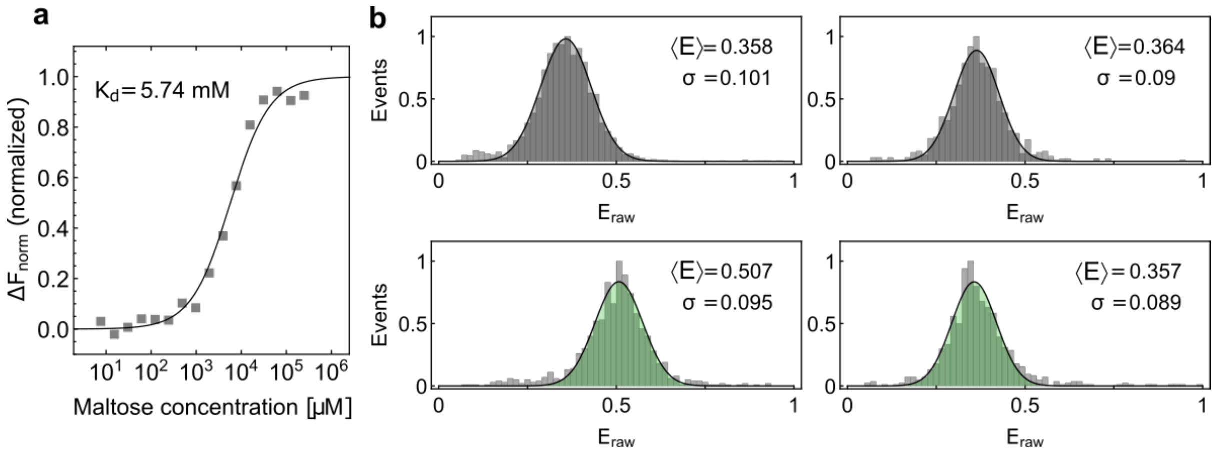

**Supplementary Figure 6: Millimolar maltose concentration does not influence fluorophore properties.** To investigate the potential influence of maltose on the photophysical properties of the used fluorophores, measurements were performed using the D65A closing deficient mutant of MalE-1. **(a)** The binding affinity of maltose to the MalE-1 mutant D65A were measured using microscale thermophoresis. The ratio of fluorescence before and after heating $\Delta F_{norm} = {}^{F_{cold}}/_{F_{hot}}$ was recorded at different maltose concentrations[88]. Data points show $\Delta F_{norm}$ normalized to minimal and maximal fluorescence intensities for the unlabeled mutants. The curves were fitted with a standard model for receptor-ligand kinetics:

$$\Delta F_{norm} = \frac{K_d + c_P + c_{malt} - \sqrt{(K_d + c_P + c_{malt})^2 - 4 c_P\, c_{malt}}}{2\, c_P},$$

where $K_d$ is the dissociation constant, $c_P$ the protein concentration (0.25 µM in the experiment) and $c_{malt}$ the maltose concentration. A $K_d$ of 5.7 mM was measured. **(b)** Raw FRET efficiency $E_{raw}$ histograms for the mutant MalE-1 labeled with Alexa546-Alexa647 without maltose (top, left) and with added 1 mM maltose (bottom, left) compared to same measurement with the MalE-1 closing deficient mutant D65A (right). No influence of the maltose is visible in the FRET efficiency histogram for the closing deficient mutant.

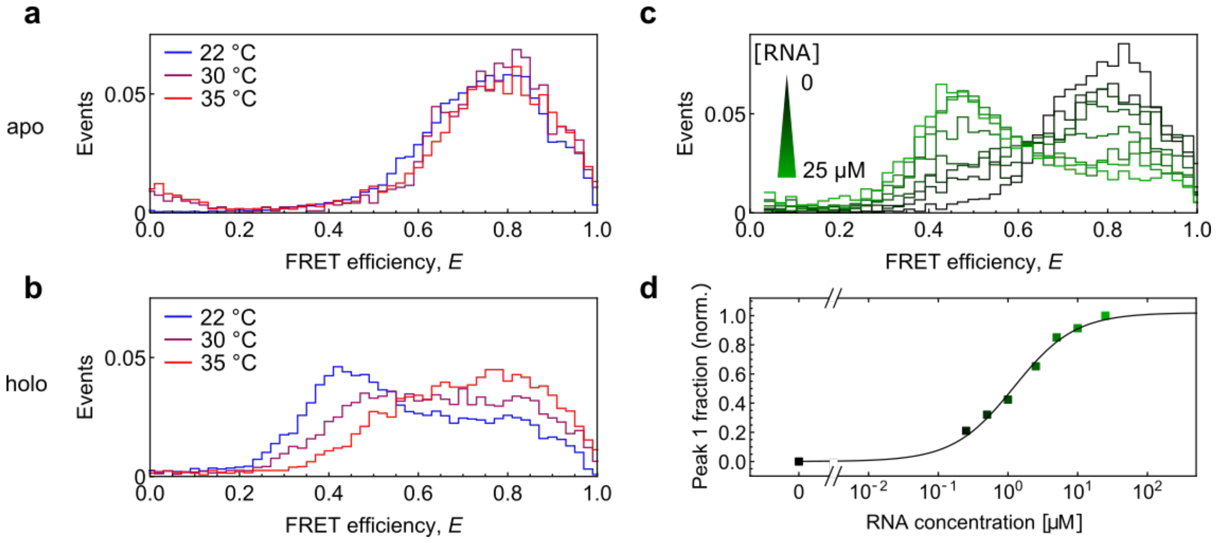

**Supplementary Figure 7: Temperature and concentration dependence of RNA binding to U2AF2.** **(a, b)** SmFRET histograms of **(a)** apo and **(b)** holo measurements at 22°C (blue), 30°C (purple) and 35°C (red). **(c)** SmFRET histograms for U9 RNA titration measurements with U2AF2 (low to high RNA concentrations are shown in a color gradient from black to light green (0, 0.25, 0.5, 1.0, 2.5, 5, 10 and 25 µM). **(d)** The area under peak 1 of the FRET histograms (0.1-0.6 FRET efficiency) from panel c is plotted versus the U9 RNA concentration to estimate the $K_d$. For normalization, the area for the apo measurement was set to zero and for the holo measurement at 25 µM was set to 1. The affinity of U9 RNA binding to U2AF62 was estimated to be ~1.2 µM using the standard model for receptor-ligand kinetics as described in Supplementary Figure 5.

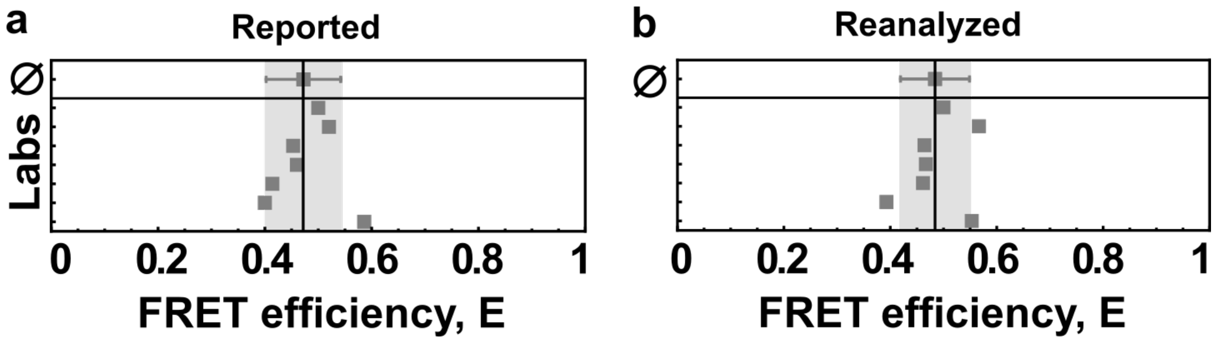

**Supplementary Figure 8: Comparison of the FRET efficiency reported for different labs and after reanalysis for the MalE-1 mutant under apo conditions. (a, b)** The determined average FRET efficiency values (**a**) reported by the different laboratories and (**b**) after reanalysis are shown as squares for measurements of MalE-1 in the apo states for 7 laboratories. The mean value (upper data point) from all data sets with the corresponding standard deviation is shown in grey. For the details of the reanalysis procedure, please refer to the Supplementary Note 3. The mean FRET Efficiency and standard deviation was 0.474 and 0.063 for the reported values and 0.487 and 0.054 after reanalysis respectively. For reanalysis purposes, we used data from the same 7 labs which measured the dynamics of U2AF.

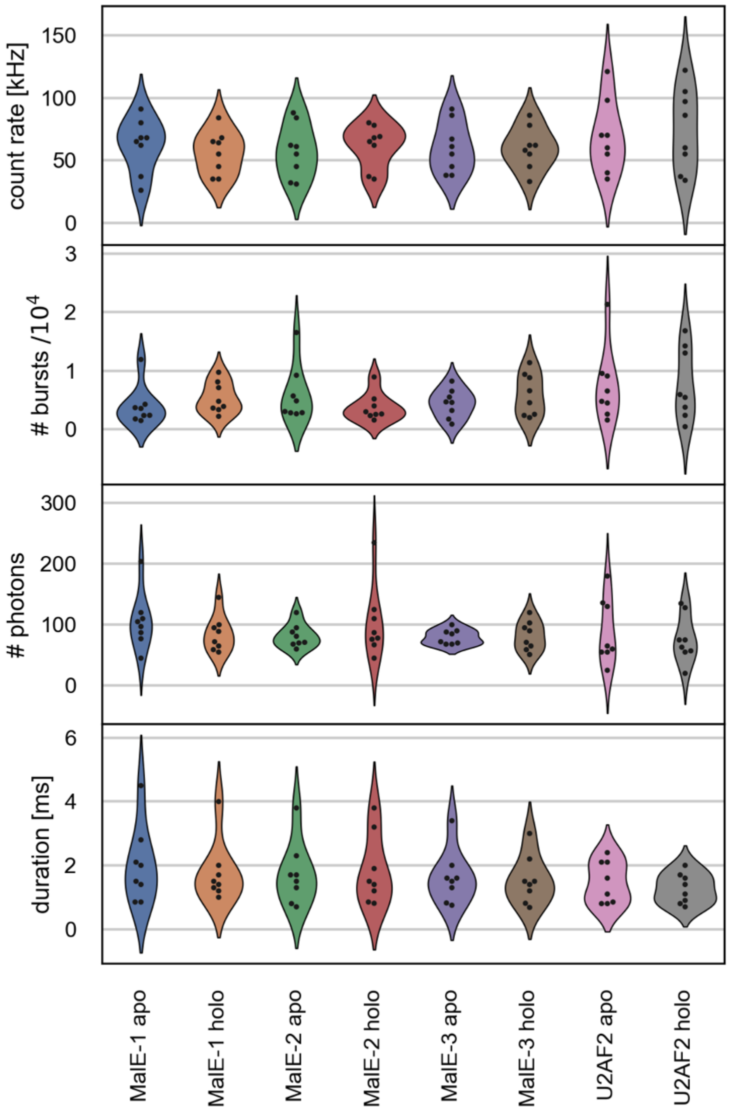

**Supplementary Figure 9: Sample-dependent setup statistics**. Sample-dependent distributions of the setup-dependent parameters are given as violin plots for (*top*) the average count rate, (*second from the top*) number of detected bursts, (*third from the top*) average number of photons per burst and (*bottom*) average burst duration. The procedure to obtain the values for the above parameters from the measurement data collected from 8 labs for both MalE and U2AF samples is as follows: The collected raw data from different labs was analyzed using the PAM software[77]. Briefly, first, a burst search was performed using an all photon burst search with a threshold of 50-100 photons per sliding time window of 500 µs depending on the dataset. For one set of measurements, a lower threshold of 20 photons per 500 µs time window was necessary. After burst selection, background subtraction and correction for crosstalk and direct excitation were performed as discussed in the data analysis section. To remove blinking and bleaching events as well as for selecting out the double-labeled molecules, an ALEX-2CDE filter[8] with a lower limit of 5 and an upper limit of 25 was used depending on the data set. The ALEX-2CDE filter values differed depending on the excitation intensities and sample concentrations used for the measurements. The values for all the parameters are a median of the values obtained for the double-labeled molecules for each measurement. These values were made available with PAM software.

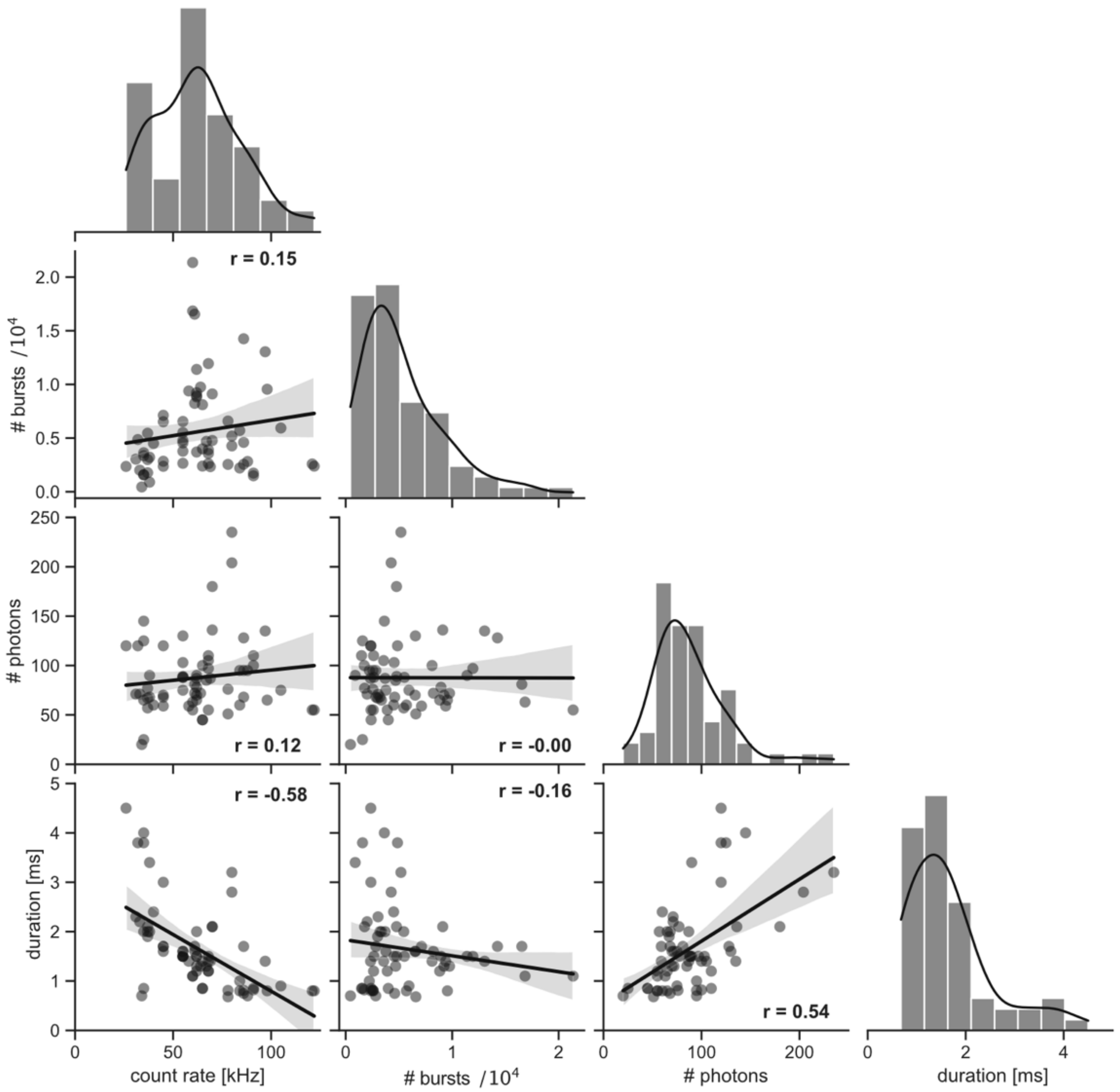

**Supplementary Figure 10: Correlations between setup-dependent parameters.** Pairwise plots of the number of detected bursts, average number of photons per burst, average burst duration and average burstwise count rate shown in Supplementary Figure 9. $r$ is the Pearson's correlation coefficient. The one-dimensional projections show the distribution of the parameters as histograms (gray bars) and kernel density estimates (black lines).

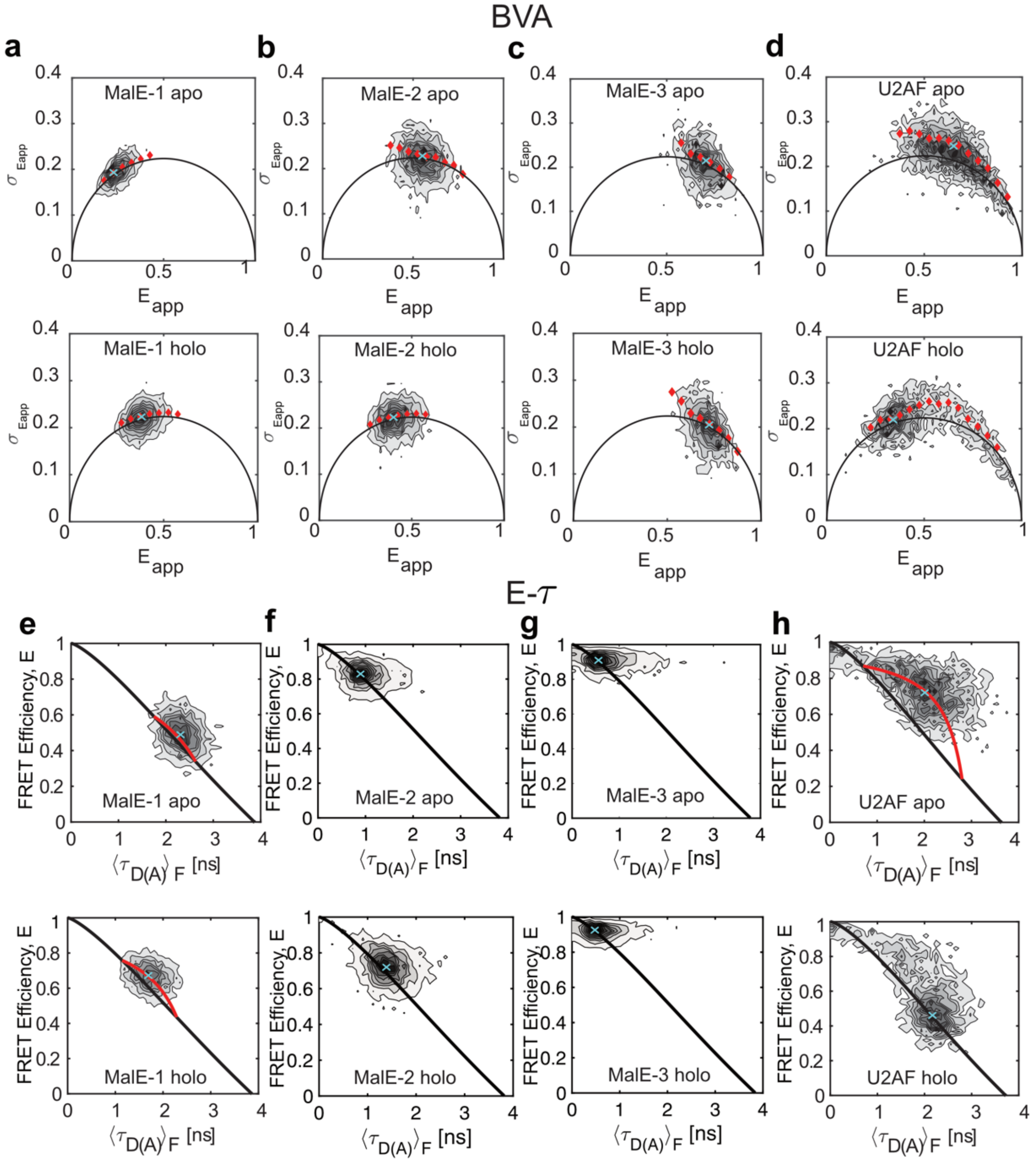

**Supplementary Figure 11: Overview of conformational dynamics and determination of the dynamic shift on the sub-millisecond timescale in MalE labeled with Alexa546-Alexa647 and U2AF2 labeled with Atto532-Atto643. (a-d)** Burst variance analysis (BVA) of MalE-1 (a), MalE-2 (b), MalE-3 (c), and U2AF (d) under both apo (upper panel) and holo (lower panel) conditions. In BVA, the standard deviation $\sigma_{E_{\text{app}}}$ of the apparent FRET efficiency $E_{\text{app}}$ is compared to the shot-noise limit given by $\sigma_{E_{\text{app}}} = \sqrt{E_{\text{app}}(1 - E_{\text{app}})/n}$ (black line, here $n = 5$). Single-molecule events with conformational dynamics show an increased variance and follow the dynamic line (red diamonds). Red diamonds indicate the average standard deviation of all bursts within a FRET efficiency range of 0.05. The mean positions of the populations (cyan crosses) were determined by fitting a two-dimensional Gaussian distribution to the data (see Supplementary Note 5). The dynamic shift, *ds*, is defined as the excess standard deviation compared to the static line. A clear deviation from the static line is observed for the apo state of U2AF2. For U2AF2 under holo conditions, the *ds* was determined for the major holo state population. Please note the leftover minor apo state population in holo condition has a similar deviation as for the apo condition. **(e-h)** Plots of the FRET efficiency $E$ versus intensity-weighted average donor lifetime $\langle\tau_{D(A)}\rangle_F$ (E-$\tau$) for MalE-1 (e), MalE-2 (f), MalE-3 (g), and U2AF (h) under both apo (upper panel) and holo (lower panel)

conditions. In the E-$\tau$ plot, the intensity-based FRET efficiency $E$ is plotted against the intensity-weighted average donor fluorescence lifetime, $\langle\tau_{D(A)}\rangle_F$. The static FRET-line is given by the Förster relation as $E = 1 - \frac{\langle\tau_{D(A)}\rangle_F}{\tau_{D(0)}}$ (black). The static lines are slightly curved as they account for the flexibility of the dye linkers. Molecules undergoing dynamics are shifted from the static line and follow a dynamic FRET-line (red). The mean positions of the populations (cyan crosses) were determined by fitting a two-dimensional Gaussian distribution to the data (see Supplementary Note 5). For a given population, the dynamic shift is defined as the displacement of the population orthogonal to the static FRET-line. A clear dynamic shift is observed for U2AF2. The slight dynamic shift observed in MalE-1 is due to the high anisotropy of attached dyes as identified later in the study (see Fig. 5e,f and Supplementary Figure 12). The end points of the dynamic FRET-line for MalE-1 and U2AF2 were determined from a sub-ensemble analysis of the fluorescence decay. For U2AF2 under holo conditions, the *ds* was determined for the major holo state population. Please note that the leftover minor apo state population in the holo measurement has a similar deviation as for the apo condition.

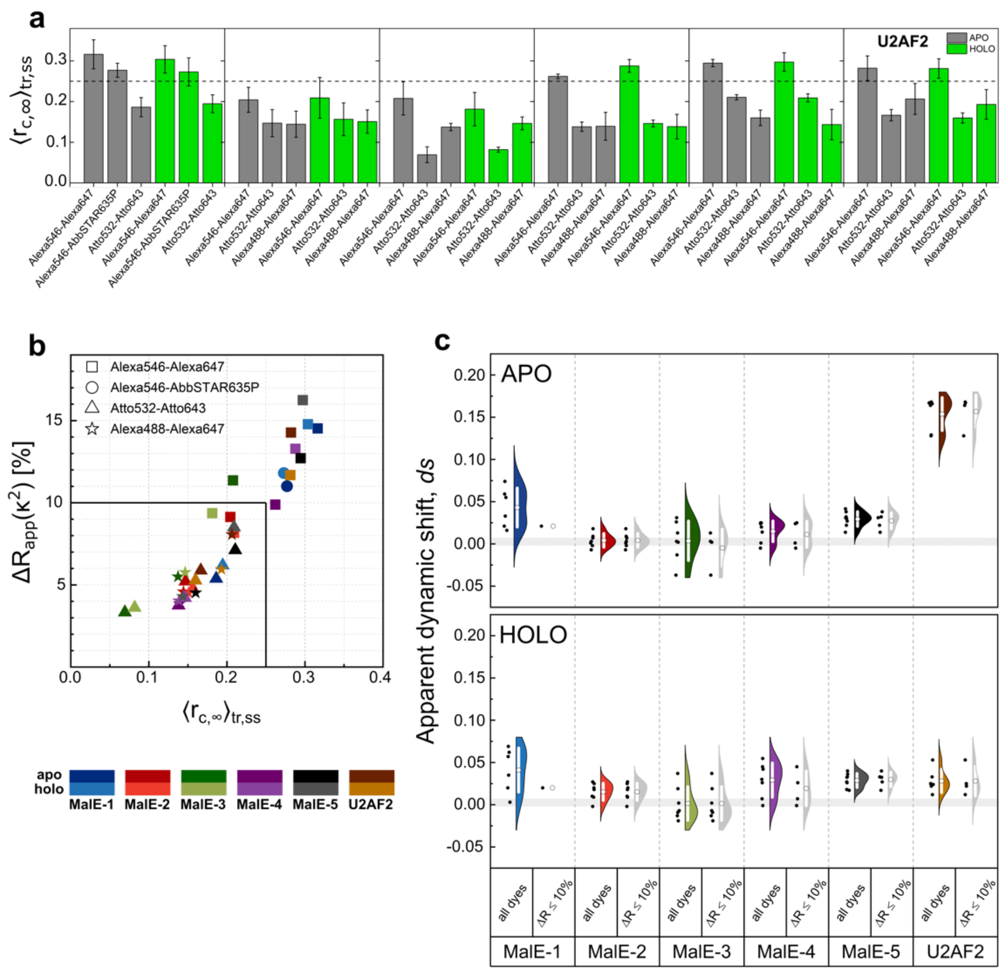

**Supplementary Figure 12: Dye-specific sticking interactions in proteins and residual apparent dynamic shift upon removal of sticky dyes. (a)** Combined residual anisotropies for different dye combinations of MalE and U2AF2 variants (see Supplementary Note 8 and Supplementary Table 13) **(b)** The distance uncertainty due to the orientation factor $\kappa^2$, $\Delta R_{\text{app}}(\kappa^2)$ (see Supplementary Note 9 and Supplementary Table 14), and its correlation with the combined residual anisotropy $\langle r_{c,\infty}\rangle_{tr,ss}$. A threshold of 10% in terms of distance uncertainty (solid black line) corresponds to $\langle r_{c,\infty}\rangle_{tr,ss}$= 0.25, which was used to filter-out sticking-prone dyes. **(c)** Violin plots of the dynamic shift values reported by three labs (see Supplementary Table 8) including all measured dye combinations and after filtering of the dyes to remove those with pronounced sticking interactions using a threshold of $\Delta R_{\text{app}}(\kappa^2) = 10\%$ corresponding to $\langle r_{c,\infty}\rangle_{tr,ss}$ = 0.25. In the violins, the white circle and whisker denote the mean ± 1σ. The shaded grey area gives the upper and lower limit of apparent dynamic shift obtained from DNA samples with different FRET efficiencies and with different dye combinations (Supplementary Table 8).

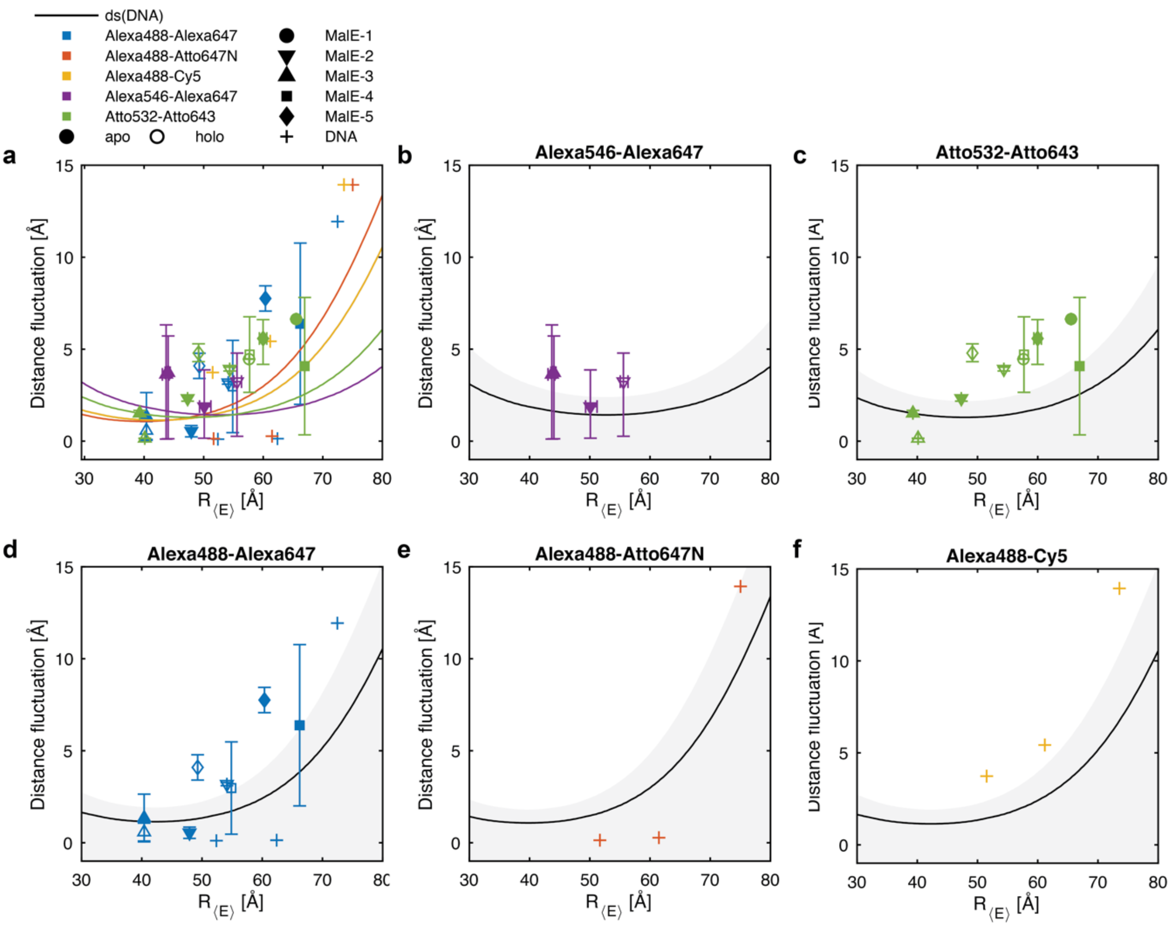

**Supplementary Figure 13: Estimated conformational flexibility of MalE based on the residual dynamic shift for different donor-acceptor pairs. (a)** The estimated distance fluctuations are plotted against the FRET-averaged interdye distance for the give MalE mutants in the apo and holo state labeled with the dye pairs Alexa546-Alexa647, Atto532-Atto643 and Alexa488-Alexa647. Additional control measurements on dsDNA are shown as crosses for the dye pairs Alexa488-Alexa647, Alexa488-Atto647N and Alexa488-Cy5. The lines indicate the apparent distance fluctuation obtained for the dsDNA control measurements, calculated based on a dynamic shift of $ds_{DNA} = 0.0026 \pm 0.0044$. **b-f)** Individual plots of the data shown in panel a for the different dye combinations. Gray areas indicate the 1$\sigma$ confidence interval for the apparent distance fluctuations obtained for the dsDNA control measurements. All values are given in Supplementary Table 8.

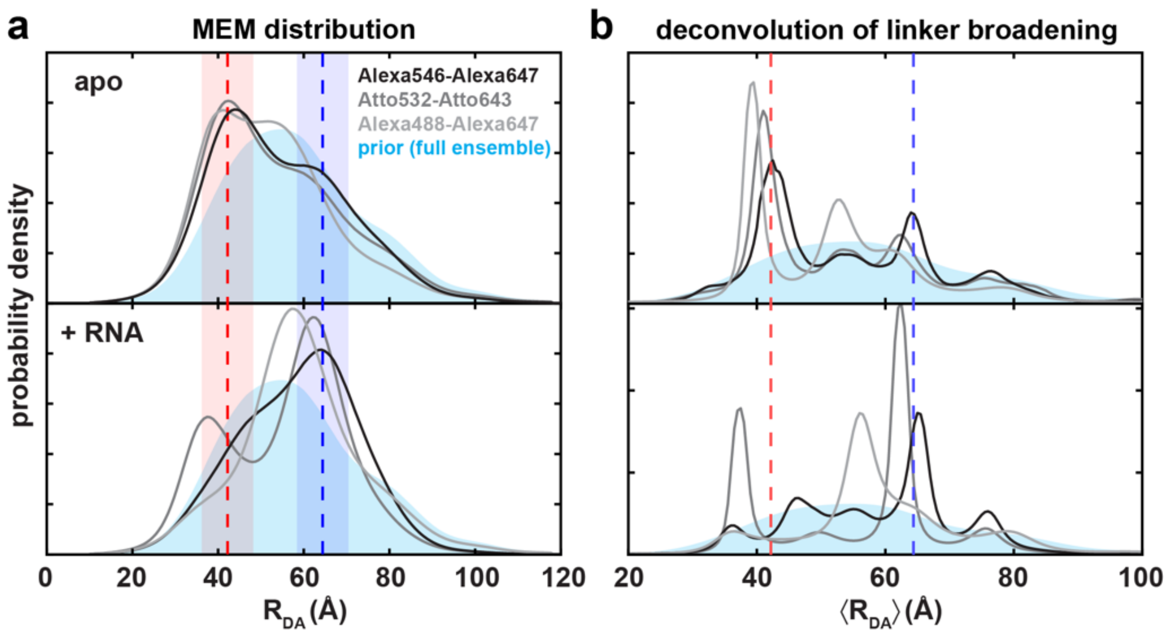

**Supplementary Figure 14: Comparison of the distance distributions obtained using different donor-acceptor dye pairs.** Comparison of the distance distributions obtained using different donor-acceptor dye pairs. **(a)** The donor-acceptor distance $R_{DA}$ distribution determined from the donor fluorescence decay using a model-free MEM approach for different dye-pairs where the distance distribution of the NMR/SAXS full ensemble of structures[44] (shown in light blue) was used as the prior distribution. The expected interdye distances for the resolved structure of the compact apo and open holo states are shown as red and blue dashed lines (PDB: 2YH0, 2YH1) with the shaded areas indicating the distance broadening due to the flexible dye linkers of 6 Å. This panel is identical to Fig. 6d. **(b)** The distribution of the mean donor-acceptor distance $R_{DA}$ obtained by the MEM in panel a deconvolved as described in Supplementary Note 15 to remove the broadening of the linker. Well-defined peaks at distances similar to the compact apo (red dashed line) and open-holo (blue dashed line) conformations become clearly visible.

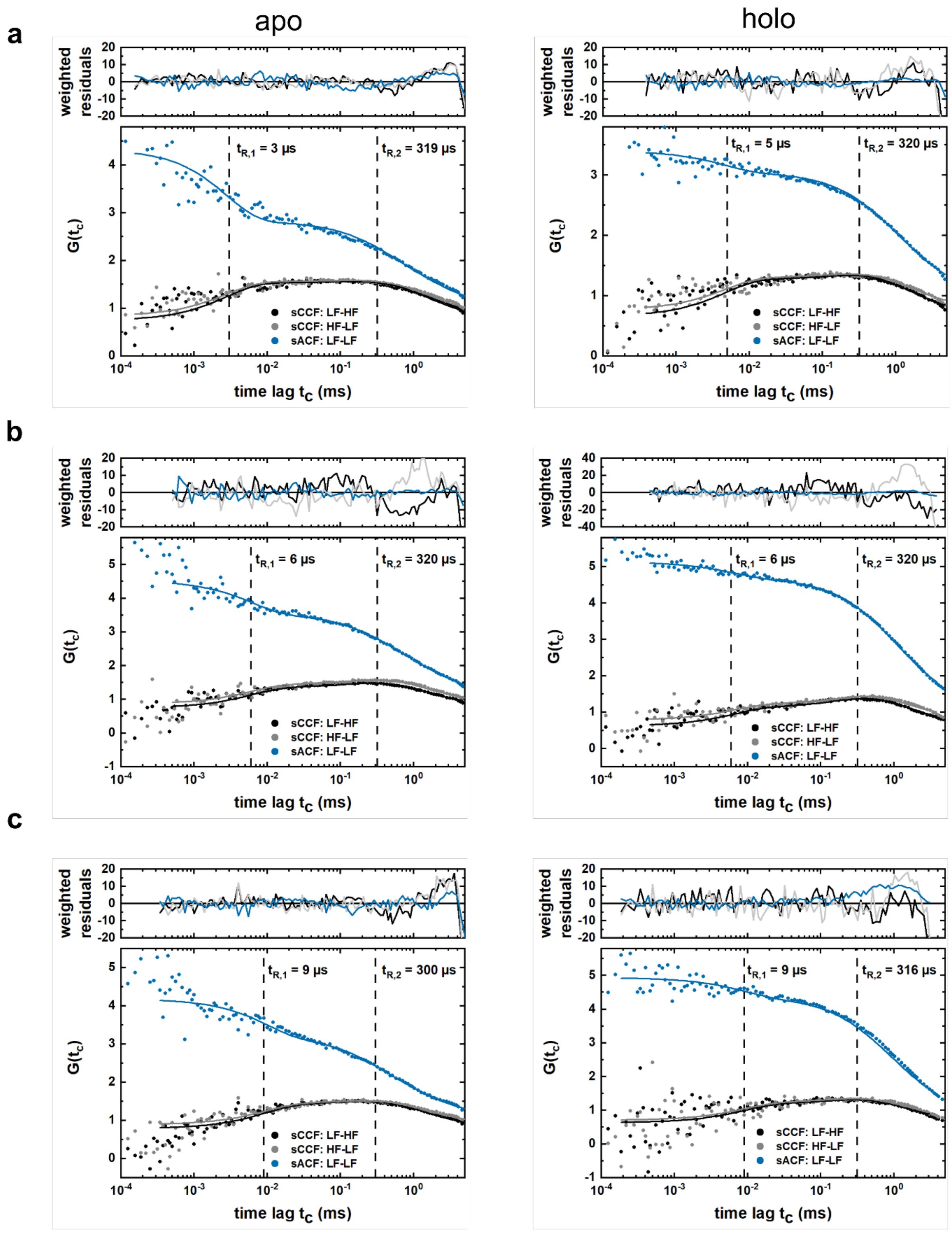

**Supplementary Figure 15: Filtered fluorescence correlation spectroscopy curves for U2AF2.** fFCS curves are shown for three different dye combinations, namely (a) Alexa546-Alexa647 (b)Atto532-Atto643 (c) Alexa488-Alexa647. In the analysis, a global fit of the two species autocorrelation functions, sACF, and two species cross-correlation functions, sCCF was performed. For simplicity, only one sACF function is shown. The fit model consisted of a diffusion term and two kinetic terms with corresponding relaxation times, $t_{R,1}$ and $t_{R,2}$ (Supplementary Note 16). The obtained relaxation times (denoted as vertical lines) are consistent between different dye combinations as well as between apo and holo states.

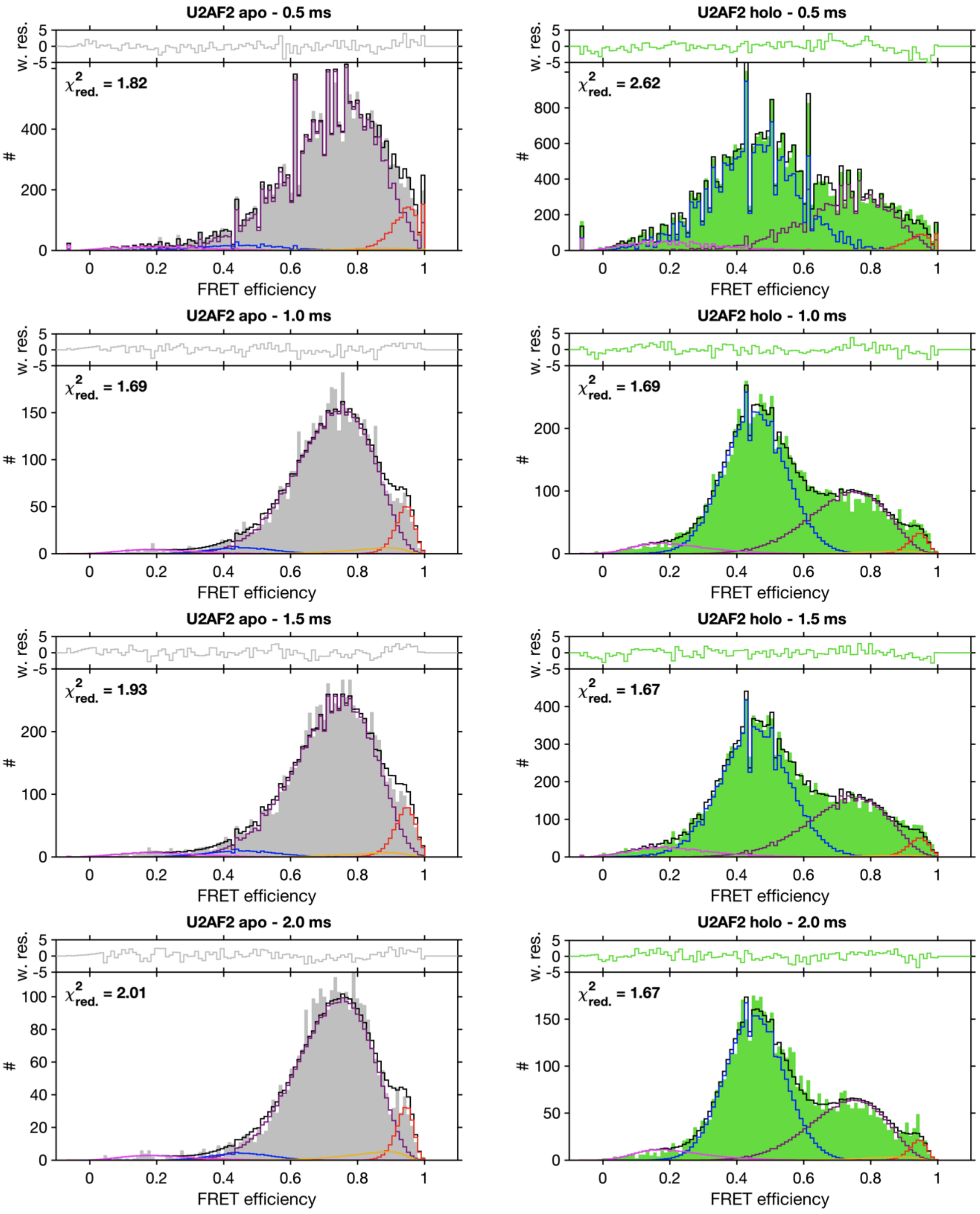

**Supplementary Figure 16: A global dynamic photon distribution analysis (PDA) of apo and holo measurements of U2AF2 labeled with the Atto532-Atto643 dye pair.** Different binning times (from top to bottom: 0.5 ms, 1 ms, 1.5 ms and 2.0 ms) for *left*: apo measurements (in grey) and *right*: holo measurement (in green) are shown. The analysis was performed globally over the apo and holo measurements using integration windows of 0.5, 1.0, 1.5 and 2.0 ms and shared distances for the compact apo conformation (red), the detached apo ensemble (purple) and the holo state (blue). The dynamic interconversion between the compact apo state and apo conformational ensemble is shown in yellow. An additional low-FRET population (magenta) had to be included, which most likely originates from photobleaching. All distances and kinetic rates for the apo ensemble are globally optimized while the amplitudes of the apo and holo populations were kept constant within the integration time windows for the apo and holo states. See Supplementary Note 17 for details.

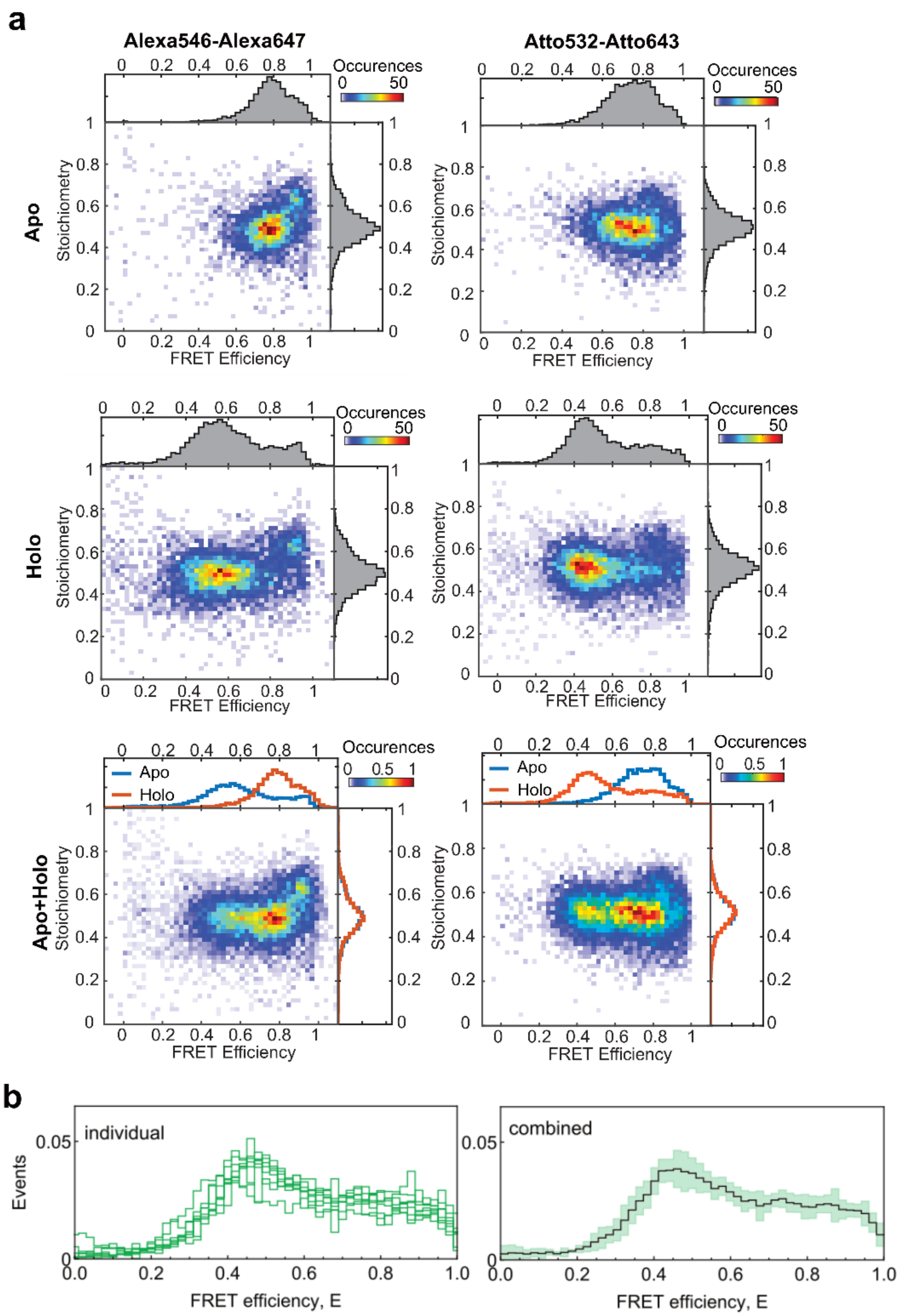

**Supplementary Figure 17: The very-high-FRET population in U2AF2. (a)** FRET efficiency versus stoichiometry plots are shown for two different dye combinations (Alexa546-Alexa647, left panel and Atto532-Atto643, right panel). These are plotted for apo (upper graph), holo (middle graph) and combined both apo and holo (lower graphs). A very-high-FRET populations with a different stoichiometry is visible in all measurements but with different amplitudes. **(b**) The FRET efficiency histograms (left) from the individual laboratories and (right) the combined histogram showing the mean (solid line) and a standard deviation (pale) after avoiding the subpopulation with slight acceptor quenching by using only bursts with a stoichiometry between 0.2-0.5 to build the smFRET histograms measured by the different labs.

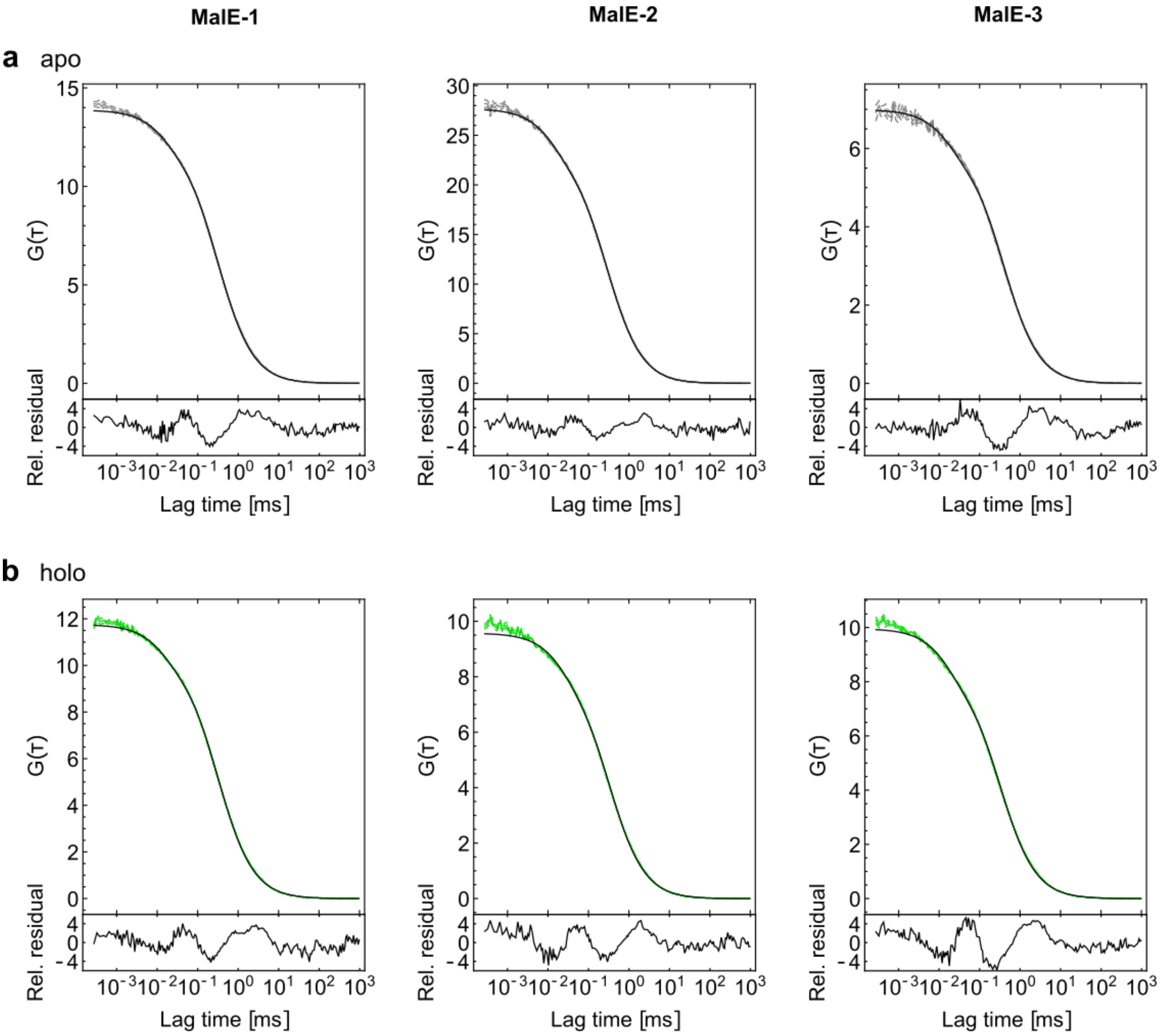

**Supplementary Figure 18: FCS experiments to rule out the presence of large aggregates.** Fluorescence Correlation Spectroscopy (FCS) measurements for the mutants MalE-1 (left), MalE-2 (middle), and MalE-3 (right) in **(a)** the apo state and **(b)** in the holo state are plotted. The two orthogonally oriented polarizations in the donor detection channel were correlated to remove the detector afterpulsing at short lag times (green line). All correlation curves were fitted with a standard model including a triplet fraction (black line):

$$G(t) = \frac{\gamma}{\langle N \rangle}\left(1 + \frac{t}{\tau_{\text{diff}}}\right)^{-1}\left(1 + \frac{t}{\tau_{\text{diff}} p^2}\right)^{-\frac{1}{2}}\left(1 + \frac{T}{1-T} e^{-\frac{t}{\tau_{\text{trip}}}}\right),$$

where $T$ is the triplet fraction, $\tau_{\text{trip}}$ the triplet lifetime, $p$ is the structural or elongation factor of the confocal volume given as the ratio of the axial and lateral dimensions, $\tau_{\text{diff}}$ is the diffusion time, and $\gamma = 2^{-3/2}$ is the geometric factor[89]. The confocal instrument was calibrated using free Alexa546 dye with a published[90] diffusion coefficient of 341 $\mu m^2 s^{-1}$ for which a diffusion time of 95±10 µs was found. The overall diffusion time for the six measurements was 325±40 µs. None of the correlation curves show any indication of the presence of protein aggregates.

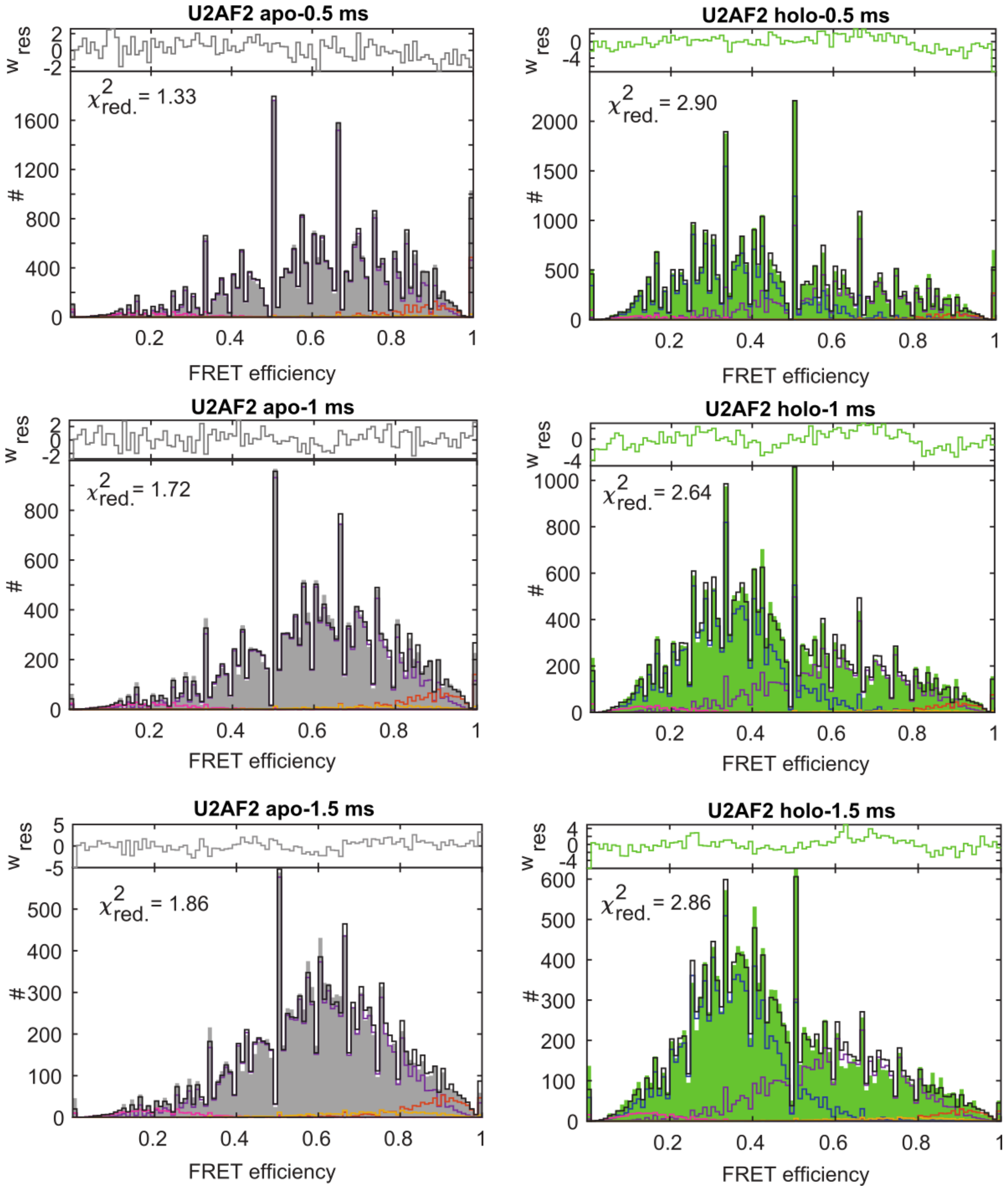

**Supplementary Figure 19: A global dynamic photon distribution analysis (PDA) of the apo and holo measurements of U2AF2 labeled with the Alexa546-Alexa647 dye-pair.** Left: apo measurement (in grey), right: holo measurement (in green). The analysis was performed globally over the apo and holo measurements for integration windows of (*top*) 0.5 ms, (*middle*) 1.0 ms, and (*bottom*) 1.5 ms using shared distances for the compact apo conformation (red), the detached apo ensemble (purple) and the holo state (blue). The dynamic interconversion between the compact apo state and the apo conformational ensemble is shown in yellow. All distances and kinetic rates for the apo ensemble are globally optimized while the amplitudes of the apo and holo populations were kept constant within the integration time windows for the apo and holo states. For details, see Supplementary Note 17.

### Supplementary Tables

**Supplementary Table 1: FRET efficiency correction factors reported by the participating labs for MalE constructs labeled with Alexa546-Alexa647.** Overview of all correction factors for MalE and the resulting change in the FRET efficiency upon application of the correction factors is shown as an example for the holo state of the MalE-2 mutant. The correction factors: α is spectral crosstalk of donor fluorescence into the acceptor channel, ß is the normalization of direct donor and acceptor excitation fluxes, $\gamma$ is the differences in donor and acceptor quantum yields and detection efficiencies and δ is the ratio of indirect and direct A excitation. $\langle E_{app} \rangle$ is the uncorrected FRET efficiency and $\langle E \rangle$ is the corrected FRET efficiency.

| Lab# | $\alpha$ | $\beta$ | $\gamma$ | $\delta$ | $\langle E_{app} \rangle$ | $\langle E \rangle$ |
|---|---|---|---|---|---|---|
| 1 | 0.03 | 2.38 | 0.23 | 0.08 | 0.43 | 0.72 |
| 2 | 0.05 | 0.5 | 0.34 | 0.32 | 0.54 | 0.73 |
| 3 | 0.05 | 1.70 | 0.51 | 0.07 | 0.56 | 0.66 |
| 4 | 0.04 | 2.65 | 0.31 | 0.06 | 0.42 | 0.66 |
| 5 | 0.04 | 1.56 | 0.34 | 0.14 | 0.54 | 0.72 |
| 6 | 0.08 | 1.59 | 0.54 | 0.12 | 0.57 | 0.66 |
| 7 | 0.07 | 0.64 | 0.63 | 0.12 | 0.59 | 0.70 |
| 8 | 0.04 | 1.53 | 0.48 | 0.11 | 0.47 | 0.63 |
| 9 | 0.06 | 1.84 | 0.47 | 0.10 | 0.57 | 0.69 |
| 10 | 0.03 | 1.88 | 0.31 | 0.11 | 0.46 | 0.72 |
| 11 | 0.03 | 2.37 | 0.32 | 0.07 | 0.43 | 0.64 |
| 12 | 0.04 | 1.99 | 0.25 | 0.07 | 0.47 | 0.74 |
| 13 | 0.06 | 0.60 | 0.34 | 0.32 | 0.58 | 0.76 |
| 14 | 0.04 | 2.00 | 0.23 | 0.09 | 0.41 | 0.77 |
| 15 | 0.05 | 1.42 | 0.33 | 0.05 | 0.57 | 0.79 |
| 16 | 0.05 | 1.31 | 0.46 | 0.14 | 0.60 | 0.72 |
| 17 * | 0.06 | 1.26 | 0.55 | 0.06 | | |
| 18 ** | 0.01 | 4.86 | 0.09 | 0.10 | 0.18 | 0.59 |
| 19 *** | 0.18 | | | | 0.59 | |

*Due to measurement problems with a large bleaching contribution, data could not be fitted with a Gaussian distribution (the FRET population was small with most bursts containing photobleaching events). Data were not considered in the evaluation of the mean and standard deviation.

**The measurements were performed in a regime with a $\gamma < 0.1$, where the error in $\gamma$ is significantly increased. Data were not considered in the evaluation of the mean and standard deviation.

***Due to measurement problems, the data could not be corrected for direct excitation ($\delta$) and $\gamma$. These data were not considered in the evaluation of mean and standard deviation.

**Supplementary Table 2: FRET efficiency correction factors reported by the 7 labs for apo and holo measurements on U2AF2 labeled with Atto532-Atto643.** α is the spectral crosstalk of donor fluorescence into the acceptor channel, ß is the normalization of direct donor and acceptor excitation fluxes, *γ* is the differences in donor and acceptor quantum yields and detection efficiencies and δ is the ratio of indirect and direct acceptor excitation.

| **Lab#** | ***α*** | ***ß*** | ***γ*** | ***δ*** |
|---|---|---|---|---|
| 1 | 0.02 | 0.78 | 0.59 | 0.06 |
| 2 | 0.06 | - | 1.1 | 0.23 |
| 3 | 0.04 | 0.64 | 0.80 | 0.05 |
| 4 | 0.03 | 1.05 | 0.83 | 0.09 |
| 8 | 0.03 | 0.91 | 0.73 | 0.05 |
| 11 | 0.05 | - | 0.64 | 0.02 |
| 14 | 0.05 | - | 0.64 | 0.09 |

**Supplementary Table 3: Reported mean FRET Efficiency and standard deviation for MalE samples by the participating laboratories:** The mean FRET efficiency values $\langle E \rangle$ and distribution widths $\sigma_E$ provided by the participating labs for all the three studied mutants of MalE labeled with Alexa546 and Alexa647 under both apo and holo conditions are listed. The difference in the reported mean FRET efficiency for individual laboratories between the apo and holo state was calculated as $\langle E_{\text{holo}} \rangle - \langle E_{\text{apo}} \rangle$. The calculated mean and standard deviation of the reported FRET efficiencies over all labs are given by $\mu_{\langle E \rangle}$ and $\sigma_{\langle E \rangle}$ and for the FRET efficiency difference, $\langle E_{\text{holo}} \rangle - \langle E_{\text{apo}} \rangle$, by $\boldsymbol{\mu}_{\langle E_{\text{holo}} \rangle - \langle E_{\text{apo}} \rangle}$and $\sigma_{\langle E_{\text{holo}} \rangle - \langle E_{\text{apo}} \rangle}$, respectively.

| | **MalE-1** | | | | | **MalE-2** | | | | | **MalE-3** | | | | |
|---|---|---|---|---|---|---|---|---|---|---|---|---|---|---|---|
| **Lab#** | **Apo** | | **Holo** | | $\langle E_{\text{holo}} \rangle - \langle E_{\text{apo}} \rangle$ | **Apo** | | **Holo** | | $\langle E_{\text{holo}} \rangle - \langle E_{\text{apo}} \rangle$ | **Apo** | | **Holo** | | $\langle E_{\text{holo}} \rangle - \langle E_{\text{apo}} \rangle$ |
| | $\langle E \rangle$ | $\sigma_E$ | $\langle E \rangle$ | $\sigma_E$ | | $\langle E \rangle$ | $\sigma_E$ | $\langle E \rangle$ | $\sigma_E$ | | $\langle E \rangle$ | $\sigma_E$ | $\langle E \rangle$ | $\sigma_E$ | |
| 1 | 0.500 | 0.128 | 0.674 | 0.098 | 0.174 | 0.832 | 0.061 | 0.724 | 0.089 | -0.108 | 0.908 | 0.039 | 0.922 | 0.035 | 0.014 |
| 2 | 0.520 | | 0.700 | | 0.180 | 0.840 | | 0.730 | | -0.11 | 0.910 | | 0.910 | | 0 |
| 3 | 0.453 | 0.096 | 0.641 | 0.077 | 0.188 | 0.794 | 0.058 | 0.654 | 0.072 | -0.14 | 0.904 | 0.040 | 0.915 | 0.039 | 0.011 |
| 4 | 0.461 | 0.169 | 0.637 | 0.119 | 0.176 | 0.785 | 0.096 | 0.658 | 0.126 | -0.127 | 0.896 | 0.066 | 0.901 | 0.064 | 0.005 |
| 5 | 0.522 | | 0.703 | | 0.181 | 0.845 | | 0.724 | | -0.121 | 0.925 | | 0.912 | | -0.013 |
| 6 | 0.509 | 0.129 | 0.644 | 0.109 | 0.135 | 0.820 | 0.073 | 0.657 | 0.090 | -0.163 | 0.900 | 0.054 | 0.911 | 0.060 | 0.011 |
| 7 | 0.454 | 0.119 | 0.641 | 0.086 | 0.187 | 0.807 | 0.058 | 0.702 | 0.068 | -0.105 | 0.917 | 0.046 | 0.914 | 0.044 | -0.003 |
| 8 | 0.414 | 0.130 | 0.602 | 0.105 | 0.188 | 0.771 | 0.057 | 0.627 | 0.101 | -0.144 | 0.881 | 0.044 | 0.890 | 0.041 | 0.009 |
| 9 | 0.451 | 0.147 | 0.622 | 0.118 | 0.171 | 0.820 | 0.069 | 0.685 | 0.095 | -0.135 | 0.911 | 0.054 | 0.911 | 0.053 | 0 |
| 10 | 0.444 | 0.180 | 0.631 | 0.144 | 0.187 | 0.834 | 0.081 | 0.713 | 0.102 | -0.121 | 0.923 | 0.062 | 0.937 | 0.073 | 0.014 |
| 11 | 0.4 | 0.112 | 0.610 | 0.087 | 0.210 | 0.781 | 0.063 | 0.644 | 0.075 | -0.137 | 0.876 | 0.054 | 0.887 | 0.047 | 0.011 |
| 12 | 0.523 | 0.118 | 0.706 | 0.081 | 0.183 | 0.849 | 0.052 | 0.74 | 0.073 | -0.109 | 0.921 | 0.039 | 0.924 | 0.037 | 0.003 |
| 13 | 0.547 | 0.151 | 0.723 | 0.105 | 0.176 | 0.872 | 0.065 | 0.761 | 0.093 | -0.111 | 0.942 | 0.051 | 0.952 | 0.043 | 0.010 |
| 14 | 0.580 | | 0.745 | | 0.165 | 0.865 | | 0.765 | | -0.1 | 0.935 | | 0.970 | | 0.035 |
| 15 | 0.621 | 0.114 | 0.76 | 0.072 | 0.139 | 0.877 | 0.047 | 0.787 | 0.066 | -0.09 | 0.936 | 0.035 | 0.936 | 0.034 | 0 |
| 16 | 0.466 | 0.100 | 0.655 | 0.082 | 0.189 | 0.831 | 0.050 | 0.716 | 0.067 | -0.115 | 0923 | 0.038 | 0.93 | 0.038 | 0.007 |
| $\boldsymbol{\mu}_{\langle E \rangle}$ or $\boldsymbol{\mu}_{\langle E_{\text{holo}} \rangle - \langle E_{\text{apo}} \rangle}$ | 0.492 | | 0.667 | | 0.177 | 0.826 | | 0.705 | | -0.121 | 0.913 | | 0.920 | | 0.007 |
| $\boldsymbol{\sigma}_{\langle E \rangle}$ or $\boldsymbol{\sigma}_{\langle E_{\text{holo}} \rangle - \langle E_{\text{apo}} \rangle}$ | 0.060 | | 0.049 | | 0.019 | 0.032 | | 0.047 | | 0.019 | 0.019 | | 0.021 | | 0.010 |

**Supplementary Table 4: Reported and reanalyzed mean FRET efficiency and standard deviations for the U2AF2 experiments:** The U2AF2 protein was labeled with Atto532-Atto643 and labs provided the number of events for FRET efficiency over 51 bins between 0 and 1. To obtain the mean FRET efficiency $\langle E \rangle$ and distribution width $\sigma_E$ values for U2AF2, the reported events were fitted with one and two Gaussian distribution functions for apo and holo conditions respectively. For reanalysis, the original datasets from the individual labs were obtained and reanalyzed according to the procedure detailed in the Supplementary Note 3. U2AF2 under holo conditions has a major fraction of holo state and a minor fraction of apo state due to incomplete saturation of the RNA ligand. Only 7 labs participated in the U2AF2 study because of its complexity. The calculated mean and standard deviation of the reported and reanalyzed FRET efficiencies over all labs for each sample are given by $\mu_{\langle E \rangle}$ and $\sigma_{\langle E \rangle}$ respectively.

| Lab # | Reported | | | | | | Reanalyzed | | | | | |
|---|---|---|---|---|---|---|---|---|---|---|---|---|
| | Apo | | +RNA | | | | Apo | | +RNA | | | |
| | | | Holo state | | Apo state | | | | Holo state | | Apo state | |
| | $\langle E \rangle$ | $\sigma_E$ | $\langle E \rangle$ | $\sigma_E$ | $\langle E \rangle$ | $\sigma_E$ | $\langle E \rangle$ | $\sigma_E$ | $\langle E \rangle$ | $\sigma_E$ | $\langle E \rangle$ | $\sigma_E$ |
| 1 | 0.748 | 0.192 | 0.436 | 0.150 | 0.756 | 0.182 | 0.736 | 0.200 | 0.405 | 0.158 | 0.769 | 0.243 |
| 2 | 0.749 | 0.202 | 0.455 | 0.168 | 0.811 | 0.174 | 0.757 | 0.207 | 0.432 | 0.157 | 0.800 | 0.208 |
| 3 | 0.708 | 0.228 | 0.506 | 0.206 | 0.785 | 0.142 | 0.738 | 0.262 | 0.449 | 0.163 | 0.767 | 0.229 |
| 4 | 0.728 | 0.205 | 0.414 | 0.197 | 0.773 | 0.152 | 0.736 | 0.211 | 0.385 | 0.158 | 0.768 | 0.179 |
| 8 | 0.795 | 0.169 | 0.521 | 0.206 | 0.857 | 0.117 | 0.742 | 0.217 | 0.414 | 0.192 | 0.814 | 0.216 |
| 11 | 0.729 | 0.204 | 0.423 | 0.133 | 0.727 | 0.231 | 0.736 | 0.196 | 0.429 | 0.112 | 0.707 | 0.269 |
| 14 | 0.718 | 0.199 | 0.430 | 0.164 | 0.743 | 0.195 | 0.749 | 0.214 | 0.448 | 0.131 | 0.755 | 0.241 |
| $\mu_{\langle E \rangle}$ | 0.74 | | 0.46 | | 0.78 | | 0.742 | | 0.423 | | 0.77 | |
| $\sigma_{\langle E \rangle}$ | 0.03 | | 0.04 | | 0.04 | | 0.008 | | 0.023 | | 0.03 | |

**Supplementary Table 5: Global fit of the polarization-resolved and magic-angle fluorescence decays from sub-ensemble data of MalE and U2AF2 samples.** The different rotational correlation times, $\rho_j$, with corresponding amplitudes, $b_j$, for the $A_{ex}|A_{em}$ channels (acceptor dye) and $D_{ex}|D_{em}$ channels (donor dye) on a sub-ensemble DA population describe different depolarization processes. The fluorescence lifetimes, $\tau_i$, and fraction of molecules with a given lifetime, $x_i$, were obtained from the magic-angle decay function, which was fitted globally with the polarization-resolved fluorescence decays, $f_{VV}(t)$ and $f_{VH}(t)$, as described in Supplementary Note 8 Eqn. 8.4 - 8.11. The fit quality was judged by the reduced Chi-squared value, $\chi_r^2$. Fitted residual anisotropies for different dye combinations are used in the calculation of relative distance uncertainties (Fig. 5e), and it is shown that they are correlated to the observed dynamic shift (Fig. 5f).

| | $A_{ex}\|A_{em}$ channels | | | | | | | | | | |
|---|---|---|---|---|---|---|---|---|---|---|---|
| **Sample** | **Dye combination** | **Condition** | $\rho_1$ [ns] | $b_1 = r_{\infty,tr}$ | $\rho_2$ [ns] | $b_2$ | $\tau_1$ [ns] | $x_1$ | $\tau_2$ [ns] | $x_2$ | $\chi_r^2$ |
| MalE-1 | Alexa546-Alexa647 | apo | >50 | 0.331 | 0.84 | 0.049 | 1.69 | 0.50 | 0.93 | 0.50 | 0.96 |
| | Alexa546-AbberiorSTAR635P | apo | 20 | 0.209 | 0.16 | 0.171 | 4.00 | 1.00 | | | 0.98 |
| | Atto532-Atto643 | apo | 20 | 0.181 | 0.27 | 0.199 | 3.95 | 1.00 | | | 0.99 |
| | Alexa546-Alexa647 | holo | 20 | 0.314 | 0.10 | 0.066 | 1.66 | 0.57 | 0.95 | 0.43 | 1.39 |
| | Alexa546-Abberior STAR635P | holo | 20 | 0.182 | 0.12 | 0.198 | 4.00 | 1.00 | | | 0.98 |
| | Atto532-Atto643 | holo | 20 | 0.212 | 0.32 | 0.168 | 3.80 | 1.00 | | | 1.15 |
| MalE-2 | Alexa546-Alexa647 | apo | 20 | 0.212 | 0.24 | 0.168 | 1.39 | 0.61 | 0.75 | 0.39 | 1.15 |
| | Atto532-Atto643 | apo | 20 | 0.160 | 0.33 | 0.220 | 3.61 | 1.00 | | | 1.08 |
| | Alexa488-Alexa647 | apo | >50 | 0.150 | 0.57 | 0.230 | 1.30 | 1.00 | | | 0.94 |
| | Alexa546-Alexa647 | holo | 20 | 0.228 | 0.25 | 0.152 | 1.48 | 0.48 | 0.91 | 0.52 | 1.10 |
| | Atto532-Atto643 | holo | 20 | 0.186 | 0.45 | 0.194 | 3.62 | 1.00 | | | 1.00 |
| | Alexa488-Alexa647 | holo | 20 | 0.150 | 0.49 | 0.230 | 1.72 | 0.21 | 1.22 | 0.79 | 1.11 |
| MalE-3 | Alexa546-Alexa647 | apo | 20 | 0.215 | 0.27 | 0.165 | 1.37 | 0.58 | 0.75 | 0.42 | 1.09 |
| | Atto532-Atto643 | apo | 20 | 0.149 | 0.31 | 0.231 | 3.78 | 1.00 | | | 1.1 |
| | Alexa488-Alexa647 | apo | 20 | 0.192 | 0.62 | 0.188 | 1.37 | 1.00 | | | 1.08 |
| | Alexa546-Alexa647 | holo | 20 | 0.201 | 0.28 | 0.179 | 1.45 | 0.49 | 0.82 | 0.51 | 1.16 |
| | Atto532-Atto643 | holo | 20 | 0.147 | 0.28 | 0.233 | 3.67 | 1.00 | | | 1.12 |
| | Alexa488-Alexa647 | holo | 20 | 0.185 | 0.44 | 0.195 | 1.33 | 1.00 | | | 1.16 |
| MalE-4 | Alexa546-Alexa647 | apo | >50 | 0.205 | 0.59 | 0.175 | 1.30 | 0.68 | 0.60 | 0.32 | 0.83 |
| | Atto532-Atto643 | apo | 20 | 0.144 | 0.42 | 0.236 | 3.84 | 1.00 | | | 1.09 |
| | Alexa488-Alexa647 | apo | 20 | 0.113 | 0.29 | 0.267 | 1.50 | 0.58 | 0.88 | 0.42 | 1.16 |
| | Alexa546-Alexa647 | holo | 20 | 0.217 | 0.32 | 0.163 | 1.67 | 0.27 | 0.91 | 0.73 | 0.9 |
| | Atto532-Atto643 | holo | 20 | 0.140 | 0.29 | 0.240 | 3.53 | 0.84 | 5.08 | 0.16 | 1.07 |
| | Alexa488-Alexa647 | holo | 20 | 0.121 | 0.30 | 0.259 | 1.84 | 0.23 | 1.12 | 0.77 | 1.16 |
| MalE-5 | Alexa546-Alexa647 | apo | >50 | 0.275 | 0.68 | 0.105 | 1.65 | 0.24 | 1.00 | 0.76 | 1.20 |
| | Atto532-Atto643 | apo | 20 | 0.198 | 0.24 | 0.182 | 3.70 | 1.00 | | | 1.04 |
| | Alexa488-Alexa647 | apo | 20 | 0.160 | 0.24 | 0.220 | 1.54 | 0.43 | 1.06 | 0.57 | 1.16 |
| | Alexa546-Alexa647 | holo | >50 | 0.296 | 1.20 | 0.084 | 1.87 | 0.16 | 1.00 | 0.84 | 1.08 |
| | Atto532-Atto643 | holo | 20 | 0.201 | 0.41 | 0.179 | 3.40 | 1.00 | | | 1.16 |
| | Alexa488-Alexa647 | holo | 20 | 0.108 | 0.34 | 0.272 | 1.68 | 0.38 | 1.00 | 0.62 | 1.11 |
| U2AF2 | Alexa546-Alexa647 | apo | 21 | 0.292 | 0.10 | 0.088 | 1.70 | 0.82 | 0.87 | 0.18 | 0.98 |
| | Atto532-Atto643 | apo | 4 | 0.128 | 0.03 | 0.252 | 4.27 | 1.00 | | | 0.99 |
| | Alexa488-Alexa647 | apo | 18 | 0.208 | 0.14 | 0.172 | 1.49 | 0.92 | 3.40 | 0.08 | 1.03 |
| | Alexa546-Alexa647 | holo | 24 | 0.230 | 0.10 | 0.150 | 1.70 | 0.72 | 1.00 | 0.28 | 1.05 |
| | Atto532-Atto643 | holo | 3.4 | 0.117 | 0.06 | 0.263 | 4.10 | 1.00 | | | 1.00 |
| | Alexa488-Alexa647 | holo | 18 | 0.204 | 0.20 | 0.176 | 1.41 | 0.92 | 2.90 | 0.08 | 1.05 |

| $D_{ex}\|D_{em}$ channels | | | | | | | | | | | | | | | |
|---|---|---|---|---|---|---|---|---|---|---|---|---|---|---|---|
| **Sample** | **Dye pair** | **Condition** | $\rho_1$ [ns] | $b_1 = r_{\infty,tr}$ | $\rho_2$ [ns] | $b_2$ | $\rho_3$ [ns] | $b_3$ | $\tau_1$ [ns] | $x_1$ | $\tau_2$ [ns] | $x_2$ | $\tau_3$ [ns] | $x_3$ | $\chi_r^2$ |
| MalE-1 | Alexa546-Alexa647 | apo | 80 | 0.340 | 0.63 | 0.040 | | | 2.73 | 0.52 | 1.12 | 0.48 | | | 1.16 |
| | Alexa546-Abberior STAR635P | apo | >50 | 0.327 | 1.60 | 0.050 | | | 2.87 | 0.49 | 1.18 | 0.51 | | | 1.03 |
| | Atto532-Atto643 | apo | 20 | 0.229 | 0.21 | 0.151 | | | 2.22 | 0.79 | 3.62 | 0.21 | | | 1.24 |
| | Alexa546-Alexa647 | holo | 46 | 0.329 | 0.16 | 0.050 | | | 1.69 | 0.48 | 0.77 | 0.41 | 3.09 | 0.11 | 1.19 |
| | Alexa546-Abberior STAR635P | holo | >50 | 0.341 | 1.61 | 0.039 | | | 2.38 | 0.38 | 0.78 | 0.62 | | | 1.19 |
| | Atto532-Atto643 | holo | 20 | 0.221 | 0.28 | 0.159 | | | 2.25 | 0.64 | 0.91 | 0.36 | | | 1.19 |
| MalE-2 | Alexa546-Alexa647 | apo | >50 | 0.252 | 4.96 | 0.128 | | | 0.95 | 0.42 | 0.30 | 0.53 | 3.06 | 0.05 | 1.2 |
| | Atto532-Atto643 | apo | 20 | 0.198 | 0.26 | 0.182 | | | 0.58 | 0.67 | 1.70 | 0.33 | | | 1.1 |
| | Alexa488-Alexa647 | apo | 20 | 0.136 | 0.19 | 0.244 | | | 0.95 | 0.54 | 2.30 | 0.46 | | | 1.1 |
| | Alexa546-Alexa647 | holo | 20 | 0.255 | 0.37 | 0.125 | | | 1.30 | 0.51 | 0.38 | 0.44 | 3.26 | 0.05 | 1.01 |
| | Atto532-Atto643 | holo | 20 | 0.207 | 0.15 | 0.173 | | | 2.02 | 0.50 | 0.77 | 0.50 | | | 1.10 |
| | Alexa488-Alexa647 | holo | 20 | 0.133 | 0.23 | 0.247 | | | 0.70 | 0.27 | 2.5 | 0.73 | | | 1.14 |
| MalE-3 | Alexa546-Alexa647 | apo | 20 | 0.255 | 0.26 | 0.125 | | | 0.33 | 0.81 | 1.04 | 0.18 | 6.30 | 0.01 | 1.03 |
| | Atto532-Atto643 | apo | 20 | 0.048 | 0.14 | 0.332 | | | 0.45 | 0.87 | 2.02 | 0.13 | | | 1.00 |
| | Alexa488-Alexa647 | apo | 20 | 0.087 | 0.10 | 0.293 | | | 0.66 | 0.69 | 2.10 | 0.31 | | | 1.14 |
| | Alexa546-Alexa647 | holo | 20 | 0.187 | 0.25 | 0.193 | | | 0.33 | 0.78 | 1.02 | 0.19 | 3.20 | 0.03 | 1.13 |
| | Atto532-Atto643 | holo | 20 | 0.044 | 0.15 | 0.336 | | | 0.45 | 0.87 | 2.24 | 0.13 | | | 1.09 |
| | Alexa488-Alexa647 | holo | 20 | 0.116 | 0.15 | 0.264 | | | 0.47 | 0.64 | 1.80 | 0.36 | | | 0.96 |
| MalE-4 | Alexa546-Alexa647 | apo | >50 | 0.344 | 0.57 | 0.036 | | | 2.60 | 0.69 | 0.80 | 0.31 | | | 1.04 |
| | Atto532-Atto643 | apo | 20 | 0.126 | 1.40 | 0.05 | 0.10 | 0.202 | 2.82 | 0.8 | 1.17 | 0.20 | | | 1.13 |
| | Alexa488-Alexa647 | apo | 20 | 0.13 | 0.15 | 0.250 | | | 3.30 | 0.87 | 1.53 | 0.13 | | | 1.18 |
| | Alexa546-Alexa647 | holo | >50 | 0.348 | 1.35 | 0.032 | | | 2.26 | 0.38 | 0.80 | 0.62 | | | 1.00 |
| | Atto532-Atto643 | holo | 20 | 0.166 | 0.10 | 0.214 | | | 1.64 | 0.57 | 0.40 | 0.23 | 3.00 | 0.2 | 1.12 |
| | Alexa488-Alexa647 | holo | 20 | 0.116 | 0.13 | 0.264 | | | 2.80 | 0.68 | 1.09 | 0.32 | | | 1.02 |
| MalE-5 | Alexa546-Alexa647 | apo | >50 | 0.335 | 0.81 | 0.04 | | | 2.35 | 0.49 | 0.80 | 0.50 | | | 1.07 |
| | Atto532-Atto643 | apo | 20 | 0.238 | 0.15 | 0.142 | | | 2.36 | 0.58 | 0.60 | 0.42 | | | 1.11 |
| | Alexa488-Alexa647 | apo | 20 | 0.143 | 0.13 | 0.236 | | | 3.00 | 0.85 | 1.11 | 0.14 | | | 1.14 |
| | Alexa546-Alexa647 | holo | >50 | 0.338 | 0.29 | 0.042 | | | 1.65 | 0.34 | 0.40 | 0.66 | | | 1.1 |
| | Atto532-Atto643 | holo | 20 | 0.233 | 0.15 | 0.147 | | | 1.70 | 0.42 | 0.50 | 0.58 | | | 1.16 |
| | Alexa488-Alexa647 | holo | 20 | 0.118 | 0.10 | 0.262 | | | 2.37 | 0.51 | 0.74 | 0.49 | | | 1.07 |
| U2AF2 | Alexa546-Alexa647 | apo | 22 | 0.276 | 0.10 | 0.104 | | | 2.50 | 0.39 | 0.60 | 0.61 | | | 1.17 |
| | Atto532-Atto643 | apo | 18 | 0.253 | 0.21 | 0.127 | | | 2.59 | 0.41 | 0.53 | 0.59 | | | 1.27 |
| | Alexa488-Alexa647 | apo | 18 | 0.200 | 0.19 | 0.180 | | | 2.95 | 0.44 | 0.67 | 0.56 | | | 1.16 |
| | Alexa546-Alexa647 | holo | 25 | 0.294 | 0.10 | 0.086 | | | 2.49 | 0.43 | 0.77 | 0.57 | | | 1.14 |
| | Atto532-Atto643 | holo | 18 | 0.25 | 0.17 | 0.130 | | | 2.48 | 0.57 | 0.65 | 0.43 | | | 1.29 |
| | Alexa488-Alexa647 | holo | 18 | 0.18 | 0.17 | 0.200 | | | 2.84 | 0.67 | 0.73 | 0.33 | | | 1.09 |

**Supplementary Table 6: The statement of the individual laboratories regarding the dynamics of MalE and U2AF2**. MalE-1: 29C/352C, MalE-2: 87C/186C; MalE-3: 134C/186C. "-" = no statement, "n/a"=not applicable due to experimental limitations (instrumentation & established evaluation routines).

| Lab# | Method | Sub-ms dynamics in MalE-1/2/3 | Sub-ms dynamics in U2AF2 (apo / holo) |
|---|---|---|---|
| 1 | BVA + E-Tau | no / no / no | yes / yes |
| 2 | E-Tau (fFCS, PDA) | yes / yes / yes | yes / yes |
| 3 | BVA + E-Tau | no / no / no | yes / yes |
| 4 | BVA + E-Tau | no / no / no | yes / yes |
| 5 | n/a | n/a | n/a |
| 6 | E-Tau (fFCS) | no / no / - | yes / yes |
| 7 | BVA | no / - / yes | n/a |
| 8 | BVA | no / no / no | yes / yes |
| 9 | BVA + E-Tau | no / no / no | n/a |
| 10 | BVA | no / no / no | n/a |
| 11 | BVA + E-Tau | no / no / no | yes / yes |
| 12 | BVA | no / no / no | n/a |
| 13 | fFCS | no / no / yes | n/a |
| 14 | BVA + E-Tau | no / no / no | n/a |
| 15 | n/a | n/a | n/a |
| 16 | BVA | no / no / no | n/a |
| 17 | n/a | n/a | n/a |
| 18 | n/a | n/a | n/a |
| 19 | n/a | n/a | n/a |

**Supplementary Table 7: The apparent dynamic shifts determined for both MalE and U2AF2 samples for the data collected from 8 labs.** MalE and U2AF2 samples were labeled with Alexa546-Alexa647 and Atto532-Atto643 dye pairs respectively. The apparent dynamic shifts for the data collected in different labs are shown below for both BVA and E-$\tau$ plots. The dynamic shift, *ds* of the peak of the population was determined graphically as explained in Supplementary Note 5. For U2AF2 in the holo state, the dynamic shift was assessed only for the low-FRET RNA-bound population. Note the negative dynamic shifts which occur due to dye artifacts.

| **Lab#** | | | **1** | **2** | **3** | **4** | **7** | **8** | **11** | **14** |
|---|---|---|---|---|---|---|---|---|---|---|
| **BVA** | **MalE-1** | **Apo** | 0.0049 | 0.0007 | 0.0128 | -0.0027 | 0.0167 | 0.0075 | 0.0042 | 0.0093 |
| | | **Holo** | 0.0083 | 0.0022 | 0.0135 | -0.001 | 0.0166 | 0.0073 | 0.0079 | 0.0098 |
| | **MalE-2** | **Apo** | 0.0083 | 0.0009 | 0.014 | 0.0023 | 0.0167 | 0.0103 | 0.0080 | 0.0105 |
| | | **Holo** | 0.0070 | 0.0025 | 0.0143 | 0.0004 | 0.021 | 0.008 | 0.0062 | 0.0097 |
| | **MalE-3** | **Apo** | 0.0127 | -0.0002 | 0.0081 | -0.0043 | 0.0166 | 0.01 | 0.0107 | 0.0293 |
| | | **Holo** | 0.0068 | -0.0036 | 0.0087 | | -0.0043 | 0.0024 | 0.0077 | 0.0028 |
| | **MalE-4** | **Apo** | 0.0022 | | -0.0031 | | | | | |
| | | **Holo** | 0.004 | | 0.0070 | | | | | |
| | **MalE-5** | **Apo** | 0.0016 | | 0.0031 | | | | | |
| | | **Holo** | 0.003 | | 0.0009 | | | | | |
| | **U2AF2** | **Apo** | 0.0283 | 0.0206 | 0.0217 | 0.026 | | 0.0261 | 0.0363 | 0.0256 |
| | | **Holo** | 0.0131 | 0.018 | 0.0259 | 0.0117 | | 0.0178 | 0.0160 | 0.0171 |
| **E-$\tau$** | **MalE-1** | **Apo** | 0.074 | 0.059 | 0.055 | 0.003 | | 0.034 | | 0.088 |
| | | **Holo** | 0.069 | 0.057 | 0.062 | 0.004 | | 0.031 | | 0.068 |
| | **MalE-2** | **Apo** | 0.003 | -0.007 | 0.018 | 0.01 | | 0.029 | | 0.042 |
| | | **Holo** | 0.027 | -0.002 | 0.025 | 0.015 | | 0.011 | | 0.05 |
| | **MalE-3** | **Apo** | 0.031 | -0.013 | 0.026 | 0.004 | | 0.044 | | 0.043 |
| | | **Holo** | 0.019 | -0.019 | 0.037 | 0.035 | | 0.027 | | 0.046 |
| | **MalE-4** | **Apo** | 0.025 | 0.014 | 0.02 | | | | | |
| | | **Holo** | 0.055 | 0.03 | 0.042 | | | | | |
| | **MalE-5** | **Apo** | 0.042 | 0.027 | 0.03 | | | | | |
| | | **Holo** | 0.036 | 0.018 | 0.026 | | | | | |
| | **U2AF2** | **Apo** | 0.167 | 0.134 | 0.097 | 0.122 | | 0.141 | | 0.159 |
| | | **Holo** | 0.022 | 0.008 | 0.019 | 0.035 | | 0.012 | | 0.029 |

**Supplementary Table 8: The apparent dynamic shift values for different dye combinations of MalE and U2AF2 FRET variants as determined by three labs.** The estimated distance fluctuation $\delta R$ and FRET-averaged distance $R_{\langle E \rangle}$ is given for the measurements that passed the filtering procedure based on the estimated distance uncertainty from the residual anisotropies. Furthermore, the *ds* value for DNA rulers for each construct (LF: low-FRET, MF: medium-FRET, HF: high-FRET) and dye labels are provided and well as the average value over all measurement with the given error being the standard error of the mean, SEM.

| | | | ***Lab#1*** | | | ***Lab#2*** | | | ***Lab#3*** | | |
|---|---|---|---|---|---|---|---|---|---|---|---|
| **Sample** | **Dye combination** | **state** | ***ds*** | $R_{\langle E \rangle}$ **[Å]** | $\delta R$ **[Å]** | ***ds*** | $R_{\langle E \rangle}$ **[Å]** | $\delta R$ **[Å]** | ***ds*** | $R_{\langle E \rangle}$ **[Å]** | $\delta R$ **[Å]** |
| MalE-1 | Alexa546-Alexa647 | apo | 0.074 | | | 0.059 | | | 0.055 | | |
| MalE-1 | Alexa546-Abb. STAR635P | apo | 0.016 | | | 0.033 | | | | | |
| MalE-1 | Atto532-Atto643 | apo | 0.021 | 65.5 | 6.6 | | | | | | |
| MalE-1 | Alexa546-Alexa647 | holo | 0.069 | | | 0.057 | | | 0.062 | | |
| MalE-1 | Alexa546-Abb. STAR635P | holo | 0.003 | | | 0.035 | | | | | |
| MalE-1 | Atto532-Atto643 | holo | 0.020 | 57.6 | 4.5 | | | | | | |
| MalE-2 | Alexa546-Alexa647 | apo | 0.003 | 49.8 | 1.6 | -0.007 | 49.3 | 0.2 | 0.018 | 51.2 | 3.9 |
| MalE-2 | Atto532-Atto643 | apo | 0.007 | 47.3 | 2.2 | 0.009 | 47.3 | 2.5 | | | |
| MalE-2 | Alexa488-Alexa647 | apo | 0.001 | 47.9 | 0.8 | -0.002 | 47.9 | 0.2 | | | |
| MalE-2 | Alexa546-Alexa647 | holo | 0.027 | 55.1 | 4.8 | -0.002 | 55.3 | 0.0 | 0.025 | 56.3 | 4.7 |
| MalE-2 | Atto532-Atto643 | holo | 0.020 | 54.4 | 4.0 | 0.018 | 54.4 | 3.8 | | | |
| MalE-2 | Alexa488-Alexa647 | holo | 0.010 | 54.1 | 3.3 | 0.009 | 54.1 | 3.1 | | | |
| MalE-3 | Alexa546-Alexa647 | apo | 0.031 | 44.2 | 5.7 | -0.013 | 44.4 | 0.0 | 0.026 | 43.6 | 5.3 |
| MalE-3 | Atto532-Atto643 | apo | 0.003 | 39.3 | 1.7 | 0.002 | 39.3 | 1.4 | | | |
| MalE-3 | Alexa488-Alexa647 | apo | -0.037 | 40.4 | 0.0 | 0.013 | 40.4 | 2.6 | | | |
| MalE-3 | Alexa546-Alexa647 | holo | 0.019 | 44.2 | 4.5 | -0.019 | 43.1 | 0.0 | 0.037 | 43.8 | 6.3 |
| MalE-3 | Atto532-Atto643 | holo | -0.009 | 40.1 | 0.0 | -0.007 | 40.1 | 0.0 | | | |
| MalE-3 | Alexa488-Alexa647 | holo | -0.013 | 40.4 | 0.0 | 0.002 | 40.4 | 1.1 | | | |
| MalE-4 | Alexa546-Alexa647 | apo | 0.025 | | | 0.014 | | | 0.02 | | |
| MalE-4 | Atto532-Atto643 | apo | -0.005 | 66.9 | 0.0 | | | | 0.025 | 66.9 | 7.8 |
| MalE-4 | Alexa488-Alexa647 | apo | 0.001 | 66.2 | 2.0 | 0.024 | 66.2 | 10.8 | | | |
| MalE-4 | Alexa546-Alexa647 | holo | 0.055 | | | 0.03 | | | 0.042 | | |
| MalE-4 | Atto532-Atto643 | holo | 0.007 | 57.7 | 2.7 | | | | 0.045 | 57.7 | 6.8 |
| MalE-4 | Alexa488-Alexa647 | holo | -0.001 | 54.9 | 0.0 | 0.026 | 54.9 | 5.5 | | | |
| MalE-5 | Alexa546-Alexa647 | apo | 0.042 | | | 0.027 | | | 0.03 | | |
| MalE-5 | Atto532-Atto643 | apo | 0.014 | 60.6 | 4.2 | 0.038 | 59.4 | 6.6 | 0.031 | 59.4 | 6.0 |
| MalE-5 | Alexa488-Alexa647 | apo | 0.032 | 60.4 | 8.4 | 0.022 | 60.4 | 7.1 | | | |
| MalE-5 | Alexa546-Alexa647 | holo | 0.036 | | | 0.018 | | | 0.026 | | |
| MalE-5 | Atto532-Atto643 | holo | 0.032 | 49.1 | 4.7 | 0.027 | 49.2 | 4.3 | 0.04 | 49.2 | 5.3 |
| MalE-5 | Alexa488-Alexa647 | holo | 0.017 | 49.3 | 3.4 | 0.033 | 49.3 | 4.8 | | | |
| U2AF | Alexa546-Alexa647 | apo | 0.166 | | | 0.129 | | | | | |
| U2AF | Atto532-Atto643 | apo | 0.167 | | | 0.164 | | | | | |
| U2AF | Alexa488-Alexa647 | apo | 0.168 | | | 0.128 | | | | | |
| U2AF | Alexa546-Alexa647 | holo | 0.033 | | | 0.024 | | | | | |
| U2AF | Atto532-Atto643 | holo | 0.022 | | | 0.025 | | | | | |
| U2AF | Alexa488-Alexa647 | holo | 0.053 | | | 0.012 | | | | | |
| DNA | Alexa488-Atto647N | LF | | | | 0.007 | 75.0 | 13.9 | | | |
| DNA | Alexa488-Cy5 | LF | | | | 0.015 | 73.6 | 13.9 | | | |
| DNA | Alexa488-Alexa647 | LF | | | | 0.012 | 72.5 | 11.9 | | | |
| DNA | Alexa488-Atto647N | MF | | | | -0.008 | 61.5 | 0.3 | | | |
| DNA | Alexa488-Cy5 | MF | | | | 0.011 | 61.2 | 5.4 | | | |
| DNA | Alexa488-Alexa647 | MF | | | | -0.003 | 62.4 | 0.0 | | | |
| DNA | Alexa488-Atto647N | HF | | | | -0.012 | 51.7 | 0.0 | | | |
| DNA | Alexa488-Cy5 | HF | | | | 0.017 | 51.5 | 3.7 | | | |
| DNA | Alexa488-Alexa647 | HF | | | | -0.016 | 52.4 | 0.0 | | | |
| DNA | average over different dye combinations | - | | | | 0.0026 ± 0.0044 | | | | | |

**Supplementary Table 9: The expected FRET efficiencies and dynamic shifts for the different experimental systems based on structural models from the PDB for the dye pair Alexa546-Alexa647.** FRET efficiencies were predicted by AV simulations using the parameters given in Supplementary Table 10 based on the following PDB IDs: MalE apo - 1OMP, MalE holo - 1ANF, U2AF2 apo - 2YHO, U2AF2 holo - 2YH1. The expected dynamic shifts were calculated as described in Supplementary Note 6 using Eq. 6.1 for the $E$-$\tau$ plot and Eq. 6.23 for BVA.

| | FRET efficiency, *E* | | Expected dynamic shift, ds | |
|---|---|---|---|---|
| **Sample** | **apo** | **holo** | **$E$-$\tau$ plot** | **BVA** |
| MalE-1 | 0.358 | 0.582 | 0.0169 | 0.0215 |
| MalE-2 | 0.827 | 0.695 | 0.0131 | 0.0089 |
| MalE-3 | 0.947 | 0.941 | 0.0002 | 0.0000 |
| MalE-4 | 0.443 | 0.680 | 0.0231 | 0.0240 |
| MalE-5 | 0.578 | 0.826 | 0.0382 | 0.0282 |
| U2AF2 | 0.882 | 0.372 | 0.1426 | 0.0984 |

**Supplementary Table 10: Parameters used for the AV and ACV calculations.** The fraction of trapped dye is computed from the fundamental and residual anisotropy as $x_{\text{trapped}} = r_{\infty,tr}/r_0$ (see Supplementary Table 5 and Supplementary Note 8, Eqn. 8.8).

| | **Dye species** | |
|---|---|---|
| | **Alexa546** | **Alexa647** |
| **Dye parameters** | | |
| $L_{\text{length}}$ / Å | 20.5 | 21.0 |
| $L_{\text{width}}$ / Å | 4.5 | 4.5 |
| $R_{\text{dye},1}$ / Å | 5.0 | 11.0 |
| $R_{\text{dye},2}$ / Å | 4.5 | 4.7 |
| $R_{\text{dye},3}$ / Å | 1.5 | 1.5 |
| **AV parameters** | | |
| Grid step / Å | 0.9 | |
| Allowed sphere radius / Å | 1.0 | |
| **ACV parameters** | | |
| Contact volume thickness / Å | 3.0 | |
| **Mutant** | **Fraction of trapped dye (apo/holo)** | |
| 29 | 0.87/0.83 | 0.33/0.36 |
| 352 | 0.72/0.73 | 0.68/0.74 |
| 87 | 0.54/0.62 | 0.44/0.44 |
| 186 | 0.38/0.42 | 0.22/0.22 |
| 134 | 0.90/0.90 | 0.45/0.43 |
| 34-205* | 0.91/0.92 | 0.54/0.57 |
| 36-205* | 0.88/0.89 | 0.72/0.78 |

*Anisotropy measurements of single mutants were not performed. Hence, trapped fractions were obtained from smFRET measurements and, as such, are an average of the two labeling positions due to the stochastic labeling of the protein.

**Supplementary Table 11: Global fit of polarization-resolved fluorescence decays from ensemble measurements of single mutants of MalE.** Rotational correlation times, $\rho_j$, with corresponding amplitudes, $b_j$, for Alexa647 and Alexa546 dyes from ensemble TCSPC measurements of single cysteine MalE mutants are reported (Supplementary Note 8, Eqn. 8.8). The rotational correlation times and fluorescence lifetimes as well their corresponding amplitudes were obtained from a global fit of the polarization-resolved fluorescence decays, $f_{VV}(t)$ and $f_{VH}(t)$, as described in Supplementary Note 8. The fit quality was judged by $\chi_r^2$. The fit results indicate strong sticking interactions for positions K29C, A134C and S352C for the Alexa546 dye while, for the Alexa647 fluorophore, pronounced sticking interactions were found only at position S352C. From the fluorescence lifetime analysis, it can be seen that positions D87C and S352C are prone to quenching for the donor dye. Fitted polarization-resolved decays are combined into anisotropy decay and displayed in Fig. 5b for two representative mutation sites, namely S352C and K29C. Furthermore, obtained residual anisotropies are used in computation of Accessible Contact Volumes (Fig. 5c), which improved the agreement between modelled and measured distances (Fig. 5d).

| Dye | Mutant | Condition | $\rho_1$ [ns] | $b_1 = r_{\infty,tr}$ | $\rho_2$ [ns] | $b_2$ | $\rho_3$ [ns] | $b_3$ | $\tau_1$ [ns] | $x_1$ | $\tau_2$ [ns] | $x_2$ | $\chi_r^2$ |
|---|---|---|---|---|---|---|---|---|---|---|---|---|---|
| Alexa546 | K29C | apo | 26 | 0.332 | 1.193 | 0.048 | | | 4.03 | 1.00 | | | 1.01 |
| | D87C | apo | 20 | 0.206 | 3.500 | 0.087 | 0.572 | 0.087 | 3.85 | 1.00 | | | 1.03 |
| | A134C | apo | 20 | 0.342 | | | 0.138 | 0.038 | 3.93 | 1.00 | | | 0.99 |
| | A186C | apo | 20 | 0.146 | 3.187 | 0.089 | 0.580 | 0.145 | 3.94 | 1.00 | | | 1.07 |
| | S352C | apo | 20 | 0.272 | 2.442 | 0.053 | 0.594 | 0.055 | 3.98 | 0.90 | 1.8 | 0.10 | 1.05 |
| | K29C | holo | 20 | 0.316 | | | 0.413 | 0.064 | 4.05 | 1.00 | | | 0.90 |
| | D87C | holo | 20 | 0.236 | 1.198 | 0.144 | | | 3.91 | 1.00 | | | 0.95 |
| | A134C | holo | 20 | 0.343 | | | 0.477 | 0.037 | 3.99 | 1.00 | | | 0.94 |
| | A186C | holo | 20 | 0.161 | | | 0.973 | 0.218 | 3.93 | 1.00 | | | 0.94 |
| | S352C | holo | 20 | 0.276 | 1.100 | 0.104 | | | 4.02 | 0.90 | 1.94 | 0.10 | 0.92 |
| Alexa647 | K29C | apo | >50 | 0.125 | | | 0.639 | 0.255 | 1.878 | 0.14 | 1.165 | 0.86 | 1.19 |
| | D87C | apo | 20 | 0.169 | | | 0.748 | 0.210 | 1.729 | 0.18 | 1.174 | 0.82 | 1.07 |
| | A134C | apo | 20 | 0.170 | | | 0.539 | 0.210 | 1.833 | 0.18 | 1.205 | 0.82 | 1.15 |
| | A186C | apo | 20 | 0.082 | 1.108 | 0.189 | 0.428 | 0.109 | 1.557 | 0.17 | 1.145 | 0.83 | 1.05 |
| | S352C | apo | 30 | 0.258 | | | 0.695 | 0.122 | 1.733 | 0.42 | 1.203 | 0.58 | 1.10 |
| | K29C | holo | 20 | 0.138 | | | 0.578 | 0.241 | 1.885 | 0.13 | 1.169 | 0.87 | 1.22 |
| | D87C | holo | 20 | 0.167 | | | 0.691 | 0.213 | 1.745 | 0.18 | 1.174 | 0.82 | 1.11 |
| | A134C | holo | 20 | 0.164 | | | 0.536 | 0.216 | 1.817 | 0.18 | 1.202 | 0.82 | 1.17 |
| | A186C | holo | 20 | 0.084 | | | 0.756 | 0.296 | 1.660 | 0.09 | 1.172 | 0.91 | 1.03 |
| | S352C | holo | 20 | 0.283 | | | 0.498 | 0.097 | 1.757 | 0.42 | 1.209 | 0.58 | 1.18 |

**Supplementary Table 12: Steady-state and residual time-resolved anisotropy values of single mutant MalE samples labelled with Alexa546 and Alexa647, respectively.** In addition to time-resolved anisotropies (Supplementary Table 11), for comparison we also give the steady-state anisotropy of the same fluorophores as free dyes and coupled to double-stranded DNA. Residual anisotropies obtained from time-resolved analysis (Supplementary Note 8, Eqn. 8.4 - 8.11) of donor and acceptor dye are used in computation of Accessible Contact Volumes (ACVs) (Fig. 5c), which improved the agreement between modelled and measured distances (Fig. 5d).

| | $D_{ex}\|D_{em}$ channels (donor dye) | | $A_{ex}\|A_{em}$ channels (acceptor dye) | |
|---|---|---|---|---|
| **Sample** | **Steady-state anisotropy, $r_{ss}$** | **Residual anisotropy, $r_{\infty,tr}$** | **Steady-state anisotropy, $r_{ss}$** | **Residual anisotropy, $r_{\infty,tr}$** |
| **Free dye** | | | | |
| Alexa546 | 0.035±0.003 | 0.01±0.02 | - | |
| Alexa647 | - | | 0.120±0.007 | 0.02±0.02 |
| **DNA-Standards** | | | | |
| 8 base-pairs | 0.114±0.003 | | 0.184±0.012 | |
| 33 base-pairs | 0.134±0.002 | | 0.159±0.011 | |
| Donor-only strand | 0.134±0.003 | | - | |
| Acceptor-only strand | - | | 0.172±0.010 | |
| **Protein single mutants** | | | | |
| K29C, apo | 0.285±0.017 | 0.332 | 0.198±0.015 | 0.125 |
| K29C, holo | 0.280±0.017 | 0.316 | 0.199±0.018 | 0.138 |
| D87C, apo | 0.231±0.012 | 0.206 | 0.217±0.016 | 0.169 |
| D87C, holo | 0.225±0.005 | 0.236 | 0.229±0.017 | 0.167 |
| A134C, apo | 0.290±0.016 | 0.342 | 0.215±0.019 | 0.170 |
| A134C, holo | 0.281±0.007 | 0.343 | 0.216±0.003 | 0.164 |
| A186C, apo | 0.176±0.018 | 0.146 | 0.186±0.014 | 0.082 |
| A186C, holo | 0.161±0.010 | 0.161 | 0.186±0.016 | 0.084 |
| S352C, apo | 0.247±0.007 | 0.272 | 0.272±0.015 | 0.258 |
| S352C holo | 0.243±0.010 | 0.276 | 0.263±0.002 | 0.283 |

**Supplementary Table 13: Combined residual anisotropies of additional dye combinations for double-labeled MalE and U2AF2 samples**. Residual anisotropies of different donor and acceptor pairs used for labeling of MalE and U2AF2 variants. Residual anisotropies are computed using two approaches: from time-resolved analysis of the polarization-resolved fluorescence decays and from steady-state anisotropy measurements using a two-component Perrin equation (Supplementary Note 8, Eqn. 8.1 - 8.3). The results from the two approaches were averaged. Furthermore, using the residual anisotropies of donor and acceptor fluorophores, a combined residual anisotropy for a given FRET pair is obtained (Supplementary Note 8, Eqn. 8.12 - 8.15).

| Sample | Dye combination | state | $\langle r_{\infty,D} \rangle_{tr,ss}$ | $\langle r_{\infty,A} \rangle_{tr,ss}$ | $\langle r_{c,\infty} \rangle_{tr,ss}$ |
|---|---|---|---|---|---|
| MalE-1 | Alexa546-Alexa647 | apo | 0.350 ± 0.014 | 0.286 ± 0.064 | 0.316 ± 0.036 |
| MalE-1 | Alexa546-Abb. STAR635P | apo | 0.336 ± 0.013 | 0.228 ± 0.027 | 0.277 ± 0.017 |
| MalE-1 | Atto532-Atto643 | apo | 0.194 ± 0.049 | 0.179 ± 0.004 | 0.186 ± 0.023 |
| MalE-1 | Alexa546-Alexa647 | holo | 0.339 ± 0.014 | 0.272 ± 0.059 | 0.304 ± 0.034 |
| MalE-1 | Alexa546-Abb. STAR635P | holo | 0.336 ± 0.006 | 0.222 ± 0.056 | 0.273 ± 0.035 |
| MalE-1 | Atto532-Atto643 | holo | 0.200 ± 0.029 | 0.189 ± 0.033 | 0.195 ± 0.022 |
| MalE-2 | Alexa546-Alexa647 | apo | 0.237 ± 0.021 | 0.176 ± 0.050 | 0.204 ± 0.031 |
| MalE-2 | Atto532-Atto643 | apo | 0.154 ± 0.063 | 0.141 ± 0.027 | 0.147 ± 0.033 |
| MalE-2 | Alexa488-Alexa647 | apo | 0.110 ± 0.037 | 0.190 ± 0.056 | 0.144 ± 0.032 |
| MalE-2 | Alexa546-Alexa647 | holo | 0.257 ± 0.002 | 0.170 ± 0.082 | 0.209 ± 0.050 |
| MalE-2 | Atto532-Atto643 | holo | 0.161 ± 0.065 | 0.152 ± 0.048 | 0.156 ± 0.040 |
| MalE-2 | Alexa488-Alexa647 | holo | 0.117 ± 0.023 | 0.195 ± 0.063 | 0.151 ± 0.029 |
| MalE-3 | Alexa546-Alexa647 | apo | 0.257 ± 0.003 | 0.168 ± 0.066 | 0.208 ± 0.041 |
| MalE-3 | Atto532-Atto643 | apo | 0.035 ± 0.019 | 0.139 ± 0.014 | 0.069 ± 0.019 |
| MalE-3 | Alexa488-Alexa647 | apo | 0.090 ± 0.004 | 0.211 ± 0.026 | 0.137 ± 0.009 |
| MalE-3 | Alexa546-Alexa647 | holo | 0.212 ± 0.036 | 0.155 ± 0.065 | 0.181 ± 0.041 |
| MalE-3 | Atto532-Atto643 | holo | 0.048 ± 0.006 | 0.139 ± 0.012 | 0.082 ± 0.006 |
| MalE-3 | Alexa488-Alexa647 | holo | 0.104 ± 0.017 | 0.206 ± 0.030 | 0.147 ± 0.016 |
| MalE-4 | Alexa546-Alexa647 | apo | 0.345 ± 0.002 | 0.199 ± 0.009 | 0.262 ± 0.006 |
| MalE-4 | Atto532-Atto643 | apo | 0.140 ± 0.020 | 0.136 ± 0.011 | 0.138 ± 0.011 |
| MalE-4 | Alexa488-Alexa647 | apo | 0.116 ± 0.020 | 0.167 ± 0.077 | 0.139 ± 0.034 |
| MalE-4 | Alexa546-Alexa647 | holo | 0.352 ± 0.006 | 0.235 ± 0.025 | 0.288 ± 0.016 |
| MalE-4 | Atto532-Atto643 | holo | 0.165 ± 0.001 | 0.130 ± 0.015 | 0.146 ± 0.008 |
| MalE-4 | Alexa488-Alexa647 | holo | 0.110 ± 0.008 | 0.174 ± 0.075 | 0.138 ± 0.030 |
| MalE-5 | Alexa546-Alexa647 | apo | 0.326 ± 0.012 | 0.266 ± 0.013 | 0.295 ± 0.009 |
| MalE-5 | Atto532-Atto643 | apo | 0.232 ± 0.009 | 0.191 ± 0.009 | 0.211 ± 0.006 |
| MalE-5 | Alexa488-Alexa647 | apo | 0.135 ± 0.012 | 0.190 ± 0.042 | 0.160 ± 0.019 |
| MalE-5 | Alexa546-Alexa647 | holo | 0.330 ± 0.011 | 0.268 ± 0.040 | 0.297 ± 0.023 |
| MalE-5 | Atto532-Atto643 | holo | 0.232 ± 0.001 | 0.188 ± 0.018 | 0.209 ± 0.010 |
| MalE-5 | Alexa488-Alexa647 | holo | 0.120 ± 0.003 | 0.171 ± 0.089 | 0.143 ± 0.037 |
| U2AF | Alexa546-Alexa647 | apo | 0.305 ± 0.041 | 0.261 ± 0.044 | 0.282 ± 0.030 |
| U2AF | Atto532-Atto643 | apo | 0.229 ± 0.034 | 0.121 ± 0.010 | 0.166 ± 0.014 |
| U2AF | Alexa488-Alexa647 | apo | 0.168 ± 0.045 | 0.253 ± 0.064 | 0.207 ± 0.038 |
| U2AF | Alexa546-Alexa647 | holo | 0.332 ± 0.054 | 0.238 ± 0.012 | 0.281 ± 0.024 |
| U2AF | Atto532-Atto643 | holo | 0.241 ± 0.013 | 0.106 ± 0.016 | 0.160 ± 0.012 |
| U2AF | Alexa488-Alexa647 | holo | 0.150 ± 0.042 | 0.248 ± 0.062 | 0.193 ± 0.036 |

**Supplementary Table 14: Computed distance uncertainties for MalE and U2AF2 samples with different dye combination using a "Diffusion with traps" (DWT) model and a "Wobbling in cone" (WIC) model.** Uncertainties in the calculated distance due to uncertainities in the orientation factor, $\kappa^2$, are computed using the residual anisotropies of donor and acceptor fluorophores for a given FRET pair. For filtering out dye combinatins with specific sticking interactions, a threshold of 10% from the DWT model in the distance uncertainty was used. Details on uncertainty calculations using the DWT model[29] can be found in Supplementary Note 9. For details of the WIC model, see ref [24].

| Sample | Dye combination | state | $\Delta R_{app}(\kappa^2)$ [%] DWT model | $\Delta R_{app}(\kappa^2)$ [%] WIC model |
|---|---|---|---|---|
| MalE-1 | Alexa546-Alexa647 | apo | 14.5 | 16.1 |
| MalE-1 | Alexa546-Abb. STAR635P | apo | 11.0 | 13.3 |
| MalE-1 | Atto532-Atto643 | apo | 5.4 | 8.7 |
| MalE-1 | Alexa546-Alexa647 | holo | 14.8 | 14.9 |
| MalE-1 | Alexa546-Abb. STAR635P | holo | 11.8 | 13.1 |
| MalE-1 | Atto532-Atto643 | holo | 6.2 | 9.0 |
| MalE-2 | Alexa546-Alexa647 | apo | 9.1 | 9.6 |
| MalE-2 | Atto532-Atto643 | apo | 5.2 | 7.2 |
| MalE-2 | Alexa488-Alexa647 | apo | 4.6 | 7.4 |
| MalE-2 | Alexa546-Alexa647 | holo | 8.2 | 9.6 |
| MalE-2 | Atto532-Atto643 | holo | 4.8 | 7.6 |
| MalE-2 | Alexa488-Alexa647 | holo | 4.5 | 7.5 |
| MalE-3 | Alexa546-Alexa647 | apo | 11.4 | 9.6 |
| MalE-3 | Atto532-Atto643 | apo | 3.3 | 5.2 |
| MalE-3 | Alexa488-Alexa647 | apo | 5.5 | 7.3 |
| MalE-3 | Alexa546-Alexa647 | holo | 9.4 | 8.5 |
| MalE-3 | Atto532-Atto643 | holo | 3.6 | 5.3 |
| MalE-3 | Alexa488-Alexa647 | holo | 5.8 | 7.4 |
| MalE-4 | Alexa546-Alexa647 | apo | 9.9 | 12.5 |
| MalE-4 | Atto532-Atto643 | apo | 3.7 | 7.0 |
| MalE-4 | Alexa488-Alexa647 | apo | 3.9 | 7.2 |
| MalE-4 | Alexa546-Alexa647 | holo | 13.3 | 14.1 |
| MalE-4 | Atto532-Atto643 | holo | 4.2 | 7.4 |
| MalE-4 | Alexa488-Alexa647 | holo | 4.0 | 7.0 |
| MalE-5 | Alexa546-Alexa647 | apo | 12.7 | 14.3 |
| MalE-5 | Atto532-Atto643 | apo | 7.1 | 9.7 |
| MalE-5 | Alexa488-Alexa647 | apo | 4.5 | 7.8 |
| MalE-5 | Alexa546-Alexa647 | holo | 16.2 | 14.4 |
| MalE-5 | Atto532-Atto643 | holo | 8.5 | 9.6 |
| MalE-5 | Alexa488-Alexa647 | holo | 4.3 | 7.2 |
| U2AF | Alexa546-Alexa647 | apo | 14.3 | 13.4 |
| U2AF | Atto532-Atto643 | apo | 5.9 | 8.2 |
| U2AF | Alexa488-Alexa647 | apo | 8.1 | 9.6 |
| U2AF | Alexa546-Alexa647 | holo | 11.7 | 13.5 |
| U2AF | Atto532-Atto643 | holo | 5.3 | 8.1 |
| U2AF | Alexa488-Alexa647 | holo | 5.9 | 9.1 |

**Supplementary Table 15: Analysis of the U2AF2 dynamics. Results collected from 5 participating labs on the dynamics of U2AF2 labelled with the Atto532-Atto643 dye-pair.** Timescales and rates were derived by dynamic PDA, filtered-FCS, and FRET-FCS as reported by the various groups. $R_1$ and $R_2$ are the distances corresponding to the compact and open states respectively for extracting the interconversion rates by dynamic PDA. Interconversion rates from closed to open and open to closed are denoted as $k_{12}$ and $k_{21}$ with their respective relaxation time $\tau_R$. See Supplementary Note 17 for more details on the dynamics of U2AF2. The two extracted relaxation times from FRET-FCS and fFCS are denoted as $t_{R1}$ and $t_{R2}$. See Supplementary Note 16 for more details on the filtered-FCS analysis. Average values of the rates and relaxation times, are given in the last row. * Only Lab#2 performed the FRET-FCS analysis and is not considered for average values for filtered-FCS.

| **Lab#** | **Method** | | | **Apo** | | | | | **Holo** | | | | |
|---|---|---|---|---|---|---|---|---|---|---|---|---|---|
| | | $R_1$ (Å) | $R_2$ (Å) | $t_{R1}$ (µs) | $t_{R2}$ (µs) | $k_{12}$ ($ms^{-1}$) | $k_{21}$ ($ms^{-1}$) | $\tau_R$ (ms) | $t_{R1}$ (µs) | $t_{R2}$ (µs) | $k_{12}$ ($ms^{-1}$) | $k_{21}$ ($ms^{-1}$) | $\tau_R$ (ms) |
| 1 | Filtered-FCS | | | 200 | 10 | | | | | | | | |
| | Dynamic PDA | 38 | 59 | | | 0.43 | 0.07 | 2 | | | 0.52 | 0.14 | 1.42 |
| 2 | Filtered-FCS | | | 320 | 6 | | | | 320 | 6 | | | |
| | FRET-FCS* | | | 32 | 2 | | | | 321 | | | | |
| | Dynamic PDA | 37 | 60 | | | 2.3 | 0.39 | 0.37 | | | 2.05 | 1.29 | 0.29 |
| 8 | Dynamic PDA | 38 | 56 | | | | | | | | 0.62 | 0.18 | 1.25 |
| 11 | Dynamic PDA | 38 | 56 | | | | | | | | 0.63 | 0.83 | 0.68 |
| 14 | Filtered-FCS | | | 370 | 12 | | | | | | | | |
| | Dynamic PDA | 37 | 60 | | | 2.3 | 4.6 | 0.14 | 617 | 15 | 0.47 | 0.15 | 1.6 |
| Average | | | | 296 | 9.3 | 1.67 | 1.68 | 0.83 | | | 0.85 | 0.51 | 1.05 |

**Supplementary Table 16: The global dynamic photon distribution analysis (PDA) of apo and holo U2AF2 labeled with the Atto532-Atto643 dye pair.** Parameters of the global dynamic PDA model are given, which is composed of a two-state dynamic system (apo ensemble) and two static states (holo and low-FRET) as described in the main text and Supplementary Note 17. The fit was performed globally over the apo and holo measurements using time windows of 0.5, 1.0, 1.5 and 2.0 ms, respectively. See Supplementary Figure 16 for an overview of the fits. The global reduced chi-square of the fit was 1.69. The analysis was performed using correction factors of: $\gamma = 0.59$ for the detection correction factor, a direct excitation correction of $\delta = 0.024$, a crosstalk value of $\alpha = 0.02$ and a Förster radius of 59 Å. Background count rates were 0.82 kHz and 0.28 kHz in the donor and FRET detection channels respectively. Fitting was performed using the *PDAFit* module of the PAM software package[7] .Errors were approximated from the covariance matrix given from the fit routine.

| Population | | Parameter | Sample apo | Sample holo |
|---|---|---|---|---|
| **apo ensemble** | compact state | $R_{compact}$ [Å] | 37.4 ± 0.1 | |
| | | $\sigma_{compact}$ [Å] | 2.1 ± 0.4 | |
| | detached ensemble | $R_{detacted}$ [Å] | 50.0 ± 0.1 | |
| | | $\sigma_{detached}$ [Å] | 4.5 ± 0.2 | |
| | kinetic rates | $k_{c\text{->}d}$ [ms$^{-1}$] | 0.14 ± 0.11 | |
| | | $k_{d\text{->}c}$ [ms$^{-1}$] | 0.013 ± 0.011 | |
| | amplitude | $A_{apo}$ | 0.95 ± 0.02 | 0.35 ± 0.05 |
| **holo** | open conformation | $R_{open}$[Å] | 60.8 ± 0.1 | |
| | | $\sigma_{open}$ [Å] | 2.9 ± 0.2 | |
| | amplitude | $A_{holo}$ | 0.03 ± 0.01 | 0.60 ± 0.03 |
| **low-FRET** | apparent distance | $R_{LF}$[Å] | 73.9 ± 4 | |
| | | $\sigma_{LF}$ [Å] | 7.2 ± 1.4 | |
| | amplitude | $A_{LF}$ | 0.02 ± 0.01 | 0.05 ± 0.03 |

**Supplementary Table 17: Correction factors obtained after reanalysis of the U2AF2 datasets from 7 different labs.** Raw datasets were collected from 8 different labs for U2AF2 labeled with Atto532-Atto643. The reanalysis procedure is described in the Supplementary Note 3.

| Lab# | $\alpha$ | $\delta$ | $\gamma$ | $\beta$ |
|---|---|---|---|---|
| 1 | 0.02 | 0.06 | 0.73 | 0.72 |
| 2 | 0.04 | 0.20 | 1.26 | 0.22 |
| 3 | 0.03 | 0.06 | 0.98 | 0.77 |
| 4 * | 0.03 | 0.08 | 0.98 | 1.15 |
| 8 | 0.03 | 0.05 | 1.15 | 0.80 |
| 11 | 0.03 | 0.07 | 0.65 | 1.21 |
| 14 | 0.03 | 0.05 | 0.92 | 1.00 |

* In these measurements, we observed a large contribution of the subpopulation with quenched acceptor lifetime (holo-high FRET) for double-labeled molecules. Hence, one population from apo and only one population from holo (holo-low FRET) conformation was used to calculate the $\gamma$ globally.

**Supplementary Table 18: Dynamic photon distribution analysis (PDA) of apo and holo U2AF2 labeled with Alexa546-Alexa647 dye-pair.** Parameters of the global PDA model are given, which is composed of a two-state dynamic system (apo ensemble) and two static states (holo and low-FRET) as described in the Supplementary Note 17. The fit was performed globally over the apo and holo measurements using time windows of 0.5, 1.0, and 1.5 ms respectively. See Supplementary Figure 19 for an overview of the fits. The analysis was performed using a detection correction factor of $\gamma = 0.32$, a direct excitation correction of $\delta = 0.10$, a crosstalk value of $\alpha = 0.036$ and a Förster radius of 65 Å. Background count rates were 0.918 kHz and 0.281 kHz in the donor and FRET detection channels respectively. Fitting was performed using the *PDAFit* module of the PAM software package[7].

<table>
<tr><th colspan="2" rowspan="2">Population</th><th rowspan="2">Parameter</th><th colspan="2">Sample</th></tr>
<tr><th>apo</th><th>holo</th></tr>
<tr><td rowspan="7">apo ensemble</td><td rowspan="2">compact state</td><td>$R_{compact}$ [Å]</td><td colspan="2">35.4</td></tr>
<tr><td>$\sigma_{compact}$ [Å]</td><td colspan="2">3.5</td></tr>
<tr><td rowspan="2">detached ensemble</td><td>$R_{detacted}$ [Å]</td><td colspan="2">50.3</td></tr>
<tr><td>$\sigma_{detached}$ [Å]</td><td colspan="2">5.0</td></tr>
<tr><td rowspan="2">kinetic rates</td><td>$k_{c\text{->}d}$ [$ms^{-1}$]</td><td colspan="2">0.16</td></tr>
<tr><td>$k_{d\text{->}c}$ [$ms^{-1}$]</td><td colspan="2">0.014</td></tr>
<tr><td>amplitude</td><td>$A_{apo}$</td><td>0.97</td><td>0.36</td></tr>
<tr><td rowspan="3">holo</td><td rowspan="2">open conformation</td><td>$R_{open}$[Å]</td><td>-</td><td>62.0</td></tr>
<tr><td>$\sigma_{open}$ [Å]</td><td>-</td><td>2.9</td></tr>
<tr><td>amplitude</td><td>$A_{holo}$</td><td>-</td><td>0.59</td></tr>
<tr><td rowspan="3">low-FRET</td><td rowspan="2">apparent distance</td><td>$R_{LF}$[Å]</td><td colspan="2">72.5</td></tr>
<tr><td>$\sigma_{LF}$ [Å]</td><td colspan="2">5.0</td></tr>
<tr><td>amplitude</td><td>$A_{LF}$</td><td>0.03</td><td>0.05</td></tr>
</table>

**Supplementary Table 19: Detailed information on dye maleimides.** Dye-maleimide conjugates were purchased from the indicated companies. Structural formulas of dyes are illustrated below, with the exception of Atto643 where the chemical structure of the dye is unknown. For dyes Atto532 and Abberior STAR635P, linkage length to the maleimide functional group is not indicated by the producer. Those dyes are therefore illustrated without the maleimide modification.

| Fluorophore | Company | Catalog Number |
|---|---|---|
| Alexa488 C5 Maleimide | ThermoFisher Scientific | A10254 |
| Alexa546 C5 Maleimide | ThermoFisher Scientific | A10258 |
| Alexa647 C2 Maleimide | ThermoFisher Scientific | A20347 |
| Atto532 Maleimide | ATTO-TEC | AD 532-45 |
| Atto643 Maleimide | ATTO-TEC | AD 643 |
| Abberior STAR635P Maleimide | Abberior | ST635P |

Alexa 488 C5 Maleimide

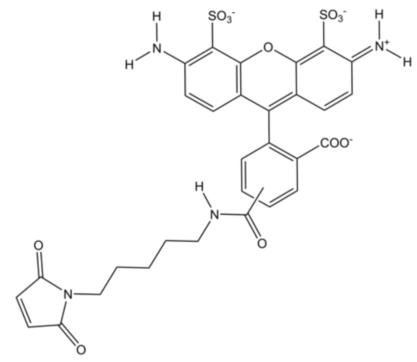

Alexa 546 C5 Maleimide

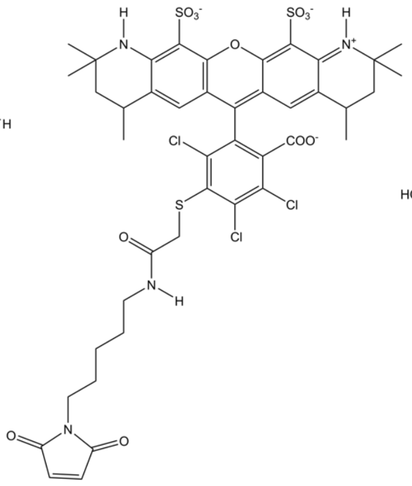

Alexa 647 C2 Maleimide

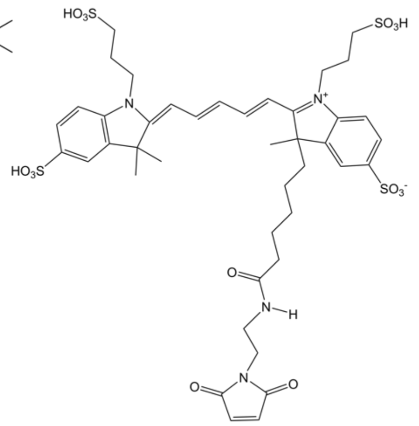

Atto 532

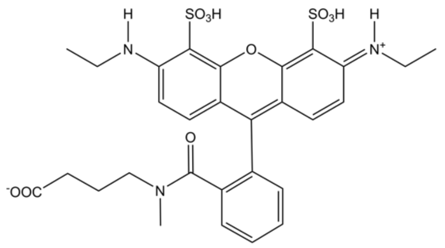

Abberior STAR635P

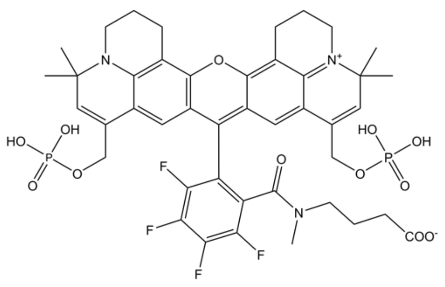

**Supplementary References:**

1. Peulen, T. O., Opanasyuk, O. & Seidel, C. A. M. Combining Graphical and Analytical Methods with Molecular Simulations to Analyze Time-Resolved FRET Measurements of Labeled Macromolecules Accurately. *J. Phys. Chem. B* **121**, 8211–8241 (2017).
2. Haas, E., Ephraim-Katchalski-Katzir & Steinberg, I. Z. Effect of the Orientation of Donor and Acceptor on the Probability of Energy Transfer Involving Electronic Transitions of Mixed Polarization. *Biochemistry* **17**, 5064–5070 (1978).
3. Kong, X., Nir, E., Hamadani, K. & Weiss, S. Photobleaching pathways in single-molecule FRET experiments. *J. Am. Chem. Soc.* **129**, 4643–4654 (2007).
4. Kudryavtsev, V. *et al.* Combining MFD and PIE for accurate single-pair Förster resonance energy transfer measurements. *ChemPhysChem* **13**, 1060–1078 (2012).
5. Eggeling, C. *et al.* Data registration and selective single-molecule analysis using multi-parameter fluorescence detection. *J. Biotechnol.* **86**, 163–180 (2001).
6. Lerner, E. *et al.* FRET-based dynamic structural biology: Challenges, perspectives and an appeal for open-science practices. *eLife* **10**, e60416 (2021).
7. Schrimpf, W., Barth, A., Hendrix, J. & Lamb, D. C. PAM: A Framework for Integrated Analysis of Imaging, Single-Molecule, and Ensemble Fluorescence Data. *Biophys. J.* **114**, 1518–1528 (2018).
8. Tomov, T. E. *et al.* Disentangling subpopulations in single-molecule FRET and ALEX experiments with photon distribution analysis. *Biophys. J.* **102**, 1163–1173 (2012).
9. Lee, N. K. *et al.* Accurate FRET measurements within single diffusing biomolecules using alternating-laser excitation. *Biophys. J.* **88**, 2939–2953 (2005).
10. Hellenkamp, B. *et al.* Precision and accuracy of single-molecule FRET measurements—a multi-laboratory benchmark study. *Nat. Methods* **15**, 669–676 (2018).
11. Ingargiola, A., Lerner, E., Chung, S. Y., Weiss, S. & Michalet, X. FRETBursts: An open source toolkit for analysis of freely-diffusing Single-molecule FRET. *PLoS One* **11**, 39198 (2016).
12. Fries, J. R., Brand, L., Eggeling, C., Köllner, M. & Seidel, C. A. M. Quantitative identification of different single molecules by selective time-resolved confocal fluorescence spectroscopy. *J. Phys. Chem. A* **102**, 6601–6613 (1998).
13. Torella, J. P., Holden, S. J., Santoso, Y., Hohlbein, J. & Kapanidis, A. N. Identifying molecular dynamics in single-molecule fret experiments with burst variance analysis. *Biophys. J.* **100**, 1568–1577 (2011).
14. Kalinin, S., Valeri, A., Antonik, M., Felekyan, S. & Seidel, C. A. M. Detection of Structural Dynamics by FRET: A Photon Distribution and Fluorescence Lifetime Analysis of Systems with Multiple States. *J. Phys. Chem. B* **114**, 7983–7995 (2010).
15. Barth, A. *et al.* Unraveling multi-state molecular dynamics in single-molecule FRET experiments. I. Theory of FRET-lines. *J. Chem. Phys.* **156**, 141501 (2022).
16. Foerster, T. Zwischenmolekulare Energiewanderung und Fluoreszenz. *Ann. Phys.* **437**, 55–75 (1948).

17. ur Rehman Malik, N. & Sadarangani, C. Brushless doubly-fed induction machine with rotating power electronic converter for wind power applications. *2011 Int. Conf. Electr. Mach. Syst.* 1–6 (2011).

18. Wiederschain, G. Y. The Molecular Probes handbook. A guide to fluorescent probes and labeling technologies. *Biochem.* **76**, 1276–1276 (2011).

19. Voith von Voithenberg, L. & Lamb, D. C. Single Pair Förster Resonance Energy Transfer: A Versatile Tool To Investigate Protein Conformational Dynamics. *BioEssays* **40**, 1–14 (2018).

20. Spectra, E. Alexa Fluor ® Dyes — Across the Spectrum. 1–3 (2014).

21. Schaffer, J. *et al.* Identification of Single Molecules in Aqueous Solution by Time-Resolved Fluorescence Anisotropy. *J. Phys. Chem. A* **103**, 331–336 (1999).

22. Perrin, F. Polarisation de la lumière de fluorescence. Vie moyenne des molécules dans l'etat excité. *J. Phys. le Radium* **7**, 390–401 (1926).

23. Sisamakis, E., Valeri, A., Kalinin, S., Rothwell, P. J. & Seidel, C. A. M. Accurate Single-Molecule FRET Studies Using Multiparameter Fluorescence Detection. *Methods Enzymol.* **475**, 455–514 (2010).

24. Sindbert, S. *et al.* Accurate distance determination of nucleic acids via Förster resonance energy transfer: Implications of dye Linker length and rigidity. *J. Am. Chem. Soc.* **133**, 2463–2480 (2011).

25. Koshioka, M., Sasaki, K. & Masuhara, H. Time-Dependent Fluorescence Depolarization Analysis in Three-Dimensional Microspectroscopy. *Appl. Spectrosc.* **49**, 224–228 (1995).

26. Schröder, G. F., Alexiev, U. & Grubmüller, H. Simulation of fluorescence anisotropy experiments: Probing protein dynamics. *Biophys. J.* **89**, 3757–3770 (2005).

27. Dale, R. E. & Eisinger, J. Intramolecular distances determined by energy transfer. Dependence on orientational freedom of donor and acceptor. *Biopolymers* **13**, 1573–1605 (1974).

28. Ivanov, V., Li, M. & Mizuuchi, K. Impact of emission anisotropy on fluorescence spectroscopy and FRET distance measurements. *Biophys. J.* **97**, 922–929 (2009).

29. Kalinin, S., Fulle, S., Hanke, C.A., Peulen, T. O., Sindbert, S., Felekyan, S., Kühnemuth, R., Gohlke, H., Seidel, C. A. M. Diffusion with traps: experiment, simulation, and theory to describe the dynamics of flexibly linked fluorophores in biomolecular FRET. *Prep.*

30. Case, D. A. *et al. AMBER 2018, University of California. University of California, San Francisco* (2018).

31. Maier, J. A. *et al.* ff14SB: Improving the Accuracy of Protein Side Chain and Backbone Parameters from ff99SB. *J. Chem. Theory Comput.* **11**, 3696–3713 (2015).

32. Horn, H. W. *et al.* Development of an improved four-site water model for biomolecular simulations: TIP4P-Ew. *J. Chem. Phys.* **120**, 9665–9678 (2004).

33. Hopkins, C. W., Le Grand, S., Walker, R. C. & Roitberg, A. E. Long-time-step molecular dynamics through hydrogen mass repartitioning. *J. Chem. Theory Comput.*

**11**, 1864–1874 (2015).

34. Ryckaert, J. P., Ciccotti, G. & Berendsen, H. J. C. Numerical integration of the cartesian equations of motion of a system with constraints: molecular dynamics of n-alkanes. *J. Comput. Phys.* **23**, 327–341 (1977).

35. Darden, T., York, D. & Pedersen, L. Particle mesh Ewald: An N·log(N) method for Ewald sums in large systems. *J. Chem. Phys.* **98**, 10089–10092 (1993).

36. McGibbon, R. T. *et al.* MDTraj: A Modern Open Library for the Analysis of Molecular Dynamics Trajectories. *Biophys. J.* **109**, 1528–1532 (2015).

37. Gopich, I. V. & Szabo, A. Single-molecule FRET with diffusion and conformational dynamics. *J. Phys. Chem. B* **111**, 12925–12932 (2007).

38. Zander, C. *et al.* Detection and characterization of single molecules in aqueous solution. *Appl. Phys. B Lasers Opt.* **63**, 517–523 (1996).

39. Hall, P. & Selinger, B. *Better estimates of exponential decay parameters*. *Journal of Physical Chemistry* **85**, (1981).

40. Livesey, A. K. & Skilling, J. Maximum entropy theory. *Acta Crystallogr. Sect. A* **41**, 113–122 (1985).

41. Brochon, J. C. Maximum entropy method of data analysis in time-resolved spectroscopy. *Methods Enzymol.* **240**, 262–311 (1994).

42. Skilling, J. & Bryan, R. K. Maximum entropy image reconstruction: general algorithm. *Mon. Not. R. Astron. Soc.* **211**, 111–124 (1984).

43. Vinogradov, S. A. & Wilson, D. F. Recursive maximum entropy algorithm and its application to the luminescence lifetime distribution recovery. *Appl. Spectrosc.* **54**, 849–855 (2000).

44. Huang, J. R. *et al.* Transient electrostatic interactions dominate the conformational equilibrium sampled by multidomain splicing factor U2AF65: A combined NMR and SAXS study. *J. Am. Chem. Soc.* **136**, 7068–7076 (2014).

45. Kalinin, S. *et al.* A toolkit and benchmark study for FRET-restrained high-precision structural modeling. *Nat. Methods* **9**, 1218–1225 (2012).

46. Bowman, A. W. & Azzalini, A. Computational aspects of nonparametric smoothing with illustrations from the sm library. *Comput. Stat. Data Anal.* **42**, 545–560 (2003).

47. Felekyan, S., Kalinin, S., Sanabria, H., Valeri, A. & Seidel, C. A. M. Filtered FCS: Species auto- and cross-correlation functions highlight binding and dynamics in biomolecules. *ChemPhysChem* **13**, 1036–1053 (2012).

48. Barth, A. *et al.* Dynamic interactions of type I cohesin modules fine-tune the structure of the cellulosome of Clostridium thermocellum. *Proc. Natl. Acad. Sci. U. S. A.* **115**, E11274–E11283 (2018).

49. Von Voithenberg, L. V. *et al.* Recognition of the 3′ splice site RNA by the U2AF heterodimer involves a dynamic population shift. *Proc. Natl. Acad. Sci. U. S. A.* **113**, E7169–E7175 (2016).

50. Naudi-Fabra, S., Tengo, M., Jensen, R., Blackledge, M. & Milles, S. Quantitative

Description of Intrinsically Disordered Proteins Using Single-Molecule FRET, NMR, and SAXS. *J. Am. Chem. Soc* **143**, 50 (2021).

51. Margittai, M. *et al.* Single-molecule fluorescence resonance energy transfer reveals a dynamic equilibrium between closed and open conformations of syntaxin 1. *Proc. Natl. Acad. Sci. U. S. A.* **100**, 15516–15521 (2003).

52. Torres, T. & Levitus, M. Measuring conformational dynamics: A new FCS-FRET approach. *J. Phys. Chem. B* **111**, 7392–7400 (2007).

53. Felekyan, S., Sanabria, H., Kalinin, S., Kühnemuth, R. & Seidel, C. A. M. Analyzing Förster resonance energy transfer with fluctuation algorithms. *Methods Enzymol.* **519**, 39–85 (2013).

54. Antonik, M., Felekyan, S., Gaiduk, A. & Seidel, C. A. M. Separating structural heterogeneities from stochastic variations in fluorescence resonance energy transfer distributions via photon distribution analysis. *J. Phys. Chem. B* **110**, 6970–6978 (2006).

55. Kalinin, S., Felekyan, S., Antonik, M. & Seidel, C. A. M. Probability distribution analysis of single-molecule fluorescence anisotropy and resonance energy transfer. *J. Phys. Chem. B* **111**, 10253–10262 (2007).

56. MacKereth, C. D. *et al.* Multi-domain conformational selection underlies pre-mRNA splicing regulation by U2AF. *Nature* **475**, 408–413 (2011).

57. Sánchez-Rico, C., Voith von Voithenberg, L., Warner, L., Lamb, D. C. & Sattler, M. Effects of Fluorophore Attachment on Protein Conformation and Dynamics Studied by spFRET and NMR Spectroscopy. *Chem. - A Eur. J.* **23**, 14267–14277 (2017).

58. Barth, A., Voith Von Voithenberg, L. & Lamb, D. C. Quantitative Single-Molecule Three-Color Förster Resonance Energy Transfer by Photon Distribution Analysis. *J. Phys. Chem. B* **123**, 6901–6916 (2019).

59. Gouridis, G. *et al.* Conformational dynamics in substrate-binding domains influences transport in the ABC importer GlnPQ. *Nat. Struct. Mol. Biol.* **22**, 57–64 (2015).

60. De Boer, M. *et al.* Conformational and dynamic plasticity in substrate-binding proteins underlies selective transport in ABC importers. *eLife* **8**, e44652 (2019).

61. De Boer, M., Gouridis, G., Muthahari, Y. A. & Cordes, T. Single-Molecule Observation of Ligand Binding and Conformational Changes in FeuA. *Biophys. J.* **117**, 1642–1654 (2019).

62. Wolf, S. *et al.* Hierarchical dynamics in allostery following ATP hydrolysis monitored by single molecule FRET measurements and MD simulations. *Chem. Sci.* **12**, 3350–3359 (2021).

63. Ambrose, B. *et al.* The smfBox is an open-source platform for single-molecule FRET. *Nat. Commun.* **11**, 5641 (2020).

64. Abdelhamid, M. A. S., Rhind-Tutt, A. V., Ambrose, B. & Craggs, T. D. Making Precise and Accurate Single-Molecule FRET Measurements using the Open-Source smfBox. *J. Vis. Exp.* e62378 (2021). doi:10.3791/62378

65. Tan, P. S. & Lemke, E. A. Probing Differential Binding Mechanisms of

Phenylalanine-Glycine-Rich Nucleoporins by Single-Molecule FRET. in *Methods in Enzymology* **611**, 327–346 (2018).

66. Müller, B. K., Zaychikov, E., Bräuchle, C. & Lamb, D. C. Pulsed interleaved excitation. *Biophys. J.* **89**, 3508–3522 (2005).

67. Robb, N. C. *et al.* The transcription bubble of the RNA polymerase-promoter open complex exhibits conformational heterogeneity and millisecond-scale dynamics: Implications for transcription start-site selection. *J. Mol. Biol.* **425**, 875–885 (2013).

68. Santoso, Y. *et al.* Conformational transitions in DNA polymerase I revealed by single-molecule FRET. *Proc. Natl. Acad. Sci. U. S. A.* **107**, 715–720 (2010).

69. Santoso, Y., Torella, J. P. & Kapanidis, A. N. Characterizing single-molecule FRET dynamics with probability distribution analysis. *ChemPhysChem* **11**, 2209–2219 (2010).

70. Robb, N. C. *et al.* Single-molecule FRET reveals the pre-initiation and initiation conformations of influenza virus promoter RNA. *Nucleic Acids Res.* **44**, 10304–10315 (2016).

71. Tomescu, A. I., Robb, N. C., Hengrung, N., Fodor, E. & Kapanidis, A. N. Single-molecule FRET reveals a corkscrew RNA structure for the polymerase-bound influenza virus promoter. *Proc. Natl. Acad. Sci.* **111**, E3335-3342 (2014).

72. Kapanidis, A. N. *et al.* Fluorescence-aided molecule sorting: Analysis of structure and interactions by alternating-laser excitation of single molecules. *Proc. Natl. Acad. Sci. U. S. A.* **101**, 8936–8941 (2004).

73. Kapanidis, A. N. *et al.* Alternating-laser excitation of single molecules. *Acc. Chem. Res.* **38**, 523–533 (2005).

74. Krainer, G. *et al.* Ultrafast Protein Folding in Membrane-Mimetic Environments. *J. Mol. Biol.* **430**, 554–564 (2018).

75. Hartmann, A., Krainer, G., Keller, S. & Schlierf, M. Quantification of Millisecond Protein-Folding Dynamics in Membrane-Mimetic Environments by Single-Molecule Förster Resonance Energy Transfer Spectroscopy. *Anal. Chem.* **87**, 11224–11232 (2015).

76. Hartmann, A. Observing Biomolecular Dynamics from Nanoseconds to Hours with Single-Molecule Fluorescence Spectroscopy. 139 (2017).

77. Kramm, K. *et al.* DNA origami-based single-molecule force spectroscopy elucidates RNA Polymerase III pre-initiation complex stability. *Nat. Commun.* **11**, 2828 (2020).

78. Kahra, D. *et al.* Conformational plasticity and dynamics in the generic protein folding catalyst SlyD unraveled by single-molecule FRET. *J. Mol. Biol.* **411**, 781–790 (2011).

79. Olofsson, L. & Margeat, E. Pulsed interleaved excitation fluorescence spectroscopy with a supercontinuum source. *Opt. Express* **21**, 3370 (2013).

80. Fuertes, G. *et al.* Decoupling of size and shape fluctuations in heteropolymeric sequences reconciles discrepancies in SAXS vs. FRET measurements. *Proc. Natl. Acad. Sci. U. S. A.* **114**, E6342–E6351 (2017).

81. Harris, P. D. *et al.* Multi-parameter photon-by-photon hidden Markov modeling. *Nat.*

*Commun.* **13**, 1000 (2022).

82. Ingargiola, A., Laurence, T., Boutelle, R., Weiss, S. & Michalet, X. Photon-HDF5: An Open File Format for Timestamp-Based Single-Molecule Fluorescence Experiments. *Biophys. J.* **110**, 26–33 (2016).

83. Nir, E. *et al.* Shot-noise limited single-molecule FRET histograms: Comparison between theory and experiments. *J. Phys. Chem. B* **110**, 22103–22124 (2006).

84. Buning, R., Kropff, W., Martens, K. & van Noort, J. spFRET reveals changes in nucleosome breathing by neighboring nucleosomes. *J. Phys. Condens. Matter* **27**, 064103 (2015).

85. Zhou, R., Schlierf, M. & Ha, T. Force-Fluorescence Spectroscopy at the Single-Molecule Level. *Methods Enzymol.* **475**, 405–426 (2010).

86. Nikaido, H. Maltose transport system of Escherichia coli: An ABC-type transporter. *FEBS Lett.* **346**, 55–58 (1994).

87. McKinney, S. A., Joo, C. & Ha, T. Analysis of single-molecule FRET trajectories using hidden Markov modeling. *Biophys. J.* **91**, 1941–1951 (2006).

88. Jerabek-Willemsen, M. *et al.* MicroScale Thermophoresis: Interaction analysis and beyond. *J. Mol. Struct.* **1077**, 101–113 (2014).

89. Widengren, J. & Schwiÿle, P. Characterization of photoinduced isomerization and back-isomerization of the cyanine dye cy5 by fluorescence correlation spectroscopy. *J. Phys. Chem. A* **104**, 6416–6428 (2000).

90. Petrášek, Z. & Schwille, P. Precise measurement of diffusion coefficients using scanning fluorescence correlation spectroscopy. *Biophys. J.* **94**, 1437–1448 (2008).
